# Dynamic evolution of *de novo* DNA methyltransferases in rodent and primate genomes

**DOI:** 10.1101/786863

**Authors:** Antoine Molaro, Harmit S. Malik, Deborah Bourc’his

## Abstract

Transcriptional silencing of retrotransposons via DNA methylation is paramount for mammalian fertility and reproductive fitness. During germ cell development, most mammalian species utilize the *de novo* DNA methyltransferases DNMT3A and DNMT3B to establish DNA methylation patterns. However, many rodent species deploy a third enzyme, DNMT3C, to selectively methylate the promoters of young retrotransposon insertions in their germline. The evolutionary forces that shaped DNMT3C’s unique function are unknown. Using a phylogenomic approach, we confirm here that *Dnmt3C* arose through a single duplication of *Dnmt3B* that occurred between 45-71Mya in the last common ancestor of muroid rodents. Most importantly, we reveal that DNMT3C is composed of two independently evolving segments: the latter two-thirds has undergone recurrent gene conversion with *Dnmt3B*, whereas the N-terminus has not undergone gene conversion and has evolved under strong diversifying selection. We hypothesize that positive selection of *Dnmt3C* is the result on an ongoing evolutionary arms race with young retrotransposon lineages in muroid genomes. Interestingly, although primates lack DNMT3C, we find that the N-terminus of DNMT3A has also evolved under diversifying selection. Thus, the N-termini of two independent *de novo* DNMT enzymes have evolved under diversifying selection in rodents and primates. We hypothesize that repression of young retrotransposons might be driving the recurrent innovation of a functional domain in the N-termini on germline DNMTs in mammals.

## Introduction

The deposition of methylation on DNA is a deeply conserved process. In mammals, it plays crucial roles in genome stability, development, genomic imprinting and chromosome-wide epigenetic silencing such as X-inactivation (1). Mammalian DNMTs (DNA methyltransferases) are enzymes that catalyze the addition of a methyl group onto cytosines (2). Most mammals encode three catalytically active enzymes (DNMT1, DNMT3A, and DNMT3B) and one non-enzymatic germ cell-specific cofactor (DNMT3L) (2-5). While DNMT1 targets hemi-methylated cytosines (maintenance DNA methyltransferase) (6-8), DNMT3A and 3B are classified as *de novo* methyltransferases that target unmodified sites (9-12). In mice, constitutive genetic knockouts of *Dnmt1, Dnmt3A*, or *Dnmt3B* are lethal, whereas *Dnmt3L* mutations lead to sterility (10, 13, 14).

Phylogenetic analyses have suggested that the DNMT enzymes belong to the clade of 5-cytosine methyltransferases, which likely predated the origin of eukaryotes (5, 15). Although both *Dnmt1* and *Dnmt3A* were present in the common ancestor of all metazoans, *Dnmt3B* is believed to have arisen by a gene duplication event close to the origin of tetrapods (5, 16). Closer phylogenetic analyses in several taxa have revealed mammalian lineage-specific duplications, including the duplication and co-retention of several *Dnmt1* paralogs in marsupials (17) and the evolution of *Dnmt3L* from *Dnmt3A* in eutherian mammals (18). Another gene duplication from *Dnmt3B* gave rise to *Dnmt3C* in muroid rodents where it has acquired a distinct, non-redundant role in retrotransposon repression during spermatogenesis (19, 20). Thus, a series of ancient and recent gene duplications have led to the current repertoires of mammalian DNMTs.

Mammalian DNA methylation plays a critical role in retrotransposon silencing in the germline (21). Germ cell development is a particularly vulnerable stage for mammalian genomes, because many epigenetic chromatin marks that otherwise repress retrotransposons -like DNA methylation -are transiently erased. However, re-establishing silencing of retrotransposons in mammalian genomes is challenging, because of their vast sequence heterogeneity. Retrotransposons can belong to many evolutionarily distinct families that each exhibit rapid sequence divergence (22). In mice, during male fetal germ cell development in mice, two distinct waves of *de novo* methylation target retrotransposons according to their age (23). During the first wave, evolutionarily old retrotransposons gain methylation together with the rest of the genome. However, evolutionarily young and transcriptionally active retrotransposons are refractory to this wave and require the piRNA pathway - a small RNA-based defense system - to target DNA methylation to their promoters (23, 24).

Two recent studies showed that DNMT3C is the crucial enzyme for this process (19, 20). *Dnmt3C* knock-out (KO) males are sterile and their germ cell methylation profiles are similar to those of piRNA mutants, with a 1% drop in genome-wide DNA methylation content that selectively affects the promoters of young copies of LINE and ERVK retrotransposons (20, 23, 25). This contrasts with germ cell-specific *Dnmt3B* KO, which has no impact on male fertility (26), whereas constitutive *Dnmt3B* KO are embryonic lethal (10). On the other hand, germ cell-specific *Dnmt3*A KO males are infertile and display mild alteration of the methylation profiles of a few older retrotransposon families (26, 27). This suggests that *Dnmt3A* might contribute to the first wave of *de novo* methylation and act non-redundantly of *Dnmt3C* for the silencing of mouse retrotransposons.

Catalytically active DNMT3s have three well-defined domains. The most C-terminal region encodes the methyltransferase domain (MTase) that includes highly conserved protein motifs that catalyze the addition of methyl groups (28, 29). The central portion encodes two chromatin-reading domains that play important roles in their targeting and regulation (30). The ADD (ATRX-DNMT3-DNMT3L) domain is repelled by methylated states of the histone modification H3K4 (31-34), preventing *de novo* DNA methylation at H3K4me3-enriched gene promoters. The PWWP (Pro-Trp-Trp-Pro) domain, helps anchor DNMT3 proteins to nucleosomes marked by histone modification H3K36me (35-39). The H3K36me mark is enriched in the body of actively transcribed genes and pericentromeric heterochromatin. Interestingly, mouse *Dnmt3C* lost the two exons coding for the PWWP domain, making it unique among catalytically active DNMT3s (19, 20). In contrast to the central and C-terminal segments, the N-terminal portion of DNMT3s remain completely uncharacterized.

Based on both its recent origin and its role in silencing young, potentially rapidly adapting retrotransposon families, we speculated that *Dnmt3C* might be participating in ongoing genetic conflicts in rodent genomes (22). We therefore performed a detailed phylogenetic survey of rodent genomes to investigate *Dnmt3C*’s age and the evolutionary forces that shape its unique function. Extending previous findings, we dated *Dnmt3C’s* evolutionary origin in the common ancestor of muroids between 45 and 71 million years ago. Unexpectedly, we found a pattern of gene conversion between *Dnmt3B* and *Dnmt3C* paralogs throughout muroid evolution. This gene conversion recurrently homogenizes the latter two-thirds of DNMT3B and DNMT3C but does not extend to their N-terminal domains. We provide evidence of strong diversifying selection in the N-terminal tail of DNMT3C, but not DNMT3B, which we hypothesize is the result of an ongoing evolutionary arms race with active retrotransposons in these lineages. Although *Dnmt3C* is not present outside rodents, we found that the N-terminal tail of *DNMT3A* has similarly evolved under diversifying selection in primates. Thus, two distinct DNMT enzymes display hallmarks of ongoing genetic conflicts with retrotransposons in two separate mammalian lineages.

## Results

### Evolutionary origins and dynamics of *Dnmt3C* in rodents

To investigate the evolutionary age and dynamics of *Dnmt3C*, we retrieved and annotated DNMT3 sequences in partially or fully assembled genomes of 19 species of Glires – which include rodents and lagomorphs (Figure 1A, see methods). Like other mammals, most species of Glires encode unique *Dnmt3A, Dnmt3L* and *Dnmt3B* genes within syntenic loci present in all placental mammals (Figure 1A). However, in the sub-group of muroid species, the syntenic locus containing *Dnmt3B* also encodes *Dnmt3C* (Figure 1A) (20).

**Figure 1:**
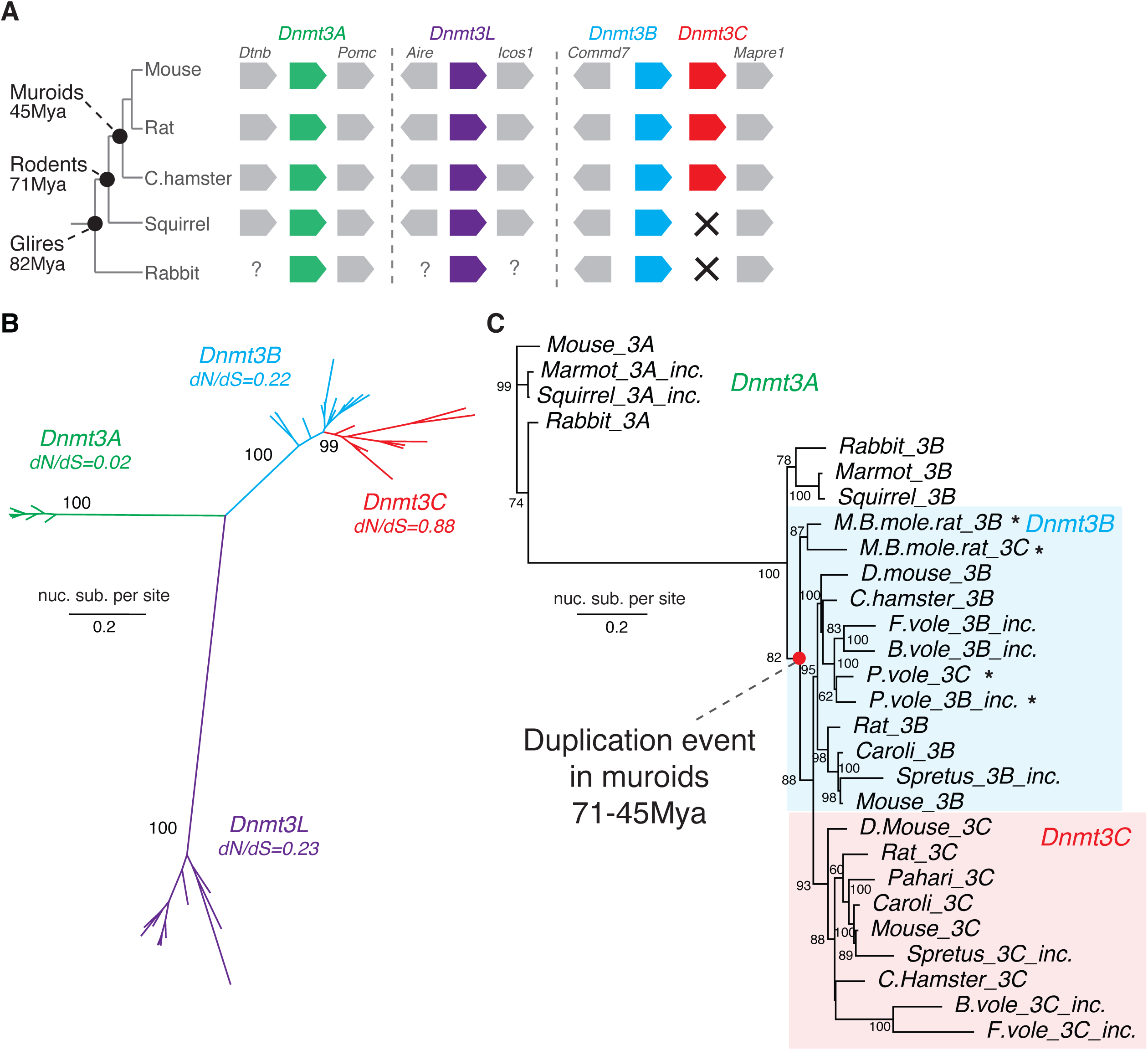
*Dnmt3C* duplicated in the last common ancestor of all muroids. (A) Schematic of the genomic locus encoding *Dnmt3* genes in representative species of Glires. Estimated divergence times based on Timetree analyses (40) are indicated at each major node. *Dnmt3* genes (colors) and corresponding neighboring genes (grey) are indicated with arrows. “?” denotes incomplete assembly while crosses denote absence of coding sequence. (B) Maximum likelihood nucleotide phylogeny of *Dnmt3* genes in Glires. Bootstrap values and average pairwise dN/dS are indicated for each clade. (C) Maximum likelihood phylogeny of all identified *Dnmt3B* and *Dnmt3C* genes. Incomplete sequences are indicated with “inc” while cases where *Dnmt3B* and *Dnmt3C* orthologs from the same species unexpectedly group together are highlighted with ‘*’. Bootstrap support values above 50% are reported. Also included are rabbit, marmot and squirrel *Dnmt3Bs* and selected *Dnmt3A*s. Abbreviations of species names: “M.B.mole.rat”: mountain blind mole rat; “D.mouse”: deer mouse; “C.hamster”: chineese hamster; “F.vole”: field vole; “B.vole”: bank vole; “P.vole”: prairie vole.

We investigated genomes from 11 muroid and 8 “outgroup” species and used available transcriptome or *de novo* gene assemblies to annotate coding sequences (CDS, see methods). In some cases, genome assemblies allowed us to tentatively assign gene orthology using shared synteny. However, in most cases, genome assemblies were too fragmented to reconstruct genomic contexts. Instead, we focused on retrieving partial or full-length sequences of putative *Dnmt* genes. We then constructed a multiple alignment and used maximum likelihood methods to build a gene phylogeny (see methods for details). Using this approach, we were able to resolve all retrieved sequences into distinct clades of DNMTs (Figure 1B).

If *Dnmt3C* arose from *Dnmt3B* in the last common ancestor of all muroids, we would expect *i) Dnmt3C* sequences to branch inside the *Dnmt3B* clade and *ii)* form two independent lineages following the split of muroids from other rodents and lagomorphs. Our first expectation was met; all putative *Dnmt3C* sequences branched within the *Dnmt3B* clade, supporting the close relatedness of these two genes relative to other *Dnmts* (Figure 1B). Moreover, a detailed phylogeny including all *Dnmt3B* and *Dnmt3C* orthologs was consistent with a single duplication event (Figure 1C). Based on the presence of *Dnmt3C* in mountain blind mole rats (*Nannospalax galili*), but not beavers or guinea pigs (*Castor canadensis* and *Cavia porcellus*), we estimate that the duplication occurred before the radiation of muroids between 45 and 71 million years ago (40).

However, the second expectation of independent evolution of *Dnmt3B* and *Dnmt3C* genes following *Dnmt3C’s* origin was not met. While most *Dnmt3B* and *Dnmt3C* genes grouped into two distinct clades according to the accepted muroid species phylogeny (41), both the prairie vole (*Microtus ochrogaster*) and the mountain blind mole rat (*Nannospalax galili*) had *Dnmt3B* and *Dnmt3C* paralogs that were more closely related to each other than to their respective orthologs (Figure 1C, asterisks). This pattern could indicate separate origins of *Dnmt3C* in these species or alternatively, recent gene conversion. It is also possible that partial gene conversion between *Dnmt3B* and *Dnmt3C* occurred in other muroid species but was not evident in phylogenetic analyses based on full-length sequences.

We therefore used a likelihood-based method, GARD, to map putative recombination breakpoints between *Dnmt3B* and *Dnmt3C* (see methods, (42)). Such analyses aim to identify recombination breakpoints based on segments of multiple alignments that have clearly discordant phylogenetic histories from each other. We identified three high confidence breakpoints in muroid *Dnmt3B* and *Dnmt3C* sequences, partitioning the aligned sequences into four segments with distinct evolutionary histories- A, B, C, and D (Figure 2A). Upon generating phylogenies of each segments independently, we observed a discordance between these segments was not limited to prairie vole and mountain blind mole rat (Figure 2B), suggesting that gene conversion between *Dnmt3B* and *Dnmt3C* occurred in many muroid lineages.

**Figure 2:**
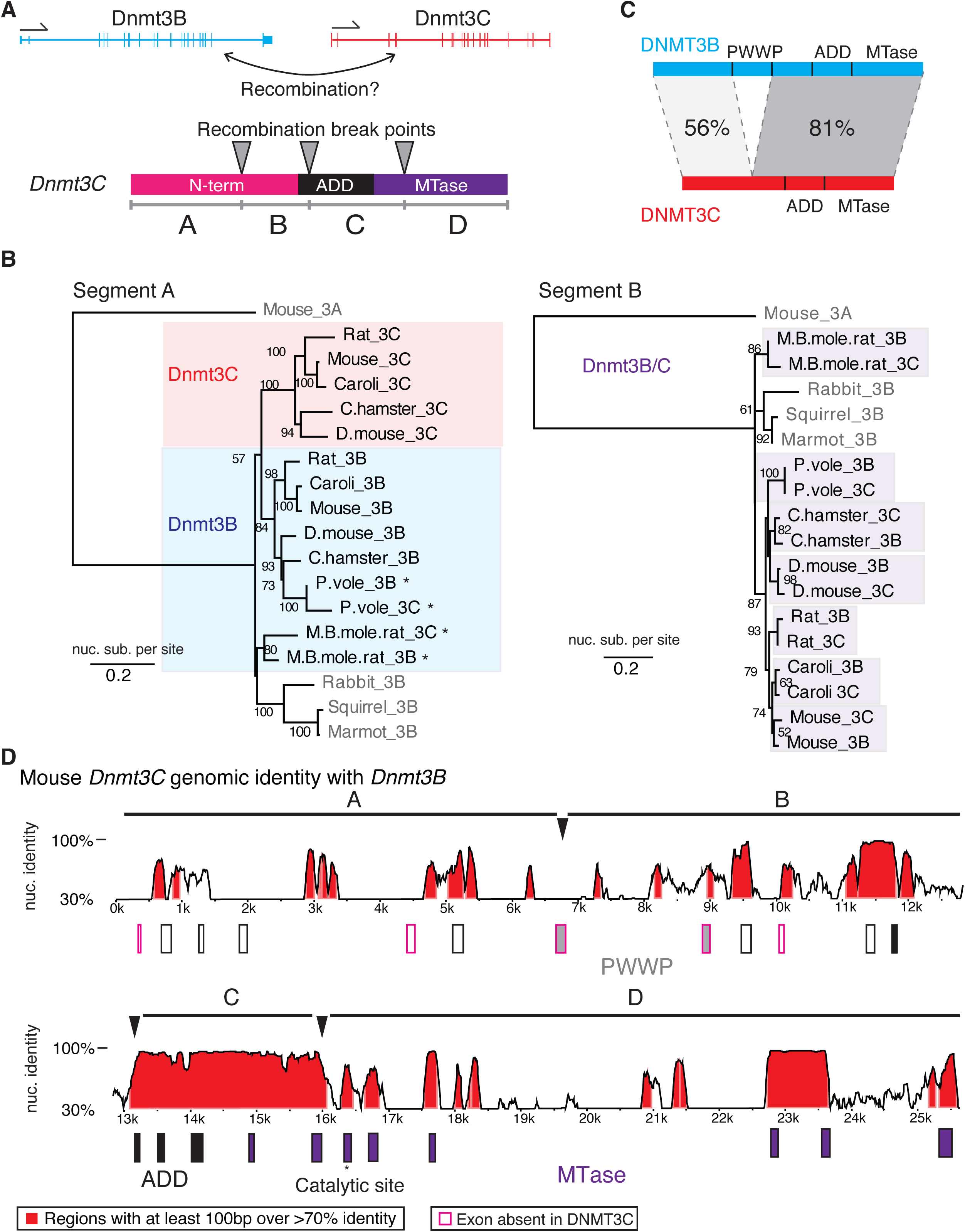
Gene conversion between *Dnmt3C* and *Dnmt3B* in muroids. (A) Schematic representation of *Dnmt3C* and *3B* genomic loci (top). Arrows indicate gene orientation in the mm10 assembly. Recombination map of *Dnmt3C* CDS is shown at the bottom. Breakpoints (arrowheads) identified by GARD (see methods) and recombination segments (“A” through “D”) are indicated on the *Dnmt3C* CDS with its predicted protein domains. (B) Maximum likelihood phylogenies of recombination segments A and B. Bootstrap supports above 50% are shown. (C) Average pairwise percent identities between muroid DNMT3C and DNMT3B proteins before and after the PWWP domain. (D) Map of the mouse *Dnmt3B* genomic locus with annotated exons (boxes). Percent nucleotide identity with *Dnmt3C* is plotted (y-axis) using mVISTA (see methods). Recombination segments (top) and predicted domains (bottom) are also shown.

Next, we investigated the individual evolutionary trajectories of the distinct recombination segments within *Dnmt3C*. Consistent with rampant gene conversion, nucleotide phylogenies showed that segment B and C - encoding the ADD and part of the MTase (Figure 2A) - grouped *Dnmt3C* and *Dnmt3B* paralogs by species (Figure 2B and Supplementary Figure S1) rather than by orthology groups. We also found evidence for gene conversion in Segment D, which encodes the rest of the MTase (Supplementary Figure S1B). With the exception of prairie voles and mountain blind mole rats, we found no evidence for gene conversion in segment A, which encodes the N-terminal tail of DNMT3C (Figure 2A). Indeed, a phylogeny based on segment A alone almost perfectly separated the *Dnmt3B* and *Dnmt3C* paralogs based on orthology groups, consistent with divergent evolution of the two genes following duplication (Figure 2B). Thus, in most muroids and in contrast to the rest of the gene, the 5’ ends of *Dnmt3B* and *Dnmt3C* do not appear to have engaged in recent gene conversion. Consistent with these findings, DNMT3B and DNMT3C protein sequences shared much higher homology in their C-terminal compared to their N-terminal domains (Figure 2C).

To further confirm our findings, we investigated the coding and non-coding genomic sequences of *Dnmt3B* and *Dnmt3C* for signatures of high sequence identity. High nucleotide identities between mouse *Dnmt3C* and *Dnmt3B* were evident not only in coding exons but also across many introns (Figure 2D). More specifically, all introns in segment C displayed more than 70% identity in segment C but not in segment A (Figure 2D), consistent with our recombination breakpoint analysis (Figure 2A). We found an even more evident pattern of sequence homogenization between *Dnmt3B* and *Dnmt3C* in genomes of rats and mountain blind mole rats (Figures S2C and S2D respectively). In particular, the high sequence identity between the *Dnmt3B* and *Dnmt3C* loci in mountain blind mole rats (Supplementary Figure S1D) supports the hypothesis that this species, as well as prairie voles (Figure 2B), engaged in gene conversion more recently that other muroids. Taken together, these results suggest that following duplication, *Dnmt3B* and *Dnmt3C* have been subject to extensive gene conversion, except in their 5’ ends. Thus, DNMT3C N-termini evolve under distinct evolutionary trajectories from their DNMT3B counterparts, while the central domains and C-termini of Dnmt3B, and Dnmt3C exchange sequences to remain similar within each genome.

### DNMT3C N-terminal domain evolve under positive selection

Gene conversion has homogenized several segments of DNMT3C and DNMTB, but not their N-terminal domains. We hypothesized that this could be to retain the functional divergence of DNMT3B and DNMT3C in their N-terminal domains. For example, loss of the ancestral PWWP domain in DNMT3C may have allowed it to specialize for functions distinct from DNMT3B. If this were the case, we might expect to find additional differences in the selective constraints that act on *Dnmt3B* versus *Dnmt3C*, especially in their N-terminal domains. We, therefore, investigated how the DNMT3B and DNMT3C N-terminal domains may have diverged in their selective constraints.

Based on branch lengths (Figure 1B), *Dnmt3C* appears to be the most divergent of all *Dnmt3* genes in muroid rodents, followed by *Dnmt3L, Dnmt3B*, and finally *Dnmt3A*, which is most highly conserved. To evaluate selective constraints, we calculated the rates of non-synonymous (amino-acid altering, dN) and synonymous (silent, dS) substitutions across orthologous sequences of all *Dnmt* genes. *Dnmt3C* displays the highest average pairwise dN/dS of all *Dnmt3* genes (0.88) compared to *Dnmt3L* (0.23), *Dnmt3B* (0.22), and *Dnmt3A* (0.02) (Figure 1B). Higher dN/dS values could reflect relaxation of selective constraint. Alternatively, these higher values could be the result of diversifying selection acting on *Dnmt3C*.

To distinguish between these possibilities, we used likelihood methods implemented in the PAML package to detect signatures of positive selection (43). Muroid species are an ideal species set for these analyses because they span a short evolutionary time (∼40 million years) with low saturation of dS (41). We separately analyzed each of the four recombination segments across all orthologs identified in muroids. Because some *Dnmt3C* genes are based on incomplete gene models, each segment alignment contained between 8 to 11 species (Table 1). We then used PAML to identify site that were subject to positive selection (see methods) (43). We found no evidence of positive selection having acted on *Dnmt3B*. In contrast, we found strong support for positive selection having acted on segment A of *Dnmt3C*, but not on segments B, C, or D (Table1).

**Table1:** Summary of selection tests across muroid *Dnmts* genes. Recombination segments of *Dnmt3C* were analyzed independently, while full-length sequences were used for other *Dnmts*. P-values are for likelihood ratio tests between substitution models allowing or not allowing for positive selection using codeml (PAML). See text and methods for details.

|  | Seg. | nb.<br>species | Length<br>(bp) | PAML - M7<br>vs M8 p-value | PAML - M8a<br>vs M8 p-value | M(0)<br>dN/dS | % sites<br>dN/dS>1<br>(avg. dN/dS) | Sites<br>BEB≥90% |
| --- | --- | --- | --- | --- | --- | --- | --- | --- |
| *Dnmt3C* | **A** | 11 | 471 | 0.002 | 0.004 | 0.886 | 49% (1.62) | 54(T),<br>57(Q),<br>95(P), 96<br>(L) |
|  | **B** | 8 | 135 | 0.509 | 0.965 | 0.176 | - | N/A |
|  | **C** | 8 | 546 | 1.000 | 0.827 | 0.114 | - | N/A |
|  | **D** | 10 | 666 | 1.000 | 0.907 | 0.184 | - | N/A |
| *Dnmt3B* | **all** | 8 | 2052 | 0.170 | 0.475 | 0.116 | 1% (1.59) | N/A |
| *Dnmt3A* | **all** | 9 | 2718 | 0.823 | 0.463 | 0.022 | - | N/A |
| *Dnmt3L* | **all** | 9 | 1218 | 0.546 | 0.732 | 0.271 | - | N/A |

In segment A of *Dnmt3C*, PAML analyses estimated 49% of sites to have evolved with an average dN/dS greater than 1 indicative of potential diversifying selection; their average dN/dS was estimated to be 1.6. Of these, four sites were highlighted with a high posterior probability of having evolved under positive selection (Bayes Empirical Bayes [P] ≥ 90%, Table1 and Figure 3B). These sites (codons 54, 57, 95 and 96 in mouse *Dnmt3C*) all cluster within the most 5’ end of the gene (first 300bp of the CDS) and display extensive diversification in both charge and hydrophobicity across muroids (Figure 3B). For site 95 and 96, rapid evolution disrupts a highly conserved arginine patch of unknown function, which is highly conserved among muroid DNMT3B proteins (Figure 3C). Thus, in addition to the loss of the PWWP domain, DNMT3B and DNMT3C differ in the selective constraints they are subject to. The signature of positive selection and loss of the PWWP domains makes *Dnmt3C* unique among all *Dnmt3* genes.

**Figure 3:**
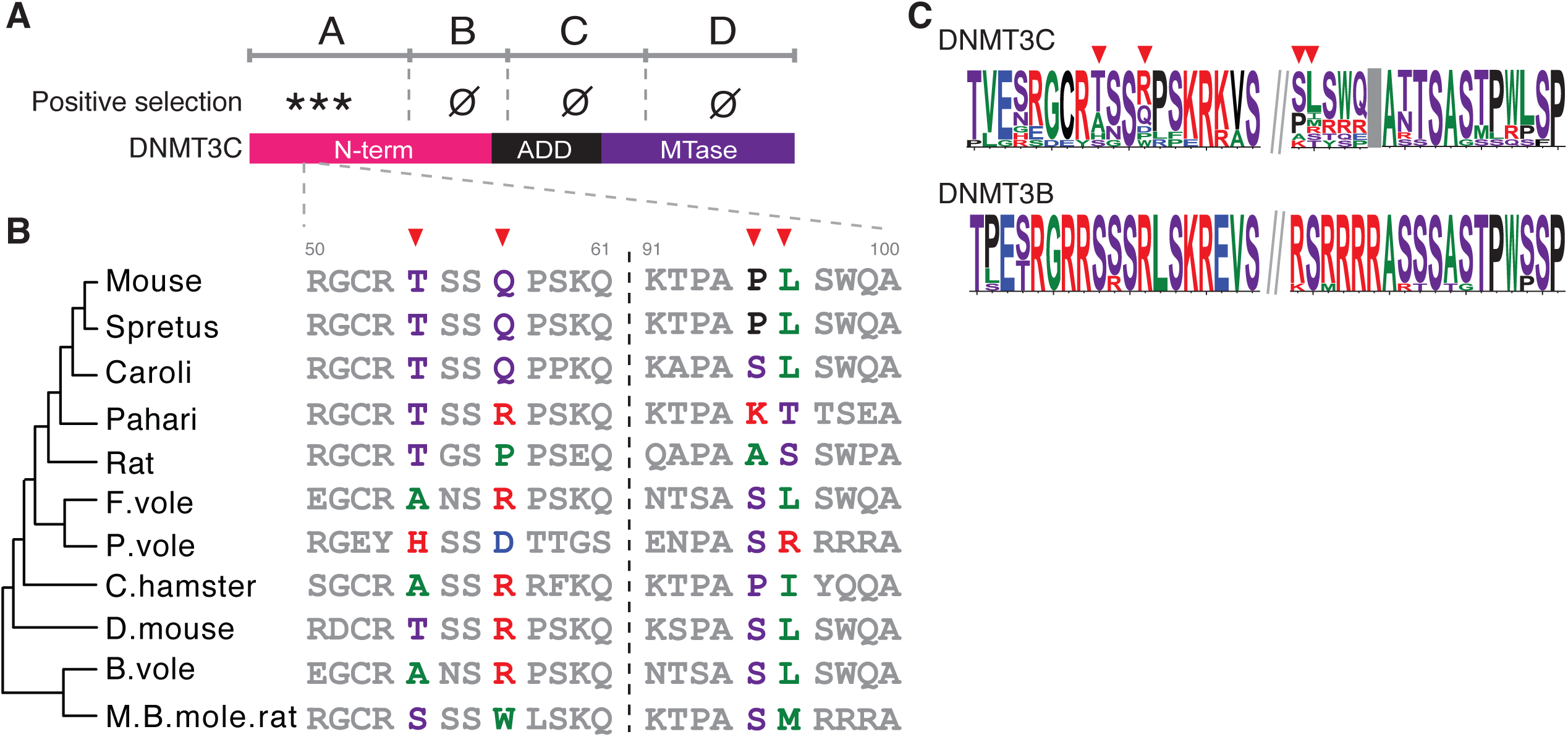
DNMT3C N-terminus is subject to positive selection. (A) Schematic representation of positive selection test results for all recombination segments of DNMT3C CDS. “***” denotes the finding of positive selection in segment A (see text and methods for details), whereas “Ø” indicates no support for positive selection. (B) Amino acid alignments (positions 50 to 61, and 91 to 100) of muroid DNMT3Cs showing four positively selected sites identified with PAML (red arrowheads). Sequences are arranged according to segment A phylogeny with species names on the right. (C) Amino acid logos of DNMT3C and 3B around the positively selected sites (arrowheads). Backslashes indicates sequences not shown. The grey box denotes an alignment gap between DNMT3C and 3B.

### Selective constraints acting on DNMT3 proteins in primate genomes

The evolutionary birth of *Dnmt3C* afforded muroid rodents a unique opportunity to silence young, active retrotransposon families by DNA methylation. However, most mammalian genomes face a similar pressure by young retrotransposon lineages and yet do not encode *Dnmt3C*. We, therefore, hypothesized that non-rodent mammalian species might deploy alternate mechanisms, possibly other DNMT enzymes, to achieve DNMT3C-like repression of their active retrotransposons. If true, we might expect *Dnmt3* genes to be locked in these molecular arms-races, and therefore subjected to similar selective pressures as *Dnmt3C* in muroid rodents *i*.*e*., diversifying selection.

To investigate this possibility, we analyzed the evolutionary constraints that act on *DNMT3* genes in primates, a distinct lineage of mammals that have substantial genomic resources across multiple species in a comparable evolutionary timespan to muroid rodents (40). Using maximum likelihood-based analyses, we found strong evidence of diversifying selection acting on *DNMT3A* and marginal evidence of positive selection in the catalytically-inactive co-factor *DNMT3L* (Table 2). In contrast, we found no evidence of diversifying selection in primate *DNMT3B* (Table 2) or muroid *DNMT3A* (Table 1).

**Table 2:** Summary of selection tests across primate DNMTs. (A) Codeml (PAML) analyses using the accepted species phylogeny. P-values are for likelihood ratio tests between substitution models allowing or not allowing for positive selection.

|  | nb.<br>species | Length<br>(bp) | PAML - M7 vs<br>M8 p-value | PAML - M8a vs<br>M8 p-value | M(0) dN/dS | % sites dN/dS>1<br>(avg. dN/dS) | Sites BEB≥90% |
| --- | --- | --- | --- | --- | --- | --- | --- |
| *DNMT3A* | 20 | 2724 | 0.022 | ≤ 0.001 | 0.047 | 2.02 (1.8) | 66 (P), 81 (A) |
| *DNMT3B* | 21 | 2481 | 0.149 | 0.430 | 0.080 | 1.81 (1.7) | N/A |
| *DNMT3L* | 20 | 1155 | 0.018 | 0.124 | 0.204 | 5.29 (1.6) | N/A |

As in muroid *Dnmt3C*, the diversifying selection signature also appeared to primarily map to the N-terminal domain of primate *DNMT3A* (codons 61 and 81, Table 2 and Figure 4). To rule out that this signature could be due to unaccounted recombination, we performed GARD analyses. This identified a single break point (within the first 1kb of the CDS), however, while a maximum likelihood phylogeny of the first segment (including the rapidly evolving sites) had strong bootstrap support, the second segment did not (Supplementary Figure S2). Further inspecting this segment showed a high rate of CpG mutations which prevents appropriate reconstruction of its evolution and accurate selection analyses. We therefore conclude that there is not sufficient evidence for gene conversion affecting *DNMT3A* evolution in primates. In spite of this, PAML analysis of *DNMT3A* putative 1^st^ (N-terminal) segment also identifies sites 61 and 81 as evolving under positive selection (not shown).

**Figure 4.**
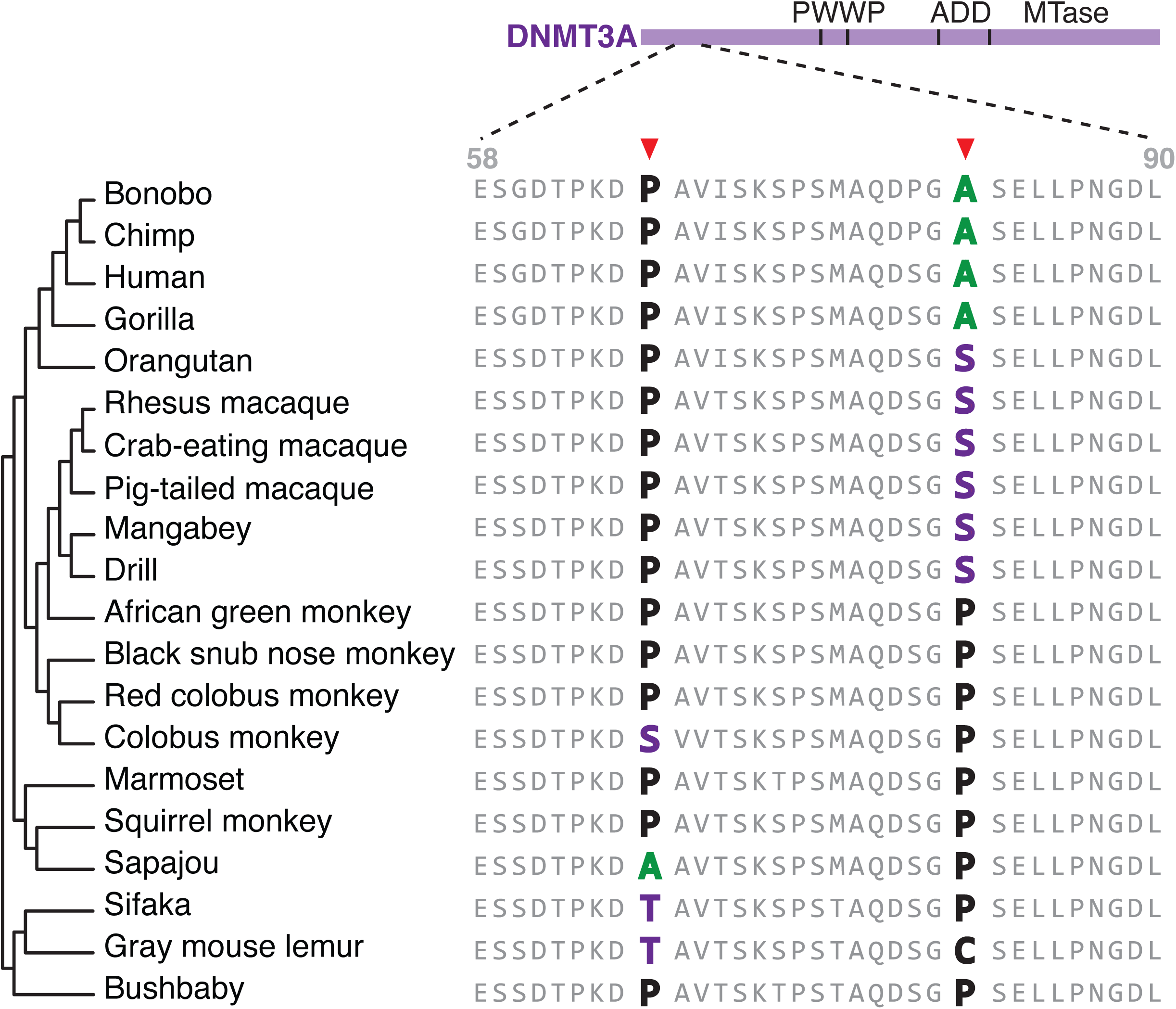
Positive selection of N-terminal domain of primate DNMT3A. Amino acid alignments (position 58 to 90) of primate DNMT3As showing the two positively selected sites identified with PAML (arrowheads). Sequences are arranged according to the accepted species phylogeny with species names on the right.

Thus, rather unexpectedly, we find evidence of diversifying selection on distinct *DNMT3* genes in rodent and primate genomes (Tables 1 and 2). Our findings suggest that the N-terminal portions of DNMT3 proteins wage evolutionary arms races for DNA methylation of young, active retrotransposons in different mammalian lineages. They further raise the possibility that DNMT3A, which is universal to all mammals, may be the original DNMT that represses young retrotransposons. The subsequent birth of DNMT3C in muroid rodents may have absolved DNMT3A of this role, which could be why we cannot detect any signatures of diversifying selection in *Dnmt3A* in rodent species.

## Discussion

Retrotransposons pose a significant fitness challenge to host genomes. To protect themselves, host genomes deploy multipronged strategies to curb retrotransposon activity. Here we identified the selective forces shaping the function of a recently duplicated DNA methyltransferase, DNMT3C, that specifically targets evolutionarily young retrotransposons in muroid rodents. We found that *Dnmt3C* has undergone recurrent gene conversion with its parental gene *Dnmt3B*, except for the N-terminal domain. These findings are reminiscent of previous studies of gene families subject to genetic conflicts (44). For example, the true evolutionary histories of the mammalian antiviral *IFIT1/IFIT1B* paralogs, which diverged 100 million years ago, were also confounded by recurrent gene conversion (45). Similarly, recurrent gene conversion affected the histone-fold domain but not the distinct N-terminal tails of centromeric histone paralogs in *Drosophila* species (46). In all these cases (*Dnmt3, IFIT, Cid*) as well as several additional examples (44), natural selection likely selects against gene conversion in the domain that likely drives the functional diversification of the paralogs. As a result, only the non-converted gene segment can recreate the true evolutionary history of the paralogs. In all these cases, the close proximity of the paralogs following gene duplication might have facilitated multiple episodes of gene conversion between them.

We found that the N-terminal domain of *Dnmt3C*, but not its parental gene *Dnmt3B*, has evolved under strong diversifying selection. Diversifying selection - especially in a host ‘defense’ gene - is a signature of an evolutionary arms race between host genomes and retrotransposons (22). As host genomes deploy repressive chromatin strategies, retrotransposons must adapt to ward off host repression, in turn spurring host adaptation. The evolutionary arms race model further makes the prediction that residues or domains that directly engage in the antagonism should be rapidly evolving. Thus, the positive selection in *Dnmt3* genes could be the result of active antagonism by a product of young retrotransposons (RNA or protein). Under this model, the positive selection would allow the DNMT3 proteins to evade binding and antagonism by young retrotransposons.

We favor the alternate model that positive selection shapes the targeting of DNMT3 proteins to young retrotransposons to mediate their silencing. This predicted activity would be similar to the KZNF (KRAB domain containing Zinc Finger) proteins, which use rapid evolution of their DNA-binding domains to keep pace with a changing nucleotide landscape of retrotransposon families (47). We hypothesize that similar evolutionary dynamics could drive the diversifying selection of the N-terminal domains in rodent DNMT3C and primate DNMT3A proteins. Interestingly, DNMT3A exists both as a long isoform, A1, and a N-terminus-less short isoform, A2 (48), as a result of alternative promoter usage. We posit that the long DNMT3A1 isoform may target young retrotransposons in male germ cells in DNMT3C-less mammalian species, such as primates.

The recurrent signature of rapid evolution within the N-termini of two different DNMT3 proteins in different mammalian lineages may highlight a novel functional domain that may be key to DNMT targeting of retrotransposons. Unlike the canonical PWWP, ADD and MTase domains, however, this domain may be characterized by its rapid evolution rather than conservation. How this domain engages with retrotransposons remains to be determined. In contrast to KZNF proteins, there is no suggestion that DNMT3 proteins have DNA sequence-binding specificity. Instead, it is possible that this region mediates interaction with components of the piRNA pathway - some of which are rapidly evolving in other animals (49, 50).

In sum, the DNMT3C N-terminal domains can be distinguished from other DNMT3 proteins in two key aspects: diversifying selection but also loss of a coding PWWP domain, which is essential for coupling *de novo* DNA methylation to local chromatin environment, via recognition of H3K36 methylated histones (36, 37, 39). In DNMT3B, the PWWP domain binds H3K36me3 marks, which are typical of transcribed gene bodies (51). In DNMT3A, the PWWP domain is intact and was recently shown to mediate DNMT3A-dependent methylation of intergenic sequences (52). We hypothesize here that DNMT3C’s N-terminal domain may be required to substitute for PWWP-dependent chromatin-targeting function. However, the mode of targeting of DNMT3C to young retrotransposon promoters remains to be determined.

In conclusion, our evolutionary studies identified a new functional domain in DNMT3C, a DNA methyltransferase that mediates full silencing of the most active, rapidly-adapting retrotransposon families in rodent genomes. Furthermore, based on our findings of diversifying selection in primate DNMT3A, we suggest that diversifying selection of enzymes that methylate retrotransposons in developing germ cells might be pervasive across mammalian genomes, although this targeting may be mediated by distinct DNMT3 paralogs.

## Methods

### Identification of DNMT3 orthologs

To identify *Dnmt* orthologs we performed tBLASTn searches on the NCBI non-redundant nucleotide database (53, 54), using reference protein sequences of mouse DNMT3A (NP_031898.1), DNMT3B (XP_006498745.1), DNMT3L (NP_001075164.1) as well as the predicted protein sequence from the *Dnmt3C* cDNA cloned from male fetal gonads (20). While most *Dnmt3s* have predicted sequences in reference databases, *Dnmt3c* genes are not annotated in most muroid genomes. In these cases we queried genomes directly using tBLASTn, and predicted gene models from contigs using GeneWise (55). CDSs were annotated based on the longest mouse gene model.

### Queried genomes

We used the following genome assemblies to predict *Dnmt3B* and *Dnmt3C* gene models. Muroids: *Mus musculus* (UCSC mm10), *Mus spretus* (Sanger, SPRET_EiJ), *Mus caroli* (Sanger, CAROLI_EiJ), *Mus pahari* (Sanger, Pahari_EiJ), *Apodemus sylvaticus* (NCBI, GCA_001305905.1_ASM130590v1), *Rattus norvegicus* (UCSC, rn6), *Peromyscus maniculatus* (NCBI, GCF_000500345.1_Pman_1.0), *Myodes glareolus* (NCBI, GCA_001305785.1_ASM130578v1), *Microtus agrestis* (NCBI, GCA_001305995.1_ASM130599v1), *Microtus ochrogaster* (NCBI, MicOch1), *Mesocricetus auratus* (NCBI, MesAur1), *Cricetulus griseus* (UCSC, criGri1), *Nannospalax galili* (NCBI, GCF_000622305.1_S.galili_v1.0).

Glires: *Castor canadensis* (NCBI, C.can genome v1.0), *Oryctolagus cuniculus* (UCSC, oryCun2), *Marmota marmota* (NCBI, GCF_001458135.1_marMar2.1), *Ictidomys tridecemlineatus* (UCSC, speTri2), *Cavia porcellus* (Broad Institute cavPor3).

### Species divergence times

Divergence time estimates were obtained from using timetree.org (40), by specifying sister taxa that belong to either Glires, rodents or muroids. Timetree outputs a range of estimated divergence times summarizing phylogenetic and fossil dating.

### Synteny analysis

Shared synteny blocks were identified using the online server Genomicus (V95.1) (16). Mouse was used as a reference locus and individual synteny blocks were inspected using the UCSC genome browser (56).

### Alignments and phylogenies

All sequence alignments are available as supplementary material. Alignments were generated using ClustalW v2.1 (IUB cost matrix, (57)) or MAFFT v7.388 (58). Maximum likelihood phylogenies were built using PHyML v3.0 with 100 bootstraps (59). Trees were visualized using the software Geneious Prime (Biomatters Ltd). In all cases we used nucleotide alignments of the CDS and the HKY85 substitution model.

### Detection of recombination

To test for recombination, we used an alignment of *Dnmt3C* and *3B* CDS from six species with nearly complete gene models (mouse, Mus caroli, rat, prairie vole, chinese hamster, and mountain blind mole rat). Assembly gaps were removed. To detect recombination breakpoints, we used GARD with the general discrete model of site to site variation and three rate classes (42). We kept breakpoints with right and left p-values below 0.01. We subsequently segmented the *Dnmt3C* alignment according to these breakpoints. Similarly, recombination in primate *DNMT3A* was tested using an alignment of all primate CDS.

### Genomic alignments

To identify region of homology between *Dnmt3C* and *Dnmt3B* genomic loci, we extracted the regions from assembled genomes of the mouse and rat and contigs of mountain blind mole rat and aligned them using mVista (60). Exon annotations were based on reference alignments with the species CDS.

### Selection analyses

We measured overall dN/dS rates with codeml, PAMLX V1.3.1 (43), under model 0 and average pairwise with SNAP V2.1.1 (61). We tested for positive selection using codon alignments generated with PAL2NAL (62) free of any gaps and stop codons and with either accepted species or gene phylogenies. We compared “NSsites” evolutionary models that either do not allow dN/dS to exceed 1 (M7 or M8a) to a model that does (M8). We tested for statistical significance using a chi-square test of the twice the difference in log-likelihoods between M8 and matched null model M7 or M8a, with the degrees of freedom reflecting the difference in number of parameters between the two models compared (43). Positively selected sites were classified as those sites with M8 Bayes Empirical Bayes posterior probability > 90%. The results we present are from codeml runs using the F3×4 codon frequency model, and initial omega 0.4. Analyses were robust to use of different starting parameters (codon frequency model F61; starting omega 1.5).

### DNMT3C and DNMT3B logo plots

Logo plots were generated using weblogo.berkeley.edu (63); using all muroid species with alignable sequences over these exons: mouse (*Mus musculus*), *Mus spretus, Mus caroli*, rat, deer mouse, field vole, prairie vole, bank vole, chinese hamster, and mountain blind mole rat.

## Acknowledgements

We would like to thank members of the Bourc’his and Malik laboratories, especially Joan Barau, Tera Levin and Janet Young for technical help and critical reading of this manuscript. This work was supported by a postdoctoral fellowship from the Damon Runyon Foundation (to A.M.), NIH grant R01 GM074108 (to H.S.M.) and an HHMI Investigator award (to H.S.M.). The laboratory of D.B. is part of the Laboratoire d’Excellence (LABEX) entitled DEEP (11-LBX0044).

## Figure legends

**Supplementary Figure S1**: (A) Maximum likelihood nucleotide phylogeny of recombination segment C. (B) Maximum likelihood nucleotide phylogeny of recombination segment D. Bootstrap support is shown for each node. Scale bars show nucleotide substitutions per site. (C) Map of the rat *Dnmt3B* genomic locus with annotated exons (boxes). Percent nucleotide identity with *Dnmt3C* is plotted (y-axis). Predicted domains are shown at the bottom. (D) as in (C) but for mountain blind mole rat sequences.

**Supplementary Figure S2**: Maximum likelihood nucleotide phylogeny of putative recombination segment A (blue, 0 to 1041bp) and B (red, 1042 to 2727bp) of primate DNMT3A found with GARD (see methods). Bootstrap support is shown for each node. Nodes with low bootstrap support are shown in red. Scale bars show nucleotide substitutions per site.

**Supplementary Files 1 to 8:** Alignments of *Dnmt* sequences used in this study.

