## Supplementary figures and images for "Dynamic evolution of *de novo* DNA methyltransferases in rodent and primate genomes"

### Suppl Figure 1

#### Supplementary Figure S1

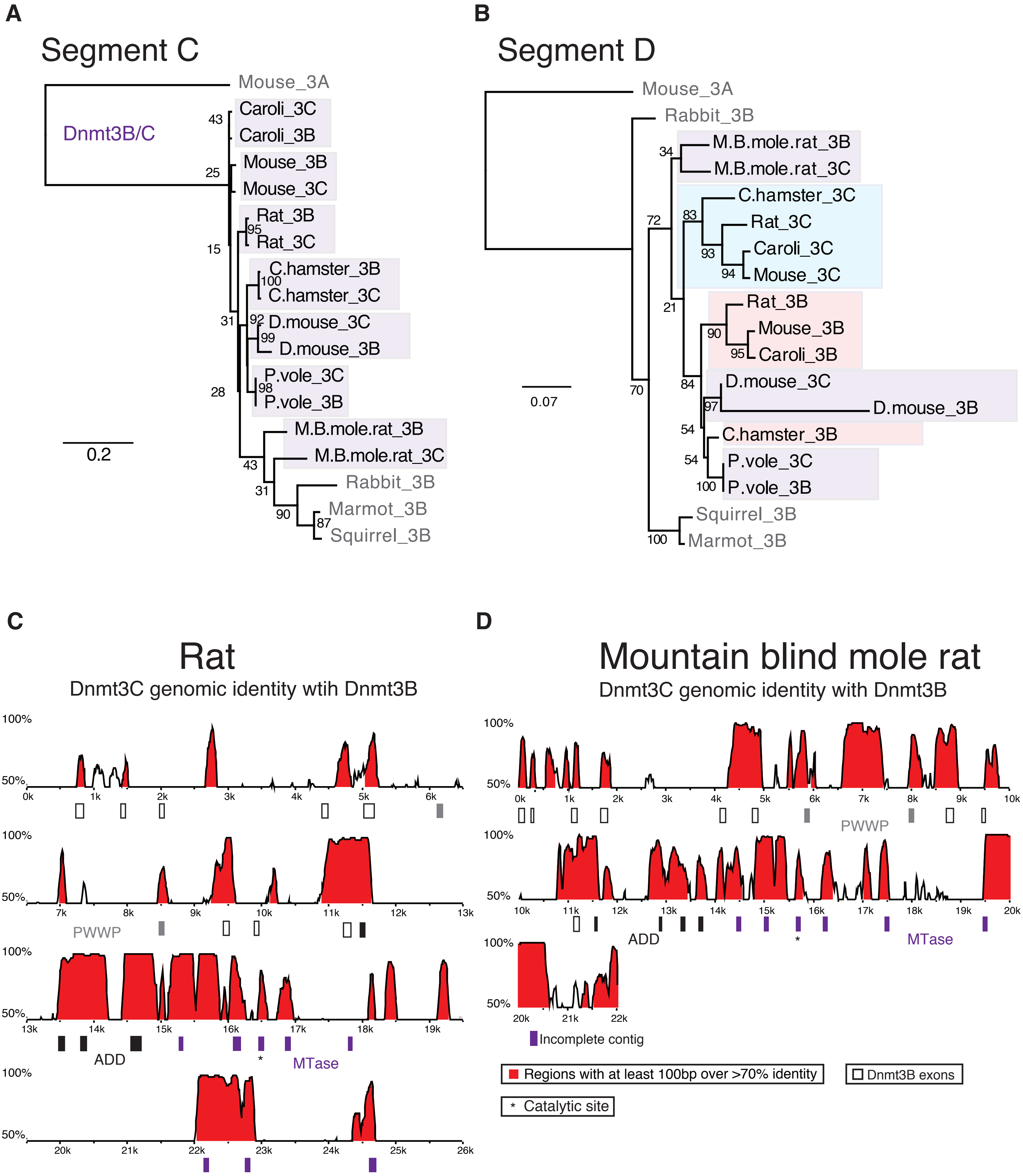

### Suppl Figure 2

**Supplementary Figure S2**

Pimate *DNMT3A*

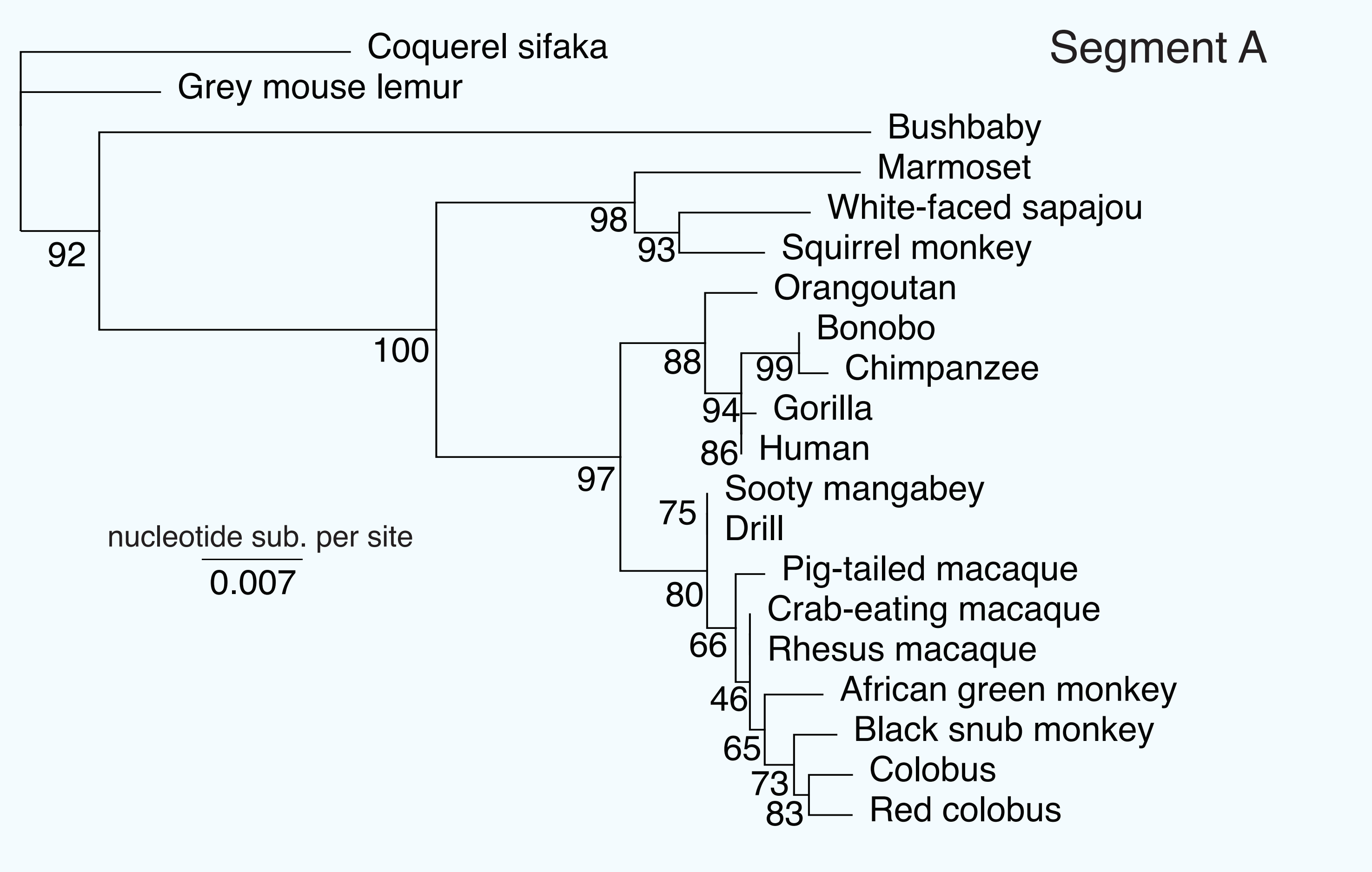


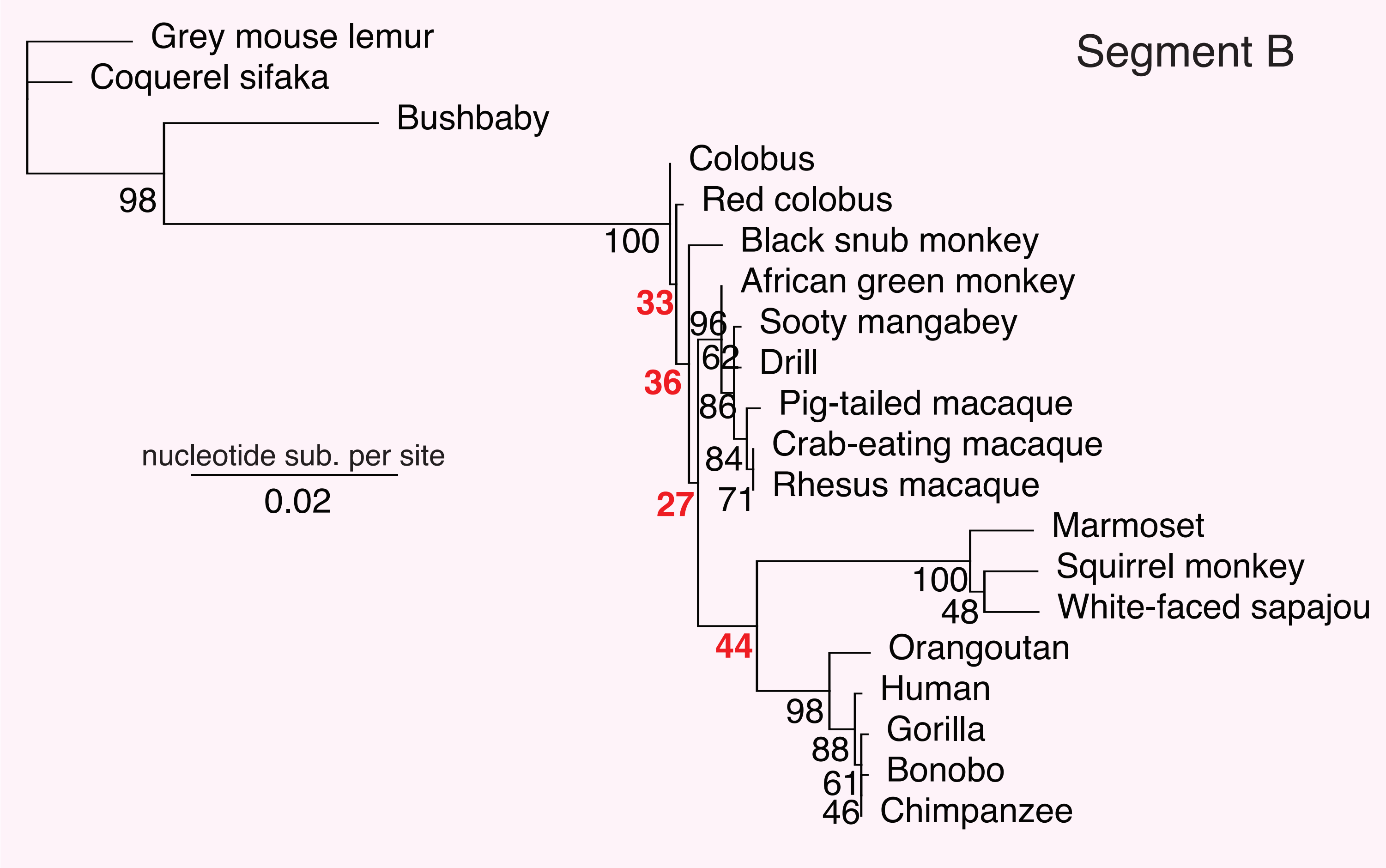
