## Supplementary material for "Dynamic evolution of *de novo* DNA methyltransferases in rodent and primate genomes": Suppl data 1

*SupFile_1:Sequences alignments used for Figure 1B*

*>Mouse3A*

*ATGCCCTCCAGCGGCCCCGGGGACACCAGCAGCTCCTCTCTGGAGCGGGAGGATGATCGAAAGGAAGGAGAGGAACAGGAGGAGAACCGTGGCAAGGAAGAGCGCCAGGAGCCCAGCGCCACGGCCCGGAAGGTGGGGAGGCCTGGCCGGAAGCGCAAGCACCCACCGGTGGAAAGCAGTGACACCCCCAAGGACCCAGCAGTGACCACCAAGTCTCAGCCCATGGCCCAGGACTCTGGCCCCTCAGATCTGCTACCCAATGGAGACTTGGAGAAGCGGAGTGAACC---CCAACCTGAGGAGGGGAGCCCAGCTGCAGGGCAGAAGGGTGGGGCCCCAGCTGAAGGAGAGGG---AACTGAGACCCCACCAGAAGCCTCCAGAGCTGTGGAGAATGGCTGCTGTGTGACCAAGGAAGGCCGTGGAGCCTCTGCAGGAGAGGGCAAAGAACAGAAGCAGACCAACATCGAATCCATGAAAATGGAGG--------GCTCCCGGGGCCGACTGCGAGGTGGCTTGGGCTGGGAGTCCAGCCTCCGTCAGCGACCCATGCCAAGACTCACCTTCCAGGCAGGGGACCCCTACTACATCAGCAAAC---------------GGAAACGGGATGAGTGGCTGGCACGTTGGAAAA------------------------GGGAGGCTGAGAAGAAAGCCAAGG-TAATTGCAGTAATGAATGCTGTGGAAGAGAAC--CAGGCCTCTGGAGAGTCTCAGAAGGTGGAGGAGGCC-AGCCCTCCTGCTGTGCAGCAGCCCACGGACCCTGCTTCTCCGACTGTGGCCACCACCCCTGAGCCAGTAGGAGGGGATGCTGGGGACAAGAATGCTACCAAAGCAGCCGACGATGAGCCTGAGTATGAGGATGGCCGGGGCTTTGGCATTGGAGAGCTGGTGTGGGGGAAACTTCGGGGCTTCTCCTGGTGGCCAGGCCGAATTGTGTCTTGGTGGATGACAGGCCGGAGCCGAGCAGCTGAAGGCACTCGCTGGGTCATGTGGTTCGGAGATGGCAAGTTCTCAGTGGTGTGTGTGGAGAAGCTCATGCCGCTGAGCTCCTTCTGCAGTGCATTCCACCAGGCCACCTACAACAAGCAGCCCATGTACCGCAAAGCCATCTACGAAGTCCTCCAGGTGGCCAGCAGCCGT--GCCGGGAAGCTGTTTCCAGCTTGCCATGACAGTGATGAAAGTGACAGTGGCAAGGC----TGTGGAAGTGCAGAACAAGCAGATGATTGAATGGGCCCTCGGTGGC--TTCC---AGCCCTCGGGTCCTAAGGGCCTGGAGCC--ACCAGAAGAAGAGAAGAATCCTTA-----------------------------------------------------CAAGGAAGTTTACACCGACATGTGGGTGGAGCCTGAAGCAGCTGCTTACGCCCCACCCCCACCAGCCAAGAAACCCAGAAA--GAGCACAACAGAGAAACCTAAGGTCAAGGAGATCATTGATGAGCGCACAAGGGAGCGGCTGGTGTATGAGGTGCGCCAGAAGTGCAGAAACATCGAGGACATTTGTATCTCATGTGGGAGCCTCAATGTCACCCTGGAGCACCCACTCTTCATTGGAGGCATGTGCCAGAACTGTAAGA---------ACTGCTTCTTGGAGTGTGCTTACCAGTA-TGACGACGAT-GGGTACCAGTCCTATTGCAC----CATCTGCTGTGGGGGGCGT-----------------GAAGTGCTCATGTGTGGGA----------------------------ACAACAAC--------------TGCTGCAGGTGC-----------------------TTTTGTGTCGAGTGTGTGGATCTCTTGG--TGGGG--CCAGGAGCTGCT-----CAGGCAGCCATTAA------GGAAGACCCCTGGAAC-TGCTACATGTGCGGGCATAAGGGCACCT---ATGGGCTGCTGCGAAGACGGGAAGACTGGCCTTCTCGACTCCAGATGTTCTTTGCCAATAACCATGACCAGGAA------TTTGACCCCCCAAAGGTTTACCCACCTGTGCCAGCTGAGAAGAGGAAGCC------------CATCCGCGTGCTGTCTCTCTTTGA-TGGGATTGCT---ACAGGGCTCCTGGTGCTGAA--GGACCTG---------GGCATCCAAGTGGACCGCTACATTGCCT----CCGAGG--TGTGTGAGGACTCCATCACGGTGGGCATGGTGCGGCACCAGGGAAAGATCATGTACGTCGGGGACGTCCGCAGCGTCACACAGAAGCATATCCAGGAGTGGGGCCCATTCGACCTGGTGATTGGAGGCA---------------------------------------GTCCCTGCAATGACCTCTCCATTGTCAACCCTGCCCGCAAGGGACTTTATGAGGGTACTGGCCGCCTCTTCTTTGAGTTCTACCGCCTCCTGCATGATGCGCGGCCCAAGGAGGGAGATGATCGCCCCTTCTTCTGGCTCTTTGAGAATGTGGTGGCCATGGGCGTTAGTGACAAGAGGGACATCTCGCGATTTCTTGAGTCTAACCCCGTGATGATTGACGCCAAAGAAGTGTCTGCTGCACACAGGGCCCGTTACTTCTGGGGTAACCTTCCTGGCATGAA---CAGGCCTTTGGCATCCACTGTGAATGATAAGCTGGAGCTGCAAGAGTGTCTGGAGCACGGCAGAATAGCCAAGTTCAGCAAAGTGAGGACCATTACCACCAGGTCAAACTCTATAAAGCAGGGCAAAGACCAGCATTTCCCCGTCTTCATGAACGAGAAGGAGGACATCCTGTGGTGCACTGAAATGGAAAGGGTGTTTGGCTTCCCCGTCCACTACACAGACGTCTCCAACATGAGCCGCTTGGCGAGGCAGAGACTGCTGGGCCGATCGTGGAGCGTGCCGGTCATCCGCCACCTCTTCGCTCCGCTGAAGGAATATTTTGCTTGTGTGTAA*

*>Caroli3A*

*ATGCCCTCCAGCGGCCCCGGGGACAGCAGCAGCTCCTCTCTGGAGCGGGAGGATGCTCGAAAGGAAGGAGAGGAACAGGAGGAGAACCGTGGCAAGGAAGAGCGCCAGGAGCCCAGTGCCACGGCCCGGAAGGTGGGGAGGCCTGGCCGGAAGCGCAAGCACCCACCGGTGGAAAGCAGTGACACCCCCAAGGACTCAGCGGTGACCACCAAGTCTCAGCCCATGGCCCAGGACTCTGGCCCCTCAGATCTGCTACCCAATGGAGACTTGGAGAAGCGGAGTGAACC---CCAACCTGAGGAGGGAAGCCCAGCTGCAGGGCAGAAGGGTGGGGCCCCAGCTGAAGGAGAGGG---AACTGAGACCCCACCAGAAGCCTCCAGAGCTGTGGAGAATGGCTGCTGTGTGACCAAGGAAGGCCGTGGAGCCTCTGCAGGAGAGGGCAAAGAACAGAAGCAGACCAACATCGAATCCATGAAAATGGAGG--------GCTCCCGGGGCCGACTGAGAGGTGGCTTGGGCTGGGAGTCCAGCCTCCGTCAGCGGCCCATGCCAAGACTCACCTTCCAGGCAGGGGACCCCTACTACATCAGCAAAC---------------GGAAACGGGATGAGTGGCTGGCACGTTGGAAAA------------------------GGGAGGCTGAGAAGAAAGCCAAGG-TAATTGCAGTAATGAATGCTGTGGAAGAGAAC--CAGGCATCTGGAGAGTCTCAGAAGGTGGAGGAGGCC-AGCCCCCCTGCTGTGCAGCAGCCCACGGACCCTGCTTCTCCGACTGTGGCCACCACCCCTGAGCCAGTAGGAGGGGATGCTGGGGACAAGAATGCTACCAAAGCAGCCGACGATGAGCCCGAGTATGAGGATGGCCGGGGCTTTGGCATTGGAGAGCTGGTGTGGGGGAAACTTCGGGGCTTCTCCTGGTGGCCAGGCCGAATTGTGTCTTGGTGGATGACAGGCCGGAGCCGAGCAGCTGAAGGCACTCGCTGGGTCATGTGGTTTGGAGATGGCAAGTTCTCAGTGGTGTGTGTGGAGAAGCTCATGCCGCTGAGCTCCTTCTGCAGTGCATTCCACCAGGCCACCTACAACAAGCAGCCCATGTACCGCAAAGCCATCTATGAAGTCCTCCAGGTGGCCAGCAGCCGT--GCCGGGAAGCTGTTTCCAGCTTGCCATGACAGTGATGAAAGTGACAGTGGCAAGGC----TGTGGAAGTGCAGAACAAGCAGATGATTGAATGGGCCCTCGGTGGC--TTCC---AGCCCTCGGGTCCTAAGAGCCTGGAGCC--ACCAGAAGAAGAGAAGAATCCTTA-----------------------------------------------------CAAGGAAGTTTACACTGACATGTGGGTGGAGCCTGAGGCAGCTGCTTACGCCCCACCCCCACCGGCCAAGAAACCCAGAAA--GAGCACAACAGAGAAACCTAAGGTCAAGGAGATCATTGATGAGCGCACAAGGGAGCGGCTCGTGTATGAGGTTCGCCAGAAGTGCAGAAACATCGAGGACATTTGTATCTCATGTGGGAGCCTCAATGTCACCCTGGAGCACCCACTCTTCATTGGTGGCATGTGCCAGAACTGTAAGA---------ACTGCTTCTTGGAGTGTGCTTACCAGTA-TGACGACGAT-GGGTACCAGTCCTATTGCAC----CATCTGCTGTGGGGGGCGT-----------------GAAGTGCTCATGTGTGGGA----------------------------ACAACAAC--------------TGCTGCAGGTGC-----------------------TTTTGTGTCGAGTGTGTGGATCTCTTGG--TGGGG--CCAGGAGCTGCC-----CAGGCAGCCATTAA------GGAAGACCCCTGGAAC-TGCTACATGTGTGGGCATAAGGGCACCT---ATGGGCTGCTGCGAAGACGGGAAGACTGGCCTTCTCGACTCCAGATGTTCTTTGCCAATAACCATGACCAGGAA------TTTGACCCCCCAAAGGTTTACCCACCTGTGCCAGCTGAGAAGAGGAAGCC------------CATCCGTGTGCTGTCTCTCTTTGA-TGGGATTGCT---ACAGGGCTCCTGGTGCTGAA--GGACCTG---------GGCATCCAAGTGGACCGCTACATTGCCT----CTGAGG--TGTGTGAGGACTCCATCACGGTGGGCATGGTGCGGCACCAGGGAAAGATCATGTACGTCGGGGACGTCCGCAGCGTCACACAGAAGCATATCCAGGAGTGGGGCCCATTCGACCTGGTGATTGGAGGCA---------------------------------------GTCCCTGCAATGACCTCTCCATCGTCAACCCTGCCCGCAAGGGACTTTACGAGGGTACTGGCCGCCTCTTCTTTGAGTTCTACCGCCTCCTGCATGATGCGCGGCCCAAGGAGGGAGATGATCGCCCCTTCTTCTGGCTCTTTGAGAATGTGGTGGCCATGGGCGTTAGTGACAAGAGGGACATCTCGCGATTTCTTGAGTCTAACCCCGTGATGATTGACGCCAAAGAAGTGTCTGCTGCACACAGGGCCCGTTACTTCTGGGGTAACCTTCCTGGCATGAA---CAGGCCTTTGGCATCCACTGTGAATGATAAGCTGGAGCTGCAAGAGTGTCTGGAGCACGGCAGAATAGCCAAGTTCAGCAAAGTGAGGACCATTACCACCAGGTCAAACTCTATAAAGCAGGGCAAAGACCAGCATTTCCCCGTCTTCATGAACGAGAAGGAGGACATCCTGTGGTGCACTGAAATGGAAAGGGTGTTTGGCTTCCCCGTCCACTACACAGACGTCTCCAACATGAGCCGCTTGGCGAGGCAGAGACTGCTGGGCCGATCGTGGAGCGTGCCGGTCATCCGCCACCTCTTCGCTCCGCTGAAGGAATATTTTGCTTGTGTGTAA*

*>Pahari3A*

*ATGCCCTCCAGCGGCCCCGGAGACACCAGCAGCTCCTCTCTGGAGCGGGAGGATGATCGAAAGGAAGGAGAGGAACAGGAGGAGAACCGTGGCAAAGAAGAGCGCCAGGAGCCCAGCGCCACGGCCCGGAAGGTGGGGAGGCCTGGCCGGAAGCGCAAGCACCCACCGGTGGAAAGCAGTGACACCCCCAAGGACCCAGCGGTGACCACCAAGTCTCAGCCCATGGCCCAGGACTCTGGCCCCTCAGATCTGCTACCCAATGGAGACTTGGAGAAGCGGAATGAACC---CCAACCTGAGGAGGGTAGCCCAGCTGCAGGGCAGAAGGGTGGGGCCCCAGCTGAGGGAGAGGG---AACTGAGACCCCACCAGAAGCCTCCAGAGCTGTGGAGAATGGCTGCTGTGTGACCAAGGAGGGCCGTGGAGCCTCTGCAGGAGAGGGCAAAGAACAGAAGCAGACCAACATCGAATCCATGAAAATGGAGG--------GCTCCCGGGGCCGACTGCGCGGTGGCTTGGGCTGGGAGTCCAGCCTCCGTCAGAGGCCCATGCCAAGACTCACCTTCCAGGCGGGGGACCCCTATTACATCAGCAAAC---------------GGAAACGGGATGAGTGGCTGGCACGTTGGAAAA------------------------GGGAGGCTGAGAAGAAAGCCAAGG-TGATTGCAGTAATGAATGCCGTGGAAGAGAAC--CAGGCCTCTGGAGAGTCTCAGAAGGTGGAGGAGGCC-AGCCCTCCTGCTGTGCAGCAGCCCACGGACCCTGCTTCTCCTACTGTGGCCACCACCCCTGAGCCAGTAGGGGGGGATGCTGGGGACAAGAATGCTACCAAAGCAGCTGATGATGAGCCCGAGTATGAGGATGGCCGGGGCTTTGGCATTGGAGAGCTGGTGTGGGGGAAACTTCGGGGCTTCTCCTGGTGGCCAGGCCGAATTGTGTCTTGGTGGATGACAGGCCGGAGCCGAGCAGCTGAAGGCACTCGCTGGGTCATGTGGTTCGGAGATGGCAAGTTCTCAGTGGTGTGTGTGGAGAAGCTCATGCCGCTGAGCTCCTTCTGCAGTGCGTTCCACCAGGCCACCTACAACAAGCAGCCCATGTACCGCAAAGCCATCTACGAAGTCCTCCAGGTGGCCAGCAGCCGC--GCCGGGAAGCTGTTTCCAGCTTGCCATGACAGTGATGAAAGTGACAGTGGCAAGGC----TGTGGAGGTGCAGAACAAGCAGATGATCGAATGGGCCCTCAGCGGC--TTCC---AGCCGTCAGGTCCTAAGGGCCTGGAGCC--ACCAGAAGAGGAGAAGAATCCTTA-----------------------------------------------------CAAGGAAGTTTACACCGACATGTGGGTTGAGCCTGAGGCAGCTGCTTATGCCCCACCCCCACCAGCCAAGAAACCCAGAAA--GAGCACAACAGAGAAGCCAAAGGTCAAGGAGATCATTGATGAACGCACAAGGGAGCGGCTCGTGTATGAGGTGCGTCAGAAGTGCAGAAACATTGAGGACATTTGTATCTCATGTGGGAGCCTCAATGTCACCCTGGAGCACCCACTCTTCATTGGTGGCATGTGCCAGAACTGTAAGA---------ACTGCTTCTTGGAGTGTGCTTACCAGTA-TGACGACGAT-GGGTACCAGTCCTATTGTAC----CATCTGCTGTGGGGGGCGT-----------------GAAGTGCTCATGTGTGGGA----------------------------ACAACAAC--------------TGCTGCAGGTGC-----------------------TTTTGTGTCGAGTGCGTGGATCTCTTGG--TGGGG--CCAGGCGCTGCC-----CAGGCAGCTATTAA------GGAAGACCCCTGGAAC-TGCTACATGTGTGGGCATAAGGGCACCT---ATGGGCTGCTGCGAAGACGGGAAGACTGGCCTTCTCGACTTCAGATGTTCTTTGCCAATAACCATGACCAGGAA------TTTGACCCCCCCAAGGTTTACCCACCTGTGCCAGCTGAGAAAAGGAAACC------------CATCCGCGTGCTGTCTCTCTTTGA-TGGGATTGCT---ACAGGGCTCCTGGTGCTGAA--GGACTTG---------GGCATCCAAGTGGACCGCTACATTGCCT----CCGAGG--TGTGTGAGGACTCCATCACAGTGGGCATGGTGCGGCACCAGGGAAAGATCATGTACGTCGGGGACGTCCGCAGCGTCACACAGAAGCATATCCAGGAGTGGGGTCCATTTGACCTGGTGATTGGAGGCA---------------------------------------GTCCCTGCAATGACCTCTCCATTGTCAACCCTGCCCGCAAGGGACTTTACGAGGGCACTGGCCGCCTCTTTTTTGAGTTCTACCGCCTCCTGCATGATGCGCGGCCCAAGGAGGGAGATGATCGCCCCTTCTTCTGGCTCTTTGAGAATGTGGTGGCCATGGGCGTTAGTGACAAGAGGGACATCTCGCGATTTCTTGAGTCTAACCCCGTGATGATCGACGCCAAAGAAGTGTCTGCAGCACACAGGGCCCGGTACTTCTGGGGTAACCTTCCCGGCATGAA---CAGGCCTTTGGCATCCACTGTGAATGATAAGCTGGAGCTGCAGGAGTGTCTGGAGCACGGCAGAGTAGCCAAGTTCAGCAAAGTGAGGACCATTACCACCAGGTCAAACTCCATAAAGCAGGGCAAAGACCAGCATTTCCCCGTCTTCATGAACGAGAAGGAGGACATCCTGTGGTGCACTGAAATGGAAAGGGTGTTTGGCTTCCCCGTCCACTATACAGATGTCTCCAACATGAGCCGCTTGGCGAGGCAGAGACTGCTGGGCCGATCGTGGAGCGTGCCAGTCATCCGCCATCTCTTCGCTCCGCTGAAGGAATATTTTGCTTGTGTGTAA*

*>Rat3A*

*ATGCCCTCCAGCGGCCCCGGGGACACCAGCATCTCCTCTCTGGAGCGGGAGGATGATCGAAAGGAAGGAGAGGAACAGGAGGAGAACCGTGGTAAGGAAGAGCGTCAGGAGCCCAGCGCCACGGCCCGGAAAGTGGGGAGGCCTGGCCGGAAGCGCAAGCACCCACCGGTGGAAAGCAGTGACACCCCCAAGGACCCAGCGGTGACCACCAAGTCTCAGCCCACAGCCCAGGACTCTGGGCCCTCAGATCTGCTACCCAATGGAGACTTGGAGAAGCGGAGTGAACC---CCAACCTGAGGAGGGGAGCCCAGCTGCAGGGCAGAAGGGTGGGGCCCCAGCTGAAGGAGAGGG---AACTGAGACCCCACCAGAAGCCTCCAGAGCAGTGGAGAATGGCTGCTGCGTAACCAAGGAAGGCCGTGGAGCCTCTGCGGGAGAGGGCAAAGAACAGAAGCAGACCAACATCGAATCCATGAAAATGGAGG--------GCTCCCGGGGCCGACTGCGTGGTGGCTTGGGGTGGGAGTCCAGCCTCCGTCAGCGGCCCATGCCAAGACTCACCTTCCAGGCAGGGGACCCCTACTACATCAGCAAAC---------------GGAAGCGGGATGAGTGGCTGGCACGTTGGAAAA------------------------GGGAGGCTGAGAAGAAAGCCAAGG-TGATTGCAGTAATGAATGCTGTGGAGGAAAGC--CAGGCCTCTGGGGAGTCCCAGAAGGTGGAGGAGGCC-AGCCCTCCTGCAGTGCAGCAGCCCACGGACCCTGCATCTCCCACTGTGGCCACCACCCCTGAGCCAGTAGGGGCTGATGCTGGGGACAAGAATGCTACCAAGGCAGCTGATGACGAGCCCGAGTATGAGGATGGCCGGGGCTTTGGCATTGGAGAGCTGGTGTGGGGGAAACTCCGGGGCTTCTCCTGGTGGCCAGGCCGAATTGTGTCTTGGTGGATGACAGGCCGGAGCCGAGCAGCCGAAGGCACTCGCTGGGTCATGTGGTTCGGAGATGGCAAATTCTCAGTGGTGTGTGTCGAGAAGCTCATGCCCCTGAGCTCCTTCTGCAGTGCGTTCCACCAGGCCACCTACAACAAGCAGCCCATGTACCGCAAAGCCATCTACGAAGTCCTCCAGGTGGCCAGCAGCCGT--GCAGGGAAGCTGTTCCCAGCATGTCATGACAGCGATGAAAGTGACACTGGCAAGGC----TGTGGAGGTGCAGAACAAGCAGATGATTGAGTGGGCCCTTGGCGGC--TTCC---AGCCCTCTGGTCCCAAAGGCCTGGAGCC--ACCAGAAGAGGAGAAGAATCCTTA-----------------------------------------------------CAAGGAAGTTTACACCGACATGTGGGTGGAGCCTGAGGCAGCTGCTTATGCCCCACCCCCACCAGCCAAGAAACCCAGAAA--GAGCACAACAGAGAAACCCAAGGTCAAGGAGATCATTGATGAACGCACAAGAGAACGGCTCGTGTATGAGGTGCGCCAGAAGTGCCGAAACATCGAGGACATTTGTATCTCATGTGGGAGCCTCAATGTTACCCTGGAGCACCCACTCTTCATTGGTGGAATGTGCCAGAACTGTAAGA---------ACTGCTTCTTGGAGTGTGCTTACCAATA-CGATGACGAT-GGGTACCAGTCCTACTGTAC----CATCTGCTGTGGGGGGCGC-----------------GAAGTGCTCATGTGTGGGA----------------------------ACAACAAC--------------TGCTGCAGGTGC-----------------------TTTTGTGTGGAGTGTGTGGATCTCTTGG--TGGGG--CCAGGGGCTGCC-----CAAGCAGCCATTAA------GGAAGACCCCTGGAAC-TGCTACATGTGTGGGCACAAGGGCACGT---ATGGGCTGCTGCGGAGACGGGAGGACTGGCCTTCTCGACTCCAGATGTTCTTCGCCAATAACCACGACCAGGAA------TTTGACCCCCCGAAGGTTTACCCACCTGTGCCAGCTGAGAAGAGGAAGCC------------CATCCGGGTGCTATCTCTCTTTGA-TGGGATTGCT---ACAGGGCTCCTGGTGCTGAA--GGACCTG---------GGCATCCAAGTGGACCGCTACATCGCCT----CCGAGG--TGTGCGAGGACTCCATCACAGTGGGCATGGTGCGGCACCAGGGAAAGATCATGTACGTCGGGGACGTCCGCAGCGTCACACAGAAGCATATCCAGGAGTGGGGCCCATTCGATCTGGTGATTGGGGGCA---------------------------------------GTCCCTGCAATGACCTCTCCATCGTCAACCCTGCCCGCAAGGGACTTTACGAGGGCACGGGCCGCCTCTTCTTTGAGTTCTACCGCCTCCTGCATGACGCGCGGCCCAAGGAGGGAGATGATCGCCCCTTCTTCTGGCTTTTTGAGAATGTGGTGGCCATGGGCGTTAGTGACAAGAGGGACATCTCGAGATTTCTTGAGTCTAACCCCGTGATGATCGACGCCAAAGAAGTGTCTGCTGCACACAGGGCCCGTTATTTCTGGGGTAACCTTCCTGGCATGAA---CAGGCCATTGGCATCCACTGTGAATGATAAGCTGGAGTTGCAAGAGTGTCTGGAACACGGCAGAATAGCCAAGTTCAGCAAAGTGAGGACCATTACCACCAGGTCAAACTCCATAAAGCAGGGCAAAGACCAGCATTTCCCTGTCTTCATGAATGAGAAGGAAGACATCCTCTGGTGCACTGAAATGGAAAGGGTGTTTGGCTTCCCTGTCCACTACACAGACGTCTCCAACATGAGCCGCTTGGCGAGGCAGAGACTGCTGGGCCGATCGTGGAGCGTGCCAGTCATCCGCCACCTCTTCGCTCCGCTGAAGGAATATTTTGCTTGTGTGTAA*

*>DeerMouse3A*

*ATGCCCTCCAGCGGCCCCGGGGACACCAGCAGCTCCGCTCTGGAGCGGGAGGATGATCGGAAGGAAGGAGAGGAACAGGAGGAGAATCGTGGCAAGGAGGAGCGCCAGGAGCCCAGCACCACGGCCCGGAAGGTGGGGAGGCCTGGCCGGAAGCGCAAGCACCCACCGGTGGAAACCAGTGACACCCCCAAGGACCCCGCAGTGACCAGCAAGTCTCAGCCCATGGCCCAGGACTCTGGCTCCTCAGATCTGTTACCCAATGGAGACTTGGAAAAGCGGAGTGAACC---CCAGCCTGAGGAGGGGAGCCCGGCCGCAGGGCAGAAGGGTGGGGCCCCAGCTGAAGGAGAGGG---AACGGAGACCCCGCCGGAATCCTCCCGAGCCGTGGAGAATGGCTGCTGTACCACCAAGGAAGGCCGCGGAGCCTCTGCGGAAGAGGGCAAAGAACAGAAGCAGACCAACATCGAATCCATGAAAATGGAGG--------GATCCCGTGGACGGCTGCGGGGTGGTCTGGGCTGGGAGTCCAGCCTCCGTCAGCGGCCCATGCCAAGACTCACCTTCCAGGCAGGGGACCCCTACTACATCAGCAAGC---------------GGAAACGGGACGAGTGGCTGGCACGTTGGAAAA------------------------GGGAGGCTGAGAAGAAAGCCAAGG-TCATTGCAGTAATGAATGCGGTGGAAGAAAAC--CAGGCCTCTGGAGAGCCTCAGAAGGTAGAGGAGGCT-AGCCCTCCTGCTGTGCAGCAGCCCACAGACCCTGCGTCCCCTACTGTGGCCACCACCCCTGAGCCAGTGGGGGCTGATGCTGGGGACAAGAATGCCACCAAAGCAGCCGACGATGAGCCCGAGTATGAGGACGGCCGTGGCTTTGGCATTGGAGAGCTGGTGTGGGGGAAGCTCCGGGGCTTCTCCTGGTGGCCAGGCCGGATCGTGTCTTGGTGGATGACAGGCCGGAGCCGGGCCGCTGAAGGCACTCGCTGGGTCATGTGGTTCGGAGATGGCAAGTTCTCGGTGGTGTGCGTCGAGAAGCTCATGCCCCTGAGCTCCTTCTGCAGCGCCTTCCACCAGGCCACCTACAACAAGCAGCCCATGTACCGCAAAGCCATCTACGAAGTCCTCCAGGTGGCCAGCAGCCGT--GCTGGGAAGCTGTTTCCGGCCTGCCACGACAGTGACGAAAGTGACACCGGGAAGGC----TGTGGAGGTGCAGAACAAGCAGATGATCGAATGGGCCCTCGGGGGC--TTCC---AGCCCTCTGGTCCCAAGGGCCTAGAGCC--ACCGGAAGAGGAGAAGAATCCATA-----------------------------------------------------CAAGGAGGTCTACACGGACATGTGGGTTGAGCCCGAGGCGGCTGCGTATGCCCCTCCCCCACCAGCCAAGAAACCCAGAAA--GAGCACAGCAGAAAAGCCTAAGGTCAAGGAGATCATTGACGAGCGCACAAGAGAGCGGCTGGTGTATGAGGTGCGGCAGAAGTGTCGGAACATCGAGGACATTTGTATCTCATGTGGAAGCCTCAATGTCACCCTGGAGCACCCACTCTTCATTGGTGGAATGTGCCAGAACTGTAAGA---------ACTGCTTCTTGGAGTGCGCCTACCAGTA-TGACGATGAT-GGGTACCAATCGTACTGCAC----CATCTGCTGTGGGGGGCGT-----------------GAAGTGCTCATGTGTGGGA----------------------------ACAATAAC--------------TGCTGCAGGTGC-----------------------TTTTGTGTCGAGTGTGTGGATCTCCTGG--TGGGG--CCAGGAGCTGCC-----CAGGCGGCCATTAA------GGAAGACCCCTGGAAC-TGCTATATGTGTGGGCACAAGGGCACCT---ATGGACTGCTGCGGAGACGGGAAGACTGGCCTTCCAGGCTCCAGATGTTCTTTGCCAATAACCATGACCAGGAA------TTTGACCCCCCAAAGGTTTACCCACCTGTTCCGGCTGAGAAGAGGAAGCC------------TATCCGGGTGCTATCTCTCTTTGA-TGGAATTGCT---ACAGGGCTCCTGGTGCTAAA--GGACCTG---------GGCATCCAAGTGGACCGCTACATCGCTT----CAGAGG--TGTGTGAGGACTCCATCACTGTGGGCATGGTGCGGCACCAAGGAAAGATCATGTACGTCGGGGACGTCCGCAGCGTCACACAGAAGCATATCCAGGAGTGGGGCCCATTCGATCTGGTGATTGGGGGCA---------------------------------------GTCCCTGCAATGACCTCTCCATCGTCAACCCTGCCCGGAAGGGACTTTATGAGGGTACCGGCCGGCTCTTCTTTGAGTTCTACCGCCTCCTCCATGATGCGCGGCCCAAGGAGGGAGATGACCGCCCCTTCTTCTGGCTCTTTGAGAACGTGGTGGCCATGGGCGTGAGCGACAAGAGGGACATCTCACGATTTCTTGAGTCCAACCCCGTGATGATTGACGCCAAAGAAGTGTCTGCTGCACACAGGGCCCGTTACTTCTGGGGTAACCTTCCTGGCATGAA---CAGGCCATTGGCATCCACTGTGAATGATAAGCTGGAGCTGCAAGAGTGTCTGGAACACGGCAGGATAGCCAAGTTCAGCAAAGTGAGGACCATTACCACCAGGTCAAACTCTATAAAGCAGGGCAAAGACCAGCATTTCCCCGTCTTCATGAATGAGAAGGAAGACATCCTCTGGTGCACTGAAATGGAAAGGGTGTTCGGCTTCCCTGTCCACTACACAGACGTCTCCAACATGAGCCGATTGGCGAGGCAGAGACTGCTGGGCCGGTCGTGGAGCGTACCGGTCATCCGCCATCTCTTCGCTCCGCTGAAGGAATATTTTGCTTGTGTGTAA*

*>CHamster3A*

*ATGCCCTCCAGCGGTCCCGGGGACACCAGCAGCTCCACTTTGGAGCGGGAGGATGATCGAAAGGAAGGAGAGGAACAGGAGGAGAGTCGTGGCAAGGAAGAGCGCCAGGAACCCAGCACCACGGCCCGGAAGGTGGGGAGGCCTGGCCGGAAGCGCAAACACCCACCGGTGGAAAGTAGTGACACACCCAAGGACTCTGCCGTGACTAGCAAGTCTCAGCCCATGGCCCAGGACTCTGGCTCCTCAGATCTGTTACCCAATGGAGACTTGGAAAAGCGGAGTGAACC---CCAACCTGAGGAGGGGAGCCCAGCTGCAGGGCAGAAGGGTGGGGCCCCTGCTGAAGGAGAGGG---AACTGAGACCCCACCAGAATCTTCCCGAGCCGTGGAAAATGGCTGCTGCACAACCAAGGAGGGCCGGGGAGCCTCTGCCGAAGAGGGCAAAGAACAGAAGCAGACCAACATTGAATCCATGAAAATGGAGG--------GCTCCCGGGGCAGACTGCGGGGTGGCTTGGGCTGGGAGTCCAGTCTCCGTCAGCGGCCAATGCCAAGACTCACCTTCCAGGCAGGGGACCCCTACTACATCAGCAAAC---------------GGAAACGGGACGAGTGGCTGGCACGTTGGAAAA------------------------GGGAGGCTGAGAAGAAAGCCAAGG-TAATTGCAGTAATGAATGCTGTGGAGGAAAAC--CAGGCCTCTGGAGAGCCTCAGAAGGTAGAGGAGGCC-AGCCCTCCTGCTGTGCAGCAGCCCACCGACCCTGCATCCCCTACTGTGGCCACCACTCCTGAGCCAGTGGGGGCTGATGCTGGGGACAAGAATGCCACCAAAGCAGCTGATGATGAGCCTGAGTATGAGGATGGCCGGGGCTTTGGCATCGGAGAGTTGGTGTGGGGGAAACTTCGAGGCTTCTCCTGGTGGCCAGGCCGAATTGTGTCTTGGTGGATGACAGGCCGGAGCCGAGCCGCGGAAGGCACTCGCTGGGTCATGTGGTTTGGAGATGGCAAGTTCTCAGTGGTGTGTGTCGAGAAACTCATGCCACTGAGCTCCTTCTGCAGTGCGTTCCACCAGGCCACCTACAATAAGCAGCCCATGTACCGCAAAGCTATCTACGAAGTTCTCCAGGTGGCCAGCAGCCGT--GCTGGGAAGCTGTTTCCAGCTTGCCATGACAGTGACGAAAGTGACACTGGCAAGGC----TGTGGAGGTGCAGAACAAGCAGATGATTGAATGGGCCCTTGGAGGG--TTCC---AGCCCTCTGGTCCCAAGGGCCTGGAGCC--ACCAGAAGAGGAGAAGAATCCATA-----------------------------------------------------CAAGGAAGTTTACACAGACATGTGGGTTGAGCCCGAGGCAGCTGCATATGCTCCACCCCCACCAGCCAAGAAACCCAGAAA--GAGCACAACAGAAAAGCCTAAGGTCAAGGAGATCATTGATGAACGCACAAGAGAGCGGCTGGTGTATGAGGTGCGTCAGAAGTGCCGGAACATCGAGGACATTTGTATCTCATGTGGAAGCCTCAATGTCACCCTGGAGCACCCACTCTTCATTGGTGGAATGTGCCAGAACTGTAAGA---------ACTGCTTCTTGGAGTGCGCGTACCAGTA-CGACGATGAC-GGGTACCAGTCCTACTGCAC----CATCTGCTGTGGGGGGCGT-----------------GAGGTGCTCATGTGTGGCA----------------------------ACAACAAC--------------TGTTGCAGGTGC-----------------------TTTTGTGTCGAGTGTGTGGATCTCTTGG--TGGGG--CCAGGAGCTGCC-----CAAGCGGCCATTAA------GGAAGACCCCTGGAAC-TGCTACATGTGTGGCCACAAGGGCACCT---ATGGGCTGCTGCGGAGACGGGAAGACTGGCCTTCCAGGCTCCAGATGTTCTTTGCCAATAACCATGACCAGGAA------TTTGACCCCCCGAAGGTTTACCCACCTGTTCCAGCTGAGAAGAGGAAGCC------------CATCCGGGTGCTGTCTCTCTTTGA-TGGAATTGCT---ACAGGGCTCCTGGTGCTGAA--GGACCTG---------GGCATCCAAGTGGACCGCTACATTGCCT----CAGAGG--TGTGTGAGGACTCCATCACGGTGGGCATGGTGCGGCACCAGGGAAAGATCATGTACGTCGGGGACGTCCGCAGCGTCACACAGAAGCATATCCAGGAGTGGGGCCCATTCGATCTGGTGATTGGGGGCA---------------------------------------GTCCCTGCAATGACCTCTCCATCGTCAACCCTGCCCGAAAGGGACTTTACGAGGGTACTGGCCGGCTCTTCTTTGAGTTCTACCGCCTCCTGCATGATGCTCGACCTAAGGAGGGAGATGACCGCCCCTTCTTCTGGCTCTTTGAGAATGTGGTGGCCATGGGCGTTAGTGACAAGAGGGACATCTCACGATTTCTTGAGTCCAACCCCGTGATGATTGACGCCAAAGAAGTGTCTGCTGCACACAGGGCCCGTTACTTCTGGGGTAACCTTCCTGGCATGAA---CAGGCCATTGGCATCCACTGTGAATGATAAGCTGGAGCTGCAAGAGTGTCTGGAACATGGCAGAATAGCCAAGTTCAGCAAAGTAAGGACCATCACCACCAGGTCAAATTCCATAAAGCAAGGCAAAGACCAGCATTTCCCCGTCTTCATGAACGAGAAGGAGGACATCCTCTGGTGCACTGAAATGGAAAGGGTGTTTGGCTTCCCTGTCCACTACACAGACGTCTCAAACATGAGCCGCTTGGCGAGGCAGAGACTGCTGGGCCGATCGTGGAGCGTGCCAGTCATCCGCCACCTCTTCGCTCCGCTGAAGGAATATTTTGCTTGTGTGTAA*

*>MBMole3A*

*ATGCCCTCCAGCGGCCCCGGGGACAACAGCAGCTCTGCTCCTGAGCGGGAGGAAGACCGGAAGGAAGAAGAG---CAGGAGGAGAATCGTGGCAAAGAGGAACGCCAGGAGCCCAGTGCCACGGCCCGGAAGGTGGGGAGGCCTGGCCGGAAGCGCAAGCACCCACTGGTGGAAAGCAGCGACACACCCAAGGACCCTGCTGTGACCTCCAAGTCCCCACCCATGGCCCAGGACTCAGGCTCTTCAGAGCTGTTACCCAATGGAGACTTGGAGAAGCGGAGTGAGCC---CCAGCCTGAGGAGGGGAGTCCTGCTGCAGGGCAAAAAGGTGGGGCCCCAGCTGAGGGAGAGGGTGCAGCTGAGACCCCACCAGAGTCCTCCAGAGCTGTGGAGAATGGCTGCTGTGTACCCAAGGAGGGCCGAGGAGCCTCTGCAGAGGAAGGTAAAGAACAGAAGGAGACCAACATCGAATCCATGAAAATGGAGG--------GCTCCCGGGGCCGACTCCGTGGTGGCTTGGGTTGGGAGTCCAGCCTCCGCCAGCGGCCCATGCCAAGGCTCACCTTCCAGGCGGGGGATCCCTACTACATCAGCAAAC---------------GCAAGCGGGATGAGTGGCTGGCACGCTGGAAAA------------------------GGGAGGCTGAGAAGAAAGCCAAGG-TAATTGCAGTAATGAATGCTGTGGAAGAGAAC--CAGGGCTCTGGGGAGCCTCAGAAGGTGGAGGAGGCC-AGCCCTCCTACTGTGCAGCAGCCTACAGACCCTGCATCCCCTACTGTGGCCACCACACCTGAGCCTGTGGGGGCTGATGCTGGGGACAAGAATGCCACCAAAGCAGCTGATGATGAGCCAGAGTATGAGGACGGCCGGGGTTTTGGCATTGGGGAGCTGGTGTGGGGGAAACTGCGAGGCTTCTCCTGGTGGCCAGGCCGAATTGTGTCTTGGTGGATGACGGGCCGGAGCCGAGCAGCTGAAGGCACCCGCTGGGTCATGTGGTTTGGCGACGGCAAGTTCTCAGTGGTGTGTGTGGAGAAGCTGATGCCCCTGAGCTCCTTCTGCAGTGCATTCCACCAGGCAACCTACAACAAGCAGCCCATGTACCGCAAAGCCATCTATGAAGTACTCCAGGTGGCCAGCAGCCGT--GCAGGGAAGCTGTTCCCGGCCTGTCATGACAGTGATGAAAGTGACACTGGCAAGGC----TGTGGAGGTGCAGAACAAGCAGATGATCGAATGGGCCCTTGGGGGG--TTCC---AGCCCTCTGGTCCCAAGGGCCTGGAGCC--ACCAGAAGAAGAGAAGAATCCCTA-----------------------------------------------------TAAGGAAGTTTACACAGACATGTGGGTTGAGCCTGAGGCAGCTGCCTATGCACCACCCCCACCAGCCAAAAAGCCCAGAAA--GAGCACAACTGAGAAGCCCAAGGTCAAGGAGATTATCGATGAACGCACAAGAGAGCGGCTGGTGTACGAGGTGCGGCAGAAGTGCCGAAACATCGAAGACATTTGTATCTCATGTGGAAGCCTCAACGTCACCCTGGAGCACCCACTTTTTATTGGTGGAATGTGCCAGAACTGCAAGA---------ACTGCTTCTTGGAGTGTGCCTACCAATA-TGATGATGAT-GGGTACCAGTCCTATTGCAC----CATCTGCTGTGGGGGCCGT-----------------GAGGTGCTCATGTGCGGGA----------------------------ACAACAAC--------------TGCTGCAGGTGC-----------------------TTTTGTGTTGAGTGTGTAGATCTCTTGG--TGGGG--CCAGGAGCTGCC-----CAAGCAGCCATTAA------GGAAGACCCCTGGAAC-TGCTACATGTGTGGGCACAAGGGCACCT---ATGGGCTGCTGCGGCGGCGAGAAGACTGGCCTTCTCGGCTCCAGATGTTCTTCGCCAATAATCATGACCAGGAA------TTTGACCCCCCAAAAGTTTACCCACCTGTCCCTGCCGAGAAAAGGAAGCC------------TATCCGGGTGCTATCTCTCTTTGA-TGGAATTGCT---ACAGGGCTCCTGGTGCTAAA--GGACCTG---------GGCATCCAAGTGGACCGCTACATTGCCT----CAGAGG--TGTGTGAGGACTCCATCACGGTGGGAATGGTGCGGCACCAGGGGAAGATCATGTACGTCGGGGACGTCCGCAGCGTCACACAGAAGCATATCCAGGAGTGGGGCCCATTTGATCTGGTGATTGGGGGCA---------------------------------------GTCCCTGCAATGACCTCTCTATTGTGAACCCTGCCCGCAAGGGACTTTACGAGGGAACTGGACGGCTCTTCTTTGAGTTCTACCGCCTCCTGCATGATGCGCGGCCCAAGGAGGGAGATGATCGCCCCTTCTTCTGGCTCTTTGAGAACGTGGTGGCCATGGGCGTTAGTGACAAGAGGGACATCTCACGATTTCTCGAGTCCAACCCTGTGATGATTGATGCCAAAGAAGTGTCAGCTGCACACAGGGCCCGCTACTTCTGGGGCAACCTTCCCGGTATGAA---CAGGCCATTGGCATCCACTGTGAATGATAAGCTGGAGCTGCAGGAGTGTCTGGAACATGGCAGAATAGCCAAGTTCAGTAAAGTGAGGACCATTACCACCAGGTCGAACTCCATAAAGCAAGGCAAAGACCAGCATTTCCCTGTCTTCATGAATGAGAAGGAGGACATCCTATGGTGCACTGAAATGGAAAGGGTGTTTGGCTTCCCTGTCCACTACACAGACGTCTCCAACATGAGCCGCTTGGCAAGGCAGAGACTGCTGGGCCGGTCATGGAGTGTGCCAGTCATCCGCCACCTCTTCGCTCCGCTGAAGGAATATTTTGCTTGTGTGTAA*

*>Marmot_3A_(Incomplete)*

*-------------------------------------------------------------------------------------------------------------------------------------------------------------------------------------------------------------------------------------------------------------------------------------------------------------------------------------------------------------------------------------------------------------------------------------------------------------------------------------------------------------------------------------------------------------------------------------CACCT---------GGCGCCCTCAGGGGA-CAGGAAGA---------------TCACCCCGGGCAGTGGGCTCC----------------------------------------GGCTGAGAAGAAAGCCAAGG-TCATTGCAGTAATGAATGCTGTGGAAGAGAAC--CAGGGCTCCGGGGAGGCTCAGAAGGTGGAGGAGGCC-AGCCCGCCTGCTGTGCAGCAGCCCACTGACCCAGCATCTCCCACTGTGGCCACCACACCGGAGCCCGTGGGGGCTGACGCTGGGGACAAGAATGCCACCAAATCAGCCGATGATGAGCCAGAGTATGAGGACGGCCGGGGCTTTGGCATTGGGGAGCTGGTGTGGGGGAAACTGCGGGGCTTCTCCTGGTGGCCAGGCCGCATTGTGTCTTGGTGGATGACGGGCCGGAGCCGTGCAGCTGAAGGCACCCGCTGGGTCATGTGGTTCGGAGACGGCAAGTTCTCAGTGGTGTGTGTAGAGAAGCTGATGCCGCTGAGCTCGTTCTGCAGCGCCTTCCACCAGGCCACCTACAACAAGCAGCCCATGTACCGCAAAGCCATCTACGAAGTCCTCCAGGTGGCCAGTAGCCGT--GCCGGGAAGCTGTTCCCAGCCTGCCATGACAGTGACGAGAGTGACACTGCCAAGGC----TGTGGAGGTGCAGAACAAGCAGATGATTGAATGGGCCCTCGGGGGG--TTCC---AGCCCTCTGGCCCGAAGGGCCTAGAGCC--ACCAGAAGAGGAGAAAAATCCCTA-----------------------------------------------------CAAGGAAGTTTACACAGACATGTGGGTTGAACCCGAGGCAGCTGCCTATGCACCACCCCCACCAGCCAAAAAGCCGCGAAA--GAGCACAACTGAGAAGCCCAAGGTCAAGGAGATTATTGATGAACGCACAAGAGAGCGGCTGGTGTATGAGGTGCGGCAGAAATGCCGGAACATTGAGGATATTTGCATCTCTTGTGGGAGCCTCAATGTTACCCTGGAGCATCCTCTGTTCATTGGTGGAATGTGCCAGAACTGCAAGA---------ACTGCTTCCTGGAGTGTGCGTACCAGTA-TGATGATGAC-GGTTATCAGTCCTATTGCAC----CATCTGCTGTGGGGGCCGT-----------------GAGGTGCTCATGTGTGGGA----------------------------ACAACAAC--------------TGCTGTAGGTGT-----------------------TTTTGTGTGGAGTGCGTGGATCTCTTGG--TGGGG--CCGGGGGCTGCC-----CAAGCAGCCATTAA------GGAAGACCCCTGGAAC-TGCTACATGTGCGGGCACAAGGGCACTT---ATGGGCTGCTGCGGCGGCGGGACGACTGGCCCTCACGGCTTCAGATGTTCTTCGCCAATAACCACGACCAGGAA------TTTGATCCTCCGAAGGTTTACCCACCTGTCCCAGCCGAGAAAAGGAAGCC------------CATCCGGGTGCTGTCTCTCTTTGA-TGGAATTGCT---ACAGGGCTCCTGGTGCTGAA--GGACTTG---------GGCATCCAGGTGGACCGCTACATAGCTT----CGGAGG--TGTGTGAGGACTCCATCACAGTGGGCATGGTGCGACATCAGGGGAAGATCATGTACGTCGGGGACGTCCGCAGTGTCACACAGAAGCATATCCAGGAGTGGGGCCCATTCGATCTGGTGATTGGGGGCA---------------------------------------GTCCCTGCAATGATCTCTCAATCGTCAACCCTGCCCGCAAGGGACTCTACGAGGGCACTGGCCGGCTCTTCTTTGAGTTCTACCGCCTCCTGCATGATGCGCGGCCCAAGGAGGGAGATGACCGCCCCTTCTTCTGGCTCTTTGAGAATGTGGTGGCCATGGGCGTTAGTGACAAGAGGGACATCTCGCGATTTCTCGAGTCCAACCCTGTGATGATTGATGCCAAAGAAGTGTCAGCCGCACATAGGGCCCGCTACTTCTGGGGTAACCTTCCCGGTATGAA---CAGGCCATTGGCATCCACTGTGAATGATAAGCTGGAGCTGCAGGAGTGTCTGGAACATGGCAGAATAGCCAAGTTCAGCAAAGTGAGGACCATTACTACTAGGTCAAACTCCATAAAACAGGGCAAAGACCAGCATTTCCCCGTCTTCATGAATGAGAAAGAGGACATCCTATGGTGCACTGAAATGGAAAGGGTTTTTGGCTTCCCTGTCCACTATACTGACGTCTCCAACATGAGCCGCTTGGCGAGGCAGAGACTGCTGGGCCGGTCGTGGAGCGTGCCAGTCATCCGCCACCTCTTCGCTCCGCTGAAGGAATATTTTGCTTGTGTGTAA*

*>GroundSquirrel_3A_(incomplete)*

*----------------------------------------------------------------------------------------------------------------------------------------------------------------------------------------------------------------------------------------------------------------------------------------------------------------------------------------------------------------------------------CCGAAGCCTCCAGAGCAGTGGAGAATGGCTGCTGTGCACCCAAGGAGGGCCGGGGAGCCTCTGCAGAAGAGGGCAAAGAACAGAAGGAGACCAACATCGAATCCATGAAAATGGAGG--------GCTCCCGGGGCCGGCTGCGGGGTGGCTTGGGCTGGGAGTCCAGCCTCCGCCAGCGGCCCATGCCGCGGCTCACCTTCCAGGCTGGGGACCCCTACTACATCAGCAAGC---------------GCAAGCGGGACGAGTGGCTGGCACGCTGGAAAA------------------------GGGAGGCTGAGAAGAAAGCCAAGG-TCATTGCAGTAATGAATGCTGTGGAAGAGAAC--CAGGGCTCCGGGGAGGCTCAGAAGGTGGAGGAGGCC-AGCCCTCCTGCTGTGCAGCAGCCCACTGACCCAGCATCTCCCACTGTGGCCACCACACCGGAGCCTGTGGGGGCTGATGCTGGGGACAAGAATGCCACCAAATCAGCTGATGATGAGCCAGAGTATGAGGACGGCCGGGGCTTTGGCATTGGGGAGCTGGTGTGGGGGAAACTGCGGGGCTTCTCCTGGTGGCCAGGCCGCATTGTGTCTTGGTGGATGACGGGCCGGAGCCGTGCAGCTGAAGGCACCCGCTGGGTCATGTGGTTCGGAGACGGCAAGTTCTCAGTGGTGTGTGTAGAGAAGCTGATGCCACTGAGCTCGTTCTGCAGCGCCTTCCACCAGGCCACCTACAACAAGCAGCCCATGTACCGCAAAGCCATCTACGAAGTCCTCCAGGTGGCCAGTAGCCGT--GCCGGGAAGCTGTTTCCAGCCTGCCATGACAGTGACGAGAGTGACACTGCCAAGGC----TGTGGAGGTGCAGAACAAGCAGATGATTGAATGGGCCCTCGGGGGT--TTCC---AGCCCTCTGGCCCGAAGGGCCTAGAGCC--ACCAGAAGAGGAGAAAAATCCCTA-----------------------------------------------------CAAGGAAGTTTACACAGACATGTGGGTTGAACCCGAGGCAGCTGCCTATGCACCACCCCCACCAGCCAAAAAGCCACGAAA--GAGCACAACTGAGAAGCCCAAGGTCAAGGAGATTATTGATGAACGCACAAGAGAGCGGCTGGTGTATGAGGTGCGGCAGAAATGCCGGAACATTGAGGATATTTGCATCTCTTGTGGGAGCCTCAATGTCACCCTGGAGCATCCTCTGTTCATTGGTGGAATGTGCCAGAACTGCAAGA---------ACTGCTTCCTGGAGTGTGCGTACCAGTA-TGACGATGAC-GGTTATCAGTCTTATTGTAC----CATCTGCTGTGGGGGCCGT-----------------GAGGTGCTCATGTGTGGGA----------------------------ACAACAAC--------------TGCTGTAGGTGT-----------------------TTTTGTGTGGAGTGCGTGGATCTCTTGG--TGGGG--CCGGGGGCTGCC-----CAAGCAGCCATTAA------GGAAGACCCCTGGAAC-TGCTACATGTGCGGGCACAAGGGCACTT---ATGGGCTTCTGCGGCGGCGGGACGACTGGCCCTCACGGCTTCAGATGTTCTTCGCCAATAACCACGACCAGGAA------TTTGATCCTCCGAAGGTTTACCCACCTGTCCCAGCCGAGAAAAGGAAGCC------------CATCCGGGTGCTGTCTCTCTTTGA-TGGAATTGCT---ACAGGGCTCCTGGTGCTGAA--GGACTTG---------GGCATCCAGGTGGACCGCTACATAGCTT----CGGAGG--TGTGTGAGGACTCCATCACAGTGGGCATGGTGCGACATCAGGGGAAGATCATGTACGTCGGGGACGTCCGCAGTGTCACACAGAAGCATATCCAGGAGTGGGGCCCATTTGATCTGGTGATTGGGGGCA---------------------------------------GTCCCTGCAATGATCTCTCAATCGTCAATCCTGCCCGCAAGGGACTCTACGAGGGCACTGGCCGGCTCTTCTTTGAGTTCTACCGCCTCCTGCATGATGCGCGGCCCAAGGAGGGAGATGACCGCCCCTTCTTCTGGCTCTTTGAGAATGTGGTGGCCATGGGCGTTAGTGACAAGAGGGACATCTCGCGATTTCTCGAGTCCAACCCTGTGATGATTGATGCCAAAGAAGTGTCAGCTGCACATAGGGCCCGCTACTTCTGGGGTAACCTTCCCGGTATGAA---CAGGCCATTGGCATCCACTGTGAATGATAAGCTGGAGCTGCAGGAGTGTCTGGAACATGGCAGAATAGCCAAGTTCAGCAAAGTGAGGACCATTACTACTAGGTCAAACTCCATAAAACAGGGCAAAGACCAGCATTTCCCCGTCTTCATGAATGAGAAAGAGGACATCCTATGGTGCACTGAAATGGAAAGGGTTTTTGGCTTCCCTGTCCACTATACTGACGTCTCCAACATGAGCCGCTTGGCGAGGCAGAGACTGCTGGGCCGGTCGTGGAGCGTGCCAGTCATCCGCCACCTCTTCGCTCCGCTGAAGGAATATTTTGCTTGTGTGTAA*

*>Rabbit_3A*

*ATGCCCTCCAGCGGCCCCGGGGACACCAGCAGCTCCGCTCCGGAGCGGGACGAGGACCGAAAGGAAGGAGAAGAGCAGGAGGAGCCTCGTGGCAAAGAGGAGCGGCAGGAGCCCAGCACCACCGCCCGGAAAGTGGGCAGGCCCGGGAGGAAGCGCAAGCACCCGCCGGTGGAAAGCAGCGACACGCCCAAGGACCCTGCGGTCACCTCCAAGTCCCCGTCCATGGCCCAGGACTCAGGCCCCTCCGAGCTGTTACCCAACGGGGACTTGGAGAAGCGGAGTGAGCC---CCAGCCGGAGGAGGGGAGCCCTGCTGGGGGGCAGAAGGGCGGGGCCCCAGCTGAGGGCGAGGGTGCAGCCGAGGCCCCGCCCGAAGCCTCCAGAGCGGTGGAGAATGGCTGCTGCACCCCGAAGGAGGGCCGAGGAGCCCCTGCAGGCGAGGGCAAAGAACAGAAGGAGACCAATGTGGAATCCATGAAAATGGAGG--------GCTCCCGGGGCCGGCTGCGGGGCGGCCTGGGCTGGGAGTCCAGCCTCCGCCAGCGGCCCATGCCGCGGCTCACCTTCCAGGCGGGCGACCCCTACTACATCAGCAAAC---------------GCAAGCGGGACGAGTGGCTGGCACGCTGGAAAA------------------------GGGAGGCTGAGAAGAAAGCCAAGG-TCATTGCCGTGATGAATGCTGTAGAAGAGAGC--CAGGGGTCCGGGGAGCCCCAGAAGGTGGAGGAGGCC-AGCCCTCCTGCTGTGCAGCAGCCCACCGACCCCGCCTCCCCCACTGTGGCCACTACGCCCGAGCCTGTGGGGGCTGATGCCGGGGACAAGAACGCCACCAAAGCAGCCGACGACGAGCCGGAGTACGAGGACGGCCGGGGCTTTGGCATTGGGGAGCTGGTGTGGGGGAAACTGCGGGGCTTCTCCTGGTGGCCAGGCCGCATAGTGTCCTGGTGGATGACGGGCCGGAGCCGTGCAGCCGAGGGGACCCGCTGGGTCATGTGGTTCGGAGACGGCAAGTTCTCAGTGGTGTGCGTGGAGAAGCTGATGCCGCTGAGTTCGTTCTGCAGCGCCTTCCACCAGGCCACCTACAACAAGCAGCCCATGTACCGCAAGGCCATCTACGAAGTCCTGCAGGTGGCCAGCAGCCGC--GCGGGCAAGCTCTTCCCGGCCTGCCATGACAGTGACGAGAGCGATACTGCCAAGGC----CGTGGAGGTGCAGAACAAGCAGATGATCGAGTGGGCCCTTGGGGGA--TTCC---AGCCCTCGGGCCCCAAGGGCCTGGAGCC--ACCAGAAGAGGAGAAGAACCCCTA-----------------------------------------------------CAAAGAAGTTTACACGGACATGTGGGTGGAACCCGAGGCAGCCGCCTACGCCCCACCCCCACCAGCCAAAAAACCACGGAA--GAGCACGACGGAGAAGCCCAAGGTCAAGGAGATCATTGATGAGCGCACACGAGAGCGGCTGGTGTACGAGGTGCGGCAAAAGTGCCGGAACATCGAGGACATCTGCATCTCGTGTGGGAGCCTCAATGTCACCCTGGAACACCCCCTCTTCATTGGAGGAATGTGCCAAAATTGCAAGA---------ACTGTTTCCTGGAATGCGCCTACCAGTA-CGACGATGAT-GGCTATCAGTCATACTGCAC----CATCTGCTGCGGGGGGCGC-----------------GAGGTGCTCATGTGTGGGA----------------------------ACAACAAC--------------TGCTGCCGGTGC-----------------------TTCTGTGTGGAGTGTGTGGACCTCTTGG--TGGGG--CCGGGGGCTGCA-----CAGGCGGCCATTAA------GGAAGACCCCTGGAAC-TGCTACATGTGTGGCCACAAGGGCACCT---ACGGGCTGCTGCGGCGTCGGGACGACTGGCCCTCTCGGCTCCAGATGTTCTTCGCCAATAACCACGACCAGGAA------TTCGACCCTCCGAAGGTTTACCCGCCTGTCCCAGCTGAAAAGAGGAAGCC------------CATCCGGGTGCTGTCTCTCTTTGA-TGGAATTGCT---ACAGGGCTCCTGGTGCTGAA--GGACTTG---------GGCATCCAGGTGGACCGCTACATCGCCT----CAGAGG--TGTGCGAGGACTCCATCACGGTGGGCATGGTGCGGCACCAGGGGAAGATCATGTACGTGGGGGACGTCCGCAGCGTCACACAGAAGCACATCCAGGAATGGGGCCCATTTGACCTTGTGATCGGGGGCA---------------------------------------GCCCCTGCAACGACCTCTCCATTGTCAACCCTGCCCGCAAGGGGCTGTATGAGGGCACTGGCCGGCTCTTCTTTGAGTTCTACCGCCTCCTGCATGACGCGCGGCCCAAGGAGGGAGATGATCGCCCCTTCTTCTGGCTCTTTGAGAATGTGGTGGCCATGGGCGTTAGTGACAAGAGGGACATCTCACGATTTCTCGAGTCCAACCCCGTCATGATTGATGCCAAAGAAGTGTCGGCCGCACACAGGGCCCGCTACTTCTGGGGTAACCTTCCCGGTATGAA---CAGGCCGTTGGCATCCACTGTGAATGATAAGCTGGAGCTGCAGGAGTGTCTGGAGCATGGCAGGATAGCCAAGTTCAGCAAAGTGAGGACCATCACTACCAGGTCAAACTCCATAAAGCAGGGCAAAGACCAGCACTTCCCCGTCTTCATGAACGAGAAGGAGGACATCCTGTGGTGCACTGAGATGGAAAGGGTGTTTGGCTTTCCTGTGCACTATACCGACGTCTCCAACATGAGCCGCTTGGCGAGACAGAGGCTGCTGGGCCGGTCGTGGAGCGTGCCCGTGATCCGCCACCTCTTCGCTCCGCTGAAGGAATACTTTGCTTGTGTGTAA*

*>Marmot_3B*

*------------------------------------------------------------------------------------------------------------------------------------ATGAAGGGGGATACCAGACACCTCAGTGA------AGAGGAGGGCGCCAGCGGGAGGGAGGACTCCATCATTCTCATCAATGGGGCCAGCAGTGACCAGTCCTCCGACTCCAAGGACGCCCCTTCGCCCCCCATCCTGGAG--GCTATCCGC---------------TCACCTGAGATCAG-----AGGCCGCAGATCCAGCTCCCGCCTGTCCAAGAGGG--------------------AGGTCTCCAGCCTAATCAGCTACACTCAGGATCTGACAGGGGAAGCAGATGGTGA------------AG---AAGATGGGGACAGCTCTGAGGCTCCTGTGATGCCAAGACTA---------TTCCGGGAAACCA---------GGACCCGCTCTGCAAGCCCAGCAGTCCGAACCCGAAATAGCAACAACAGTGCCTCCAGTCGGGAGAGGTACAAGTCCTTCCCACA---------TCCCACCCGAGGCCGCCAGGGCCGCAGTCACGTGGACGAGTCTCCCGTGGAGTTCCCGGCTACCAGGTCCCTAAGGCGGCGGTCAACA---GCCTCACCAG-GTACGCCGTGGCCATCCCCGTCCACTCCTTACCTCACCATCGACCTCACGGAGGACGATGTGACACCACAGAGCAGCAGT------ACCCCTT-GTGCCAGCCTAGCTGAG---AGCC-AGCAGGAGAACTCGGAGACCTCCCAGGTGGACGCAGAGGGGAGAGACTCTGACAACACTGAGTACCAGGATGGGAAGGAGTTTGGAATAGGGGACCTGGTATGGGGCAAGATCAAGGGCTTCTCCTGGTGGCCTGCCATGGTGGTTTCTTGGAAGGCTACCTCCAAGCGCCAGGCCATGTCCGGCATGCGATGGGTCCAGTGGTTTGGTGATGGCAAGTTCTCTGAGGTCTCTGCAGACAAACTGGTGGCCTTGGGGCTGTTCAGTCAGCATTTTAACCAGGCCACCTTCAATAAACTGGTGTCATACAGGAAGGCCATGTACCAGGCCCTGGAGAAAGCCAGGGTACGA--GCTGGCAAGACCTTCCCCA----GCA-----------------GCCCTGGAGACTC----TTTGGAGGACCAGCTGAAGCCCATGTTGGAGTGGGCCCACGGGGGC--TTCA---AGCCCACTGGGATCGAGGGCCTCAAACCCAACAACAAC---CAACCAGTGGTTAATAAATCGAAGGTGCGTCGTGCAGGCAGTAGGAACTTAGAATCAAGGAAACACGAGAACAAGAGTCGAAGACGCACAACT----GATGATTCAGCTGCCTCTGACCACTGTCCCCCAAAC-AAGCGCCTCAAGACAAACTGCTATAACAATGGCAAGGACCGTGGAGAGGAA-GATCAG---AGCCGAGAACAAATGGTTTCTGATGTTACCAACAACAAGAACAATCTGGAAGATAGCTGTTTGTCCTGTGGTAGAAAAAACCCCGTGTCCTTCCACCCGCTCTTTGAGGGGGGGCTTTGCCAGACGTGCCGGG---------ATCGCTTCCTTGAGTTGTTCTACATGTA-CGACGACGAT-GGTTATCAGTCCTACTGCAC----CGTGTGCTGTGAGGGCCGT-----------------GAGCTGCTTCTCTGCAGCA----------------------------ACACAAGC--------------TGTTGCCGGTGC-----------------------TTCTGTGTGGAGTGTCTGGAGGTGCTGG--TGGGC--AAGGGCACAGCG-----GAGGACGC--CAAG----CTGCAGGAGCCCTGGAGC-TGCTACATGTGTCTCCCCCAGCGCTGCC---ACGGAGTCCTGCGGCGCCGGAAGGACTGGAATGTCCGCCTGCAGACCTTCTTCACCAGCGACATGGGGCTTGA---ATAC---GAGGCCCCCAAGTTGTACCCTGCGATTCCCGCAGCCAAAAGGCGGCC------------CATTAGAGTCCTGTCACTATTCGA-TGGAATTGCC---ACAGGTTACTTGGTCCTCAA--AGACTTG---------GGCATTAAAGTGGAAAAGTATGTCGCCT----CAGAAG--TGTGCGAAGAATCCATCGCCGTTGGAACCGTGAAGCACGAGGGAAACATCAAATATGTGAATGACGTCAGGAACATCACAAAGAAAAATATTGAAGAGTGGGGTCCATTTGATTTGGTGATCGGTGGAA---------------------------------------GCCCATGCAATGACCTATCAAATGTGAATCCTGCCCGGAAAGGCCTGTATGAGGGCACTGGCCGACTCTTCTTCGAATTCTACCACCTGCTCAATTACTCACGCCCCAAAGAAGGGGATGACCGGCCATTCTTCTGGATGTTTGAGAATGTTGTAGCCATGAAGGTCGGCGACAAAAGGGACATCTCACGGTTCCTGGAGTGTAACCCAGTGATGATTGATGCCATCAAAGTGTCTGCTGCTCACAGGGCTAGATACTTCTGGGGAAACCTACCTGGGATGAA---CAGGCCTGTGATAGCATCAAAGAATGATAAGCTCGAGCTGCAGGACTGCTTGGAATTCAGTAGGACTGCAAAGTTAAAGAAAGTGCAGACAATAACCACCAAGTCCAACTCCATCAAACAGGGGAAAAACCAACTTTTCCCTGTTGTCATGAATGGCAAAGAAGATGTTTTGTGGTGCACTGAGCTCGAAAGGATCTTCGGTTTTCCTGTGCACTACACGGATGTGTCCAACATGGGCCGCGGTGCCCGCCAGAAGCTGCTTGGGCGGTCCTGGAGTGTGCCCGTCATCCGACACCTCTTCGCCCCCCTGAAGGACTACTTTGCATGTGAATAG*

*>Squirel_3B*

*------------------------------------------------------------------------------------------------------------------------------------ATGAAGGGGGACACCAGACACCTCAGTGA------AGAGGAGGGCGCCAGCGGGAGGGAGGACTCCATCATTCTCATCAACGGGGCCAGCAGTGACCAATCCTCCGACTCCAAGGACGCCCCTTCGCCCCCCATCCTGGAG--GCTATCCGC---------------TCGCCCGAGATCAG-----AGGCCGCAGATCCAGCTCCCGCCTGTCCAAGAGGG--------------------AGGTCTCCAGCCTAATCAGCTACACTCAGGATCTGACAGGGGAAGCAGATGGTGA------------AGGAGAAGATGGGGACAACTCCGAGGCTCCTGTGATGCCAAGACTA---------TTCCGGGAAACCA---------GGACCCGCTCTGCAAGCCCAGCAGTCCGAACCCGAAATAGCAACAACAGTGCTTCCA---------------AGTCCTTCCCACA---------TCCCACCCGAGGCCGCCAGGGCCGCAGTCATGTGGACGAGTCTCCCGTGGAGTTCCCGGCTACCAGGTCCCTAAGGCGGCGGTCAACAACAGCCTCGCCAG-GTACGCCATGGCCATCCCCATCCACTCCTTACCTCACCATCGACCTCACGGAGGACGATGTGACACCACAGAGCAGCAGT------ACCCCCT-GTGCCAGCCTAGCTGAG---AGCC-AGCAGGAGAACTCGGAGATCTCCCAGGTGGACGCAGAGGGGAGAGACTCCGACAACACTGAGTACCAGGATGGGAAGGAGTTTGGAATAGGGGACCTGGTATGGGGCAAGATCAAGGGCTTCTCCTGGTGGCCTGCCATGGTGGTTTCTTGGAAGGCTACCTCCAAGCGCCAGGCCATGTCCGGCATGCGATGGGTCCAGTGGTTTGGTGATGGCAAGTTCTCTGAGGTCTCTGCAGACAAACTGGTGGCCTTGGGGCTGTTCAGTCAGCATTTTAACCAGGCCACCTTCAATAAACTGGTGTCCTACAGGAAGGCCATGTACCAGGCCCTGGAGAAAGCCAGGGTACGA--GCTGGCAAGACCTTCCCCA----GCA-----------------GCCCTGGAGACTC----TTTGGAGGACCAGCTGAAGCCCATGTTGGAGTGGGCCCACGGGGGC--TTCA---AGCCCACTGGGATCGAGGGCCTCAAACCCAACAACAAC---CAACCAG------------------------------------------------------------AGAACAAGAGTCGAAGACGCACAGTT----GATGATTCAGCCGCCTCTGACCACTGTCCTCCAAAC-AAGCGCCTCAAGACAAACTGCTATAACAATGGCAAGGACCGGGGAGAGGAA-GATCAG---AGCCGAGAACAAATGGTTTCTGATGTTGCCAACAACAAGAACAATCTGGAAGATAGCTGTTTGTCCTGTGGTAGGAAAAACCCAGTGTCCTTCCATCCCCTCTTTGAGGGTGGGCTTTGCCAGACGTGCCGGG---------ATCGCTTCCTTGAGCTCTTCTACATGTA-CGACGATGAT-GGTTACCAGTCCTACTGCAC----CGTGTGCTGTGAGGGCCGT-----------------GAGCTGCTTCTCTGCAGCA----------------------------ATACAAGC--------------TGTTGCCGGTGC-----------------------TTCTGTGTGGAGTGTCTGGAGGTGCTGG--TGGGC--AAGGGCACAGCG-----GAGGACGC--CAAG----CTGCAGGAGCCCTGGAGC-TGCTACATGTGTCTCCCCCAGCGCTGCC---ACGGAGTCCTGCGGCGCCGGAAGGACTGGAATGTCCGCCTGCAGACCTTCTTCACCAGCGACCCGGGGCTCGA---ATAT---GAGGCCCCCAAGTTGTACCCTGCGATTCCCGCAGCCAAAAGGCGGCC------------CATTAGAGTCCTGTCACTATTCGA-TGGAATTGCC---ACAGGTTACTTGGTCCTCAA--AGACTTG---------GGCATTAAAGTGGAAAAGTATGTTGCCT----CGGAAG--TGTGCGAAGAATCCATCGCCGTTGGAACCGTGAAGCACGAGGGCAACATCAAATATGTGAATGACGTTAGGAACATCACAAAGAAAAATATTGAAGAGTGGGGTCCATTTGATTTGGTGATCGGTGGAA---------------------------------------GCCCGTGCAATGACCTGTCAAATGTGAATCCTGCCCGGAAAGGCCTGTATGAGGGCACTGGCCGACTCTTCTTCGAATTCTACCACCTGCTCAATTACTCACGCCCCAAAGAAGGGGATGACCGGCCGTTCTTCTGGATGTTTGAGAATGTTGTAGCCATGAAGGTCGATGACAAGAGGGACATCTCACGGTTCTTGGAGTGCAACCCAGTGATGATTGATGCTATCAAAGTGTCTGCTGCTCACAGGGCTCGATACTTCTGGGGAAACCTACCTGGGATGAA---CAGGCCTGTGATAGCATCAAAGAATGATAAGCTCGAGCTGCAGGACTGCTTGGAATTCAGTAGGACTGCAAAGTTAAAGAAAGTGCAGACAATAACTACCAAGTCCAACTCCATCAAACAGGGGAAAAACCAACTTTTCCCTGTTGTCATGAATGGCAAAGAAGATGTTTTGTGGTGCACTGAGCTCGAAAGGATCTTCGGTTTTCCTGTGCACTACACGGATGTGTCCAACATGGGCCGCGGTGCCCGCCAGAAGCTGCTTGGGCGGTCCTGGAGTGTGCCAGTCATTCGACACCTCTTCGCCCCCCTGAAGGACTACTTTGCCTGTGAATAG*

*>Rabbit_3B*

*------------------------------------------------------------------------------------------------------------------------------------ATGAAGGGAGACACCAGACACCTCAATGG------AGAGGAGGACGCCAGCGGGAGGGAAGACTCAATTGTCCTCGTCAACGGGGCCTGCAGCGACCAGTCCTCTGATTCCAAGGACGCCCCCTCGCCCCCCATCCTGGAG--GCTATCCGC---------------ACCCCGGAGATCAG-----AGGCCGCCGATCGAGCTCCCGACTGTCCCGGAGGG--------------------AGGTGTCCAGCCTGCTGAGTTACACCCAGGATCTGACGGGCGATGGAGAT---GA------------AGGCGAGGACGGGGATGGGTCAGACACTCCAGTGATGCCTAAACTG---------TTCCGTGAGACCA---------GAACGCGCTCTGAAAGCCCAGCAGTCCGAACCCGAAATAGCAACA---GTGCCTCCAGCCGGGAGAGGTACAGGCCCTCCCCACGGTCCACCCGTTCCACCCGAGGCCGGCAGGGCCACAGTCACGTGGACGAGTCCCCCGTGGAGTTCCCAGCAACCAGGTCCCTGAGGCGCAGGGCCACA---GCATCCACAG-GCACGCCGTGGCCATCCCCTGCCAGTCCCTACCTCACCATCGACCTCACTGATGACGACGTGACACCCCAGAGTAGCAGT------ACCCCCT-ACACTGGCCTGGCCCAGGACAGCC-AGCTGGAGAGCCTGGAGTCCTCACAGGTGGATGGAGA---CAGAGATGCGGATGGAGCGGAGTACCAGGATGGGAAGGAGTTTGGAATAGGGGACCTCGTGTGGGGAAAGATCAAGGGCTTCTCCTGGTGGCCAGCCATGGTGGTGTCCTGGAAGGCCACCTCCAAGCGCCAGGCCATGGCCGGCATGCGATGGGTGCAATGGTTTGGCGATGGAAAATTCTCTGAGGTCTCTGCAGACAAGCTTGTTGCTCTGGGGTTGTTCAGCCAGCACTTCAACCTTGCAACGTTTAATAAACTGGTCTCCTATAGGAAGGCCATGTTCCAGGCTCTGGAGAAAGCCAGGGTCCGT--GCTGGCAAGACCTTCTCCA----CCA-----------------GCCCTGGAGACCC----GCTGGAGGACCAGCTGAAGCCCATGCTGGAGTGGGCCCACGGGGGC--TTCA---AGCCCACCGGCACCGAGGGCCTCAAGCCCGACAACAGCAAGCAACCAGTGGTTAATAAGTCGAAGGTGCGCCGTGCAGGCAATAGGAACTTAGAAGCACGGAGAAACGAGAATAAGAGTCGGAGACGCACGGCC----GAGGACTCCACTGCCTCAGAGCACTGCCCCCCACCC-AAGCGCCTCAAGACCAACTGCTACAACAACGGCAAAGACCGAGGAGAGGAA-GATCAG---AGCCGAGAGCAAATGGCTGCCGATGTGACCAACAACAAGAGCAACCTCGAAGACAGCTGCTTGTCCTGCGGGAGGAAAAACCCCGTGTCCTTCCACCCCCTCTTTGAGGGTGGCCTCTGCCAGACCTGCCGGG---------ATCGCTTTCTCGAGCTGTTCTACATGTA-TGATGATGAC-GGCTATCAGTCATACTGCAC----CGTGTGCTGCGAGGGCCGG-----------------GAGCTGCTACTGTGTAGCA----------------------------ACACGAGC--------------TGCTGCCGGTGC-----------------------TTCTGCGTGGAGTGCCTGGAGGTGCTGG--TGGGC--AGCGGCACCGCA-----GCCGACGC--TAAA----CTACAGGAGCCCTGGAGC-TGCTACATGTGCGTCCCGCAGCGCTGCC---ATGGGGTCCTGCGGCGCCGCAAGGACTGGAACGTGCGCCTGCAGGCCTTCTTCACCAGCGACATGGGGCACGA---ATAC---GAAGCCCCCAAGCTGTACCCTGCCATTCCTGCAGCCCGTAGGCGGCC------------CATTCGAGTCCTGTCCTTGTTCGA-TGGCATTGCG---ACAGGGTACTTGGTCCTCAA--AGAATTG---------GGCATAAAGGTGGAAAAGTACGTCGCTT----CAGAAG--TGTGTGAAGAATCCATCGCAGTGGGGACCGTGAAGCACGAGGGCAACATCAAATATGTGAATGACGTCAGGAACATCACGAAGAAAAACATTGATGAATGGGGGCCCTTTGACTTGGTGATTGGGGGCA---------------------------------------GCCCGTGCAACGACCTCTCCAATGTGAATCCTGCCCGGAAAGGCCTGTATGAAGGCACAGGCCGGCTCTTCTTTGAGTTCTACCACCTGCTGAATTACACCCGGCCCAAGGAGGGTGACGACCGGCCCTTCTTCTGGATGTTTGAGAATGTTGTAGCCATGAAGGTTGGAGACAAGAGGGACATCTCAAGGTTCCTGGAGTGTAACCCGGTGATGATCGATGCCATCAAAGTTTCTGCTGCTCACAGGGCCCGGTACTTTTGGGGCAACCTGCCTGGGATGAA---CAGGCCCGTGATAGCATCAAAGAATGATAAGCTCGAGCTGCAGGACTGCTTGGAATACAGTAGGACAGCAAAGTTAAAGAAAGTACAGACAATAACCACCAAGTCGAACTCGATCAAACAGGGGAAAAACCAACTTTTCCCTGTTGTCATGAATGGCAAAGAAGATGTTTTGTGGTGCACTGAGCTCGAAAGGATCTTCGGCTTTCCCGTACACTACACGGACGTGTCCAACATGGGCCGCGGCGCCCGCCAGAAGCTGCTCGGGAGGTCGTGGAGCGTGCCCGTCATCCGACACCTCTTCGCCCCGCTGAAGGACTACTTTGCCTGTGAATAG*

*>BlindMountainMole_3B*

*------------------------------------------------------------------------------------------------------------------------------------ATGAAGGGAGACAGCAGACACCTCAATGA------GGAAGAAGGTGCCAGTGGGTGTGAGGACTCCATCATT---GTCAACGGAACCTGCAGTGACCAGTCCTCAGACACCAAGGATGCTCCCTCACCCCCAGTCTTGGAG--GCAATCTGC---------------ACACCTGAGAGCAG-----AGGCCGCAGATCAAGCTCACGATTGTCCAAGAGGG--------------------AGGTCTCCAGCCTGCTAAGTTACACTCAGGACCTGACTGGAGATGGAGATGGAGATGAT---GAAGTAGAGGAGGATGGGGATGGCACTGAAATTCCAGTCATGCCAAAACTC---------ACTCGGGAGACCA---------GGATACCCTCTGAAAGCCCAGCTGTGCGAACCCGAAATAGCAATA---GTACCTCCAGCCGGGAGAGGCGCAGAGCCTCCCGGAGA---------GTCACCCGGGGCCGGCGGGGCCACCACCAAGTGCACGATTCCCCTGTGGAGTTTCCAGCTACCAGGTCCATGAGGCGGAGGGCAAGA---ACATCAGCAG-GCACGCCGTGGCCATCCCCATCCAGCCCCTACCCCACCATCGACCTCACAGATGAAGAAGTGACACCTGGGAGCAGCAGG------ACCCCCT-CCGTTGACCTAGGCCAGGACAGCC-AGCAGGAGAGCATGGACTCCACACAGGTGGATACAGAAAGCAGGGATGGAGACAGCACTGAGTATCAGGACGGAAAGGAGTTTGGAATAGGCGACCTTGTGTGGGGAAAGATCAAGGGCTTCTCCTGGTGGCCTGCCATGGTGGTTTCCTGGAGGGCCACCTCTAAGCGCCAGGCCATGCCAGGCATGAGATGGGTACAGTGGTTTGGTGATGGCAAGTTCTCTGAGGTCTCTGCAGACAAACTGGTGGCCTTGGGGCTGTTCAGTCAGCACTTTAATCTGGCCACCTTCAATAAACTGGTTTCTTACAGGAAGGCTATGTACCACACTCTGGAGAAAGCCAGGGTTCGA--GCTGGCAAGACCTTCCCCA----CCA-----------------GCCCTGCAGACTC----ACTGGAGGACCAGCTGAAGCCCATGTTGGAGTGGGCCCACGGAGGC--TTCA---AGCCCACTGGGATTGAGGGCCTCAAACCCAACAACAAG---CAACCAGTGGTTAATAAGTCGAAGGTGCGTCGTACAGGCAGTAAGGACTTAGAATCCAGGAAACACGAGAGCAAAAGTCGAAGACGCACAACA----GATGACTCTGCAACTTCTGACTATTGTCCCCTACCC-AAGCGCCTCAAGACAAACAGCTGT---GGTGGCAAGGACCGTGGGGAGGAT-GATCAG---AGCCGAGAGCAAATGGCTTCCGATGTTACCAACAACAAGAGCAACCTGGAAGACAGCTGTTTGTCCTGTGGGAGGAAAAACCCTGTGTCCTTCCACCCACTCTTTGAGGGTGGGCTCTGCCAGACATGCCGGG---------ATCGCTTCCTCGAGCTGTTTTACATGTA-CGATGATGAT-GGCTATCAGTCATACTGCAC----CGTGTGCTGTGAGGGCCGT-----------------GAGCTACTTCTGTGCAGCA----------------------------ACACAAGC--------------TGCTGCCGGTGC-----------------------TTCTGTGTTGAGTGTCTGGAGGTGCTGG--TGGGC--ACAGGCACAGCT-----GAAGAGGC--CAAA----ATGCAGGAACCCTGGAGC-TGCTACATGTGTCTCCCTCAGCGCTGCC---ATGGTGTCCTCCGACGCAGAAAGGACTGGAACACACGCCTGCAGGATTTCTTTACCACTGTTCCCGATCTGGAGGAATTT---GAGCCTCCCAAGTTGTATCCAGCTATTCCTGCAGCCCGACGACGGCC------------AATTAGAGTTCTGTCCCTGTTTGA-TGGAATTGCG---ACAGGCTACTTGGTCCTCAA--AGAATTG---------GGTATTAAAGTGGACAGGTACGTCGCCT----CTGAAG--TCTGTGCAGAATCCATCGCTGTGGGAACTGTGAAGCATGAAGGGCAAATCAAATACGTGAATGACGTCAGGAAAATCACAAAGAGAAATATTGAAGAATGGGGCCCCTTCGACTTGGTGATTGGTGGAA---------------------------------------GCCCGTGCAATGATCTCTCTAATGTCAATCCTGCCAGGAAAGGACTGTATGAGGGTACTGGCCGACTCTTTTTTGAGTTTTACCACTTGCTGAATTATACCCGTCCCAAAGAGGGTGACAACCGTCCGTTTTTCTGGATGTTTGAGAATGTGGTTGCCATGAAGGTTAATGACAAGAAGGACATCTCACGGTTCCTGGCGTGTAACCCAGTGATGATTGATGCCATCAAGGTTTCTGCGGCTCACAGGGCTCGATACTTCTGGGGCAACCTACCTGGGATGAA---CAGGCCCGTGATAGCATCAAAGAATGATAAACTCGAGCTGCAGGACTGCTTGGAGTTCAGTAGGACAGCAAAGTTGAAGAAAGTACAGACAATAACCACCAAGTCGAACTCGATCAGACAGGGGAAAAACCAACTTTTCCCTGTAGTCATGAATGGCAAAGAAGATGTTTTGTGGTGCACTGAGCTCGAAAGGATCTTCGGCTTTCCTGTACACTACACAGACGTGTCCAACATGGGTCGTAGTGCCCGCCAGAAGCTGCTGGGAAGGTCCTGGAGTGTGCCTGTCATCAGACATCTCTTTGCCCCCTTGAAGGACTACTTTGCATGTGAATAG*

*>Mouse_3B*

*------------------------------------------------------------------------------------------------------------------------------------ATGAAGGGAGACAGCAGACATCTGAATGA------AGAAGAGGGTGCCAGCGGGTATGAGGAGTGCATTATC---GTTAATGGGAACTTCAGTGACCAGTCCTCAGACACGAAGGATGCTCCCTCACCCCCAGTCTTGGAG--GCAATCTGCACAGAGCCAGTCTGCACACCAGAGACCAG-----AGGCCGCAGGTCAAGCTCCCGGCTGTCTAAGAGGG--------------------AGGTCTCCAGCCTTCTGAATTACACGCAGGACATGACAGGAGATGGAGACAGAGATGA------TGAAGTAGATGATGGGAATGGCTCTGATATTCT---AATGCCAAAGCTC---------ACCCGTGAGACCAAGGACACCAGGACGCGCTCTGAAAGCCCGGCTGTCCGAACCCGACATAGCAATG---GGACCTCCAGCTTGGAGAGGCAAAGAGCCTCCCCCAGA---------ATCACCCGAGGTCGGCAGGGCCGCCACCATGTGCAGGAGTACCCTGTGGAGTTTCCGGCTACCAGGTCTCGGAGACGTCGAGCATCA---TCTTCAGCAA-GCACGCCATGGTCATCCCCTGCCAGCGTCGACT------------TCATGGAAGA---AGTGACACCTAAGAGCGTCAGT------ACCCCAT-CAGTTGACTTGAGCCAGGATGGAG-ATCAGGAGGGTATGGATACCACACAGGTGGATGCAGAGAGCAGAGATGGAGACAGCACAGAGTATCAGGATGATAAAGAGTTTGGAATAGGTGACCTCGTGTGGGGAAAGATCAAGGGCTTCTCCTGGTGGCCTGCCATGGTGGTGTCCTGGAAAGCCACCTCCAAGCGACAGGCCATGCCCGGAATGCGCTGGGTACAGTGGTTTGGTGATGGCAAGTTTTCTGAGATCTCTGCTGACAAACTGGTGGCTCTGGGGCTGTTCAGCCAGCACTTTAATCTGGCTACCTTCAATAAGCTGGTTTCTTATAGGAAGGCCATGTACCACACTCTGGAGAAAGCCAGGGTTCGA--GCTGGCAAGACCTTCTCCA----GCA-----------------GTCCTGGAGAGTC----ACTGGAGGACCAGCTGAAGCCCATGCTGGAGTGGGCCCACGGTGGC--TTCA---AGCCTACTGGGATCGAGGGCCTCAAACCCAACAAGAAG---CAACCAGTGGTTAATAAGTCGAAGGTGCGTCGTTCAGACAGTAGGAACTTAGAACCCAGGAGACGCGAGAACAAAAGTCGAAGACGCACAACC----AATGACTCTGCTGCTTCTGAGT---CCCCCCCACCC-AAGCGCCTCAAGACAAATAGCTAT---GGCGGGAAGGACCGAGGGGAGGAT-GAGGAG---AGCCGAGAACGGATGGCTTCTGAAGTCACCAACAACAAGGGCAATCTGGAAGACCGCTGTTTGTCCTGTGGAAAGAAGAACCCTGTGTCCTTCCACCCCCTCTTTGAGGGTGGGCTCTGTCAGAGTTGCCGGG---------ATCGCTTCCTAGAGCTCTTCTACATGTA-TGATGAGGAC-GGCTATCAGTCCTACTGCAC----CGTGTGCTGTGAGGGCCGT-----------------GAACTGCTGCTGTGCAGTA----------------------------ACACAAGC--------------TGCTGCAGATGC-----------------------TTCTGTGTGGAGTGTCTGGAGGTGCTGG--TGGGC--GCAGGCACAGCT-----GAGGATGC--CAAG----CTGCAGGAACCCTGGAGC-TGCTATATGTGCCTCCCTCAGCGCTGCC---ATGGGGTCCTCCGACGCAGGAAAGATTGGAACATGCGCCTGCAAGACTTCTTCACTACTGATCCTGACCTGGAAGAATTTCAGGAGCCACCCAAGTTGTACCCAGCAATTCCTGCAGCCAAAAGGAGGCC------------CATTAGAGTCCTGTCTCTGTTTGA-TGGAATTGCA---ACGGGGTACTTGGTGCTCAA--GGAGTTG---------GGTATTAAAGTGGAAAAGTACATTGCCT----CCGAAG--TCTGTGCAGAGTCCATCGCTGTGGGAACTGTTAAGCATGAAGGCCAGATCAAATATGTCAATGACGTCCGGAAAATCACCAAGAAAAATATTGAAGAGTGGGGCCCGTTCGACTTGGTGATTGGTGGAA---------------------------------------GCCCATGCAATGATCTCTCTAACGTCAATCCTGCCCGCAAAGGTTTATATGAGGGCACAGGAAGGCTCTTCTTCGAGTTTTACCACTTGCTGAATTATACCCGCCCCAAGGAGGGCGACAACCGTCCATTCTTCTGGATGTTCGAGAATGTTGTGGCCATGAAAGTGAATGACAAGAAAGACATCTCAAGATTCCTGGCATGTAACCCAGTGATGATCGATGCCATCAAGGTGTCTGCTGCTCACAGGGCCCGGTACTTCTGGGGTAACCTACCCGGAATGAA---CAGGCCCGTGATGGCTTCAAAGAATGATAAGCTCGAGCTGCAGGACTGCCTGGAGTTCAGTAGGACAGCAAAGTTAAAGAAAGTGCAGACAATAACCACCAAGTCGAACTCCATCAGACAGGGCAAAAACCAGCTTTTCCCTGTAGTCATGAATGGCAAGGACGACGTTTTGTGGTGCACTGAGCTCGAAAGGATCTTCGGCTTCCCTGCTCACTACACGGACGTGTCCAACATGGGCCGCGGCGCCCGTCAGAAGCTGCTGGGCAGGTCCTGGAGTGTACCGGTCATCAGACACCTGTTTGCCCCCTTGAAGGACTACTTTGCCTGTGAATAG*

*>Caroli_3B*

*------------------------------------------------------------------------------------------------------------------------------------ATGAAGGGAGACAGCAGACATCTGAATGA------AGAAGAGGGAGCCAGCGGATTTGAGGAGTGCATAATT---GTTAATGGGAACTTCAGTGACCAGTCCTCAGACACGAAGGATGCTCCCTCACCCCCAGTCTTGGAG--GCAATCTGCACAGAGCCAGTCTGCACACCAGAGACCAG-----AGGCCGCAGGTCAAGCTCCCGGCTGTCTAAGAGGG--------------------AGGTCTCCAGCCTTCTGAATTACACGCAGGACATGACAGGAGATGGAGATGGAGATGA------TGAAGTGGATGATGGGAATGGCTCTGATATTCT---AATGCCAAAGCTC---------ACCCGTGAGACCAAGGACACCAGGACGATCTCTGAAAGCCCGGCTGTCCGAACCCGACATAGCAATG---GGACCTCCAGCTTGGAGAGGCAAAGAGCCTCCCCCAGA---------ATCACCCGAGGCCGGCAGGGCCGCCACCATGTGCAGGAGTACCCCGTGGAGTTCCCAGCTACCAGGTCTCGGAGACGTCGAGCATCG---TCTTCAGCAA-GCACGCCGTGGTCATCCCCTGCCAGCGTCGACT------------TCATGGAAGA---AGCGACACCTAAGAGCGTCAGT------ACCCCAT-CAGTTGACTTGAGCCAGGATGGAG-ATCAGGAGGGCATAGATACCACACAAGTGGATGCAGAGAGCAGAGATGGAGACAGCACAGAGTATCAGGATGATAAAGAGTTTGGAATAGGTGACCTCGTGTGGGGAAAGATCAAGGGCTTCTCCTGGTGGCCTGCCATGGTGGTGTCCTGGAAAGCCACCTCCAAGCGCCAGGCCATGCCCGGAATGCGCTGGGTACAGTGGTTTGGCGATGGCAAGTTTTCTGAGATCTCTGCTGACAAACTGGTGGCTCTGGGGCTGTTCAGCCAGCACTTTAATCTGGCTACCTTCAATAAGCTGGTTTCTTATCGGAAGGCCATGTACCACACTCTGGAGAAAGCCAGGGTACGA--GCTGGCAAGACCTTCTCCA----GCA-----------------GTCCTGGAGAGTC----ACTGGAGGACCAGCTGAAGCCCATGCTGGAGTGGGCCCACGGCGGC--TTCA---AGCCCACTGGGATCGAGGGCCTCAAACCCAACAACAAG---CAACCAG------------------------------------------------------------AGAACAAAAGTCGAAGGCGCACAGCC----AATGACTCTGCTGCTTCTGAGTATTCCCCCCCACCC-AAGCGCCTCAAGACCAACAGCTAT---GGTGGGAAGGACCGAGGGGAGGAT-GAGGAG---AGCCGAGAACGGATGGCTTCTGATGTCACCAACAACAAGGGCAATCTGGAAGACCGCTGTTTGTCCTGTGGAAAGAAGAACCCTGTGTCCTTCCACCCCCTCTTTGAGGGTGGGCTCTGTCAGAGTTGCCGGG---------ATCGCTTCCTAGAGCTCTTCTACATGTA-TGACGAGGAC-GGCTATCAGTCCTACTGCAC----CGTGTGCTGTGAGGGCCGT-----------------GAACTGCTGCTGTGCAGTA----------------------------ACACAAGT--------------TGCTGCAGATGC-----------------------TTCTGTGTGGAGTGTCTGGAGGTGCTGG--TGGGC--GCAGGCACAGCT-----GAGGATGC--CAAG----CTGCAGGAACCCTGGAGC-TGCTATATGTGCCTCCCTCAGCGCTGCC---ATGGGGTCCTCCGACGCAGGAAGGATTGGAACATGCGCCTGCAAGACTTCTTCACTACTGATCCTGACCTGGAAGAATTTCAGGAGCCGCCCAAGTTGTACCCAGCAATTCCTGCTGCCAAAAGGAGGCC------------CATTAGAGTCCTGTCTCTGTTTGA-TGGAATTGCA---ACAGGGTACTTGGTGCTCAA--GGAGTTG---------GGTATTAAAGTGGAGAAGTACATTGCCT----CCGAAG--TCTGTGCAGAGTCCATCGCTGTGGGAACCATTAAGCATGAAGGCCAAATCAAATATGTCAATGACGTCCGGAAAATCACCAAGAAAAATATTGAAGAGTGGGGCCCGTTCGACTTGGTGATTGGTGGAA---------------------------------------GCCCATGCAACGATCTCTCTAATGTCAATCCTGCCCGCAAAGGTTTATATGAGGGCACGGGAAGGCTCTTCTTCGAGTTTTACCACTTGCTGAATTATACCCGCCCCAAGGAGGGCGACAACCGTCCATTCTTCTGGATGTTCGAGAATGTTGTGGCCATGAAAGTGAATGACAAGAAAGACATCTCCCGATTCCTGGCATGTAACCCAGTGATGATTGATGCCATCAAGGTGTCTGCTGCTCACAGGGCCCGGTACTTCTGGGGTAACCTACCGGGAATGAA---CAGG---------------------------------------------------------------------------------------------------------------------------------------------------------------------------------------------ATCTTCGGCTTCCCTGCTCACTACACGGACGTGTCTAACATGGGCCGCGGCGCCCGTCAGAAGCTGCTGGGCAGGTCCTGGAGTGTGCCGGTCATCAGACACCTGTTTGCCCCCTTGAAGGACTACTTTGCCTGTGAATAG*

*>Spretus_PredictedCDS_(incomplete)*

*------------------------------------------------------------------------------------------------------------------------------------ATGAAGGGAGACAGCAGACATCTGAATGA------AGAAGAGGGTGCCAGCGGGTATGAGGAGTGCATAATC---GTTAGTGGGAACTGCAGTGACCAGTCCTCAGACACGAAGGATGTTCCCTCACCCCCCGTCTTGGAG--GCAATCTGCACAGAGCCAGTCTGCACACCAGAGACCAG-----AGGCCGCAGGTCAAGCTCCCGGCTGTCTAAGAGGG--------------------AGGTCTCCAGCCTTCTGAATTACACGCAGGACATGACAGGAGATGGAGACGGAGATGA------TGAAGTGGATGATGGGAATGGCTCTGATATTCT---AATGCCAAAGCTC---------ACCCGTGAGACCAAGGACACCAGGACGCGCTCTGAAAGCCCAGCTGTCCGAGCCCGACATAGCAATG---GGACCTCCAGCTTGGAGAGGCAAAGAGCCTCCCCCAGA---------ATCACCCGAGGTCGGCAGGGCCGCCACCATGTGCAGGAGTACCCCGTGGAGTTTCCGGCTACCAGGTCTCGGAGACGTCGAGCATCG---TCTTCAGCAA-GCACGCCATGGTCATCTCCTGCCAGTGTTGACT------------TCATGGAAGA---AGTGACACCTAAGAGCGTCAGT------ACCCCAT-CAGTTGACTTGAGTCAGGATGGAG-ATCAGGAGGGCATGGATACCACACAGGTGGATGCAGAGAGCAGAGATGGAGACAGCACAGAGTATCAGGATGATAAAGAGTTTGGAATAGGTGACCTCGTGTGGGGAAAGATCAAGGGCTTCTCCTGGTGGCCTGCCATGGTGGTCTCCTGGAAAGCCACCTCCAAGCGCCAGGCCATGCCCGGAATGCGCTGGGTACAGTGGTTTGGTGATGGCAAGTTTTCTGAGATTTCTGCTGACAAACTGGTGGCTCTGGGGCTGTTCAGCCAGCACTTTAATCTGGCTACCTTCAATAAGCTGGTTTCTTATAGGAAGGCCATGTACCACACTCTGGAGAAAGCCAGGGTTCGA--GCTGGCAAGACCTTCTCCA----GCA-----------------GTCCTGGAGAGTC----ACTGGAGGACCAGCTGAAGCCCATGCTGGAGTGGGCCCACGGTGGC--TTCA---AGCCCACTGGGATCGAGGGCCTCAAACCCAACAAGAAG---CAACCAGT-----------------------------------------------------------GGCTAATAAGTCGAAGGTGCGTCGTT----CA-GAC-------------AGT---------------AGGAACTTAGAACCCAGGAGACGC---GGTATTACCTTCCCA---------------------TCTTTAAGTGGTTTTCTTTGCTCTGTCT---------TGGAATTTT----ACCGCTGTTTGTCGTGTGGAAAGAAGAACCCTGTGTCCTTCCACCCCCTCTTTGAGGGTGGGCTCTGTCAGAGTTGCCGGG---------TAAGTCTTCTCTTGCTCTCCAGCATGCA---------GT-GCTCCCCAGCATT----CAG----TGCTCCCTATCAGG-CAGT-----------------GCTCT-CTATCAGGCAGTG----------------------------CTCCCTAT--------------TAG-GCAG-TGC-----------------------TCTCCTCTCCAGCA---GGCAGTGCTCT--TCAGT--CCTAGCTCTCTC-----TATTTTGT--ATAC----CTGCAGGAATC----AGT-CAACACACACCACACTCTCAGGTCTGCTTCAATGGGGCCT--GAGTTCCCCTGGGGTTGACACAGAT-CCTCTGAGGTT--------ACAGGGATGGACT--GAGGGTCCTCATTGTCCTGCAGGAGGACACCGTGGGGCTGATGGCCCCACACTT-GGCC------------TCTTGGCCTCACATTTCTCTTTGA-TC-----------ACAGGGTACTTGGTGCTCAA--GGAGTTG---------GGTATTAAAGTGGAAAAGTACATTGCCT----CCGAAG--TCTGTGCAGAGTCCATCGCTGTGGGAACTGTTAAGCATGAAGGCCAAATCAAATATGTCAATGACGTCCGGAAAATCACCAAGAAAAATATTGAAGAGTGGGGCCCGTTCGACTTGGTGATTGGTGGAA---------------------------------------GCCCATGCAATGATCTCTCTAACGTCAATCCTGCCCGCAAAGGTTTATATGAGGGCACAGGAAGGCTCTTCTTCGAGTTTTACCACTTGCTGAATTACACCCGCCCCAAGGAGGGCGACAACCGTCCATTCTTCTGGATGTTCGAGAATGTTGTGGCCATGAAAGTGAATGACAAGAAAGACATCTCAAGATTCCTAGCATGTAACCCAGTGATGATCGATGCCATCAAGGTGTCTGCTGCTCACAGGGCCCGGTACTTCTGGGGTAACCTACCCGGAATGAA---CAGGCCCGTGATGGCTTCAAAGAATGATAAGCTCGAGCTGCAGGACTGCCTGGAGTTCAGTAGGACAGCAAAGTTAAAGAAAGTGCAGACAATAACCACCAAGTCGAACTCCATCAGACAGGGCAAAAACCAGCTTTTCCCTGTAGTCATGAATGGCAAGGACGACGTTTTGTGGTGCACTGAGCTCGAAAGGATCTTCGGCTTCCCTGCTCACTACACGGACGTGTCCAACATGGGCCGCGGCGCCCGTCAGAAGCTGCTGGGCAGGTCCTGGAGTGTGCCGGTCATCAGACACCTGTTTGCCCCCTTGAAGGACTACTTTGCCTGTGAA---*

*>Rat_3B*

*------------------------------------------------------------------------------------------------------------------------------------ATGAAGGGAGACAGCAGACATCTTAATGA------AGAAGAGGGTGCTAGTGGGTATGAGGACTGTATCATT---GTTAATGGGAACTGTAGTGACCAGTCCTCGGACACGAAGGATGCTCCCTCACCCCCAGTCTTGGAG--GCAATCTGCACAGAGCCAGTCTGTACACCAGAGACCAG-----AGGCCGCAGATCAAGCTCACGGCTGTCTAAGAGGG--------------------AGGTCTCCAGCCTTCTGAATTACACGCAGGACGTGGTAGGAGATGGAGATGG------------TGAAGCGGATGATGGAGATGGCTCTGATATTCTCATGATGCCAAAGCTC---------ACCCGAGAGACCAAGGATGCCAGGACTCCCTCTGAAAGCCCAGCCGTCCGAACCCGAAACAGCAACA---GTATCTCCAGCCTGGAGAGGCAAAGAACCTCCCCCAGA---------ATCACGCGAGGCCGGCAGGGCCGCTACCACGTTCAGGAGTACCCCGTGGAGTTCCCAGCTACCAAGTCTCGGAGGCGGCGAGCATCG---TCTTCAGCAA-GCACGCCATGGTCATCCCCTGCCAGCATCGAGC------------TCATGGAAGA---TGTGACACCTAAGAGCAGCAGT------ACGCCAT-CGGTTGACTTGAGCCAGGACGGCC-CTCAGGAGGGCATGGATGCCACACAGGTGGATGCGGAGAGCAGGGATGGTGACAGCACTGAGTATCAGGATGACAAGGAGTTTGGAATAGGTGACCTTGTGTGGGGAAAGATCAAGGGCTTCTCCTGGTGGCCTGCCATGGTGGTGTCCTGGAAAGCCACCTCCAAGCGCCAGGCCATGCCCGGAATGCGCTGGGTACAGTGGTTTGGCGATGGCAAGTTTTCTGAGATCGCTGCTGACAAGCTGGTGGCTCTGGGTCTGTTCAGCCAGCACTTTAATCTGGCCACCTTCAATAAGCTGGTTTCTTATAGGAAGGCCATGTACCACACTCTGGAGAAAGCCATGGTGCGA--GCTGGCAAGACCTTCCCCA----GCC-----------------GCCCTGGAGACTC----ACTGGAGGACCAGCTGAAGCCCATGCTGGAGTGGGCCCACGGGGGC--TTCA---AGCCCACTGGGATCGAGGGCCTCAAACCCAACAACAAG---CAACCAGAGGTTCATAAGTCGAAGGTGCGTCGTTCAGGCAGTAGGAACTTAGAAGCCAGGAGACGCGAGAACAAAAGTCGAAGACGCACAACC----ATTGACTTTGCCGCTTCTGAGTACTCCACACCCCCT-AAGCGCCTCAAGACAAATAGCTAT---GGTGGGAAGGACCGAGGGGAAGAT-GAGGAG---AGCCGAGAACGGATGGCTTCTGATGTCACTAACAACAAGGGCAATCTGGAAGACCGCTGTCTGTCCTGCGGTAAGAAGAACCCTGTGTCCTTCCACCCCCTCTTTGAGGGTGGGCTCTGTCAGAGTTGCCGGG---------ATCGCTTCCTGGAGCTCTTCTACATGTA-CGATGAGGAC-GGCTATCAGTCCTACTGCAC----CGTGTGCTGTGAGGGCCGT-----------------GAACTGCTTCTGTGCAGTA----------------------------ACACAAGC--------------TGCTGCAGGTGC-----------------------TTCTGTGTGGAGTGTCTGGAGGTGCTGG--TGGGT--ACAGGCACAGCG-----GAGGATGC--CAAG----CTGCAGGAACCCTGGAGC-TGCTATATGTGCCTCCCGCAGCGCTGCC---ATGGGGTCCTGCGGCGCAGGAAGGACTGGAACATGCGCCTGCAAGACTTCTTCACTACTGATCCTGACCTGGAAGAGTTT---GAGCCGCCCAAGTTGTACCCAGCGATTCCTGCAGCCAAAAGGAGGCC------------CATTAGGGTCCTGTCCCTGTTTGA-TGGAATTGCA---ACAGGGTACTTGGTGCTCAA--GGACTTG---------GGTATTAAAGTGGAGAAGTACGTTGCCT----CTGAAG--TCTGTGCAGAGTCCATTGCTGTAGGAACCATTAAGCATGAAGGCCAAATCAAATATGTCAATGACGTCCGGAAAATCACCAAGAAAAATATTGAAGAGTGGGGCCCATTTGACTTGGTGATTGGCGGAA---------------------------------------GCCCATGCAATGATCTTTCTAATGTCAATCCTGCCAGGAAAGGCCTGTATGAGGGCACAGGAAGGCTCTTCTTTGAGTTTTACCACTTGCTGAATTATACACGCCCCAAGGAGGGCGACAACCGTCCATTCTTCTGGATGTTCGAGAATGTTGTGGCCATGAAAGTGAATGACAAGAAAGACATCTCCCGATTCCTGGCATGTAACCCAGTGATGATCGACGCCATCAAGGTTTCTGCTGCTCACAGGGCCCGGTACTTCTGGGGTAATCTACCTGGAATGAA---CAGGCCTGTGATAGCTTCAAAGAATGATAAGCTCGAGCTACAAGACTGCTTGGAGTTCAGTAGGACAGCAAAGTTAAAGAAAGTACAGACAATAACCACCAAGTCGAACTCCATCAGACAGGGGAAAAACCAGCTTTTCCCTGTAGTCATGAATGGCAAGGATGACGTTCTGTGGTGCACTGAGCTCGAAAGGATCTTCGGGTTCCCAGCTCACTACACAGACGTGTCCAACATGGGCCGGGGCGCCCGCCAGAAGCTACTGGGAAGGTCCTGGAGTGTGCCAGTCATCAGACACCTGTTTGCCCCCTTGAAGGACTACTTTGCATGTGAATAG*

*>FieldVole_GeneWiseORF_3B_(incomplete)*

*------------------------------------------------------------------------------------------------------------------------------------ATGAAGGGAGACAGCAGACATCTGAATGA------AGAAGAGGGTGCCAGTGGGGGTGAGGAATGTGTCATT---GTCAATGGGAACTGTAGTGACCATTCCTCAGACACTAAGGATGCTCCCTCACCCCCAGTCTTGGAG--GCAATCTGCACAGAGGCTGACAGCACACTGGAGAGCAG-----AGGCCGCAGATCAAGATCAAGGCTGTCAAAGAGGG--------------------AGGTCTCCGGCCTGCTGAATTATACTCAGGACCTGACAGGAAATGGAGATGG------------TGAGGCAGAGGATGGGGATGGTGATGATGTTCTACTGATGCCAAAGCTC---------ACACGTGAGACCAAGGAGACCAGGTCACCCTCTGAAAGCCCAGCTATGCGAACCCGAAATAACAACA---GTACCTCCAGGCTAGGGAGGCAAAGAGCCTCCCCCAGA---------ATCACCCGAGGCCGCCAGGGACGACACCATGTGCAGGAATACCCCGTGGAGTTCCCAGCTACCAGGTCTCGGAGGCGTCGGGCATCA---TCGTCAGCAA-GCACGCCATGGTCGTCCCCTGCCAGCCCTTCCT------------TGGTAGAAGA---AGGTACACCTCAGAACAGCAGT------ATCCCAT-CAGTTGACTTGACCCAGGACATCA-ATCAGGAGAGCATGGACACTACACAGCTGGAGGCAGAAGGCAAAGATGGAGACAGCACTGAGTATCAGGATGACAAGGAATTTGGAATAGGTGACCTTGTGTGGGGAAAAATCAAGGGATTTTCCTGGTGGCCTGCTATGGTGGTGTCCTGGAAAGCCACCTCTAAGCGCCAGGCCATGCCCGGCATGCGATGGGTACAGTGGTTTGGTGACGGCAAGTTTTCAGAGGTCTCTGCTGACAAACTCATGGCCCTGGGGCTATTCAGCCAGCACTTTAACCTGACCACCTTCAATAAGCTGGTTTCCTATAGGAAAGCCATATACCACACTCTGGAGGTAACATGGGTGTGGGTGCTAACTAGGCTTCTGTGG--TTGCATATCTCTT--------ATCTCCCTGAATAT----ACCCATGATTCCACAGAA--CAAAGTCAAGGTTG---TACCGGAAC----------ACCCCCTAATAT--AATGTTTTTGTTTTGTTTTCCTG---CAAA-AGTGGTTAATAAGTCGAAGGTGCATCGTTCAGGCAGTAGGAAGTTAGAACGCAGGAGACACGGGCATTGTTNNNGGAAAGG-----------AGACTCCCCACCTCATCTTACACCTGTTCCTCTTTC------CTTCAC-------------------------------------------------------AGAACGGATGGCTTCTGATGTTGCCAATAACAAAGTCAGTCTAGAAGACCGCTGTTTGTCCTGCGGTAGGAAGAACCCTGTGTCCTTCCACCCCCTCTTTGAGGGTGGGCTCTGTCAGAGTTGCCGGG---------ACCGCTTCCTGGAGCTCTTCTACATGTA-CGATGAGGAC-GGCTATCAGTCCTACTGCAC----CGTGTGCTGCGAGGGCCGC-----------------GAGCTGCTTCTGTGCAGCA----------------------------ACACAAGC--------------TGCTGCAGGTGC-----------------------TTCTGTGTGGAGTGTCTGGAGGTGCTGG--TGGGT--ACAGGGACAGCT-----GAGGATGC--CAAG----CTGCAGGAACCCTGGAGC-TGCTATATGTGCCTC----------------------------------------------------------------------------------------------------------------------------------------------------------------------------------------------------------------------------------------------CCACAG----------CGCTGCCA----TGG---------------GTCCATCGCCGTGGGAACTGTTAAGCATGAAGGCCAAATCAAATATGTGAACGACGTCAGGAAAATCACAAAGAAAAATGT----GAGGGTGACCTTTTT--CCTGGA---------------------------------------------------CCCTTGCAATGACCTCTCCAATGTCAATCCTGCCAGGAAAGGCCTATATGAGGGTACCGGCAGGCTCTTCTTTGAGTTTTACCACTTGCTGAATTATACACGCCCCAAGGAGGGCGACAACCGTCCGTTCTTCTGGATGTTTGAGAATGTGGTTGCCATGAAGGTTAATGACAAGAAAGACATCTCCAGATTCTTGGCGTGTAACCCAGTGATGATTGATGCCATCAAAGTTTCTGCTGCTCACAGGGCCCGATACTTCTGGGGCAACCTGCCTGGGATGAA---CAGG------------------------------------------------------------------------------------------------------------------------------------------------------------------------------------------------------------------------------------------------------------------------------------------------------------------------------------------*

*>BankVolePredictedCDS_3B_(incomplete)*

*------------------------------------------------------------------------------------------------------------------------------------ATGAAGGGAGACAGCAGACATCTGAATGA------AGAAGAGGGCGCCAGTGGGGGTGAGGAATGTGTCATT---GTCAATGGGAACTGTAGTGACCATTCCTCAGACACTAAGGATGCTCCCTCACCCCCAGTCTTGGAG--GCAGTCTGCGCAGAGGCAGACAGCACACCAGAGAGCAG-----AGGCCGCAGATCAAGTTCAAGGCTGTCAAAGAGGG--------------------AGGTCTCCGGCCTGCTGAGTTACACTCAGGACCTGACAGGAAATGGAGATGA------------TGAGGCAGAGGATGGGGACGGCGATGATGTCCTACTGATGCCAAAGCTC---------ACACGCGAGACCAAGGAGACCAAGTCATTCTCTGAAAGCCCGGCTATGCGAACCCGAAATAACAACA---GTACCTCCAGCCTAGGGAGGCAAAGAACCTCCCTCAGA---------ATCACCCGAGGCCGCCAGGGCCGACACCATGTGCAGGAATACCCCGTGGAGTTCCCAGCTACCAGGTCTCGGAGGCGTCGGGCATC------GTCAGCAA-GCACGCCATGGTCGTCTCCTGCCAGCCCTTCCC------------TCGTAGAAGA---AGGGACACCTCAGAACAGCAGT------ACCCCAT-CAATTGACTTGAGCCAGGACATCT-ATCAGGAGGACATAGACACTACACAGCTGGAGGCAGAAGGCAAAGATGGAGACAGCACTGAGTACCAGGATGATAAGGAGTTTGGAATAGGCGATCTTGTGTGGGGAAAAATCAAAGGATTCTCCTGGTGGCCTGCCATGGTGGTGTCCTGGAAAGCCACCTCCAAGCGCCAGGCCATGCCCGGCATGCGATGGGTACAGTGGTTTGGTGACGGCAAGTTTTCAGAGATCTCTGCTGACAAACTCATGGCCCTGGGGCTATTCAGCCAGCACTTTAACCTGGCCACCTTCAATAAGCTGGTGTCCTATAGGAAGGCCATGTACCACACTCTGGAGGTAACATGGGTGCGGGTGCTAACTAGGCTGGTGCGAGCTGGCAAGACCTTCCCCA--GCAGCCCCGGAGACTC----ACTGGAGGACCAGCTGAAGCCCATGCTAGAGTGGGCCCACGGGGGC--TTCA---AGCCCACTGGCATTGAGGGCCTCAAACCGAACAACAAG---CAACCAGTGGTTAATAAGTCGAAGGTGCATCGTTCAGGCAGTAGGAAGTTAGAACCCAGGAGACACGAGAACAAAAGTCGAAGACGTACAACC----AACGACTCTGCCTCTTCTGAGTACTGTCCCCCACCC-AAGCGCCTCAAGACAAATAGCTAT---GGCGGGAAAGACCGAGGGGAGGAT-GATGAG---AGCCGAGAACGGATGGCTTCTGACGTTGCCAACAACAAAGGCAATCTGGAAGACCGCTGTTTGTCCTGCGGTAGGAAGAACCCTGTGTCCTTCCACCCCCTCTTTGAGGGTGGGCTCTGTCAGAGTTGCCGGATAAGAGGACAACCTCTCCTGTGTGATTTACCCAGGTATCAATCAGGCTTGGTGACAAGTTCCTCTGCCCGCCACATTCCCTCCTTGGCCCTTTTCTATGTTTCTTGTTTGACTTGTTTTCCTTCGGCTCAAGTATCTGGAAGTTTCCTGGCTACACACACAAGTGTTGTGATAGATAATGACGCAGTGACCCTCAGTCTTGGCTCCTTTCTAGTTTTGTTTTCAGTGTGTATGCGCACAGGATTCAGTCAACACGCACAGCCCTCTCAGGTCTGC--TGTGGCTCCTGCAGGTTTCTTGGTGCCTGGTACTTGGTGCTCA----------------------------------------------------------------------------------------------------------------------------------------------------------------------------------------------------------------------A--AGACTTG---------GGTATCAAAGTGGAAAAGTACGTTGCCT----CGGAAG--TCTGTGCAGAGTCCATCGCCGTGGGAACCGTTAAGCATGAAGGCCAAATCAAATATGTGAATGACGTCAGGAAAATCACAAAGAAAAATGT----GAGGGTGACCTTATT--CTTGGATGGAGTTCATCTCATCTTCCTTATCCTTCTTCTATACCCTGTGCCCTCCTGCCCTTGCAATGACCTCTCCAATGTCAATCCTGCCAGGAAAGGCCTATATGAGGGTACCGGCCGGCTCTTCTTTGAGTTTTACCACTTGCTGAATTATACACGCCCCAAGGAGGGCGACAATCGTCCGTTCTTCTGGATGTTTGAGAATGTGGTGGCCATGAAGGTTAATGATAAGAAAGACATCTCCAGATTCTTGGCGTGTAACCCAGTGATGATCGATGCCATCAAGGTTTCTGCCGCTCACAGGGCCCGGTACTTCTGGGGCAACCTGCCCGGGATGAA---CAGG------------------------------------------------------------------------------------------------------------------------------------------------------------------------------------------------------------------------------------------------------------------------------------------------------------------------------------------*

*>ParirieVole_3B_(incomplete)*

*------------------------------------------------------------------------------------------------------------------------------------ATGAAGGGAGACAGCAGACATCTGAATGA------AGAAGAGGGTGCCAGTGGGGGTGAGGAAAGTGTCATT---GTCAATGGGAACTGTAGTGACCATTCCTCAGACACTAAGGATGCTCCCTCACCCCCAGTCTTGGAG--GCAATCTGCACAGAGGCTGACAGCACACTGGAGAGCAG-----AGGCCGCAGATCAAGATCAAGGCTGTCAAAGAGGG--------------------AGGTCTCCGGCCTGCTGAGTTACACTCAGGACCTGACAGGAAATGGAGATGG------------TGAGGCAGAGGATGGGGATGGCGATGATGTTCTACCGATGCCAAAGCTC---------ACACGTGAGACCAAAGAGACCAGGTCACCCTCGGAAAACCCAGCTATGCGAACCCGAAATAACAACA---GTACCTCCAGGCTAGGGAGGCAAAGAGCCTCCCCCAGA---------ATCACCCGAGGCCGCCAGGGCCGACACCATGTGCAGGAATACCCCGTGGAGTTCCCAGCTACCAGGTCTCGGAGGCGTCGGGCATCA---TCGTCAGCAA-GCACGCCATGGTCGTCCCCTGCCAGCCCTTCCT------------TCATGGAAGA---AGGGACACCTCAGAACAGCAGT------ATCCCAT-CAATTGACTTGACCCAGGACATCA-ATCAGGAGAGCATGGACACTACACAGCTGGAGGCAGAAGGCAAAGATGGAGACAGCACTGAGTATCAGGATGACAAGGAGTTTGGAATAGGTGACCTTGTGTGGGGAAAAATCAAGGGATTCTCCTGGTGGCCTGCCATGGTGGTATCCTGGAAAGCCACCTCTAAGCGCCAGGCCATGCCCGGCATGCGATGGGTACAGTGGTTTGGTGACGGCAAGTTTTCAGAGGTCTCTGCTGACAAACTCATGGCCCTGGGGCTATTCAGCCAGCACTTTAACCTGACTACCTTTAATAAGCTGGTTTCCTATAGGAAAGCCATATACCACACTCTGGAGAGAGCCAGGGTGCGA--GCTGGCAAGACCTTCCCCA----GCA-----------------GCCCGGGAGATTC----CCTGGAGGACCAGCTGAAGCCCATGCTGGAATGGGCCCATGGGGGC--TTCA---AGCCCACTGGCATCGAGGGCCTCAAACCCAACAACAAG---CAACCAGTGGTTAATAAGTCGAAGGTGCATCGTTCAGGCAATAGGAAGTTAGGACCCAGGAGACACGAGAACAAAAGTCGAAGACGCACAACC----ACCGACTCTGCCTCTGCTGAGTACTGTCCCCCACCC-AAGCGCCTCAAGACCAATAGCTAT---GGCGGGAAAGACCGAGGGGAGGAT-GATGAG---AGCCGAGAACGGATGGCTTCTGATGTTGCCAATAACAAAGTCAGTCTAGAAGACCGCTGTTTGTCCTGTGGTAGGAAGAACCCTGTGTCCTTCCACCCCCTCTTTGAGGGTGGGCTCTGTCAGAGTTGCCGGG---------ACCGCTTCCTGGAGCTCTTCTACATGTA-CGATGAGGAC-GGCTATCAGTCCTACTGCAC----CGTGTGCTGCGAGGGCCGC-----------------GAGCTGCTTCTGTGCAGCA----------------------------ACACAAGC--------------TGCTGCAGGTGC-----------------------TTCTGTGTGGAGTGTCTGGAGGTGCTGG--TGGGT--ACAGGGACAGCT-----GAGGATGC--CAAG----CTGCAGGAACCCTGGAGC-TGCTATATGTGCCTCCCACAGCGCTGCC---ATGGGGTCCTCAGGCGCAGGAAGGATTGGAACACGCGCCTGCAAGACTTCTTCACTACTGACCCTGATCTCGAGGAATTT---GAGCCGCCCAAGGTGTTCCCAGCAATTCCTGCAGCCAGGAGGAGACC------------CATTAGAGTCCTGTCCCTGTTTGA-TGGAATTGCG---ACAGGGTACTTGGTGCTCAA--AGACTTG---------GGTATCAAAGTGGAAAAGTACGTTGCCT----CCGAAG--TCTGTGCCGAGTCCATCGCCGTGGGAACTGTTAAGCATGAAGGCCAAATCAAATATGTGAACGACGTCAGGAAAATCACAAAGAAAAACATTGAAGAGTGGGGCCCTTTTGACTTGGTGATTGGTGGAA---------------------------------------GCCCTTGCAATGACCTCTCCAATGTCAATCCTGCCAGGAAAGGTCTATATGAGGGTACCGGCCGGCTCTTCTTTGAGTTTTACCACTTGCTGAATTATACACGCCCCAAGGAGGGCGACAACCGTCCGTTCTTCTGGATGTTTGAGAATGTGGTGGCCATGAAGGTTAATGACAAGAAAGACATCTCCAGATTCTTGGCGTGTAACCCAGTGATGATCGATGCCATCAAGGTTTCTGCCGCTCACAGGGCCCGATACTTCTGGGGCAACCTGCCCGGGATGAA---CAGGATCTTCGGCTTTCCTGCACACTACACAGACGTGTCCAACATGGGCCGTGGTGCCCGCCAGAAGCTGCTGGGAAGGTCCTGGAGTGTGCCAGTCATCAGACACCTCTTTGCCCCCTTGAAGGACTACTTTGCATGTGAATAG---------------------------------------------------------------------------------------------------------------------------------------------------------------------------------------------*

*>Deer_mouse_3B*

*------------------------------------------------------------------------------------------------------------------------------------ATGAAGGGAGACAGCAGACATCTGAATGA------GGAAGAGGGAGCCAGTGGGTGTGAGGACTCTGTCATC---GTCAACGGGAACTGCAGTGACCAGTCCTCAGACACTAAGGATGCCCCCTCACCCCCCGTCTTGGAG--GCGATCTGCACAGAGGCAGTCAGCACACCAGAGAGCCG-----AGGCCGCAGATCAAGCTCACGGCTGTCAAAGAGGG--------------------AGGTCTCCAGCCTGCTGTCTTACACTCAGGACTCAGCAGGAGATGGAGATGG------------TGAGGCAGAGGATGGGGACGGCTCTGACATTGTA---ATGCCAAAGCTC---------ACGCGTGAGACCAAGGAGCCCAGGTCACCCTCTGAAAGTCCAGCGGTGCGAACCCGAAATAGCAACA---GTACCTCCAGCCTGGAGAGGCAAAGAGCCTCCCCTAGA---------ATCACCCGAGGCCGCCAGGGCCGCCACCATGTGCAGGAGTACCCCGTGGAGTTCCCAGCTACCAGGTCTCGGAGACGCCGAGCATCG---TCTTCAGCAA-GCACACCATGGCCATCCCCTGCCAGCCCGTACCCCAGCATTGACCTCATAGAAGA---AGTGACACCTCAGAGCAGCAGC------ACCCCAT-CGATTGACCTGAGCCGGGACAGCC-CGCAGGAGAGCATGGATGCTACACAGCTGGATGCAGATAGCAAAGACGGAGACAGCACGGAGTATCAGGATGATAAGGAGTTTGGAATAGGTGACCTCGTGTGGGGAAAAATCAAGGGCTTCTCCTGGTGGCCTGCCATGGTGGTATCCTGGAAAGCTACCTCCAAGCGCCAGGCCATGCCCGGCATGCGATGGGTACAGTGGTTTGGTGACGGCAAGTTTTCTGAGGTCTCAGCTGACAAACTCATGGCTCTGGGGCTGTTCAGCCAGCACTTTAACCTGGCCACCTTCAATAAGCTGGTGTCTTATAGGAAGGCCATGTACCACACTCTGGAGAGAGCCAGGGTGCGA--GCTGGCAAGACCTTCCCCA----GCA-----------------GCCCGGGAGACTC----GCTGGAGGACCAGCTGAAGCCCATGCTGGAGTGGGCCCATGGGGGC--TTCA---AGCCCACTGGCATCGAGGGCCTCAAACCAAACAACAAA---CAACCAGTGGTTAATAAGTCGAAGGTGCGTCGGGCAGGCAGTAGGAACTTAGAACCCAGGAGACACGAGAACAAAGGTCGAAGACGCACCACA----AATGACTCTGCCGCTTCTGAGTACTGTCCCACACCC-AAGCGCCTCAAGACAAACAGCTGC---GGAGGGAAAGACCGAGGGGAGGAT-GACGAG---AGCCGAGAACGGATGGCTTCTGATGTCACCAACAACAAGGGCAATCTGGAAGACCGCTGTTTGTCCTGCGGTAGGAAGAACCCTGTGTCCTTCCATCCCCTCTTTGAGGGTGGGCTCTGTCAGAGTTGCCGGG---------ATCGCTTCCTGGAGCTCTTCTACATGTA-TGATGAGGAC-GGCTATCAGTCCTACTGCAC----CGTGTGCTGCGAGGGCCGG-----------------GAGCTGCTTCTCTGCAGCA----------------------------ACACCAGC--------------TGCTGCAGGTGC-----------------------TTCTGTGTGGAGTGTCTGGAGGTGCTGG--TGGGG--ACAGGGACAGCC-----GAGGATGC--CAAG----CTGAAGGAACCCTGGAGC-TGCTATATGTGCATCCCACAGCGCTGCC---ATGGTGTCCTCAGGCGCAGGAAGGATTGGAACATGCGCCTGCAAGACTTCTTCACGACTGATCCTGACCTCGAAGAATTT---TGCCTTCTGAGGAGGGAACAAGA---------------------------------------------------------CGG-AATGTTTGGA---GCCTGGTACTTGGTGCTCAA--AGAGTTG---------GGTATCAAAGTGGAAAAGTACGTCGCCT----CCGAAG--TCTGTTCAGAGTCCATCGCTGTGGGAACCGTTAAGCATGAAGGCCATATCAAATACGTGGATGACGTCAGGAAAATCACAAAGAAAAATATTGAGGAGTGGGGCCCATTTGACTTGGTGATTGGTGGAA---------------------------------------GCCCTTGCAATGACCTCTCCAATGTCAATCCCGCCAGGAAAGGCCTATATGAGGGCACCGGCAGGCTCTTCTTTGAGTTTTACCACTTGCTGAATTATACACGCCCCAAGGAGGGCGACAACCGTCCATTCTTCTGGATGTTTGAGAATGTGGTAGCCATGAAGGTCAACGACAAGAAAGACATCTCCCGATTCTTGGCGTGTAACCCAGTGATGATCGATGCCATCAAGGTTTCTGCCGCTCACAGGGCCCGATACTTCTGGGGCAACCTCCCTGGGATGAA---CAGGCCCGTGATAGCTTCAAAGAATGATAAGCTCGAGCTGCAGGACTGCTTGGAGTTCAGTAGGACAGCAAAGGTAAGATGCTGAGTGGTCTATGCTGCACCTTCTGGAAAACTGACCTTGGCTCTATCCACGTGGCTC---------------------------------------------------------------------------------------------------------------------------------------------------------------------------------------------------*

*>ChineeseHAmster_3B*

*------------------------------------------------------------------------------------------------------------------------------------ATGAAGGGAGATAGCAGACATCTGAATGA------GGAGGAGGGTGCCAGCGGGTGTGAGGAATGTCTCATC---GTCAATGGGAACTGTAGTGACCAGGCCTCAGATACTAAGGATGCTCCTTCACCCCCAGTCTTGGAG--GCAATGTGCACAGAGGCAGTCAACACATCAGAGAGCAG-----AGGCCGAAGATCAAGCTCACGGCTGTCAAAGAGGG--------------------AGGTCTCCAACCTGCTGAGTTACACTCAGGACCTGGCAGGAGATGGAGATGGAGATGG------TGAAGCAGAGGATGGGGATGGCTCAGACATTCTACTAATGCCAAAGCTC---------ACGCGTGAGACCAAGGAGACAAGGTCACCCTCGGAAAGTCCAGCTGTTCGAACCCGAAACAGCAACA---TTACCTCCAGCCTGGAGAGGCAAAGAGCCTCGCCCAGA---------ATCACCAGAGGCCGCCAGGGCCGCCACCATGTGCAGGAATACCCCGTGGAATTCCCAGCTACCAGGTCTCGGAGAAGGCGAGCATCG---TCTTCTACAA-GCACACCATGGTCGTCCCCTGCCAGCCCTTATCCCAGTATCGACCTCATAGAAGA---AGGGACACCTCAGAGGAGCAGT------ACCCCAT-CAACTGACTGGAGCCAGGACAGCC-AGCAAGAGAGTATGGATGCCACACAACTGTATGCAGAGAGCAAAGATGGAGACAGCACTGAGTACCAGGATGATAAGGAGTTTGGAATAGGCGACCTTGTATGGGGAAAAATCAAGGGCTTCTCCTGGTGGCCTGCCATGGTGGTTTCCTGGAAAGCCACCTCCAAGCGCCAGGCCATGCCTGGCATGCGATGGGTACAGTGGTTTGGTGACGGCAAGTTTTCTGAGGTCTCTGCTGACAAACTCGTGGCCCTTGGGCTGTTCAGCCAGCACTTTAACCTGGCCACCTTCAATAAGCTGGTTTCTTATAGGAAGGCCATGTACCACACTCTGGAGAGAGCCAGGGTGCGA--GCTGGCAAGACCTTCCCCA----GCA-----------------GCCCTGGAGACTC----ACTGGAGGACAAGCTGAAGCCCATGCTGGAGTGGGCCCACGGGGGC--TTCA---AGCCCACTGGCATCGAGGGCCTCAAACCGAATAACAAG---CAACCAGTGATTAATAAGTCGAAGGTGCGTCGTGCAGGCAGTAGCAAGTTAGAACCCAGGAAACACGAGAACAAAAGTCGAAGAGGCACAACCAAC-AACGACTCTGCAGCTTCTGAGTCCTATCCCCCTCCC-AAGCGCCTCAAGACCAACAGCTAC---GGTGGGAAAGACCGAGGGCAGGAT-GATGAG---AGCCGAGAACGGATGGCTTCTGATGTCACCAACAACAAGGGCAATCTGGAAGAACGCTGCTTGTCCTGTGGTAGGAAGAACCCTGTATCCTTCCATCCCCTCTTTGAGGGTGGGCTCTGTCAGAGTTGCCGGG---------ATCGCTTCCTGGAGCTGTTCTACATGTA-CGATGAGGAC-GGCTATCAGTCCTACTGCAC----TGTGTGCTGCGAGGGCCGA-----------------GAGCTGCTTCTGTGCAGCA----------------------------ACACAAGC--------------TGCTGCAGGTGC-----------------------TTCTGTGTGGAGTGTCTGGAGGTGCTGG--TGGGT--ACAGGGACAGCT-----GAGGATGC--CAAG----CTGCAGGAACCCTGGAGC-TGCTATATGTGCCTTCCACAGCGCTGCC---ATGGGGTCCTCAGGCGCAGGAAGGATTGGAACATGCGCCTGCAAGACTTCTTCACTTCTGATCCTGACCTCGAAGAATTC---GAGCCACCCAAGTTGTACCCAGCAATTCCTGCAGCCAAGAGGAGACC------------TATTAGAGTCCTGTCCCTGTTTGA-TGGAATTGCA---ACAGGGTACTTGGTGCTCAA--AGAGCTG---------GGTATCAAAGTGGAAAAGTATGTTGCGT----CCGAAG--TGTGTACAGAGTCCATTGCTGTGGGAACTGTTAAGCATGAAGGTCAAATCAAATATGTGAATGACGTCAGGAAAATCACAAAGAAAAATATTGAAGAGTGGGGCCCTTTTGACTTGGTGATTGGTGGAA---------------------------------------GCCCTTGCAATGACCTCTCCAATGTCAATCCAGCCAGGAAAGGCCTATATGAGGGCACCGGCAGGCTCTTCTTTGAGTTTTACCACTTGCTGAATTATACCCGCCCCAAGGAGGGCGACAACCGTCCGTTCTTCTGGATGTTTGAGAATGTGGTGGCCATGAAGGTTAACGACAAGAAAGACATCTCCCGATTCCTGGCGTGTAACCCAGTGATGATCGATGCCATCAAGGTTTCTGCTGCACATAGGGCCCGATATTTCTGGGGCAACCTACCTGGGATGAA---CAGGCCTGTGATAGCTTCCAAGAATGATAAGCTCGAGCTGCAGGACTGCTTGGAGTTCAGTAGGACAGCAAAGTTAAAGAAAGTACAGACAATAACCACCAAGTCGAACTCCATCAGACAGGGGAAAAACCAGCTTTTCCCTGTAGTCATGAATGGCAAGGACGATGTTTTGTGGTGTACTGAGCTCGAAAGGATCTTCGGCTTTCCTGCACACTACACGGATGTATCCAACATGGGCCGTGGTGCCCGCCAGAAGCTGCTGGGAAGGTCCTGGAGTGTGCCAGTCATCAGACACCTCTTTGCCCCCTTGAAGGACTACTTTGCATGTGAATAG*

*>FieldVolePredicted_3C_(incomplete)*

*------------------------------------------------------------------------------------------------------------------------------------ATGGAGGGGGAGAGCAGACAGCTCAATGC------TGATGAGGCTGCCGGTGGATGTGAGGATTGTGTCATC---ATCAGTGGGAACTGCAGTGACCAGTCCTCAGACGCCAAGAATGTTCCCTTGACCCAAGTCATGGAG--GCAATCTGC---------------ACAGTGGAGCACGA-----AGGCTGCAGAGCAAACTCACGACCGTCCAAGAGGA--------------------AGGTCTCCAGCACGATTGCTTACATCCAGGACCTGACTGAAGATGGAGATGAAGGTAGAGGTGGCGAGGCAGAGGACAGCAATGGCTCTGGTTCTCTGGTGACGCCCACACTC---------TCCTGGGAGACCA---------GCGCACCCTCTAATACCTCAGC------------------------------------------------------------------------------------------------------------------------------------------ATCCCTGAGCTGGCAAGCAACC---ACTTCAGCAA-GCACACCACAGCTGTCCCCTGCCAGCCCTTACCCCACCATTGACCTCACAGATGAAGAAGCGACACCCTGGAGCATCAGT------TCCCCAC-TGGTTGAGCTGAGCCAGGACAGCC-ACCAGGAGGGCATGGACACTACCCAGGTGGATGTGGAAAGCAGAGAGGGAGCCAACACTGAGTATCAGGT---------------------------------------------------------------------------------------------------------------------------------------------------------------------------------------------------------------------------------------------------------------------------ACC------------ACTGAGAAGGCAGTCT--G----ACT-----------------GTCTTAGTGTTTCTGTTGCTGTGAAGAGACACCATGACCATGGCAACTCTTATAAAGGAAGCCATTTCATTGGACCCACTGGCATCGAGGGCCTCAAACCCAACAACAAG---CAACCAGA------------------------------------------------------------GAACAAAAGTCGAAGACGCACAACCA---CCGA-CTCTGCCTCTGCTGAGTACTGTCCCCCACCC-AAGCGCCTCAAGACCAATAGCTAT---GGCGGGAAAGACAAAAGGGAGGAT-GATGAG---AGCCGAGGTGAATT---------------------TGGGG----------------------GTGGTGCAGCAGCAGGAGCAGCAGGTCCTCAGGACCACTGCTCTGGGGTTCAGAGGTGCATGGCTGCATGG---------GAAAG-----GAGACTTCCTTAC---------------------------CTTCCTGTCC----CTTTTTCCTCCACA-------------------------------------------------------------------------------------------------------------------------------------------------------------------GAAAGGATGGCT-----TCTGATGTCGCCAA----CAACAAAGGCAATCTGGA-AGGTACTATGTCTCTTTGCCCTCCTTTT---TGGGTGCCCCAGGCCTGCCTTAGCCTGGCTTTGACCTTCTGCATGGA-TTTTAATCTGGAGCACTGGGCCCAATCAGAACT-TGTGCCTCAGTATCTGAGATTTAATGTTTATGAAAGATGGAGATGCCAAATCCAAAACAATATTCAGCCTCTTGTTTTGCCTGGCCAGGCCTTGGCTTACCTTGTGCTTCCTGTTCGTTTGAACTTGTCCTTCCCAGACAACAAAGTGGGATGGCGCTTTAAACAAGGTCGAGTCTTCTGTGCCGAGTCCATCGCCGTGGGCACCATTAAGCATGAAGGCCAAATCAAATACGTGGATGACGTCAGGAACATCACAAAGGAAAATATTGATGAGTGGGGCCCATTCGACTTGGTGATTGGTGGAA---------------------------------------GCCCTTGCAATGACCTCTCTTGTGTGAACCCTGTCCGAAAAGGCCTGTTTGAGGGCACTGGCCGACTCTTCTTTGAGTTTTATCGATTGCTAAACCACTCACGCCCTGAGGAGTGTGATGACCGTCCATTCTTCTGGATGTTTGAGAATGTGGTAGCCATGGATGTCGGTGACAAGCAGGATATCTCACGATTCCTGGAGTGTAACCCAGTGATGATCGATGCCATCAAGGTTTCTGCCGCTCACAGGGCCCGATACTTTTGGGGCAACCTGCCCGGGATGAA---CAGG------------------------------------------------------------------------------------------------------------------------------------------------------------------------------------------------------------------------------------------------------------------------------------------------------------------------------------------*

*>BankVolePredicted_3C_(incomplete)*

*------------------------------------------------------------------------------------------------------------------------------------ATGGAGGGGGAGAGCAGACAGCTCAGTGA------TGATGAGGGTGTCGGTGGATGTGAGGATTGTGTCATC---ATCAGTGGGAACTGCAGTGACCAGTCCTCAGACGCCAGGAACGTTCCCTTGACCCAAGTCATGGAG--GCGATCTGC---------------ACAGTGGGACGCGA-----AGGCTGCAGAGCAAACTCACGACCGTCCAAGAGGA--------------------GGGTCTCCAGCACGGTTGCTTACATCCAGGACCTGACAGAAGATGAAGATGAAGGTAGAGATGGCGAGGTGGAGGACAGCAGTGGCTCTGGCTCTCCGGTGTCGCCCACACTC---------TGCTGGGAGACCA---------GGGCACCCTCTAATACCTCAGC------------------------------------------------------------------------------------------------------------------------------------------ATCCCTGAGTTGGCAAGCAACC---ACTTCAGCAA-GCACACCACGGCTGTCCCCTGCCAGCCCTTACCCCATCATTGACCTCACAGATGAAGAAGTGATACCCGGGAGCATCAGT------GCCCCAT-TGGTTGACCTGAACCAGGACAGCC-ACCAGGAGGGCATGGACACCACACAGGCGAATGTGGAAAGCAGAGATGGAGCCAGCACCGAGGTGCGAG--------------------------------------------------------------------------------------------------------------------------------------------------------------------------------------------------------------------------------------------------------------------------------------------CTGGCAAGACCTTCCCCA----GCA-----------------GCCCGGGAGACTC----ACTGGAAGATCAGCTGAAGCCCATGCTGGAGTGGGCCCACGGGGGC--TTCA---AGCCCCCTGGCATCGAGGGCCTCCAACCGAACAACAAG---CAACCAGGT---------------------------------------------------------GGGAATGAATGGCACAGACGTAGGAACT---TAGAACCCAGGAGATACGGTATTCCCTCACCATCTTTGAGTGTTCTGTTTTCTGT-GCTCT---GTCCTGCGGTTTCTCGGTGTCCAC-GGTGCTC--AGTGAAGAAGAATC---------------------CAA-------------------------TCTTCTA--AGCTG----AGCTGAATCTCTCACCCTCACCTATGACCCTCAGGCTAACCTGGGTGCACAT---------CCAGCCTCCTAAAACCTTAGGAC---------------------------TCTTCCCTTC----TCTTTTGTTGGTTA-------------------------------------------------------------------------------------------------------------------------------------------------------------------ATATTAATGGCA-----CAT-ATGCAAGCAT----TGGCAGATGTCACTTTGT-CACCA--AAGACTCCTGGTGTTTCTGTT---T------TCTGAGACAAC---AGTCTCACTCTGCAGTCCTTTCTGG--TCTCA---TGGGGAATTGTAGCCG----------TGTGTCACCTCATCTGGCTCCTCATGACTTTAGAAG-----GGTGTC----------------TGAGTTTCTTACCTCGTGTG--TAGGAATGGG--GAAGAGGTACTTGGTGCTCAA--AGAGTTG---------GGTATCAAAGTAGAAAAGTACGTTGCCT----CCGAAA--TCTGTGCAGAGTCCATCACCGTGGGCACCATTAAGCATGAAGGCCAAATCAAATACGTGGATGACGTCAGGAACATCACAAAGGGAAATATTGATGAGTGGGGCCCATTCGACTTGGTGATTGGTGGAA---------------------------------------GCCCTTGCAATGACCTCTCTTGTGTGAACCCTGTCCGAAAAGGCCTGTTTGAGGGCACTGGCCGACTCTTCTTTGAGTTTTATCGATTGCTAAATCACTCACGCCCTGAGGAGTGTGATGACCGTCCATTCTTCTGGATGTTTGAGAATGTGGTAACCATGGAGGTCGGTGACAAGCGGGACATCTCACAATTCCTGGAGTGTAACCCAGTGATGATCGATGCCATCAAGGTTTCTGCCGCTCACAGGGCCCGGTACTTTTGGGGCAACCTGCCCGGGATGAA---CAGGCCTCCTTGTGCCGTCCTTTGTGAT----------TTCAGGAACAGCTTG--TCGGCTAAGCTGGGAAATGCCACTGTAGTGACCATCCATATATGCAAGAGTCACACTGGGAAACACTTTCTGAAGGATCTTTTCCCTGTAGTGATGAATGGCAAGGACGATGATTTGTGGTGCACTGAGCTCGAAAGGATCTTCGGCTTTCCTGCACACTACACAGACGTGTCCAACATGGGCCGTGGTGCCCGCCAGAAGCTGCTGGGGAAGTCCTGGAGTGTGCCAGTCATCAGACACCTCTTTGCCCCCCTGAAGGACTACTTTGCATGTGAA---*

*>Mus_spretus_Dnmt3C_CDS_(incomplete)*

*------------------------------------------------------------------------------------------------------------------------------------ATGAGGGGAGGTGGCAGACACCTCAGTAA------TGAGGAGGATGTCAGTGGATGTGAGGACTGTATTATC---ATCAGTGGGACCTGCAGTGACCAGTCTTCAGACCCCAAGACTGTTCCTTTGACCCAAGTCTTGGAG--GCAGTCTGC---------------ACAGTGGAGAGCAG-----AGGATGCAGAACAAGCTCACAACCATCCAAGAGGA--------------------AAGCATCCAGCCTGATTAGTTATGTTCAGGACCTCACAGGAGATGGAGATGAAGATAGGGATGGTGAGGTGGGGGGCAGCAGTGGCTCTGGCACTCCAGTGATGCCCCAACTC---------TTCTGTGAGACCA---------GGATACCCTCTAAAACCCCAGC------------------------------------------------------------------------------------------------------------------------------------------ACCCCTCAGTTGGCAAGCAAAC---ACTTCAGCAA-GCACGCCCTGGTTGTCCCCTGCCAGCCCTTACCCCATCATTGACCTCACAGATGAAGATGTGATACCCCAGAGCATCAGT------ACCCCAT-CGGTTGACTGGAGCCAGGACAGCC-ATCAGGAGGGCATGGATACCACACAGGTGGATGCAGAGAGCAGAGATGGAGGCAACATTGAGTATCAGGT-------------------------------------------------------------------------------------------------------------------------------------------------------------------------------------------------------------------------------------------------------------------------------------------CTCGGCTGACAAACT-------GTT-----------------GCTCAGCCAGTCC-------------------TGTATCCTGGCCGCCTTCTATAAACTGGTTCCTTACA----------------GGAAGTCTATATACC---------G---TACTCTGG----------------------------------------------------------------AGAAAGCCAGGGTG-AGAGCTG---GCAAGGCCTGCCCCAGCAGT-------------CCTGGAGAGTC------------------------------------------------------ACTGGAGG----------------------------------------------------------------ACCAG---------------------------CTGAAGCCCATGCTGGAGTGGGCCCACG--------------------GTGGCTTCA--AG---------------------------CCTACTG----------------------------------------------------------------------------------------------------------------------------------------------------------------------------------------GGATCGAGGGCC-----TCAAAC----CCAA----CA--AGAAGCAACC---------------------------------------------------------------------------------------------AGAGAACAAAAGTCG-------------------------------------------AAGA----------------------------------------CGCACAACCA-----------ACAGGGTACTTGGTGCTCAA--GGAGTTG---------GGTATTAAAGTGGAAAAGTACATTGCCT----CCGAAG--TCTGTGCAGAGTCCATCGCTGTGGGAACCATTAAGCATGAAGGCCAGATCAAATACGTGGATGACATCAGGAACATTACAAAGGAACATATTGACGAGTGGGGCCCGTTCGACCTGGTGATTGGTGGAA---------------------------------------GCCCCTGCAATGATCTTTCCTGTGTGAATCCTGTCAGGAAAGGCCTGTTTGAGGGTACTGGCCGGCTCTTCTTTGAGTTTTACCGATTGCTAAATTACTCATGCCCTGAGGAAGAGGATGACCGCCCATTCTTTTGGATGTTTGAGAATGTGGTAGCCATGGAGGTCGGTGACAAGAGGGACATCTCACGATTCCTGGAGTGTAACCCAGTGATGATCGATGCCATCAAGGTGTCTGCTGCTCACAGGGCCCGGTACTTCTGGGGTAACCTACCCGGAATGAA---CAGGCCCGTGATGGCTTCAAAGAATGATAAGCTCGAGCTGCAGGACTGCCTGGAGTTCAGTAGGACAGCAAAGTTAAAGAAAGTGCAGACAATAACCACCAAGTCGAACTCCATCAGACAGGGCAAAAACCAGCTTTTCCCTGTAGTCATGAATGGCAAGGACGACGTTTTGTGGTGCACTGAGCTCGAAAGGATCTTCGGCTTTCCTGAACACTACACAGACGTGTCCAACATGGGCCGTGGCGCCCGTCAGAAGCTGCTGGGCAGGTCTTGGAGTGTGCCAGTCATCAGACACCTGTTTGCCCCCTTGAAGGACCACTTTGCCTGTGAA---*

*>ChineeseHamster_DNMT3C_ORF*

*------------------------------------------------------------------------------------------------------------------------------------ATGAAGAGAAACGGCAGACACCTCAGTGA------TGAAGAGGGTGTCAGTGGGTGTGAGGACTGTGTCATC---ATCACTGGGACCTGCAGTGACCAGTCCTCAGACGCCAAGGATGTCACCTTGACCCAAGTCTTGGAG--GCATTCTGC---------------CCAGTGGAGAACAG-----TGGCTGCAGAGCAAGCTCACGACGGTTCAAGAGGA--------------------GGGTCTCCAACACGGTTACTTACTTTCAGGACCTGACAGGAGATGAAGATGA---------TGTCGAGGTGGAGGGCAGCAGTGGCTCTTGCAGTCCAGCGATGCCCAAATTC---------TTCTGGGAGACCA---------GGACACCCTCTAAAACCCCAGC------------------------------------------------------------------------------------------------------------------------------------------ACCCATTTACCAGCAAGCAACC---ACTTCAGCAA-GCACACCGTGGCTGTTCCCTACCAGCCCTTACCCAACCATTGACCTCACAGATGAAGAAGTGATACTCCAGAGCATCAGT------ACCCCAC-TGGTTGACTTGAGCCAGGACAGCC-ATCAGGAAAGCATGGATACCACACAGGTAGATGTGGGAAGCAGAGATGAAGATACCACCAAGTATCAGAG---------------------------------------------------------------------------------------------------------------------------------------------------------------------------------------------------------------------------------------------------------------------------AGCCAGGGTGCGA--GCTGGCAAGACCTTCCCCA----GCA-----------------GCCCTGGAGACTC----ACTGGAGGACAAGCTGAAGCCCATGCTGGAGTGGGCCCACGGGGGC--TTCA---AGCCCATTGGCATCGAGGGCCTCAAACCAAACAAGGAC---CGCTGCTTG---------------------------------------------------------TCCTGTGGTAGGAAGAACCCTGTATCCT---TCCATCCCCTCTTTGAGGGTGGGCTCTGTCAGAGTTGCCGGGAT-CGCTTCCTGGAGCTGT---TCTACATG----TACGATGAGGAC-GGCTATC--AGTCCTACTGCACT---------------------GTG-------------------------TGCTGCAAGGGCCG------------CGAGCTGCTTCTGTGCAGCAACACAAGCTGCTGCAGGTGCTTCT---------GTG------TGGAGTGTCTGGAG---------------------------GTGCTGGTGG----GT-------------------------------------------------------------------------------------------------------------------------------------------------------------------------------ACAGGGACAGCT-----GAGGATGC--CAAG----CTGCAGGAACCCTGGAGC-TGCTATATGTGCCTTCCACAGCGCTGCC---ATGGGGTCCTCAGGCGCAGGAAGGATTGGAACATGCGCCTGCAAGACTTCTTCACTTCTGATCCTGACCTCGAAGAA------GAGCCACCCAAGTTGTACCCAGCAATTCCTGCAGCCAAGAGGAGACC------------TATTAGAGTCCTGTCCCTGTTTGA-TGGAATTGCA---ACAGGGTACTTGGTGCTCAA--AGAGCTG---------GGTATCAAAGTGGAAAAGTATGTTGCGT----CTGAAG--TGTGTACAGAGTCCATTGCTGTGGGAACTGTTAAGCATGAAGGTCAAATCAAATATGTGGATGACGTCAGGAACATCACAAAGGGAAATATTGATGAGTGGGGCCCATTCGACTTGGTGATTGGTGGAA---------------------------------------GCCCTTGCAATGATCTCTCCTGTGTGAATCCTGGCAGGAAAGGGCTGTTTGAGGGCACTGGTCGGCTCTTCTTTGAATTTTATCGATTGCTAAATTACGCACGTCCCGAGGAGTGTGATAACCGCCCATTCTTCTGGATGTTTGAGAATGTGGTAGCTATGGAGGTTGGTGACAAGAGGGACATCTCACTATTCCTGGAGTGTAACCCAGTGATGATCGATGCCATCAAGGTTTCTGCTGCACACAGGGCCCGATATTTCTGGGGCAACCTACCTGGGATGAA---CAGGCCTGTGATAGCTTCCAAGAATGATAAGCTCGAGCTGCAGGACTGCTTGGAGTTCAGTAGGACAGCAAAGTTAAAGAAAGTACAGACAATAACCACCAGGTCGAACTCCATCAGACAGGGGAAAAACCAGCTTTTCCCTGTAGTCATGAATGGCAAGGACGATGATTTGTGGTGTACTGAGCTTGAAAAGATCTTTGGCTTTCCTTCACACTACACGGATGTGTCTAACATGGGCCGTGGTGCCCGCCAGAAGCTGCTGGGAAAGTCCTGGAGTGTGCCAGTCATCAGACACCTCTTTGCCCCGCTGAAGGACTACTTTGCATGTGAA---*

*>Rnovergicus_DNMT3C_ORF*

*------------------------------------------------------------------------------------------------------------------------------------ATGAATGGAGGTAGCAGACTACTCAGTGA------TGAGGAGGGTATCAGTGGATGCCAGGACTGTATCAGC---ATCAGTGGGACCTGCAGTGACCGGTCTTCAGACACCAAGACTCTTCCCTTGACCCAAGTCTTGGAG--GTACTCGGC---------------ACAGTGGAGAGCAG-----AGGCTGCAGAACAGGCTCACCACCATCCGAGAGAA--------------------AAGTCTCCAGCCTGATTAGTTACATTCAGGACCTCACGGGAGATGGAGATGA---------TGGTGAGGTCGGGGACAGCAGTGGCTCTGACACTCCAGTGATGCCCGAACTC---------TTCTGGGAGACCA---------GGACACCTTCTCAAGCCCCAGC------------------------------------------------------------------------------------------------------------------------------------------AGCTTCCAGTTGGCCAGCAACC---ACTTCAGCCA-GCTCACCCTGGCCGTCCCCTGCCAGCCCTTACCCCATCATTGACCTCACAGATGAAGATGTGATACCCCAGAGCATCAGC------ACCTCAT-CCAGTGACTTGAGCCTGGACGGCC-CTCAGGAGGACATGGATACCACACAGGTGGATGTAGAGAGCAGAGACGGGGACAGCCCTGAGTATCAGAA---------------------------------------------------------------------------------------------------------------------------------------------------------------------------------------------------------------------------------------------------------------------------AGCCATGGTGCGA--GCTGGCAAGACCTTCCCCA----GCC-----------------GCCCTGGAGACTC----ACTGGAGGACCAGCTGAAGCCCATGCTGGAGTGGGCCCACGGGGGC--TTCA---AGCCCACTGGGATCGAGGGCCTCAAACCCAACAACGAC---CACTGTTTG---------------------------------------------------------TCCTGCGGTAGGAAGAACCCTGTGTCCT---TCCACCCCCTCTTTGAGGGTGGGCTCTGTCAGAGTTGCCGGGAT-CGCTTCCTGGAGCTCT---TCTACATG----TACGATGAGGAC-GGCTATC--AGTCCTACTGCACC---------------------GTG-------------------------TGCTGTGAGGGCCG------------TGAGCTGCTTCTGTGCAGTAACACAAGCTGCTGCAGGTGCTTCT---------GTG------TGGAGTGTCTGGAG---------------------------GTGCTGGTGG----GT-------------------------------------------------------------------------------------------------------------------------------------------------------------------------------ACAGGCACAGCG-----GAGGATGC--CAAG----CTGCAGGAACCCTGGAGC-TGCTATATGTGCCTCCCGCAGCGCTGCC---ATGGGGTCCTGCGGCGCAGGAAGGACTGGAACATGCGCCTGCAAGACTTCTTCACTACTGATCCTGACCTGGAAGAG------GAGCCGCCCAAGTTGTACCCAGCGATTCCTGCAGCCAAAAGGAGGCC------------CATTAGGGTCCTGTCCCTGTTTGA-TGGAATTGCA---ACAGGGTACTTGGTGCTCAA--GGACTTG---------GGTATTAAAGTGGAGAAGTACGTTGCCT----CCGAAG--TCTGTGCAGACTCCATTGCTGTAGGAACCATTAAGCATGAAGGACAAATCAAATACGTGGATGACATCCAGAACATTGCAAAGGAACATATTGACGAGTGGGGCCCGTTCGACCTGGTGATTGGTGGAA---------------------------------------GCCCATGCAATGATCTTTCCTGTGTGAATCCCATCAGGAAAGGCCTGTTTGAGGGCACTGGCCGGCTCTTCTTTGAGTTTTACCGATTGCTAAATTACTCACGTCCTGAGGAGGAGGATGACCGCCCATTCTTTTGGATGTTTGAGAATGTGGTAGCCATGGAAGTTGGTGACAAGAGGGACATTTCACGATTCCTGGAGTGTAACCCAGTGATGATCGATGCCATCAAGGTTTCTGCTGCTCACAGGGCCCGATACTTCTGGGGCAACCTACCTGGAATGAA---CAGGCCTGTGATAGCTTCAAAGAATGATAAGCTCGAGCTACAAGACTGCTTGGAGTTCAGTAGGACAGCAAAGTTAAAGAAAGTGCAGACAATAACCACCAAGTCGAACTCCATCAGACAGGGGAAAAACCAGCTTTTCCCTGTAGTCATGAACGGCAAGGATGACGTTCTGTGGTGCACTGAGCTCGAAAGGATCTTCGGCTTCCCGGCTCACTACACGGATGTGTCCAACATGGGCCGTAGTGCCCGCCAGAAGCTGCTGGGAAAGTCCTGGAGTGTGCCAGTCATCAGACACCTGTTTGCCCCCTTGAAGGACTACTTTGCATGTGAA---*

*>Pahari_3C*

*------------------------------------------------------------------------------------------------------------------------------------------GGAAGTAGCAGACACCTCAGCGA------TGAGGAGGATGTCAGTGGATGTGAGGACTGTATCATC---ATCAGTGGGACCCGCAGTGACCAGTCTTCAAACCCCAAGACTGTTCCTTTGACCCAACTCTTGGAG--GCAGTCTGC---------------ACAGTGGAGAGCAG-----AGGATGCAGGACAAGCTCACGGCCATCCAAGAGGA--------------------AAGTATCCAGCCTGATTAGTTCCGTTCAGGACCTCACAGGAGATGGAGATGGAGACAGGGATGGTGAGATGGGGAACAGCAGTGGCTCTGACACTCCAGTGATG------------------TTCTGCGAGACCA---------GGACACCCTCTAAAACCCCAGC------------------------------------------------------------------------------------------------------------------------------------------AAAAACCACTTCAGAAGCAACC---ACTTCAGCAA-GCACACCCAGGCTGTCCCCTGTCAGCCCTTACCCCATCATTGACCTCACAGATGAAGATACGATACCCCAGAGCATCAGT------ACCCCAT-CGGTTGACTGGAGCCAGGACAGCC-ATCAGGAGGGCATGGATACCACACAGGTGGATGCAGAAAGCAGAGATGGACATAACACTGAGTATCAGAA---------------------------------------------------------------------------------------------------------------------------------------------------------------------------------------------------------------------------------------------------------------------------AGCCAGGGTTCGA--GCGGGCAAGACCTTCCCCA----GCA-----------------GTCCCGGAGAGTC----ACTGGAGCACCAGCTGAAGCCCGTGCTGGAGTGGGCCCACAGCAGC--TTCA---AGCCCACTGGGATTGAGGGCCTCCAACCCAACAGCAAG---CAGCCAGA------------------------------------------------------------GGACAAAAGTCAAAGACGCACAACCA---ATGA-CTCTGCCGTTTCTGAGTACTCTACCCCACCC-AAGCGCCTCAAGACGAATAGCTAT---GGTGGGAAGGACCTAAAGGGGGAT-GAGGAG---AGCCGAGGTGATTT---------------------TGGGGAGATGCTGGAAGAACACTGCTTGTCCTGCGGTAGGAGGGACCCTGTGTCCTTCCACCCCCTCTTTGAGGGTGGGCTCTGTCAGAGTTGCCGGG---------ACCGCTTCCTGGAGCTCTTCTACATGTA-TGACGAGGAC-GGCTATCAGTCCTACTGCAC----CGTGTGCTGTGAGGGCTAT-----------------GAATTGCTGCTGTGCAGTA----------------------------ACACAAGC--------------TGCTGCAGGTGC-----------------------TTCTGTGTGGAGTGTCTGGAGGTGCTGG--TGGGT--GCAGGCACAGCT-----GAGGATGC--CAAG----CTGCAGGAACCCTGGAGT-TGCTATATGTGCCTCCCCCAGCGCTGCC---ATGGGTTCCTCCGACGCAGGAAGGATTGGAACATACGCCTGCAGGACTTCTTCACTACTGATCCTGACCTGGAAGAATTTCAGGAGCCACCCAAGTTGTACCCAGCAATTCCTGCAGCCAAAAGAAGGCC------------CATTAGAGTCCTGTCTCTGTTTGA-TGGAATTGCG---ACAGGGTACTTGGTGCTCAA--GGAGTTG---------GGTATTAAAGTGGAGAAGTACGTTGCCT----CCGAAG--TCTGTGCAGAGTCCATCGCTGTGGGAACCGTTAAGCATGAAGGCCAAATCAAATATGTGGATGACATCAGGAACATTACAAAGGAACATATCGACGAGTGGGGCCCATTCGACCTGGTGATTGGTGGAA---------------------------------------GCCCCTGCAATGATCTTTCCTGTGTGAATCCTGTCAGGAAAGGCCTGTTTGAGGGTACTGGTCGGCTCTTCTTTGAGTTTTATCGATTGCTAAATTACTCACGCCCTGAGGAGGAGGATGACCGCCCATTCTTTTGGATGTTTGAGAATGTGGTAGCCATGAAGGTTGGTGACAAGAGGGACATCTCACGATTCCTGGAGTGTAACCCAGTGATGATCAATGCCATCAAGGTTTCTGCTGCTCACAGGGCCCGGTACTTCTGGGGCAACCTACCTGGAATGAA---CAGGCCCGTGATAGCTTCAAAGAATGATAAGCTCGAGCTGCAGGACTGCCTGGAGTTCAGTAGGACAGCAAAGTTAAAGAAAGTGCAGACAATAACCACCAAGTCGAACTCCATCAGACAGGGGAAAAAACAGCTTTTCCCTGTAGTCATGAATGGCAAAGACGACATTCTGTGGTGCACTGAGCTCGAAAGGATCTTCGGCTTCCCTGCTCACTACACGGATGTGTCCAACATGGGCCGTGGTGCCCGCCAGAAGCTGCTGGGCAGGTCCTGGAGTGTGCCAGTCATCAGACACCTGTTGGCCCCCTTAAAGGACCACTTTGCCTGTGAA---*

*>Mouse_Dnmt3c_long_isoform_CDS*

*------------------------------------------------------------------------------------------------------------------------------------ATGAGGGGAGGTAGCAGACACCTCAGTAA------TGAGGAGGATGTCAGTGGATGTGAGGACTGTATTATC---ATCAGTGGGACCTGCAGTGACCAGTCTTCAGACCCCAAGACTGTTCCTTTGACCCAAGTCTTGGAG--GCAGTCTGC---------------ACAGTGGAGAACAG-----AGGATGCAGAACAAGCTCACAACCATCCAAGAGGA--------------------AAGCATCCAGCCTGATTAGTTACGTTCAGGACCTCACAGGAGATGGAGATGAAGATAGGGATGGTGAGGTGGGGGGCAGCAGTGGCTCTGGCACTCCAGTGATGCCCCAACTC---------TTCTGTGAGACCA---------GGATACCCTCTAAAACCCCAGC------------------------------------------------------------------------------------------------------------------------------------------ACCCCTCAGTTGGCAAGCAAAC---ACTTCAGCAA-GCACGCCCTGGCTGTCCCCTGCCAGCCCTTACCCCATCATTGACCTCACAGATGAAGATGTGATACCCCAGAGCATCAGT------ACCCCAT-CGGTTGACTGGAGCCAGGACAGCC-ATCAGGAGGGCATGGATACCACACAGGTGGATGCAGAGAGCAGAGATGGAGGCAACATTGAGTATCAGAA---------------------------------------------------------------------------------------------------------------------------------------------------------------------------------------------------------------------------------------------------------------------------AGCCAGGGTGAGA--GCTGGCAAGGCCTGCCCCA----GCA-----------------GTCCTGGAGAGTC----ACTGGAGGACCAGCTGAAGCCCATGCTGGAGTGGGCCCACGGTGGC--TTCA---AGCCCACTGGGATCGAGGGCCTCAAACCCAACAAGAAG---CAACCAGA------------------------------------------------------------GAACAAAAGTCGAAGACGCACAACCA---ATGA-CCCTGCTGCTTCTGAGTCCTC---CCCACCC-AAGCGCCTCAAGACAAATAGCTAT---GGCGGGAAGGACCGAGGGGAGGAT-GAGGAG---AGCCGAGAACAGATGGCTTCTGATGTCACCAACAACAAGGGCAATCTGGAAGACCACTGTTTGTCGTGCGGTAGGAAGGACCCTGTGTCCTTCCACCCCCTCTTTGAGGGTGGGCTCTGTCAGAGTTGCCGGG---------ACCGCTTCCTAGAGCTCTTCTACATGTA-TGACGAGGAC-GGCTATCAGTCCTACTGCAC----CGTGTGCTGTGAGGGCCGT-----------------GAACTGCTGCTGTGCAGTA----------------------------ACACAAGC--------------TGCTGCAGATGC-----------------------TTCTGTGTGGAGTGTCTGGAGGTGCTGG--TGGGT--GCAGGCACAGCT-----GAGGATGT--CAAG----CTGCAGGAACCCTGGAGC-TGCTATATGTGCCTCCCTCAGCGCTGCC---ATGGGGTCCTCCGACGCAGGAAAGATTGGAACATGCGCCTGCAAGACTTCTTCACTACTGATCCTGACCTGGAAGAATTTCAGGAGCCGCCCAAGTTGTACCCAGCGATTCCTGCAGCCAAAAGGAGGCC------------CATTAGAGTCCTGTCTCTGTTTGA-TGGAATTGCA---ACAGGGTACTTGGTGCTCAA--GGAGTTG---------GGTATTAAAGTGGAAAAGTACATTGCCT----CCGAAG--TCTGTGCAGAGTCCATCGCTGTGGGAACCGTTAAGCATGAAGGCCAAATCAAATATGTGGATGACATCAGGAACATTACAAAGGAACATATTGACGAGTGGGGCCCGTTCGACCTGGTGATTGGTGGAA---------------------------------------GCCCCTGCAATGATCTTTCCTGTGTGAATCCTGTCAGGAAAGGCCTGTTTGAGGGTACTGGCCGGCTCTTCTTTGAGTTTTACCGATTGCTAAATTACTCATGCCCTGAGGAGGAGGATGACCGCCCCTTCTTCTGGATGTTTGAGAATGTGGTAGCTATGGAGGTCGGTGACAAGAGGGACATCTCACGATTCCTGGAGTGTAACCCAGTGATGATCGATGCCATCAAGGTGTCTGCTGCTCACAGGGCCCGGTACTTCTGGGGTAACCTACCCGGAATGAA---CAGGCCCGTGATGGCTTCAAAGAATGATAAGCTCGAGCTGCAGGACTGCCTGGAGTTCAGTAGGACAGCAAAGTTAAAGAAAGTGCAGACAATAACCACCAAGTCGAACTCCATCAGACAGGGCAAAAACCAGCTTTTCCCTGTAGTCATGAATGGCAAGGACGACGTTTTGTGGTGCACTGAGCTCGAAAGGATCTTCGGCTTTCCTGAACACTACACAGACGTGTCCAACATGGGCCGTGGCGCCCGTCAGAAGCTGCTGGGCAGGTCTTGGAGTGTGCCAGTCATCAGACACCTGTTTGCCCCCTTGAAGGACCACTTTGCCTGTGAATAG*

*>Caroli_3C*

*------------------------------------------------------------------------------------------------------------------------------------------GGAGGTAGCAGACACCTCAATAA------TGAGGAGGATGTCAGTGGATGTGAGGACTGTATTATC---ATCAGTGGGACCTGCAGTGACCAGTCTTCAGACCCCAAGACTGTTCCTTTGATCCAAGTCTTGGAG--GCATTCTGC---------------ACAGTGGAGAGCAG-----AGGATGCAGAACAAGCTCACAACCACCGAAGAGGA-----------------------------------CTAGTTACGTTCAGGACCTCACAGGAGATGGAGATGAAGATAGGGATGGTGAGGTGGGGGGCAGCAGTGGCTCTGGCACTCCAGTGATGCCCCAATTC---------CTCTGCGAGACCA---------GGATACCCTCTAAAGCCCCAGC------------------------------------------------------------------------------------------------------------------------------------------ATCCCTCAGTTGGCAAGCAAAC---ACTTCAGCAA-GCACACCCTGGCTGTCCCCTGCCAGCCCTTACCCCATCATTGACCTCACAGATGAAGATGTGATACCCCAGAGCGTCAGT------ACCCCAT-CGGTTGACTGGAGCCAGGACAGCC-ATCAGGAGGGCATGGATACCACACAGGCAGATGCAGAGAGCAGAGATGGAGGCAACATTGAGTATCAGAA---------------------------------------------------------------------------------------------------------------------------------------------------------------------------------------------------------------------------------------------------------------------------AGCCAGGGTGCGA--GCTGGCAAGGCCTGCCCCA----GCA-----------------GTCCTGGAGAGTC----ACTGGAGGACCAGCTGAAGCCCATGCTGGAGTGGGCCCACGGTGGT--TTCA---AGCCCACTGGGATCGAGGGCCTCAAACCCAACAACAAG---CAACCAGA------------------------------------------------------------GAACAAAAGTCGAAGGCGCACAGCCA---ATGA-CTCTGCTGCTTCTGAGTATTCCCCCCCACCC-AAGCGCCTCAAGACCAACAGCTAT---GGTGGGAAGGACCGAGGGGAGGAT-GAGGAG---AGCCGAGAACGGATGGCTTCTGATGTCACCAACAACAAGGGCAATCTGGAAGACCACTGTTTGTCCTGTGGTAGGAAAGACCCTGTGTCCTTCCACCCCCTCTTTGAGGGTGGGCTCTGTCAGAGTTGCCGGG---------ATCGCTTCCTAGAACTCTTCTACATGTA-TGACGAGGAC-GGCTATCAGTCCTACTGCAC----CGTGTGCTGTGAGGGCCGT-----------------GAACTGCTGCTGTGCAGTA----------------------------ACACAAGC--------------TGCTGCAGATGC-----------------------TTCTGTGTGGAGTGTCTGGAGGTGCTGG--TGGGC--GCAGGCACAGCT-----GAGGATGC--CAAG----CTGCAGGAACCCTGGAGC-TGCTATATGTGCCTCCCTCAGCGCTGCC---ATGGGGTCCTCCGACGCAGGAAGGATTGGAACATGCGCCTGCAAGACTTCTTCACTACTGATCCTGACCTGGAAGAATTTCAGGAGCCGCCCAAGTTGTACCCAGCAATTCCTGCTGCCAAAAGGAGGCC------------CATTAGAGTCCTGTCTCTGTTTGA-TGGAATTGCA---ACAGGGTACTTGGTGCTCAA--GGAGTTG---------GGTATTAAAGTGGAGAAGTACATTGCCT----CCGAAG--TCTGTGCAGAGTCCATCGCTGTGGGAACCGTTAAGCATGAAGGCCAAATCAAATACGTGGATGACATCAGGAACATTACAAAGGAACATATTGATGAGTGGGGCCCGTTCGACCTGGTGATTGGTGGAA---------------------------------------GCCCCTGCAATGATCTTTCCTGTGTGAATCCTGTCAGGAAAGGCCTGTTTGAGGGTACTGGCCGGCTCTTCTTTGAGTTTTACCGATTGCTAAATTACTCATGCCCTAAGGAGGAGGATGACCGCCCATTCTTCTGGATGTTTGAGAATGTGGTAGCCATGGAGGTCGGTGACAAGAGGGACATCTCACGATTCCTAGAGTGTAACCCAGTGATGATCGATGCCATCAAGGTGTCTGCTGCTCACAGGGCCCGGTACTTCTGGGGTAACCTACCCGGAATGAA---CAGGCCCGTGATGGCTTCAAAGAATGATAAGCTCGAGCTGCAGGACTGCCTGGAGTTCAGTAGGACAGCAAAGTTAAAGAAAGTGCAGACAATAACCACCAAGTCGAACTCCATCAGACAGGGGAAAAACCAGCTTCTCCCTGTAGTCATGAATGGCAAGGACGACGTTCTGTGGTGCACTGAGCTCGAAAGGATCTTCGGCTTTCCTGAACACTACACAGACGTGTCCAACATGGGCCGTGGCGCCCGTCAGAAGCTGCTGGGCAGGTCCTGGAGTGTGCCAGTTATCAGACACCTGTTTGCCCCCTTGAAGGACCACTTTGCCTGTGAA---*

*>MountainBlindMole_DNMT3C_ORF*

*------------------------------------------------------------------------------------------------------------------------------------ATGACTGGAGACAGCAAATACCTCAGTGA------GGAAGAGGGTGCCAATGGGTGTGAGGACTCCATCATC---ATCATTGGGAACTTCAGTGACCAGTCCTCAGACACCAAGGATGTTCCCTCACCCCCTGCCTTGGAG--GCGGTCTGC---------------ACACTGGAAGGCAG-----AGGCTGCAGATCAAGCTCATGGCTGTCCAAGAGGA--------------------AGGTCTCCAGGCTGCTAAGTTACAATCAGGATCTGATGGGAGATGGAGATGA---------TGGTGAGGTAGAGGACAGGGTTGGAGCTGATACTCCAGTGATGCCAAAACTC---------TTCTGGGAGACTG---------GGACACCCTCTAAAACCCCAGC------------------------------------------------------------------------------------------------------------------------------------------ATCCATGAGGCGGAGGGCAAGA---ACATCAGCAG-GCATGCTGTGGCCATCCCCTTCCAGCCCCTACCCCACCATCGACCTCACAGATGAAGAAGTGACACCTGGGAGCAGCAGG------ACAGTTT-ATGTTGACCTAGACGAGGACAGCC-AGCAGGAGAGCATGGACTCCACACAGGTGGATACAGAAAGCAGAGATGGAGACAGCACTGAGTATCAGAA---------------------------------------------------------------------------------------------------------------------------------------------------------------------------------------------------------------------------------------------------------------------------AGCCAGGGTTCGA--GCTGGCAAGACCTTCCCCA----CCA-----------------GCCCTGCAGACTC----ACTGGAGGACCAGCTGAAGCCTATATTGGAGTGGGCCCATGGAGGC--TTCA---AGCCCACTGGGATTGAGGGCCTCAAACCCAACAGCGAC---AGCTGTTTG---------------------------------------------------------TCTTGTGGAAGGAAAAACCCCGTGTCCT---TCCACCCACTCTTTGAGGGTGGGCTTTGCCAGACATGCCGGGAT-CGCTTCCTTGAGCTGT---TCTACATG----TACGACGACGAT-GGCTATC--AGTCCTACTGCACCG----------------------------------------------TGTGCTGCGATGGCGG------------TGAACTACTTCTCTGCAGCAACATAAGCTGCTGCCGGTGCTTCT---------GTG------TCGAGTGTCTGGAC---------------------------GTGTTGGTGG----GC-------------------------------------------------------------------------------------------------------------------------------------------------------------------------------ACAGGCACGGCT-----GCAGATGC--CAAG----CAGCAGGCGTTCTGGAGC-TGCTACATGTGTGTGCGAAGGTGCAGCC---ATGGTTTTCTGCGGCGCCGGAAGGACTGGAACTTGCGCCTGCAGGCCTTCTTCTCCAGCATTCTTGACATTGAATA------TGAGGCCCCCAAGTTGTACCCAGCAATTCTTGCAGCCCAAAGGCGGCC------------CATTAGAGTCCTGTCCCTGTTTGA-TGGAATTGCG---ACAGGCTACTTGGTCCTCAA--AGAATTG---------GGTATTAAAGTGGACAGGTACGTCGCCT----CTGAAG--TCTGCGCAGAATCCATCGCTGTGGGAACCGTGAAGCATGAAGGGCAAATCAAATACGTGAATGACGTCAGGAAAATCACAAAGAGAAATATTGATGAGTGGGGCCCATTCGACCTGGTGATTGGTGGAA---------------------------------------GCCCATGCAATGATCTTGCCTCTGTGAATCCTGTCAGGAAAGGCCTGTTTGAGGGTACTGGCCAACTCTTCTTTGAGTTTTACCGCTTGCTAAATTACTCACGCCCCAGAGAGAGTGATGACCGTCCATTCTTCTGGATGTTTGAGAATGTGGTAGCCATGAAGATCTGTGACAAGAGGGACATCTCACGGTTCCTGGAGTGTAACCCAGTGATGCTTGATGCCATCAAGGTTTCCGCTGCTCACAGGGCCCGATACTTCTGGGGCAACCTACCTGGGATGAA---CAGNNNNNTGATAGCATCAAAGAATGATAAACTCGAGCTGCAGGACTGCTTGGAGTTCAGTAGGACAGCAAAGTTGAAGAAAGTACAGACAATAACCACCAAGTCGAACTCGATCAGACAGGGGAAAAACCAACTTTTCCCTGTAGTCATGAATGGCAAAGAAGATGTTTTGTGGTGCACTGAGCTCGAAAGGATCTTCGGCTTTCCTGTACACTACACAGATGTGTCCAACATGGGCCGTAGTGCCCGCCAGAGGCTGCTGGGAAGGTCCTGGAGTGTGCCCGTCATCAGACACCTCTTCGCCCCCTTGAAGGAATACTTTGCATGTGAA---*

*>DeerMouse3C*

*------------------------------------------------------------------------------------------------------------------------------------ATGGAGGGAGAGAGTAGATACCTCCAGGA------TGAAGAGGGTGTCACTGGATGTGAGGACTGTGTCGTC---ATCAATGGGACCTGCAGTGGCCAGTCCTCAGACACTGCCAATGGTCCCTTGATCCACGTCTTGGAG--GCAGTCTGC---------------ACGGTGGAGAGCAG-----AGACTGTAGAACAAGCTCACGACCATCCAAGAGGA--------------------AGGTCTCCAGCACGATTGCAGACATTCAGGACCTGACAGGAGATGGAGATGAAGGTAGAGATGACGAGGTGGAGGACAGCCGTGGCTCTGGCACCCCAGTGACACCCAAACTC---------TTCTGGAAGGCCA---------GGACACCCTCTAAGTCCCCAGC------------------------------------------------------------------------------------------------------------------------------------------ATCCCTGAGTTGGCAAGCAACC---ACTTCAGCAA-GCATGTCATGGCTGTCCCCTGCCAGCCCTTACCCTACCATTGACCTCACAGATGAAGAAGTGATACCCCAGAGCATCAGT------ACCCCAT-TGGTTGACTTGAGCCAGGACAGCT-ATCAGGAGAGCATGGATACCACACAGGTGGATGTGAAAAGCAGAGATGGAGATAACACCAAGTATCAGAG---------------------------------------------------------------------------------------------------------------------------------------------------------------------------------------------------------------------------------------------------------------------------AGCCAGGGTGCGA--GCTGGCAAGACCTTCCCCA----GCA-----------------GCCCGGGAGACTC----GCTGGAGGACCGGCTGAAGCCCGTGCTGGAGTGGGCCCACGGCGTC-----------------------GAGGGCCTCAAACCAAACAGCAAG---CAACCAGA------------------------------------------------------------GAACAAAGGTCGAAGACGCACCACAA---ATGA-CTCTGCCGCTTCTGAGTACTGTCCCACACCC-AAGCGCCTCAAGACAAACAGCTGC---GGAGGGAAAGACCGAGGGGAGGAT-GACGAG---AGCCGAGAGCGGATGGCTTCTGACGTCACCAACAACAAGGGCAATCTGGAAGACCGCTGTTTGTCCTGCGGTAGGAAGAACCCTGTGTCCTTCCATCCCCTCTTTGAGGGTGGGCTCTGTCAGAGTTGCCGGG---------ATCGCTTCCTGGAGCTCTTCTACATGTA-TGATGAGGAC-GGCTATCAGTCCTACTGCAC----CGTGTGCTGCGAGGGCCGG-----------------GAGCTGCTTCTCTGCAGCA----------------------------ACACCAGC--------------TGCTGCAGGTGC-----------------------TTCTGTGTGGAGTGTCTGGAGGTGCTGG--TGGGG--ACAGGGACAGCG-----GAGGATGC--CAAG----CTGAAGGAACCCTGGAGC-TGCTATATGTGCATCCCACAGCGCTGCC---ATGGTGTCCTCAGGCGCAGGAAGGATTGGAACATGCGCCTGCAAGACTTCTTCACGACTGATCCTGACCTCGAAGAA---TTTGAGCCGCCCAAGTTGTACCCAGCAATTCCCGCAGCCAGGAGGAGGCC------------CATTAGAGTCCTGTCCCTGTTTGA-TGGAATTGCG---ACAGGGTACTTGGTGCTCAA--AGAGTTG---------GGTATCAAAGTGGAAAAGTACGTCGCCT----CCGAAG--TCTGTTCAGAGTCCATCGCCGTGGGAACCGTTAAGCATGAAGGCCATATCAAATACGTGGATGACGTCAGGAAAATCACAAAGAAAAATATTGAGGAGTGGGGCCCATTTGACTTGGTGATTGGTGGAA---------------------------------------GCCCTTGCAATGACCTCTCCAATGTCAATCCCGCCAGGAAAGGCCTATATGAGGGCACCGGCAGGCTCTTCTTTGAGTTTTACCACTTGCTGAATTATACACGCCCCAAGGAGGGCGACAACCGTCCATTCTTCTGGATGTTTGAGAATGTGGTAGCCATGAAGGTCAACGACAAGAAAGACATCTCCCGATTCTTGGCGTGTAACCCAGTGATGATCGACGCCATCAAGGTTTCTGCCGCTCACAGGGCCCGATACTTCTGGGGCAACCTCCCTGGGATGAA---CAGGCCCGTGATAGCTTCAAAGAATGATAAGCTCGAGCTGCAGGACTGCTTGGAGTTCAGTAGGACAGCAAAGTTAAAGAAAGTACAGACAATAACCACCAAGTCGAACTCCATCAGACAGGGGAAAAACCAGCTTTTCCCTGTAGTCATGAATGGCAAGGACGATGTTTTGTGGTGTACTGAGCTCGAAAGGATCTTCGGCTTTCCTGCCCACTACACTGACGTCTCCAACATGGGCCGTGGCGCCCGCCAGAAGCTGCTGGGAAGGTCCTGGAGTGTGCCAGTCATCAGACATCTCTTTGCCCCCTTGAAGGACTACTTTGCATGTGAA---*

*>PrairieVole3C*

*------------------------------------------------------------------------------------------------------------------------------------ATGAAGGGAGACAGCAGACATCTGAATGA------AGAAGAGGGTGCCAGTGGGGGTGAGGAAAGTGTCATT---GTCAATGGGAACTGTAGTGACCATTCCTCAGACACTAAGGATGCTCCCTCACCCCCAGTCTTGGAG--GCAATCTGC---------------ACAGAGGCTGACAGC----ACACTGGAGAGCAGA--GGTGAGTATCATAG----------------------------CTCTGACACTACAGGCTCC------GACCTGACAGGAAATGGAGATG------------GTGAGGCAGAGGATGGGGATGGCGATGATGTTCTACCGATGCCAAAGCTCACACGTGAGACCAAAGAGACCA---------GGTCACCCTCGGAAAACCCAGC------------------------------------------------------------------------------------------------------------------------------------------TTCTCGGAGGCGTCGGGCATCA---TCGTCAGCAA-GCACGCCATGGTCGTCCCCTGCCAGCCCTTCCTTCATG---------------GAAGAAGGGACACCTCAGAACAGCAGT------ATCCCAT-CAATTGACTTGACCCAGGACATCA-ATCAGGAGAGCATGGACACTACACAGCTGGAGGCAGAAGGCAAAGATGGAGACAGCACTGAGTATCAGAG---------------------------------------------------------------------------------------------------------------------------------------------------------------------------------------------------------------------------------------------------------------------------AGCCAGGGTGCGA--GCTGGCAAGACCTTCCCCA----GCA-----------------GCCCGGGAGATTCC----CTGGAGGACCAGCTGAAGCCCATGCTGGAATGGGCCCATGGGGGC--TTCA---AGCCCACTGGCATCGAGGGCCTCAAACCCAACAACAAG---CAACCAGA------------------------------------------------------------GAACAAAAGTCGAAGACGCACAACCA---CCGA-CTCTGCCTCTGCTGAGTACTGTCCCCCACCC-AAGCGCCTCAAGACCAATAGCTAT---GGCGGGAAAGACCGAGGGGAGGAT-GATGAG---AGCCGAGAACGGATGGCTTCTGATGTTGCCAATAACAAAGTCAGTCTAGAAGACCGCTGTTTGTCCTGTGGTAGGAAGAACCCTGTGTCCTTCCACCCCCTCTTTGAGGGTGGGCTCTGTCAGAGTTGCCGGG---------ACCGCTTCCTGGAGCTCTTCTACATGTA-CGATGAGGAC-GGCTATCAGTCCTACTGCAC----CGTGTGCTGCGAGGGCCGC-----------------GAGCTGCTTCTGTGCAGCA----------------------------ACACAAGC--------------TGCTGCAGGTGC-----------------------TTCTGTGTGGAGTGTCTGGAGGTGCTGG--TGGGT--ACAGGGACAGCT-----GAGGATGC--CAAG----CTGCAGGAACCCTGGAGC-TGCTATATGTGCCTCCCACAGCGCTGCC---ATGGGGTCCTCAGGCGCAGGAAGGATTGGAACACGCGCCTGCAAGACTTCTTCACTACTGACCCTGATCTCGAGGAA---TTTGAGCCGCCCAAGGTGTTCCCAGCAATTCCTGCAGCCAGGAGGAGACC------------CATTAGAGTCCTGTCCCTGTTTGA-TGGAATTGCG---ACAGGGTACTTGGTGCTCAA--AGACTTG---------GGTATCAAAGTGGAAAAGTACGTTGCCT----CCGAAG--TCTGTGCCGAGTCCATCGCCGTGGGAACTGTTAAGCATGAAGGCCAAATCAAATATGTGAACGACGTCAGGAAAATCACAAAGAAAAACATTGAAGAGTGGGGCCCTTTTGACTTGGTGATTGGTGGAA---------------------------------------GCCCTTGCAATGACCTCTCCAATGTCAATCCTGCCAGGAAAGGTCTATATGAGGGTACCGGCCGGCTCTTCTTTGAGTTTTACCACTTGCTGAATTATACACGCCCCAAGGAGGGCGACAACCGTCCGTTCTTCTGGATGTTTGAGAATGTGGTGGCCATGAAGGTTAATGACAAGAAAGACATCTCCAGATTCTTGGCGTGTAACCCAGTGATGATCGATGCCATCAAGGTTTCTGCCGCTCACAGGGCCCGATACTTCTGGGGCAACCTGCCCGGGATGAA---CAGGCCCGTGATAGCTTCCAAGAATGATAAGCTCGAGCTGCAGGACTGCCTGGAGTTCAGTAGGACAGCAAAGTTAAAGAAAGTACAGACAATAACCACCAAGTCGAACTCCATCAGACAGGGGAAAAACCAGCTTTTCCCTGTAGTGATGAATGGCCAGGACGATGTTTTGTGGTGTACTGAGCTCGAAAGGATCTTCGGCTTTCCTGCACACTACACAGACGTGTCCAACATGGGCCGTGGTGCCCGCCAGAAGCTGCTGGGAAGGTCCTGGAGTGTGCCAGTCATCAGACACCTCTTTGCCCCCTTGAAGGACTACTTTGCATGTGAA---*

*>Mouse_3L*

*----------------------------------------------------------------------------------------------------------------------------------------------------------------------------------------------------------------------------------------------------------------------------------------------------------------------------------------------------------------------------------------------------------------------------------------------------------------------------------------------------------------------------------------------------------------------------------------------------------------------------------------------------------------------------------------------------------------------------------------------------------------------------------------------------------------------------------------------------------------------------------------------------------------------------------------------------------------------------------------------------------------------------------------------------------------------------------------------------------------------------------------------------------------------------------------------------ATGGGTTCCCGGGAGACACCTTCTTCTTGCTCTAAGACCCT---------TGAA-----------ACCTTGGACCTGGAGACTTC----------CGACAGCTCTAGCCCTGATGCTGACAGTCCTCTGGAAGAG---CA----ATGGCTG---------AAATCCTCCCCAGCCCTGAAG----GAGGA---------------------------------------------------------------------CAGTGTGGATGTGGTACTG---GAAGACTGCAAAGAGCCTCTGTCCCCCTCCTCGCCT------CCGACAG--------------------------------------------------GCAGAGAGATGATCAGGTACGAAGTCAAAGTGAACCGACGGAGCATTGAAGACATCTGCCTCTGCTGTGGAACTCTCCAGGTGTACACTCGGCACCCCTTGTTTGAGGGAGGGTTATGTGCCCCATGTAAGG---------ATAAGTTCCTGGAGTCCCTCTTCCTGTA-TGATGATGAT-GGACACCAGAGTTACTGCAC----CATCTGCTGTTCCGGGGGT-----------------ACCCTGTTCATCTGTGAGA----------------------------GCCCCGAC--------------TGTACCAGATGC-----------------------TACTGTTTCGAGTGTGTGGACATCCTGG--TGGGC--CCCGGGACCTCA-------GAGAGGATCAAT-----GCCATGGCCTGCTGGGTTTGCTTCCTGTGCCTGCCCTTCTCACGGA---GTGGACTGCTGCAGAGGCGCAAGAGGTGGCGGCACCAGCTGAAGGCCTTCC------ATGATCAAGAGGGAGCG---------GGCCCTATGGAGATATACAAGACAGTGTCTGCATGGAAGAGACAGCC------------AGTGCGGGTACTGAGCCTTTTTAG---------------------------------------------------AAATATTGATA--AAGTACTAAAGAGTTT-------------------GGGCTTTTTGGAAAGCGGTTCTGGTTCTGGGGGAGGAACGCTGAAGTACGTGGAAGATGTCACAAATGTCGTGAGGAGAGACGTGGAGAAATGGGGCCCCTTTGACCTGGTGTACGGCTCGA----------------------------------------CGCAGCCCCTAGGCAGCTCT-TGTGATCGCTGTC----------------------CCGGCTGGTACATGTTCCAGTTCCACCGGATCCTGCAGTATGCGCTGCCTCGCCAGGAGAGTCAGCGGCCCTTCTTCTGGATATTCATGGACAATCTGCTGCTGACTGAGGATGACCAAGAGACAACTACCCGCTTCCTTCAGACAGAGGCTGTGACCCTCCAGGATGTCCGTGGCAGAGACTACCAGAATGCTATGCGGGTGTGGAGCAACATTCCAGGGCTGAAGAGCAAGCATGCGCCCCTGACCCCAAAGGAAGAAGAGTATCTGCAAGCCC------AAGTCAGAAGCAGGAGCAAGCTG------------------GACGCCCCG---AAAGTTGACCTCCTGGTGAAGAACTGCCTTCTCCCGCT--------------GAGAGAGTACTTCA---AGTATTTTTCTCAAAACTCACTTCCTCTTTAGAAATGA-------------------------------------------------------------------------------------------------------------------------*

*>Caroli3L*

*----------------------------------------------------------------------------------------------------------------------------------------------------------------------------------------------------------------------------------------------------------------------------------------------------------------------------------------------------------------------------------------------------------------------------------------------------------------------------------------------------------------------------------------------------------------------------------------------------------------------------------------------------------------------------------------------------------------------------------------------------------------------------------------------------------------------------------------------------------------------------------------------------------------------------------------------------------------------------------------------------------------------------------------------------------------------------------------------------------------------------------------------------------------------------------------------------ATGGGTTCCCGGGAGACACCTTCTTCTTTCTCTAAGACCCT---------TGAA-----------ACCTTGGACCTGGAGACTTC----------CGACAGCTCTAGCCCTGACGCTGACAGTCCTCTGGAAGAG---CA----ATGGCTG---------AAATCCTCCCCAGCCCTGAAG----GAGGA---------------------------------------------------------------------CAATGTGGATATGGTACTG---GAAGACTGCAAAGAGCCTCTGTCCCCCTCCTCACCC------CCGACAG--------------------------------------------------GCAGAGAGATGATCAGGTACGAAGTCAAAGTGAACCGACGGAGCATTGAAGACATCTGCCTCTGCTGCGGAACTCTCCAGGTGTACACTCAGCACCCCTTGTTTGAGGGGGGGATATGTGCCCCATGTAAGG---------ATAAGTTCCTGGAGTCCCTCTTCCTGTA-CGATGATGAT-GGACACCAGAGTTACTGCAC----CATCTGCTGTTCCGGGGGT-----------------ACCCTGTTCATCTGTGAGA----------------------------GCCCCGAC--------------TGTACCAGATGC-----------------------TACTGTTTCGAGTGTGTGGATATCCTGG--TGGGC--CCAGGGACCTCA-------GAGCGGATCAAT-----GCCATGGCCTGCTGGGTTTGCTTCCTGTGCCTGCCCTTCTCACGGA---GTGGACTCCTGCAGAGGCGCAAGAGGTGGCGGCACCAGCTGAAGGCCTTCC------ATGATCAAGAGGGAGCA---------GGCCCTATGGAGATATACAAGACAGTGTCCACATGGAAGAGACAGCC------------AGTGCGGGTACTGAGCCTTTTTGG---------------------------------------------------GAATATTGATA--AAGTACTAAAGAGTTT-------------------GGGCTTTTTGGAAAGCGGTTCTGGTTCTGGGGGAGGAACGCTGAAGTACGTGGAAGATGTCACAAATGTCGTGAGGAGAGACGTGGAGAAATGGGGCCCCTTTGACCTGGTGTACGGCTCGA----------------------------------------CGCAGCCCCTAGGCAGCTCT-TGTGATCGCTGTC----------------------CTGGCTGGTACATGTTCCAATTCCACCGGATCCTGCAGTATGCACTGCCTCGCCAGGAGAGTCAGCGGCCCTTCTTCTGGATATTTATGGACAATCTACTGATGACTGAGGATGACCAAGAGACAACTGCCCGCTTCCTTCAGACAGAGGCTGTGACCCTCCAGGATGTCCGTGGTAGAGACTACCAGAATGTTATGCGGGTGTGGAGCAACATTCCAGGGCTGAAGAGCAAGCATGTGCCCTTGACCCCAAAGGAAGAAGAGTATCTGCAAGCCC------AAGTCAGGACCAGAAGCAAGCTG------------------GACGCCCAG---AAAGTTGACCTCCTGGTGAAGAACTGCCTTCTCCCCCT--------------GAGAGAGTACTTCA---AGTATTTTTCTTAA-------------------------------------------------------------------------------------------------------------------------------------------------*

*>Pahari3L*

*----------------------------------------------------------------------------------------------------------------------------------------------------------------------------------------------------------------------------------------------------------------------------------------------------------------------------------------------------------------------------------------------------------------------------------------------------------------------------------------------------------------------------------------------------------------------------------------------------------------------------------------------------------------------------------------------------------------------------------------------------------------------------------------------------------------------------------------------------------------------------------------------------------------------------------------------------------------------------------------------------------------------------------------------------------------------------------------------------------------------------------------------------------------------------------------------------ATGGGTTCCCGGGAGACACCTTCTTCTTGCTCTAAGACCTT---------TGAA-----------ACCGTGAACCTGGAGACTTC----------CGACAGCTCTAATCCCGACGCTGACAGTCCTCTGGAAGAG---CA----ATGGCTG---------AAATCCTCCCCAGACCTGAAG----GAGGAGG---------------------------------------------------------------A---CAGTGTGGATGTGGTACTG---GAAGACTGCGAGGAGCCTCTGTCCCCCTCCTCACCC------CCCGAAG--------------------------------------------------GCAGAGAGATCATCAGGTACGAAGTCAAAGTGAACCAACGGAACATTGAAGACATCTGCCTCGGCTGCGGAACTCTCCAGGTGTACACCCAGCACCCCTTGTTTGAGGGGGGGATATGTGCCCTGTGTAAGG---------ATAAGTTCCTGGAGTCCCTCTTCCTGTA-TGATGATGAT-GGACACCAGAGTTACTGCAC----CATCTGCTGTTCCGGGGGT-----------------ACCCTGTTCATCTGTGAGA----------------------------GCCCTGAC--------------TGTACCAGATGC-----------------------TACTGTTTCGAGTGTGTGGACATTCTGG--TGGCC--CCCGGAACCTCA-------GAGCGGATCAAT-----GCCATGGCCTGCTGGGTTTGCTTCCTGTGCCTGCCCTTCTCACGGA---GCGGACTGCTGCAGAGGCGCAAGAGGTGGCGGCACCAGCTGAAGGCCTTCC------ATGATCAAGAGGTAGCA---------GGCCCTATAGAGATATACAAGACAGTGTCCGCGTGGAAGAGACAGCC------------AGTGCGGGTACTGAGCCTTTTTGG---------------------------------------------------GAATATTGATA--AAGTACTAAAGAGTTT-------------------GGGCTTTTTGGAAAGCGGTTCTGGTTCTGAGGGAGGAATGCTGAAGTACGTGGAAGATGTCACAAATGTCGTGAGGAGAGACGTGGAGAAATGGGGCCCCTTTGACCTGGTGTATGGCTCGA----------------------------------------CGCAGCCCCTAGGCAGCTCT-TGTGATCGCTGTC----------------------CTGGCTGGTACATGTTCCAATTCCACCGGATCCTGCAGTATGCGCTGCCTCGCCAGGAGAGTCAGCGGCCCTTCTTCTGGATATTTATGGACAATCTGTTGCTGACTGAGGATGACCAGGAGACAACTGTCCGCTTCCTTCAGACAGAGACTGTGACCCTCCAGTATGTCCGTGGCAGAGTCCTCCAGAATGCTGTGCGGGTGTGGAGCAACATTCCTGGGCTGAAGAGCAAGCACGCGGTCCTGACCCCACAGGAAGAAGAGACTCTGCAAGCCC------AAATCAGAACCAGAAACAAGCCG------------------GACACCCAG---AAAGTTGACCCCCTGGTGAAGAGCTGCCTTCTGCCCCT--------------GAGAGAGTACTTCA---AGTATTTTTCTCAGAACTCACTTCCTCTTTAG-------------------------------------------------------------------------------------------------------------------------------*

*>Rat3L*

*----------------------------------------------------------------------------------------------------------------------------------------------------------------------------------------------------------------------------------------------------------------------------------------------------------------------------------------------------------------------------------------------------------------------------------------------------------------------------------------------------------------------------------------------------------------------------------------------------------------------------------------------------------------------------------------------------------------------------------------------------------------------------------------------------------------------------------------------------------------------------------------------------------------------------------------------------------------------------------------------------------------------------------------------------------------------------------------------------------------------------------------------------------------------------------------------------ATGGGTTCCCGGGAGACACCTTCTTCCTGCTCTAAGACCCA---------TGAA-----------ACCTTGAACCTGGAGACTCC----------GGAGAGCTCTAGCACTGACCCTGACAGTCCCCTGGAAGAG---CA----ATGGCCG---------AAATCTTCCCCAGATCTGAAA----GAGGAAG---------------------------------------------------------------A---CAGCGTGGATATGGTACTG---GAAGACTCCAAGGAGCCTCTAACCCCTTCCTCACCG------CCGACAG--------------------------------------------------GCAGAGAGGTCATCAGGTACGAAGTCAACGTGAACCAGCGGAACATCGAAGACATCTGCCTCTGTTGCGGATCTCTCCAGGTGTACGCTCAGCACCCCTTGTTTGAGGGGGGAATTTGTGCCCCGTGTAAGG---------ACAAGTTCCTGGAGACCCTCTTCCTATA-CGACGAGGAT-GGACACCAGAGCTACTGTAC----CATCTGCTGCTCCGGGCAT-----------------ACCCTGTTCATCTGCGAGA----------------------------GCCCCGAC--------------TGTACCAGATGC-----------------------TACTGTTTCGAGTGTGTGGATATCCTGG--TGGGC--CCCGGGACCTCG-------GAGCGCATCAAT-----GCCATGGCCTGCTGGGTTTGCTTCCTGTGCCTGCCTTTCTCCCGGA---GCGGACTGCTGCAGAGGCGCAAGAAGTGGCGTCACCAGCTGAAGGCCTTCC------ATGACCGAGAGGGGGCA---------AGCCCTGTGGAGATATACAAGACTGTGTCTGCATGGAAAAGACAGCC------------AGTGAGGGTGCTGAGTCTTTTTGG---------------------------------------------------GAATATTGATA--AAGAACTAAAGAGTTT-------------------GGGCTTTTTGGAAAGCAGTTCTGGTTCTGAGGGAGGAACGCTGAAGTACGTGGAAGATGTCACCAATGTCGTGAGGAGAGAAGTGGAGAAATGGGGTCCCTTTGACTTGGTGTATGGCTCAA----------------------------------------CCCAGCCCCTAGGCTATTCT-TGTGACCGCTGTC----------------------CTGGCTGGTACATGTTCCAGTTCCACCGGATCCTGCAGTATGCCCGGCCTCGCCAAGACAGCCAGCAGCCCTTCTTCTGGATATTTGTGGACAATCTGCTGCTGACTGAAGATGACCAAGAGACAACTGTCCGCTTCCTTCAGACAGAGGCTGTGACCCTCCAGGATGTCCGCGGCAGAGTCCTCCAGAATGCTATGAGGGTGTGGAGCAACATTCCAGGGCTGAAGAGCAAGCACGCGGACCTGACCCCTAAGGAAGAGCAGTCTCTGCAAACCC------AAGTCAGAACCAGAAGCAAGCTG------------------GCCGCCCAG---AAAGTTGACTCCCTAGTGAAGTACTGCCTTCTCCCCCT--------------GAGAGAGTACTTCA---AGTATTTTTCTCAGAACTCACTTCCTCTTTAGAAATGA-------------------------------------------------------------------------------------------------------------------------*

*>CHamster3L*

*----------------------------------------------------------------------------------------------------------------------------------------------------------------------------------------------------------------------------------------------------------------------------------------------------------------------------------------------------------------------------------------------------------------------------------------------------------------------------------------------------------------------------------------------------------------------------------------------------------------------------------------------------------------------------------------------------------------------------------------------------------------------------------------------------------------------------------------------------------------------------------------------------------------------------------------------------------------------------------------------------------------------------------------------------------------------------------------------------------------------------------------------------------------------------------------------------ATGGGTTCCCGGGAGA---ATTCTTCTTGCTCCAAGGCCCT---------TGAA-----------ACTTGGGACCTGGAGACGCT----------GGACAGCTCTAGCCGGGACTCTGCCAGCCACCTGGAAGAG---CA----ATGGCGG---------AAATTCTCCCCCGTCCTCAAT----GAAGAGG---------------------------------------------------------------A---CAGCACGGATATTGTACTG---GAAGACTCCCGGGAGCTGCTGTCCTTAACTTCACCC------CCGCCAG--------------------------------------------------GCAGAGAGATCATCAGGTATGAAGTCAGTGTGAACCGACGGAACATCGAAGACATCTGCCTAGGTTGTGGAAGTCTCCAGGTGTACACGCAGCACCCCTTATTTGAGGGGGGGATATGTGCCTCGTGTAAGG---------ACAATTTCCTGGAAACCCTCTTCCTTTA-TGATGAGGAT-GGGCACCAGAGCTACTGTAC----CATCTGCTGCTCTGGGGAC-----------------ACCCTGTTCATCTGTGAGA----------------------------GCCCCGAC--------------TGTACCAGATGC-----------------------TACTGTTTCGAGTGTGTGGATATCCTGG--TAGGC--CCTGGGACCTCA-------GAACGAATCAAT-----GCCATGGCCTGCTGGGTTTGCTTCCTGTGCCTGCCTTTCGCTCGTA---GCGGACTGCTGCAGAGGCGAAGACAGTGGCGTCACCAGCTGAAGGCCTTCT------ATGATCTAGAAGGGGCA---------AGCCCTCTGGAGATGTACAAGACAGTGTCTGCGTGGAAGAGACAGCC------------CATGCGGGTGCTGAGCCTTTTTGG---------------------------------------------------GAATATTGATA--AAGAGCTAAAGAGTTT-------------------GGGCTTTTTGGAAAGTGGCTCTCGCACTGAGGAAGGAAGACTGAAGTACTTGGATGATGTCACAAATGTTGTGAGGAGAGACGTGGAGAGATGGGGTCCCTTTGACCTTGTGTATGGCTCAA----------------------------------------CACAGCCCCTAGGCTACTCT-TGTGACCACTGCC----------------------CTGGATGGTACATGTTCCAATTTCACCGGATACTGCAATACGCCCGGCCTCACCCAGGGAGTCAGCAGCCTTTCTTCTGGATATTTGTGGACAATCTGCTGCTGTCTGAGGATGACCAAGTCACAGCAGCCCGCTTCTTTCAGACGGAGGCTGTGACCCTCCAGGATGTCCGCGGCAGAGTCCTCCAGAATGCTGTGCGGGTATGGAGCAACATCCCAGGCTTGAAGAGCAAACACTCGGCCCTGACACCAAAGGAGGAGCAGTCCCTGGAAGGAC------ACGTTAGAACCAGAGCCAAGGTG------------------GCCGCCCAG---AAAACTGACGCCCTGGTGAAGAACTGCCTTCTCCCTCT--------------GAGAGAGTATTTCA---AGTATTTTTCTCAGAACCCACTTCCTCTTTACAAATGA-------------------------------------------------------------------------------------------------------------------------*

*>DeerMouse3L*

*----------------------------------------------------------------------------------------------------------------------------------------------------------------------------------------------------------------------------------------------------------------------------------------------------------------------------------------------------------------------------------------------------------------------------------------------------------------------------------------------------------------------------------------------------------------------------------------------------------------------------------------------------------------------------------------------------------------------------------------------------------------------------------------------------------------------------------------------------------------------------------------------------------------------------------------------------------------------------------------------------------------------------------------------------------------------------------------------------------------------------------------------------------------------------------------------------ATGGGTTCCCAGGAGA---ATTCTTCTTGCCCCAAGGCTCT---------TGGA-----------ACCTGGGACCTGGAGACTCT----------GGACAGTTCTAGCCCGGACTCTGCCAGCCACCTGGAAGAG---CA----GTGGCCG---------AAACCCTCCCCGAGCCTGAAT----GACGAGG---------------------------------------------------------------A---TAGCATGGATGTTGTACTG---GAAGACTCTGGGGAGATACGGTCCTTAGCCTCACCT------CCACCAG--------------------------------------------------GCAGAGAGATCATCAGGTATGAAGTCACTGTGAACCGACGGAACATCGAAGACATCTGCCTCTGTTGTGGAAGTCTCCAGGTGCATACGCAGCACCCCTTATTTGAGGGGGGCATATGTGCCTCGTGTAAGG---------ACACGTTCCTGGAGACCATCTTCCTTTA-TGATGAGGAT-GGGCATCAGAGCTACTGTTC----CATCTGCTGCTCAGGGCAC-----------------ACCTTGTTCATCTGTGAGA----------------------------GCCCCGAC--------------TGTACCAGATGT-----------------------TACTGTTTTGAGTGTGTGGATATCCTGG--TGGGC--CCCGGGACTTCA-------GAGCGAATCAAC-----GCCATGGCGTGCTGGGTTTGCTTCCTGTGCCTGCCTTTCACTCGGA---GCGGGCTGCTGCAGAGGCGAAAGAGGTGGCGTCACCAGCTGAAGGCCTTCC------ATGATCTAGAGGGGGCA---------AGCCCTCTGGAGATGTATAAGCCAGTGTCTGCTTGGAAGAGACAGCC------------AGTGCAGGTACTGAGCCTCTTTGG---------------------------------------------------GAATATTGAGA--AAGAGCTAGAGAGTTT-------------------GGGCTTTCTGGAAAGAGGTTCTGGTTCTGAAGGAGGAAGGCTGAAGTACTTGGAAGATGTCACGAATGTTGTGAGGAGAGATGTGGAGAGATGGGGCCCCTTTGACCTTGTATATGGCTCAA----------------------------------------CACAGCCTCTAGGTCGCTCT-TGTGACCGCTGCC----------------------CTGGCTGGTACATGTTCCAGTTCCACCGGATCCTGCAGTATGCTCGCCCACACCCAGGGAGTCAGCGGCCTTTCTTCTGGCTGTTTGTGGACAGTCTGCTGCTGACTGAGGATGACCAAGTCACCACAACCCGCTTCCTCCAGATGGAGCCTGTGACCCTCCAGAATATTCGCGGCAGAGTCCTCCAGAATGCTGTGCGGGTGTGGAGCAACATCCCAGGGTTGAAGAGCAAACACTTGGCCCTGACACCAAAGGAGCAACAGTCCTTGGAAGGCC------AAGTCAGAACCAGAGCCAAGATG------------------GCCTCCCAG---AAAGACGACCCCCTGGTGAAGAACTGCCTTCTCCCCCT--------------GAGAGAGTATTTCA---AGTATTTTTCTCAGAACCCACTTCCTCTTTACAAATGA-------------------------------------------------------------------------------------------------------------------------*

*>PVole3L*

*----------------------------------------------------------------------------------------------------------------------------------------------------------------------------------------------------------------------------------------------------------------------------------------------------------------------------------------------------------------------------------------------------------------------------------------------------------------------------------------------------------------------------------------------------------------------------------------------------------------------------------------------------------------------------------------------------------------------------------------------------------------------------------------------------------------------------------------------------------------------------------------------------------------------------------------------------------------------------------------------------------------------------------------------------------------------------------------------------------------------------------------------------------------------------------------------------ATGGGTTCCCGGAAAA---ATTCT---TGCTCCAAGACAGG---------TGAA-----------ACATGGGACCTGGAGACTCT----------GGACAGCTCTAGCCCGGACTCTGCCAATCACCTGGAAGAG---CA----ATGGCTG---------AAATCCACCCGTAGCCTCAGT----GACGATG---------------------------------------------------------------AGCACAGCGTGGAGGTCGTACTG---GAAGATTCCCGGGAGCTGCTGTCCTTAACCTCACCC------CCGCCAG--------------------------------------------------GCAGAGAGATCATCAGGTATGAAGTCACTGTGAACCAACGGAACATCGAAGACATCTGCCTCTGTTGTGGAAGTCTCCAGGTGTATACGCAGCACCCCTTATTTGAGGGGGGGATGTGTGCCCCATGTAAGG---------ACATATTCCTGGAGACCATCTTCCTTTA-CGATATGGAT-GGGCACCAGAGTTACTGCTC----CATATGCTGCTTCGGGAAA-----------------ACCCTGTTTATCTGTGAGA----------------------------GCCCCGAC--------------TGTACCAGATGC-----------------------TACTGCTTTGAGTGTGTGGATATCCTGG--TGGGT--CCCGGGACCTCA-------GAGCGAATCAGC-----ACCATGGCCTGCTGGGTTTGCTTCCTGTGCCTGCCTTTCACTCGGA---GCGGACTGCTGCAGAGGCGAAGGAAGTGGCGTCACCAGCTGAAGGCCTTCC------ACGATCTCGAGGGGTCA---------AATCCTGTGAAGATGTACAAGATAGTGTCTGCCTGGAAGAGACAGCC------------CGTGAGGGTGCTTAACCTTTTTGG---------------------------------------------------GAAGATTGATA--AAGAGCTCGAGAGTTT-------------------GGGCTTTCTGGAAAGTGGTTCTGGTTCGGAAGAGGGAAGACTGAAGTACTTGGAAGACGTCACCAATGTTGTGAGAAGAGATGTGGAGAGATGGGGCCCCTTTGACCTCGTGTATGGCTCGA----------------------------------------CACAGCCCCTAGGCTACTCT-TGTAACCGCTGCC----------------------CTGGCTGGTACATGTTCCAATTCCACCGGATCCTGCAGTATGCTCAACCACCCTCAGGGAGTCAGCAGCCTTTCTTCTGGCTCTTTATGGACAATCTACTGCTGACAGAGGATGACCAGGCCACGGCAACCCGCTTCTTCCAGGTGGAGGCTGTGACCCTCCAGGATGTCCGCAGCAGAGTCCTCCAGAATGCTGTGCGGGTGTGGAGCAACATCCCAGGGTTGAAGAGCAAACACTCGGCCCTAACACCTAAGGAGCTGCAGTCCCTGGAAACCC------AGACCAGAACCAGAGGCAAGATG------------------GCTGCCCAG---AAAGTTGACCTCCTGGTGAAGACCTGCCTTCTCCCCCT--------------GAGAGAGTATTTCA---AGTATTTTTCTCAGAACCCACTTCCTCTTTACAAATGA-------------------------------------------------------------------------------------------------------------------------*

*>MBMole3L*

*----------------------------------------------------------------------------------------------------------------------------------------------------------------------------------------------------------------------------------------------------------------------------------------------------------------------------------------------------------------------------------------------------------------------------------------------------------------------------------------------------------------------------------------------------------------------------------------------------------------------------------------------------------------------------------------------------------------------------------------------------------------------------------------------------------------------------------------------------------------------------------------------------------------------------------------------------------------------------------------------------------------------------------------------------------------------------------------------------------------------------------------------------------------------------------------------------ATGGGTTCCTGGGAGGCACTGTCCTCTTGTTCCAAGGCCCTCTCCTCCCCTGGA-----------ACATGGAACTTGGAGACTCC----------AGACAACTCGAGCCCTGACCCAGCCGGCCACCTGGAAGAA---CA----GAAGCCC------------TCACCTCCACCTCTGAAG----GCCGAGA---------------------------------------------------------------A---CAACATGGATGTAGTACTG---GTGGACTCCAGGGAGCTGCTGTGCCCCACTTCACCC------CCTCCAG--------------------------------------------------GTAGAGATCGCATCGCCTATGAAGTGACCGTCAACCACCGGAACATAGAAGAGCTCTGCCTGTGCTGTGGAAGTCTCCAGGTGTACACACAGCACCCCCTGTTCCAGGGAGGAATGTGTGCCCCATGTAAGG---------ACAAATTCTTGGAGACCCTCTTCCTGTA-TGACGATGAC-GGGTATCAGTGTTCCTGCTC----CATCTGCTGCTCGGGGGAC-----------------ACGCTGTTCATCTGCGAGA----------------------------GTCCAGAC--------------TGTACCCGGTGC-----------------------TACTGTTTTGAGTGTGTGGATGTCCTGG--TGGGC--CCTGGGACCTCA-------GAGCGAATCCAC-----GCCATGAACTGCTGGGTGTGCTTCCTGTGCCTGCCTTTCTCTCAAA---GCGGGCTGCTCCTGAGGCGGAGGAAGTGGCGCCACCGGCTGAAGGCCTTCT------ATGACCTCGAGGCGGCA---------AGCCCTCTGGAAATGTACAAAACAGTGCCTGTGTGGAAGAGAGAGGC------------AGTGCGGGTGCTGACCCTTTTTGA---------------------------------------------------AGACATTCAGA--AAGAACTAAGGAGCTT-------------------AGGCTTTTTGGAAAGTGGTTCCGGTTCTGAGAGAGGAAGACTGAAACACTTGGATGATGTCACAAACGTAGTGCGAAGAGATGTGGAGGGATGGGGCCCCTTTGACCTTGTGTATGGCTCAA----------------------------------------CACAGCCCCTGGGCTATTCC-TGCAACCGTTGTC----------------------CTGGCTGGTATCTGTTCCAGTTTCACCGGATCCTGCAGTATGCACGGTCACACCGGGGCAGCCAGCAACCCTTTTTCTGGATGTTTGTGGACAACTTGCTGCTGACTGAGGCTGACCAGGCCACAGCAGCCCGCTTCCTTGAGATGGAGCCCATGACCCTCCAGGACATCCGAGGCAGAGTCCTGCAGAACGCGGTTCACGTGTGGAGCAACATCCCAGGGGTGAAGAGCAAGCACATGGCCCTGACAGCCGAGGAGGAACAGTTTCTGCAGGCCC------AAGGTGATGCCCGAGCCAAGCTG------------------GTTGCCAAG---GGACTGGCCCCGCTGGTGAAGAAGTGCTTTCTCCCCCT--------------TAGAGAGTATTTCA---AGTATTTTTCTCAGAACTCACTTCCTCTTTACAAATGA-------------------------------------------------------------------------------------------------------------------------*

*>Beaver3L*

*---------------------------------------------------------------------------------------------------------------------------------------------------------------------------------------------------------------------------------------------------------------------------------------------------------------------------------------------------------------------------------------------------------------------------------------------------------------------------------------------------------------------------------------------------------------------------------------------------------------------------------------------------------------------------------------------------------------------------------------------------------------------------------------------------------------------------------------------------------------------------------------------------------------------------------------------------------------------------------------------------------------------------------------------------------------------------------------------------------------------------------------------------------------------------------------------------------------------------------------------------------------------------------------------------------------------------------------ATGG---TA----GCACCC-------------CCAGACTCAGACCTGGAG----CCTGAGC---------------------------------------------------------------G---CAGCATGGACGTGATCCTG---GTGGGCTCCACCGAGCTGCTGCCCTCCCCTCCAACT------CCACCAG--------------------------------------------------GCAGAGATCTCATTGCCTATGAAGTCACGGTGAAACAGCGGAACATAGAAGACATCTGCCTCTGCTGTGGAAGTTTCCAGGTGCAGACACAGCACCCCCTGTTTGATGGAGGAATGTGCGCCCCGTGCAAGG---------ACAAGTTCCTGGAGAGCCTCTTCCTGTA-TGACGATGAT-GGGTACCAGTGCTACTGCTC----CATCTGCGGCTTGGGGGAT-----------------ACACTGCTTATCTGTGAGA----------------------------GCCCTGAC--------------TGCACCCGATGC-----------------------TACTGTTTTGAGTGTGTGGACATCCTGG--TGGGC--CCCGGGACCTCG-------GAGCGAGTCCAA-----GGCATGAGCTGCTGGGTGTGCTTCCTGTGCCTGCCTTTCTCTCGCA---GTGGGCTGCTGCAGAGGAGGAGGAAGTGGCGCGACCGGCTAAAGGCCTTCC------ATGACCGCGAGGCGGAG---------AGTCCCCTGGAGATGTACAAAACAGTGCCTGTGTGGAAGAGAGAGCC------------AATGCGGGTGCTGTCCCTTTTTGG---------------------------------------------------GGACATTAAGA--AAGAGTTAACGAGTTT-------------------GGGCTTTTTAGAAAATGGTTCCGGTTCTGAT---GGAAAACTGAAGCACTTGGACGACGTCACAAACATAGTGCGGAGAGATGTGGAGGAGTGGGGCCCCTTCGATCTCCTGTACGGCTCGA----------------------------------------CACACCCCCTAGGCCATACC-TGTGACCGTTCTC----------------------CTGGCTGGTACATGTTCCAGTTCCACCGGCTGCTGCAGTACGCACGGCCCCGGCTGGGCAGCCAGCAACCCTTCTTCTGGATGTTCGTGGATAACCTGCTGCTGACCCAGGATGACCAGGCCACAGCAACACGCTTCTTTGAGATGGAGCCCGTGACCCTGCAGGACGTCCGTGGCAGGGTCCTCCATAACGCCGTACGTGTGTGGAGTAACATCCCAGCTGTGAAGAGCAAACATGAGGCTCTGGACCCTGAAGAGGAACTGTCACTGCTGTCCC------AAGCCACACAGAAAGCAAAGCTG------------------GCCACCCAG---AGACCGGCCACACTTGTGAAGAACTGCTTTCTTCCACT--------------AAGAGAATATTTCA---AGTATTTTTCTCAGAATTCAGTTCCTCTTTACAAATGA-------------------------------------------------------------------------------------------------------------------------*

*>Squirrel3L*

*----------------------------------------------------------------------------------------------------------------------------------------------------------------------------------------------------------------------------------------------------------------------------------------------------------------------------------------------------------------------------------------------------------------------------------------------------------------------------------------------------------------------------------------------------------------------------------------------------------------------------------------------------------------------------------------------------------------------------------------------------------------------------------------------------------------------------------------------------------------------------------------------------------------------------------------------------------------------------------------------------------------------------------------------------------------------------------------------------------------------------------------------------------------------------------------------------CCCACTTACTGGGAGGCGCTGTGGCCCGACCTCATGGCCCTCTCCCGCTCTGGG-----------ACCCTGACTCCAGACAGCCT----------GG---GCTCTGACCCGGACCCAGACAGACTCCTGGACGGGGCGCA----GTGGCAG---------CAGCGCTTCCTGGACCTGGAG----GCTGAGC---------------------------------------------------------------A---CAGCATGGATGTCATCCTG---GTGGGCTCCAGCGGGCGGTCATCCCCCTCTTCAGTC------CAGCTGG--------------------------------------------------GCAGAGACCTCATTGCCTATGAAGTCACTGTGAACCAGCGAAACATGGAAGACATCTGCCTCTGCTGTGGAAGCTTCCAGGTGCACACGCAGCACCCTCTGTTTGAGGGGGGGATCTGTGCCCCGTGCAAGG---------ACAGGTTCCTGGAGGCCCTGTTCCAGTA-CGACGAGGAC-GGGTACCAGTCCTCCTGCTC----CATCTGCGGCTCAGGGGAG-----------------ACGCTGCTCATCTGCGAGA----------------------------ACCCGGAC--------------TGCACACGGTGC-----------------------TACTGTCTGGAGTGTGTGGACACCCTGG--TGGGC--CCAGGGACCGCA-------GGCAGAATCCAC-----GCCATGAGCAGCTGGGTGTGCTTCCTGTGCCTGCCCTTCTCGCGCA---GTGGCCTGCTGCAGCGCAGGAGGAAGTGGCGTGAGCGGCTGAAGGCCTTCC------AGGATCGCGAGGTGGCG---------AGTCCCCAGGAGATCTATAAAACCCTGCCAGCCTGGAAAAGGGAGCC------------AGTGCGGGTGCTGTCCCTCTTTGG---------------------------------------------------GGACATCGGGA--AAGAGCTAACCAGTCT-------------------GGGGTTTCTGGAA------CCCGGCTCCGAGGCGGGAAGACTGAGGCACCTGGAGGACGTCACCGACATTGTGCGCAGAGATGTGGAGGAATGGGGCCCCTTCGACCTCGTGTACGGCTCGA----------------------------------------CGCCCGCCCTGGGCCACGCC-TGTGACCACTCTC----------------------CTGGGTGGTACCTGTTCCAGTTCCACCGCCTCCTGCAGTACGCGCGGCCGCGGCCGGGCAGCCCCCAGGCCTTCTTCTGGATGTTTGTGGACAACCTGCAGCTGACCGGGGAAGAGCAGGCCATAGCCGCTCGCTTCCTGGAGACGGAGCCCGTGATCCTCCAGGACGTCCGCGGGAGCGCCCTGCAGAATGCCGTGCGCGTGTGGACCAACATCCCCGCTGTGAAGAGCAGGCACTCGGCCCTGGCTTCAGAAGAGGAGCTCCTGCTGCTGGCTC------AAGATGGACAGCGAGGGACGCTG------------------CCCGCCCAG---GGCCCATCTGCGCTCGTGAAGAACTGCTTTCTTCCCCT--------------ACGAGAATATTTCA---AGTATTTCTCTCAGAACGCACTTCCTCTTTATAAATGA-------------------------------------------------------------------------------------------------------------------------*

*>Pika3L*

*-------------------------------------------------------------------------------------------------------------------------------------------------------------------------------------------------------------------------------------------------------------------------------------------------------------------------------------------------------------------------------------------------------------------------------------------------------------------------------------------------------------------------------------------------------------------------------------------------------------------------------------------------------------------------------------------------------------------------------------------------------------------------------------------------------------------------------------------------------------------------------------------------------------------------------------------------------------------------------------------------------------------------------------------------------------------------------------------------------------------------------------------------------------------------------------------------------------------------------------ATGGCTCT--------------------------CCCCTCCCCGGAGACCCT----------GGACAGTCTGGATCGTGTCCCCGCCAGCCACCCGGACGAGCAGCA----CTGGACAGTCTGTGACAACTCTGACCCCATCCTGGAG----GTGGAAGC------------------------------------------------------------TGAGGGCAGCATGGATGTGATCCTG---GTGGACGAC------------------TCTCCAGCC------CCCTCAG--------------------------------------------------GCAGAGACCGTATTGAGCTCGAGGTCAAAGTGAACCAGCGGAGTATAGAGGACCTCTGCCTGTGCTGCGGGAGCTCCCAGGTGCACAGGCAGCACCCCCTGTTCCAGGGCGGCCTGTGCGCGCCTTGCAAGG---------ACAAGTTCCTGGAAGCACTCTTCCTGTA-CGATGAGGAC-GGCTACCAGTCCTACTGCTC----CATTTGCGGCCTGGGGGAC-----------------ACCCTGCTCGTTTGCGAGA----------------------------GCCCGGAC--------------TGCACCCGGGGC-----------------------TATTGTTTCGCCTGTGTGGATGGCCTGG--TGGGC--GCTGGGAGCTCA-------GGGCACATGCAC-----ACCGTGAGCCCCTGGGTTTGTTTCCTGTGTGTGCCGGGCTCCCGCC---ATGGGCTCCTGCAGCGCCGGCGGAGGTGGCGGACCCAGCTGAAGGTCTTCC------ATGAGCAGGAGGCGGCC---------CAGCCCCTTGAGATATATGAGACGGTGCCAGCGTGCAGGAGGAAGCC------------ATTAAGGGTGCTGTCCCTTTTTGA---------------------------------------------------GCACATCGAGA--AAGAGCTGGCAAGCCT-------------------GGGCTTCCTGGAGACCGGGTCCAGCCC------GGGCCGAATAAGGCATCTGGACGATGTGACGGACGTAGTGAGGAGGGACGTGGAGCAGTGGGGCCCTTTTGACCTGGTGTATGGCTCAA----------------------------------------CCCCTCCGCTCGGCCACGCC-AGTCCACGGTCAC----------------------CGGGCTGGTACCTGTTCCAGTTCCACCGCATGCTGCAGTACACACAGCCCACTGCGAGCACACAGCGGCCCTTCTTCTGGATGTTCGTGGACAATCTGCTGCTGACCAGGGATGACCTGGTCACGGCGACCCGCTTCTTGGAGGTAGAGCCGGCGACCCTGCAGGACGTTCGTGGCCGTGTCCTCCAGGGCGCCATGCGTGTCTGGAGCAACATCCCGGCCGTGAACAGCAGGCACACGGAGCTGGCTCCGGAGGCGGAGACGGCCCTGCTGGCGC------AGAGCTGTCGGCGAGCAAAGGCC------------------TCGGGCGAG---GGGCTGGCCAGGCTGCTGAAGAGCTGCTTCCTCCCGCT--------------CAGAGAATATTTCA---AGTATTTTCCCCAGAGCCCACTTCCTCTCCGCAAATGA-------------------------------------------------------------------------------------------------------------------------*
