## Supplementary material for "Dynamic evolution of *de novo* DNA methyltransferases in rodent and primate genomes": Suppl data 2

*SupFile_2: Muroid* Dnmt3C *sequence alignments*

>ChineeseHamster_DNMT3C_ORF

ATGAAGAGAAACGGCAGACACCTCAGTGATGAAGAGGGTGTCAGTGGGTGTGAGGACTGTGTCATCATCACTGGGACCTGCAGTGACCAGTCCTCAGACGCCAAGGATGTCACCTTGACCCAAGTCTTGGAGGCATTCTGCCCAGTGGAGAACAG-TGGCTGCAGAGCAAGCTCACGACGGTTCAAGAGGAGGGTCTC-CAACACGGTTACTTACTTTCAGGACCTGACAGGAGATGAAGAT---------GATGTCGAGGTGGAGGGCAGCAGTGGCTCTTGCAGTCCAGCGATGCCCAAATTCTTCTGG---------GAGACCAGGACACCCTCTAAAACCCCAGCACCCATTTACCAGCAAGCAACCACTTCAGCAAGCACACCGTGGCTGTTCCCTACCAGCCCTTACCCAACCATTGACCTCACAGATGAAGAAGTGATACTCCAGAGCATCAGTACCCCACTGGTTGACTTGAGCCAGGACAGCCATCAGGAAAGCATGGATACCACACAGGTAGATGTGGGAAGCAGAGATGAAGATACCACCAAGTATC------------------------------------------------------------------------------------------------AGAGAGCCAGGGTGCGAGCTGGCAAGAC-------CTTCCCCAGCAGCCC--------TGGAGACTCACTGGAGGACAAGCTGAAGCCCATGCTGGAGTGGGCCCACGGGGGCTTCAAGCCCATTGGCATCGAGGGCCTCAAACCAAACAAG------------------------------------------------------------------------------------------------------------------------------------------------------------------------------------GACCGCTGCTTGTCCTGTGGTAGGAAGAACCCTGTATCCTTCCATCCCCTCTTTGAGGGTGGGCTCTG-TCAGAGTTGCCGGGATCGCTTCCTGGAGCTGTTCTACATGTACGA----TGAGGACGGCTATCAGTCCTACTGCACTGTGTG----------CTGCAAGGGCCGCGA--GCTGCTTCTGT--GCAGCAACACAAGCTGCTGCAGGTGCTTCTGTGTGGAGTGTCTGGAGGTGCTGGTGGGTACAGGGACAGCTGAGGATGCCAAGCTGC-----AGGAACCCTGGAGCTGCTATATGTGCCT---TCCACAGCGCTGCCATGGGGTCCTCA---GGCGCAGGAAGGATTGGAACATGCGCCTGCAAGACTTC---TTCACTTCTGATCCTGACCTCGAAGAA------GAGCCACCCAAGTTGTACCCAGCAATTCCTGCAGCCAAGAGGAGACCTA--TTAGAGTCCTGTCCCTGTTTGATGGAATTGCAACAGGGTACTTGGTGCTCAAAGAGCTGGGTATCAAAGTGGAAAAGTAT----------GTTGCGTCTGAAGTGTGTACAGAGTCCATTGCTGTGGGAACTGTTAAGCATGAAGGTCAAATCAAATATGTGGATGACGTCAGGAACATCACAAAGGGAAATATTGATGAGTGGGGCCCATTCGACTTGGTGATTGGTGGAAGCCCTTGCAATGATCTCTCCTGTGTGAATCCTGGCAGGAAAGGGCTGTTTGAGGGCACTGGTCGGCTCTTCTTTGAATTTTATCGATTGCTAAATTACGCACGTCCCGAGGAGTGTGATAACCGCCCATTCTTCTGGATGTTTGAGAATGTGGTAGCTATGGAGGTTGGTGACAAGAGGGACATCTCACTATTCCTGGAGTGTAACCCAGTGATGATCGATGCCATCAAGGTTTCTGCTGCACACAGGGCCCGATATTTCTGGGGCAACCTACCTGGGATGAACAGGCC----TGTGATAGCTTCCAAGAATGATAAGCTCGAGCTGCAGGACTGCTTGGAGTTCAGTAGGACAGCAAAGTTAAAGAAAGTACAGACAATAACCACC-----------AGGTCGAACTCCATCAGACAGGGGAAAAACC---------AGCTTTTCCCTGTAGTCATGAATGGCAAGGACGATGATTTGTGGTGTACTGAGCTTGAAAAGATCTTTGGCTTTCCTTCACACTACACGGATGTGTCTAACATGGGCCGTGGTGCCCGCCAGAAGCTGCTGGGAAAGTCCTGGAGTGTGCCAGTCATCAGACACCTCTTTGCCCCGCTGAAGGACTACTTTGCATGTGAA---

>MountainBlindMole_DNMT3C_ORF

ATGACTGGAGACAGCAAATACCTCAGTGAGGAAGAGGGTGCCAATGGGTGTGAGGACTCCATCATCATCATTGGGAACTTCAGTGACCAGTCCTCAGACACCAAGGATGTTCCCTCACCCCCTGCCTTGGAGGCGGTCTGCACACTGGAAGGCAG-AGGCTGCAGATCAAGCTCATGGCTGTCCAAGAGGAAGGTCTC-CAGGCTGCTAAGTTACAATCAGGATCTGATGGGAGATGGAGAT---------GATGGTGAGGTAGAGGACAGGGTTGGAGCTGATACTCCAGTGATGCCAAAACTCTTCTGG---------GAGACTGGGACACCCTCTAAAACCCCAGCATCCATGAGGCGGAGGGCAAGAACATCAGCAGGCATGCTGTGGCCATCCCCTTCCAGCCCCTACCCCACCATCGACCTCACAGATGAAGAAGTGACACCTGGGAGCAGCAGGACAGTTTATGTTGACCTAGACGAGGACAGCCAGCAGGAGAGCATGGACTCCACACAGGTGGATACAGAAAGCAGAGATGGAGACAGCACTGAGTATC------------------------------------------------------------------------------------------------AGAAAGCCAGGGTTCGAGCTGGCAAGAC-------CTTCCCCACCAGCCC--------TGCAGACTCACTGGAGGACCAGCTGAAGCCTATATTGGAGTGGGCCCATGGAGGCTTCAAGCCCACTGGGATTGAGGGCCTCAAACCCAACAGC------------------------------------------------------------------------------------------------------------------------------------------------------------------------------------GACAGCTGTTTGTCTTGTGGAAGGAAAAACCCCGTGTCCTTCCACCCACTCTTTGAGGGTGGGCTTTG-CCAGACATGCCGGGATCGCTTCCTTGAGCTGTTCTACATGTACGA----CGACGATGGCTATCAGTCCTACTGCACCGTGTG----------CTGCGATGGCGGTGA--ACTACTTCTCT--GCAGCAACATAAGCTGCTGCCGGTGCTTCTGTGTCGAGTGTCTGGACGTGTTGGTGGGCACAGGCACGGCTGCAGATGCCAAGCAGC-----AGGCGTTCTGGAGCTGCTACATGTGTGT---GCGAAGGTGCAGCCATGGTTTTCTGC---GGCGCCGGAAGGACTGGAACTTGCGCCTGCAGGCCTTC---TTCTCCAGCATTCTTGACATTGAATAT------GAGGCCCCCAAGTTGTACCCAGCAATTCTTGCAGCCCAAAGGCGGCCCA--TTAGAGTCCTGTCCCTGTTTGATGGAATTGCGACAGGCTACTTGGTCCTCAAAGAATTGGGTATTAAAGTGGACAGGTAC----------GTCGCCTCTGAAGTCTGCGCAGAATCCATCGCTGTGGGAACCGTGAAGCATGAAGGGCAAATCAAATACGTGAATGACGTCAGGAAAATCACAAAGAGAAATATTGATGAGTGGGGCCCATTCGACCTGGTGATTGGTGGAAGCCCATGCAATGATCTTGCCTCTGTGAATCCTGTCAGGAAAGGCCTGTTTGAGGGTACTGGCCAACTCTTCTTTGAGTTTTACCGCTTGCTAAATTACTCACGCCCCAGAGAGAGTGATGACCGTCCATTCTTCTGGATGTTTGAGAATGTGGTAGCCATGAAGATCTGTGACAAGAGGGACATCTCACGGTTCCTGGAGTGTAACCCAGTGATGCTTGATGCCATCAAGGTTTCCGCTGCTCACAGGGCCCGATACTTCTGGGGCAACCTACCTGGGATGAACAGNNN----NNTGATAGCATCAAAGAATGATAAACTCGAGCTGCAGGACTGCTTGGAGTTCAGTAGGACAGCAAAGTTGAAGAAAGTACAGACAATAACCACC-----------AAGTCGAACTCGATCAGACAGGGGAAAAACC---------AACTTTTCCCTGTAGTCATGAATGGCAAAGAAGATGTTTTGTGGTGCACTGAGCTCGAAAGGATCTTCGGCTTTCCTGTACACTACACAGATGTGTCCAACATGGGCCGTAGTGCCCGCCAGAGGCTGCTGGGAAGGTCCTGGAGTGTGCCCGTCATCAGACACCTCTTCGCCCCCTTGAAGGAATACTTTGCATGTGAA---

>Rnovergicus_DNMT3C_ORF

ATGAATGGAGGTAGCAGACTACTCAGTGATGAGGAGGGTATCAGTGGATGCCAGGACTGTATCAGCATCAGTGGGACCTGCAGTGACCGGTCTTCAGACACCAAGACTCTTCCCTTGACCCAAGTCTTGGAGGTACTCGGCACAGTGGAGAGCAG-AGGCTGCAGAACAGGCTCACCACCATCCGAGAGAAAAGTCTC-CAGCCTGATTAGTTACATTCAGGACCTCACGGGAGATGGAGAT---------GATGGTGAGGTCGGGGACAGCAGTGGCTCTGACACTCCAGTGATGCCCGAACTCTTCTGG---------GAGACCAGGACACCTTCTCAAGCCCCAGCAGCTTCCAGTTGGCCAGCAACCACTTCAGCCAGCTCACCCTGGCCGTCCCCTGCCAGCCCTTACCCCATCATTGACCTCACAGATGAAGATGTGATACCCCAGAGCATCAGCACCTCATCCAGTGACTTGAGCCTGGACGGCCCTCAGGAGGACATGGATACCACACAGGTGGATGTAGAGAGCAGAGACGGGGACAGCCCTGAGTATC------------------------------------------------------------------------------------------------AGAAAGCCATGGTGCGAGCTGGCAAGAC-------CTTCCCCAGCCGCCC--------TGGAGACTCACTGGAGGACCAGCTGAAGCCCATGCTGGAGTGGGCCCACGGGGGCTTCAAGCCCACTGGGATCGAGGGCCTCAAACCCAACAAC------------------------------------------------------------------------------------------------------------------------------------------------------------------------------------GACCACTGTTTGTCCTGCGGTAGGAAGAACCCTGTGTCCTTCCACCCCCTCTTTGAGGGTGGGCTCTG-TCAGAGTTGCCGGGATCGCTTCCTGGAGCTCTTCTACATGTACGA----TGAGGACGGCTATCAGTCCTACTGCACCGTGTG----------CTGTGAGGGCCGTGA--GCTGCTTCTGT--GCAGTAACACAAGCTGCTGCAGGTGCTTCTGTGTGGAGTGTCTGGAGGTGCTGGTGGGTACAGGCACAGCGGAGGATGCCAAGCTGC-----AGGAACCCTGGAGCTGCTATATGTGCCT---CCCGCAGCGCTGCCATGGGGTCCTGC---GGCGCAGGAAGGACTGGAACATGCGCCTGCAAGACTTC---TTCACTACTGATCCTGACCTGGAAGAG------GAGCCGCCCAAGTTGTACCCAGCGATTCCTGCAGCCAAAAGGAGGCCCA--TTAGGGTCCTGTCCCTGTTTGATGGAATTGCAACAGGGTACTTGGTGCTCAAGGACTTGGGTATTAAAGTGGAGAAGTAC----------GTTGCCTCCGAAGTCTGTGCAGACTCCATTGCTGTAGGAACCATTAAGCATGAAGGACAAATCAAATACGTGGATGACATCCAGAACATTGCAAAGGAACATATTGACGAGTGGGGCCCGTTCGACCTGGTGATTGGTGGAAGCCCATGCAATGATCTTTCCTGTGTGAATCCCATCAGGAAAGGCCTGTTTGAGGGCACTGGCCGGCTCTTCTTTGAGTTTTACCGATTGCTAAATTACTCACGTCCTGAGGAGGAGGATGACCGCCCATTCTTTTGGATGTTTGAGAATGTGGTAGCCATGGAAGTTGGTGACAAGAGGGACATTTCACGATTCCTGGAGTGTAACCCAGTGATGATCGATGCCATCAAGGTTTCTGCTGCTCACAGGGCCCGATACTTCTGGGGCAACCTACCTGGAATGAACAGGCC----TGTGATAGCTTCAAAGAATGATAAGCTCGAGCTACAAGACTGCTTGGAGTTCAGTAGGACAGCAAAGTTAAAGAAAGTGCAGACAATAACCACC-----------AAGTCGAACTCCATCAGACAGGGGAAAAACC---------AGCTTTTCCCTGTAGTCATGAACGGCAAGGATGACGTTCTGTGGTGCACTGAGCTCGAAAGGATCTTCGGCTTCCCGGCTCACTACACGGATGTGTCCAACATGGGCCGTAGTGCCCGCCAGAAGCTGCTGGGAAAGTCCTGGAGTGTGCCAGTCATCAGACACCTGTTTGCCCCCTTGAAGGACTACTTTGCATGTGAA---

>PrairieVole3C

ATGAAGGGAGACAGCAGACATCTGAATGAAGAAGAGGGTGCCAGTGGGGGTGAGGAAAGTGTCATTGTCAATGGGAACTGTAGTGACCATTCCTCAGACACTAAGGATGCTCCCTCACCCCCAGTCTTGGAGGCAATCTGCACAGAGGCTGACAGCACACTGGAGAGC-----------------AGAGGTGAGTATCATAGCTCTGACACTACAGGCTCCGACCTGACAGGAAATGGAGAT------------GGTGAGGCAGAGGATGGGGATGGCGATGATGTTCTACCGATGCCAAAGCTCACACGTGAGACCAAAGAGACCAGGTCACCCTCGGAAAACCCAGCTTCTCGGAGGCGTCGGGCATCATCGTCAGCAAGCACGCCATGGTCGTCCCCTGCCAGCCCTTCCTTCAT---------------GGAAGAAGGGACACCTCAGAACAGCAGTATCCCATCAATTGACTTGACCCAGGACATCAATCAGGAGAGCATGGACACTACACAGCTGGAGGCAGAAGGCAAAGATGGAGACAGCACTGAGTATC------------------------------------------------------------------------------------------------AGAGAGCCAGGGTGCGAGCTGGCAAGAC-------CTTCCCCAGCAGCCC--------GGGAGATTCCCTGGAGGACCAGCTGAAGCCCATGCTGGAATGGGCCCATGGGGGCTTCAAGCCCACTGGCATCGAGGGCCTCAAACCCAACAACAAGCAACCA---GAGAACAAAAGTCGAAGACGCACAACCACCGACTCTGCCTCTGCTGAGTACTGTCCCCCACCCAAGCGCCTCAAGACCAATAGCTATGGCGGGAAAGACCGAGGGGAGGATGATGAGAGCCGAGAACGGATGGCTTCTGATGTTGCCAATAACAAAGTCAGTCTAGAAGACCGCTGTTTGTCCTGTGGTAGGAAGAACCCTGTGTCCTTCCACCCCCTCTTTGAGGGTGGGCTCTG-TCAGAGTTGCCGGGACCGCTTCCTGGAGCTCTTCTACATGTACGA----TGAGGACGGCTATCAGTCCTACTGCACCGTGTG----------CTGCGAGGGCCGCGA--GCTGCTTCTGT--GCAGCAACACAAGCTGCTGCAGGTGCTTCTGTGTGGAGTGTCTGGAGGTGCTGGTGGGTACAGGGACAGCTGAGGATGCCAAGCTGC-----AGGAACCCTGGAGCTGCTATATGTGCCT---CCCACAGCGCTGCCATGGGGTCCTCA---GGCGCAGGAAGGATTGGAACACGCGCCTGCAAGACTTC---TTCACTACTGACCCTGATCTCGAGGAATTT---GAGCCGCCCAAGGTGTTCCCAGCAATTCCTGCAGCCAGGAGGAGACCCA--TTAGAGTCCTGTCCCTGTTTGATGGAATTGCGACAGGGTACTTGGTGCTCAAAGACTTGGGTATCAAAGTGGAAAAGTAC----------GTTGCCTCCGAAGTCTGTGCCGAGTCCATCGCCGTGGGAACTGTTAAGCATGAAGGCCAAATCAAATATGTGAACGACGTCAGGAAAATCACAAAGAAAAACATTGAAGAGTGGGGCCCTTTTGACTTGGTGATTGGTGGAAGCCCTTGCAATGACCTCTCCAATGTCAATCCTGCCAGGAAAGGTCTATATGAGGGTACCGGCCGGCTCTTCTTTGAGTTTTACCACTTGCTGAATTATACACGCCCCAAGGAGGGCGACAACCGTCCGTTCTTCTGGATGTTTGAGAATGTGGTGGCCATGAAGGTTAATGACAAGAAAGACATCTCCAGATTCTTGGCGTGTAACCCAGTGATGATCGATGCCATCAAGGTTTCTGCCGCTCACAGGGCCCGATACTTCTGGGGCAACCTGCCCGGGATGAACAGGCC----CGTGATAGCTTCCAAGAATGATAAGCTCGAGCTGCAGGACTGCCTGGAGTTCAGTAGGACAGCAAAGTTAAAGAAAGTACAGACAATAACCACC-----------AAGTCGAACTCCATCAGACAGGGGAAAAACC---------AGCTTTTCCCTGTAGTGATGAATGGCCAGGACGATGTTTTGTGGTGTACTGAGCTCGAAAGGATCTTCGGCTTTCCTGCACACTACACAGACGTGTCCAACATGGGCCGTGGTGCCCGCCAGAAGCTGCTGGGAAGGTCCTGGAGTGTGCCAGTCATCAGACACCTCTTTGCCCCCTTGAAGGACTACTTTGCATGTGAA---

>FieldVolePredicted_3C_(incomplete)

ATGGAGGGGGAGAGCAGACAGCTCAATGCTGATGAGGCTGCCGGTGGATGTGAGGATTGTGTCATCATCAGTGGGAACTGCAGTGACCAGTCCTCAGACGCCAAGAATGTTCCCTTGACCCAAGTCATGGAGGCAATCTGCACAGTGGAGCACGA-AGGCTGCAGAGCAAACTCACGACCGTCCAAGAGGAAGGTCTC-CAGCACGATTGCTTACATCCAGGACCTGACTGAAGATGGAGATGAAGGTAGAGGTGGCGAGGCAGAGGACAGCAATGGCTCTGGTTCTCTGGTGACGCCCACACTCTCCTGG---------GAGACCAGCGCACCCTCTAATACCTCAGCATCCCTGAGCTGGCAAGCAACCACTTCAGCAAGCACACCACAGCTGTCCCCTGCCAGCCCTTACCCCACCATTGACCTCACAGATGAAGAAGCGACACCCTGGAGCATCAGTTCCCCACTGGTTGAGCTGAGCCAGGACAGCCACCAGGAGGGCATGGACACTACCCAGGTGGATGTGGAAAGCAGAGAGGGAGCCAACACTGAGTATC------------------------------------------------------------------------------------------------AGGTACC----------ACTGAGAAGGCAGTCTGACTGTCTTAGTGTTTCTGTTGCTGTGAAGAGACACC--ATGACCA--TGGCAACTCTTATAAAGGAAGCC----ATTTCATTGGACCCACTGGCATCGAGGGCCTCAAACCCAACAACAAGCAACCA---GAGAACAAAAGTCGAAGACGCACAACCACCGACTCTGCCTCTGCTGAGTACTGTCCCCCACCCAAGCGCCTCAAGACCAATAGCTATGGCGGGAAAGACAAAAGGGAGGATGATGAGAGCCGAGGTGAATT-------------------------------------------TGGGGGTGGTGCAGCAGCAGGAGCAGCAGGTCCTCAGGACCACTGCTCTGGGGT---------TCAGAGGTGCATGG-------------------CTGCATGGG-------AAAGGAGACTTCCTTACCTTCCTGTCCCTTTTT----------CCTCCACAGAAAGGATGGCTTCTGATGTCGCCAACAACAAAGGC-AATCTGGAAGGTACTATGTCTCTTTGCCCTCCTTTTTGGGTGCCCCAGGCCTGCCTTAGCCTGGC-------------------TTTGACCTTCTGCATGGATTTTAATCTGGAGCACTG-----------------GGCCCAATCAGAACTTGTGCCTCAGTATCTGAGATTTAATGTTTATGAAAGATGGAGATGCCAAAT--------------CCAAAACAATATTCAGCCTCTTGTTTTGCCTGGCCAGGCCTTGGCTTACCTTGTGCTTCCTGTTCGTTTGA-------------ACTTGTCCTTC------CCAGACAACAAAGTGGGATGGCGCTTTAAACAAGGTCGAGTCT----TCTGTGCCGAGTCCATCGCCGTGGGCACCATTAAGCATGAAGGCCAAATCAAATACGTGGATGACGTCAGGAACATCACAAAGGAAAATATTGATGAGTGGGGCCCATTCGACTTGGTGATTGGTGGAAGCCCTTGCAATGACCTCTCTTGTGTGAACCCTGTCCGAAAAGGCCTGTTTGAGGGCACTGGCCGACTCTTCTTTGAGTTTTATCGATTGCTAAACCACTCACGCCCTGAGGAGTGTGATGACCGTCCATTCTTCTGGATGTTTGAGAATGTGGTAGCCATGGATGTCGGTGACAAGCAGGATATCTCACGATTCCTGGAGTGTAACCCAGTGATGATCGATGCCATCAAGGTTTCTGCCGCTCACAGGGCCCGATACTTTTGGGGCAACCTGCCCGGGATGAACAGG------------------------------------------------------------------------------------------------------------------------------------------------------------------------------------------------------------------------------------------------------------------------------------------------------------------------------------------------------------------

>BankVolePredicted_3C_(incomplete)

ATGGAGGGGGAGAGCAGACAGCTCAGTGATGATGAGGGTGTCGGTGGATGTGAGGATTGTGTCATCATCAGTGGGAACTGCAGTGACCAGTCCTCAGACGCCAGGAACGTTCCCTTGACCCAAGTCATGGAGGCGATCTGCACAGTGGGACGCGA-AGGCTGCAGAGCAAACTCACGACCGTCCAAGAGGAGGGTCTC-CAGCACGGTTGCTTACATCCAGGACCTGACAGAAGATGAAGATGAAGGTAGAGATGGCGAGGTGGAGGACAGCAGTGGCTCTGGCTCTCCGGTGTCGCCCACACTCTGCTGG---------GAGACCAGGGCACCCTCTAATACCTCAGCATCCCTGAGTTGGCAAGCAACCACTTCAGCAAGCACACCACGGCTGTCCCCTGCCAGCCCTTACCCCATCATTGACCTCACAGATGAAGAAGTGATACCCGGGAGCATCAGTGCCCCATTGGTTGACCTGAACCAGGACAGCCACCAGGAGGGCATGGACACCACACAGGCGAATGTGGAAAGCAGAGATGGAGCCAGCACCGA---------------------------------------------------------------------------------------------------------------GGTGCGAGCTGGCAAGAC-------CTTCCCCAGCAGCCC--------GGGAGACTCACTGGAAGATCAGCTGAAGCCCATGCTGGAGTGGGCCCACGGGGGCTTCAAGCCCCCTGGCATCGAGGGCCTCCAACCGAACAACAAGCAACCAGGTGGGAATGAATGGCACAGACGTA------------------------------------------GGAACTTAGAACCCAGGAGATACGGTAT--------------------------------------------------------------------TCCCTCACCATCTTTGAGTGTTCTGTTTT----------CTGTGCTCTGTCCTGCGGTTTCTCGGTGTCCACGGTGCTCAGTGAAGAAGAATCCAATCTTCTAAGCTGAGCTGAATCTCTCACCCTCACCTATGACCCTCAGGCTAAC----CTGGGTGCACATCCAGCCTCCTAAAACCTTAG--GACTCTTCCCT--TC----------TCTTTTGTTGGTTAATATTAATGGCACATATGCAAGCATTGGC------------------AGATGTCACTTTGTCACCAAAGACTCCTGGTGTTTCTGTTTTCTGAG---ACAACAGTCTCACTCTGCAGTCCTTTCTGGTCTCATGGGGAATTGTAGC-----CGTGTGTCACCTC---ATC-TGGCTCCTCATGACTTTAGA---------AGGGTGTCTGAGT--------------------------------------------TTCTTACCTCGTGTGTAGGAATGGGGAAGAGGTACTTGGTGCTCAAAGAGTTGGGTATCAAAGTAGAAAAGTAC----------GTTGCCTCCGAAATCTGTGCAGAGTCCATCACCGTGGGCACCATTAAGCATGAAGGCCAAATCAAATACGTGGATGACGTCAGGAACATCACAAAGGGAAATATTGATGAGTGGGGCCCATTCGACTTGGTGATTGGTGGAAGCCCTTGCAATGACCTCTCTTGTGTGAACCCTGTCCGAAAAGGCCTGTTTGAGGGCACTGGCCGACTCTTCTTTGAGTTTTATCGATTGCTAAATCACTCACGCCCTGAGGAGTGTGATGACCGTCCATTCTTCTGGATGTTTGAGAATGTGGTAACCATGGAGGTCGGTGACAAGCGGGACATCTCACAATTCCTGGAGTGTAACCCAGTGATGATCGATGCCATCAAGGTTTCTGCCGCTCACAGGGCCCGGTACTTTTGGGGCAACCTGCCCGGGATGAACAGGCCTCCTTGTGCCGTCCTTTGTGA--------------TTTCAGGAA-----------CAGCTTGTCGGCTAAGCTGGGAAATGCCACTGTAGTGACCATCCATATATGCAAGAGTCACACT-----------GGGAAACACTTTCTGAAGGATCTTTTCCCTGTAGTGATGAATGGCAAGGACGATGATTTGTGGTGCACTGAGCTCGAAAGGATCTTCGGCTTTCCTGCACACTACACAGACGTGTCCAACATGGGCCGTGGTGCCCGCCAGAAGCTGCTGGGGAAGTCCTGGAGTGTGCCAGTCATCAGACACCTCTTTGCCCCCCTGAAGGACTACTTTGCATGTGAA---

>DeerMouse3C

ATGGAGGGAGAGAGTAGATACCTCCAGGATGAAGAGGGTGTCACTGGATGTGAGGACTGTGTCGTCATCAATGGGACCTGCAGTGGCCAGTCCTCAGACACTGCCAATGGTCCCTTGATCCACGTCTTGGAGGCAGTCTGCACGGTGGAGAGCAG-AGACTGTAGAACAAGCTCACGACCATCCAAGAGGAAGGTCTC-CAGCACGATTGCAGACATTCAGGACCTGACAGGAGATGGAGATGAAGGTAGAGATGACGAGGTGGAGGACAGCCGTGGCTCTGGCACCCCAGTGACACCCAAACTCTTCTGG---------AAGGCCAGGACACCCTCTAAGTCCCCAGCATCCCTGAGTTGGCAAGCAACCACTTCAGCAAGCATGTCATGGCTGTCCCCTGCCAGCCCTTACCCTACCATTGACCTCACAGATGAAGAAGTGATACCCCAGAGCATCAGTACCCCATTGGTTGACTTGAGCCAGGACAGCTATCAGGAGAGCATGGATACCACACAGGTGGATGTGAAAAGCAGAGATGGAGATAACACCAAGTATC------------------------------------------------------------------------------------------------AGAGAGCCAGGGTGCGAGCTGGCAAGAC-------CTTCCCCAGCAGCCC--------GGGAGACTCGCTGGAGGACCGGCTGAAGCCCGTGCTGGAGTGGGCCCACGGCG------------------TCGAGGGCCTCAAACCAAACAGCAAGCAACCA---GAGAACAAAGGTCGAAGACGCACCACAAATGACTCTGCCGCTTCTGAGTACTGTCCCACACCCAAGCGCCTCAAGACAAACAGCTGCGGAGGGAAAGACCGAGGGGAGGATGACGAGAGCCGAGAGCGGATGGCTTCTGACGTCACCAACAACAAGGGCAATCTGGAAGACCGCTGTTTGTCCTGCGGTAGGAAGAACCCTGTGTCCTTCCATCCCCTCTTTGAGGGTGGGCTCTG-TCAGAGTTGCCGGGATCGCTTCCTGGAGCTCTTCTACATGTATGA----TGAGGACGGCTATCAGTCCTACTGCACCGTGTG----------CTGCGAGGGCCGGGA--GCTGCTTCTCT--GCAGCAACACCAGCTGCTGCAGGTGCTTCTGTGTGGAGTGTCTGGAGGTGCTGGTGGGGACAGGGACAGCGGAGGATGCCAAGCTGA-----AGGAACCCTGGAGCTGCTATATGTGCAT---CCCACAGCGCTGCCATGGTGTCCTCA---GGCGCAGGAAGGATTGGAACATGCGCCTGCAAGACTTC---TTCACGACTGATCCTGACCTCGAAGAATTT---GAGCCGCCCAAGTTGTACCCAGCAATTCCCGCAGCCAGGAGGAGGCCCA--TTAGAGTCCTGTCCCTGTTTGATGGAATTGCGACAGGGTACTTGGTGCTCAAAGAGTTGGGTATCAAAGTGGAAAAGTAC----------GTCGCCTCCGAAGTCTGTTCAGAGTCCATCGCCGTGGGAACCGTTAAGCATGAAGGCCATATCAAATACGTGGATGACGTCAGGAAAATCACAAAGAAAAATATTGAGGAGTGGGGCCCATTTGACTTGGTGATTGGTGGAAGCCCTTGCAATGACCTCTCCAATGTCAATCCCGCCAGGAAAGGCCTATATGAGGGCACCGGCAGGCTCTTCTTTGAGTTTTACCACTTGCTGAATTATACACGCCCCAAGGAGGGCGACAACCGTCCATTCTTCTGGATGTTTGAGAATGTGGTAGCCATGAAGGTCAACGACAAGAAAGACATCTCCCGATTCTTGGCGTGTAACCCAGTGATGATCGACGCCATCAAGGTTTCTGCCGCTCACAGGGCCCGATACTTCTGGGGCAACCTCCCTGGGATGAACAGGCC----CGTGATAGCTTCAAAGAATGATAAGCTCGAGCTGCAGGACTGCTTGGAGTTCAGTAGGACAGCAAAGTTAAAGAAAGTACAGACAATAACCACC-----------AAGTCGAACTCCATCAGACAGGGGAAAAACC---------AGCTTTTCCCTGTAGTCATGAATGGCAAGGACGATGTTTTGTGGTGTACTGAGCTCGAAAGGATCTTCGGCTTTCCTGCCCACTACACTGACGTCTCCAACATGGGCCGTGGCGCCCGCCAGAAGCTGCTGGGAAGGTCCTGGAGTGTGCCAGTCATCAGACATCTCTTTGCCCCCTTGAAGGACTACTTTGCATGTGAA---

>Pahari_3C

------GGAAGTAGCAGACACCTCAGCGATGAGGAGGATGTCAGTGGATGTGAGGACTGTATCATCATCAGTGGGACCCGCAGTGACCAGTCTTCAAACCCCAAGACTGTTCCTTTGACCCAACTCTTGGAGGCAGTCTGCACAGTGGAGAGCAG-AGGATGCAGGACAAGCTCACGGCCATCCAAGAGGAAAGTATC-CAGCCTGATTAGTTCCGTTCAGGACCTCACAGGAGATGGAGATGGAGACAGGGATGGTGAGATGGGGAACAGCAGTGGCTCTGACACTCCAGTGATG---------TTCTGC---------GAGACCAGGACACCCTCTAAAACCCCAGCAAAAACCACTTCAGAAGCAACCACTTCAGCAAGCACACCCAGGCTGTCCCCTGTCAGCCCTTACCCCATCATTGACCTCACAGATGAAGATACGATACCCCAGAGCATCAGTACCCCATCGGTTGACTGGAGCCAGGACAGCCATCAGGAGGGCATGGATACCACACAGGTGGATGCAGAAAGCAGAGATGGACATAACACTGAGTATC------------------------------------------------------------------------------------------------AGAAAGCCAGGGTTCGAGCGGGCAAGAC-------CTTCCCCAGCAGTCC--------CGGAGAGTCACTGGAGCACCAGCTGAAGCCCGTGCTGGAGTGGGCCCACAGCAGCTTCAAGCCCACTGGGATTGAGGGCCTCCAACCCAACAGCAAGCAGCCA---GAGGACAAAAGTCAAAGACGCACAACCAATGACTCTGCCGTTTCTGAGTACTCTACCCCACCCAAGCGCCTCAAGACGAATAGCTATGGTGGGAAGGACCTAAAGGGGGATGAGGAGAGCCGA-----GGTGATTTTGG----------------GGAGATGCTGGAAGAACACTGCTTGTCCTGCGGTAGGAGGGACCCTGTGTCCTTCCACCCCCTCTTTGAGGGTGGGCTCTG-TCAGAGTTGCCGGGACCGCTTCCTGGAGCTCTTCTACATGTATGA----CGAGGACGGCTATCAGTCCTACTGCACCGTGTG----------CTGTGAGGGCTATGA--ATTGCTGCTGT--GCAGTAACACAAGCTGCTGCAGGTGCTTCTGTGTGGAGTGTCTGGAGGTGCTGGTGGGTGCAGGCACAGCTGAGGATGCCAAGCTGC-----AGGAACCCTGGAGTTGCTATATGTGCCT---CCCCCAGCGCTGCCATGGGTTCCTCC---GACGCAGGAAGGATTGGAACATACGCCTGCAGGACTTC---TTCACTACTGATCCTGACCTGGAAGAATTTCAGGAGCCACCCAAGTTGTACCCAGCAATTCCTGCAGCCAAAAGAAGGCCCA--TTAGAGTCCTGTCTCTGTTTGATGGAATTGCGACAGGGTACTTGGTGCTCAAGGAGTTGGGTATTAAAGTGGAGAAGTAC----------GTTGCCTCCGAAGTCTGTGCAGAGTCCATCGCTGTGGGAACCGTTAAGCATGAAGGCCAAATCAAATATGTGGATGACATCAGGAACATTACAAAGGAACATATCGACGAGTGGGGCCCATTCGACCTGGTGATTGGTGGAAGCCCCTGCAATGATCTTTCCTGTGTGAATCCTGTCAGGAAAGGCCTGTTTGAGGGTACTGGTCGGCTCTTCTTTGAGTTTTATCGATTGCTAAATTACTCACGCCCTGAGGAGGAGGATGACCGCCCATTCTTTTGGATGTTTGAGAATGTGGTAGCCATGAAGGTTGGTGACAAGAGGGACATCTCACGATTCCTGGAGTGTAACCCAGTGATGATCAATGCCATCAAGGTTTCTGCTGCTCACAGGGCCCGGTACTTCTGGGGCAACCTACCTGGAATGAACAGGCC----CGTGATAGCTTCAAAGAATGATAAGCTCGAGCTGCAGGACTGCCTGGAGTTCAGTAGGACAGCAAAGTTAAAGAAAGTGCAGACAATAACCACC-----------AAGTCGAACTCCATCAGACAGGGGAAAAAAC---------AGCTTTTCCCTGTAGTCATGAATGGCAAAGACGACATTCTGTGGTGCACTGAGCTCGAAAGGATCTTCGGCTTCCCTGCTCACTACACGGATGTGTCCAACATGGGCCGTGGTGCCCGCCAGAAGCTGCTGGGCAGGTCCTGGAGTGTGCCAGTCATCAGACACCTGTTGGCCCCCTTAAAGGACCACTTTGCCTGTGAA---

>Caroli_3C

------GGAGGTAGCAGACACCTCAATAATGAGGAGGATGTCAGTGGATGTGAGGACTGTATTATCATCAGTGGGACCTGCAGTGACCAGTCTTCAGACCCCAAGACTGTTCCTTTGATCCAAGTCTTGGAGGCATTCTGCACAGTGGAGAGCAG-AGGATGCAGAACAAGCTCACAACCACCGAAGAGGA----------------CTAGTTACGTTCAGGACCTCACAGGAGATGGAGATGAAGATAGGGATGGTGAGGTGGGGGGCAGCAGTGGCTCTGGCACTCCAGTGATGCCCCAATTCCTCTGC---------GAGACCAGGATACCCTCTAAAGCCCCAGCATCCCTCAGTTGGCAAGCAAACACTTCAGCAAGCACACCCTGGCTGTCCCCTGCCAGCCCTTACCCCATCATTGACCTCACAGATGAAGATGTGATACCCCAGAGCGTCAGTACCCCATCGGTTGACTGGAGCCAGGACAGCCATCAGGAGGGCATGGATACCACACAGGCAGATGCAGAGAGCAGAGATGGAGGCAACATTGAGTATC------------------------------------------------------------------------------------------------AGAAAGCCAGGGTGCGAGCTGGCAAGGC-------CTGCCCCAGCAGTCC--------TGGAGAGTCACTGGAGGACCAGCTGAAGCCCATGCTGGAGTGGGCCCACGGTGGTTTCAAGCCCACTGGGATCGAGGGCCTCAAACCCAACAACAAGCAACCA---GAGAACAAAAGTCGAAGGCGCACAGCCAATGACTCTGCTGCTTCTGAGTATTCCCCCCCACCCAAGCGCCTCAAGACCAACAGCTATGGTGGGAAGGACCGAGGGGAGGATGAGGAGAGCCGAGAACGGATGGCTTCTGATGTCACCAACAACAAGGGCAATCTGGAAGACCACTGTTTGTCCTGTGGTAGGAAAGACCCTGTGTCCTTCCACCCCCTCTTTGAGGGTGGGCTCTG-TCAGAGTTGCCGGGATCGCTTCCTAGAACTCTTCTACATGTATGA----CGAGGACGGCTATCAGTCCTACTGCACCGTGTG----------CTGTGAGGGCCGTGA--ACTGCTGCTGT--GCAGTAACACAAGCTGCTGCAGATGCTTCTGTGTGGAGTGTCTGGAGGTGCTGGTGGGCGCAGGCACAGCTGAGGATGCCAAGCTGC-----AGGAACCCTGGAGCTGCTATATGTGCCT---CCCTCAGCGCTGCCATGGGGTCCTCC---GACGCAGGAAGGATTGGAACATGCGCCTGCAAGACTTC---TTCACTACTGATCCTGACCTGGAAGAATTTCAGGAGCCGCCCAAGTTGTACCCAGCAATTCCTGCTGCCAAAAGGAGGCCCA--TTAGAGTCCTGTCTCTGTTTGATGGAATTGCAACAGGGTACTTGGTGCTCAAGGAGTTGGGTATTAAAGTGGAGAAGTAC----------ATTGCCTCCGAAGTCTGTGCAGAGTCCATCGCTGTGGGAACCGTTAAGCATGAAGGCCAAATCAAATACGTGGATGACATCAGGAACATTACAAAGGAACATATTGATGAGTGGGGCCCGTTCGACCTGGTGATTGGTGGAAGCCCCTGCAATGATCTTTCCTGTGTGAATCCTGTCAGGAAAGGCCTGTTTGAGGGTACTGGCCGGCTCTTCTTTGAGTTTTACCGATTGCTAAATTACTCATGCCCTAAGGAGGAGGATGACCGCCCATTCTTCTGGATGTTTGAGAATGTGGTAGCCATGGAGGTCGGTGACAAGAGGGACATCTCACGATTCCTAGAGTGTAACCCAGTGATGATCGATGCCATCAAGGTGTCTGCTGCTCACAGGGCCCGGTACTTCTGGGGTAACCTACCCGGAATGAACAGGCC----CGTGATGGCTTCAAAGAATGATAAGCTCGAGCTGCAGGACTGCCTGGAGTTCAGTAGGACAGCAAAGTTAAAGAAAGTGCAGACAATAACCACC-----------AAGTCGAACTCCATCAGACAGGGGAAAAACC---------AGCTTCTCCCTGTAGTCATGAATGGCAAGGACGACGTTCTGTGGTGCACTGAGCTCGAAAGGATCTTCGGCTTTCCTGAACACTACACAGACGTGTCCAACATGGGCCGTGGCGCCCGTCAGAAGCTGCTGGGCAGGTCCTGGAGTGTGCCAGTTATCAGACACCTGTTTGCCCCCTTGAAGGACCACTTTGCCTGTGAA---

>Mus_spretus_Dnmt3C_CDS_(incomplete)

ATGAGGGGAGGTGGCAGACACCTCAGTAATGAGGAGGATGTCAGTGGATGTGAGGACTGTATTATCATCAGTGGGACCTGCAGTGACCAGTCTTCAGACCCCAAGACTGTTCCTTTGACCCAAGTCTTGGAGGCAGTCTGCACAGTGGAGAGCAG-AGGATGCAGAACAAGCTCACAACCATCCAAGAGGAAAGCATC-CAGCCTGATTAGTTATGTTCAGGACCTCACAGGAGATGGAGATGAAGATAGGGATGGTGAGGTGGGGGGCAGCAGTGGCTCTGGCACTCCAGTGATGCCCCAACTCTTCTGT---------GAGACCAGGATACCCTCTAAAACCCCAGCACCCCTCAGTTGGCAAGCAAACACTTCAGCAAGCACGCCCTGGTTGTCCCCTGCCAGCCCTTACCCCATCATTGACCTCACAGATGAAGATGTGATACCCCAGAGCATCAGTACCCCATCGGTTGACTGGAGCCAGGACAGCCATCAGGAGGGCATGGATACCACACAGGTGGATGCAGAGAGCAGAGATGGAGGCAACATTGAGTATCAGGTCTCGGCTGACAAACTGTTGCTCAGCCAGTCCTGTATCCTGGCCGCCTTCTATAAACTGGTTCCTTACAGGAAGTCTATATACCGTACTCTGGAGAAAGCCAGGGTGAGAGCTGGCAAGGC-------CTGCCCCAGCAGTCC--------TGGAGAGTCACTGGAGGACCAGCTGAAGCCCATGCTGGAGTGGGCCCACGGTGGCTTCAAGCCTACTGGGATCGAGGGCCTCAAACCCAACAAGAAGCAACCA---GAGAACAAAAGTCGAAGACGCACAAC----------------------------------------------------------------------------------------------------------------------------------------------------------------------------------------------------------------------------------------------------------------------------------------------------------------------------------------------------------------------------------------------------------------------------------------------------------------------------------------------------------------------------------------------------------------------------------------------------------------------------------------------------------------CAACAGGGTACTTGGTGCTCAAGGAGTTGGGTATTAAAGTGGAAAAGTAC----------ATTGCCTCCGAAGTCTGTGCAGAGTCCATCGCTGTGGGAACCATTAAGCATGAAGGCCAGATCAAATACGTGGATGACATCAGGAACATTACAAAGGAACATATTGACGAGTGGGGCCCGTTCGACCTGGTGATTGGTGGAAGCCCCTGCAATGATCTTTCCTGTGTGAATCCTGTCAGGAAAGGCCTGTTTGAGGGTACTGGCCGGCTCTTCTTTGAGTTTTACCGATTGCTAAATTACTCATGCCCTGAGGAAGAGGATGACCGCCCATTCTTTTGGATGTTTGAGAATGTGGTAGCCATGGAGGTCGGTGACAAGAGGGACATCTCACGATTCCTGGAGTGTAACCCAGTGATGATCGATGCCATCAAGGTGTCTGCTGCTCACAGGGCCCGGTACTTCTGGGGTAACCTACCCGGAATGAACAGGCC----CGTGATGGCTTCAAAGAATGATAAGCTCGAGCTGCAGGACTGCCTGGAGTTCAGTAGGACAGCAAAGTTAAAGAAAGTGCAGACAATAACCACC-----------AAGTCGAACTCCATCAGACAGGGCAAAAACC---------AGCTTTTCCCTGTAGTCATGAATGGCAAGGACGACGTTTTGTGGTGCACTGAGCTCGAAAGGATCTTCGGCTTTCCTGAACACTACACAGACGTGTCCAACATGGGCCGTGGCGCCCGTCAGAAGCTGCTGGGCAGGTCTTGGAGTGTGCCAGTCATCAGACACCTGTTTGCCCCCTTGAAGGACCACTTTGCCTGTGAA---

>Mouse_Dnmt3c_long_isoform_CDS

ATGAGGGGAGGTAGCAGACACCTCAGTAATGAGGAGGATGTCAGTGGATGTGAGGACTGTATTATCATCAGTGGGACCTGCAGTGACCAGTCTTCAGACCCCAAGACTGTTCCTTTGACCCAAGTCTTGGAGGCAGTCTGCACAGTGGAGAACAG-AGGATGCAGAACAAGCTCACAACCATCCAAGAGGAAAGCATC-CAGCCTGATTAGTTACGTTCAGGACCTCACAGGAGATGGAGATGAAGATAGGGATGGTGAGGTGGGGGGCAGCAGTGGCTCTGGCACTCCAGTGATGCCCCAACTCTTCTGT---------GAGACCAGGATACCCTCTAAAACCCCAGCACCCCTCAGTTGGCAAGCAAACACTTCAGCAAGCACGCCCTGGCTGTCCCCTGCCAGCCCTTACCCCATCATTGACCTCACAGATGAAGATGTGATACCCCAGAGCATCAGTACCCCATCGGTTGACTGGAGCCAGGACAGCCATCAGGAGGGCATGGATACCACACAGGTGGATGCAGAGAGCAGAGATGGAGGCAACATTGAGTATC------------------------------------------------------------------------------------------------AGAAAGCCAGGGTGAGAGCTGGCAAGGC-------CTGCCCCAGCAGTCC--------TGGAGAGTCACTGGAGGACCAGCTGAAGCCCATGCTGGAGTGGGCCCACGGTGGCTTCAAGCCCACTGGGATCGAGGGCCTCAAACCCAACAAGAAGCAACCA---GAGAACAAAAGTCGAAGACGCACAACCAATGACCCTGCTGCTTCTGAGTCCT---CCCCACCCAAGCGCCTCAAGACAAATAGCTATGGCGGGAAGGACCGAGGGGAGGATGAGGAGAGCCGAGAACAGATGGCTTCTGATGTCACCAACAACAAGGGCAATCTGGAAGACCACTGTTTGTCGTGCGGTAGGAAGGACCCTGTGTCCTTCCACCCCCTCTTTGAGGGTGGGCTCTG-TCAGAGTTGCCGGGACCGCTTCCTAGAGCTCTTCTACATGTATGA----CGAGGACGGCTATCAGTCCTACTGCACCGTGTG----------CTGTGAGGGCCGTGA--ACTGCTGCTGT--GCAGTAACACAAGCTGCTGCAGATGCTTCTGTGTGGAGTGTCTGGAGGTGCTGGTGGGTGCAGGCACAGCTGAGGATGTCAAGCTGC-----AGGAACCCTGGAGCTGCTATATGTGCCT---CCCTCAGCGCTGCCATGGGGTCCTCC---GACGCAGGAAAGATTGGAACATGCGCCTGCAAGACTTC---TTCACTACTGATCCTGACCTGGAAGAATTTCAGGAGCCGCCCAAGTTGTACCCAGCGATTCCTGCAGCCAAAAGGAGGCCCA--TTAGAGTCCTGTCTCTGTTTGATGGAATTGCAACAGGGTACTTGGTGCTCAAGGAGTTGGGTATTAAAGTGGAAAAGTAC----------ATTGCCTCCGAAGTCTGTGCAGAGTCCATCGCTGTGGGAACCGTTAAGCATGAAGGCCAAATCAAATATGTGGATGACATCAGGAACATTACAAAGGAACATATTGACGAGTGGGGCCCGTTCGACCTGGTGATTGGTGGAAGCCCCTGCAATGATCTTTCCTGTGTGAATCCTGTCAGGAAAGGCCTGTTTGAGGGTACTGGCCGGCTCTTCTTTGAGTTTTACCGATTGCTAAATTACTCATGCCCTGAGGAGGAGGATGACCGCCCCTTCTTCTGGATGTTTGAGAATGTGGTAGCTATGGAGGTCGGTGACAAGAGGGACATCTCACGATTCCTGGAGTGTAACCCAGTGATGATCGATGCCATCAAGGTGTCTGCTGCTCACAGGGCCCGGTACTTCTGGGGTAACCTACCCGGAATGAACAGGCC----CGTGATGGCTTCAAAGAATGATAAGCTCGAGCTGCAGGACTGCCTGGAGTTCAGTAGGACAGCAAAGTTAAAGAAAGTGCAGACAATAACCACC-----------AAGTCGAACTCCATCAGACAGGGCAAAAACC---------AGCTTTTCCCTGTAGTCATGAATGGCAAGGACGACGTTTTGTGGTGCACTGAGCTCGAAAGGATCTTCGGCTTTCCTGAACACTACACAGACGTGTCCAACATGGGCCGTGGCGCCCGTCAGAAGCTGCTGGGCAGGTCTTGGAGTGTGCCAGTCATCAGACACCTGTTTGCCCCCTTGAAGGACCACTTTGCCTGTGAATAG
