## Supplementary material for "Dynamic evolution of *de novo* DNA methyltransferases in rodent and primate genomes": Suppl data 3

*SupFile_3: Muroid* DNMT3A *sequence alignments*

>Mouse3A

ATGCCCTCCAGCGGCCCCGGGGACACCAGCAGCTCCTCTCTGGAGCGGGAGGATGATCGAAAGGAAGGAGAGGAACAGGAGGAGAACCGTGGCAAGGAAGAGCGCCAGGAGCCCAGCGCCACGGCCCGGAAGGTGGGGAGGCCTGGCCGGAAGCGCAAGCACCCACCGGTGGAAAGCAGTGACACCCCCAAGGACCCAGCAGTGACCACCAAGTCTCAGCCCATGGCCCAGGACTCTGGCCCCTCAGATCTGCTACCCAATGGAGACTTGGAGAAGCGGAGTGAACCCCAACCTGAGGAGGGGAGCCCAGCTGCAGGGCAGAAGGGTGGGGCCCCAGCTGAAGGAGAGGG---AACTGAGACCCCACCAGAAGCCTCCAGAGCTGTGGAGAATGGCTGCTGTGTGACCAAGGAAGGCCGTGGAGCCTCTGCAGGAGAGGGCAAAGAACAGAAGCAGACCAACATCGAATCCATGAAAATGGAGGGCTCCCGGGGCCGACTGCGAGGTGGCTTGGGCTGGGAGTCCAGCCTCCGTCAGCGACCCATGCCAAGACTCACCTTCCAGGCAGGGGACCCCTACTACATCAGCAAACGGAAACGGGATGAGTGGCTGGCACGTTGGAAAAGGGAGGCTGAGAAGAAAGCCAAGGTAATTGCAGTAATGAATGCTGTGGAAGAGAACCAGGCCTCTGGAGAGTCTCAGAAGGTGGAGGAGGCCAGCCCTCCTGCTGTGCAGCAGCCCACGGACCCTGCTTCTCCGACTGTGGCCACCACCCCTGAGCCAGTAGGAGGGGATGCTGGGGACAAGAATGCTACCAAAGCAGCCGACGATGAGCCTGAGTATGAGGATGGCCGGGGCTTTGGCATTGGAGAGCTGGTGTGGGGGAAACTTCGGGGCTTCTCCTGGTGGCCAGGCCGAATTGTGTCTTGGTGGATGACAGGCCGGAGCCGAGCAGCTGAAGGCACTCGCTGGGTCATGTGGTTCGGAGATGGCAAGTTCTCAGTGGTGTGTGTGGAGAAGCTCATGCCGCTGAGCTCCTTCTGCAGTGCATTCCACCAGGCCACCTACAACAAGCAGCCCATGTACCGCAAAGCCATCTACGAAGTCCTCCAGGTGGCCAGCAGCCGTGCCGGGAAGCTGTTTCCAGCTTGCCATGACAGTGATGAAAGTGACAGTGGCAAGGCTGTGGAAGTGCAGAACAAGCAGATGATTGAATGGGCCCTCGGTGGCTTCCAGCCCTCGGGTCCTAAGGGCCTGGAGCCACCAGAAGAAGAGAAGAATCCTTACAAGGAAGTTTACACCGACATGTGGGTGGAGCCTGAAGCAGCTGCTTACGCCCCACCCCCACCAGCCAAGAAACCCAGAAAGAGCACAACAGAGAAACCTAAGGTCAAGGAGATCATTGATGAGCGCACAAGGGAGCGGCTGGTGTATGAGGTGCGCCAGAAGTGCAGAAACATCGAGGACATTTGTATCTCATGTGGGAGCCTCAATGTCACCCTGGAGCACCCACTCTTCATTGGAGGCATGTGCCAGAACTGTAAGAACTGCTTCTTGGAGTGTGCTTACCAGTATGACGACGATGGGTACCAGTCCTATTGCACCATCTGCTGTGGGGGGCGTGAAGTGCTCATGTGTGGGAACAACAACTGCTGCAGGTGCTTTTGTGTCGAGTGTGTGGATCTCTTGGTGGGGCCAGGAGCTGCTCAGGCAGCCATTAAGGAAGACCCCTGGAACTGCTACATGTGCGGGCATAAGGGCACCTATGGGCTGCTGCGAAGACGGGAAGACTGGCCTTCTCGACTCCAGATGTTCTTTGCCAATAACCATGACCAGGAATTTGACCCCCCAAAGGTTTACCCACCTGTGCCAGCTGAGAAGAGGAAGCCCATCCGCGTGCTGTCTCTCTTTGATGGGATTGCTACAGGGCTCCTGGTGCTGAAGGACCTGGGCATCCAAGTGGACCGCTACATTGCCTCCGAGGTGTGTGAGGACTCCATCACGGTGGGCATGGTGCGGCACCAGGGAAAGATCATGTACGTCGGGGACGTCCGCAGCGTCACACAGAAGCATATCCAGGAGTGGGGCCCATTCGACCTGGTGATTGGAGGCAGTCCCTGCAATGACCTCTCCATTGTCAACCCTGCCCGCAAGGGACTTTATGAGGGTACTGGCCGCCTCTTCTTTGAGTTCTACCGCCTCCTGCATGATGCGCGGCCCAAGGAGGGAGATGATCGCCCCTTCTTCTGGCTCTTTGAGAATGTGGTGGCCATGGGCGTTAGTGACAAGAGGGACATCTCGCGATTTCTTGAGTCTAACCCCGTGATGATTGACGCCAAAGAAGTGTCTGCTGCACACAGGGCCCGTTACTTCTGGGGTAACCTTCCTGGCATGAACAGGCCTTTGGCATCCACTGTGAATGATAAGCTGGAGCTGCAAGAGTGTCTGGAGCACGGCAGAATAGCCAAGTTCAGCAAAGTGAGGACCATTACCACCAGGTCAAACTCTATAAAGCAGGGCAAAGACCAGCATTTCCCCGTCTTCATGAACGAGAAGGAGGACATCCTGTGGTGCACTGAAATGGAAAGGGTGTTTGGCTTCCCCGTCCACTACACAGACGTCTCCAACATGAGCCGCTTGGCGAGGCAGAGACTGCTGGGCCGATCGTGGAGCGTGCCGGTCATCCGCCACCTCTTCGCTCCGCTGAAGGAATATTTTGCTTGTGTGTAA

>Caroli3A

ATGCCCTCCAGCGGCCCCGGGGACAGCAGCAGCTCCTCTCTGGAGCGGGAGGATGCTCGAAAGGAAGGAGAGGAACAGGAGGAGAACCGTGGCAAGGAAGAGCGCCAGGAGCCCAGTGCCACGGCCCGGAAGGTGGGGAGGCCTGGCCGGAAGCGCAAGCACCCACCGGTGGAAAGCAGTGACACCCCCAAGGACTCAGCGGTGACCACCAAGTCTCAGCCCATGGCCCAGGACTCTGGCCCCTCAGATCTGCTACCCAATGGAGACTTGGAGAAGCGGAGTGAACCCCAACCTGAGGAGGGAAGCCCAGCTGCAGGGCAGAAGGGTGGGGCCCCAGCTGAAGGAGAGGG---AACTGAGACCCCACCAGAAGCCTCCAGAGCTGTGGAGAATGGCTGCTGTGTGACCAAGGAAGGCCGTGGAGCCTCTGCAGGAGAGGGCAAAGAACAGAAGCAGACCAACATCGAATCCATGAAAATGGAGGGCTCCCGGGGCCGACTGAGAGGTGGCTTGGGCTGGGAGTCCAGCCTCCGTCAGCGGCCCATGCCAAGACTCACCTTCCAGGCAGGGGACCCCTACTACATCAGCAAACGGAAACGGGATGAGTGGCTGGCACGTTGGAAAAGGGAGGCTGAGAAGAAAGCCAAGGTAATTGCAGTAATGAATGCTGTGGAAGAGAACCAGGCATCTGGAGAGTCTCAGAAGGTGGAGGAGGCCAGCCCCCCTGCTGTGCAGCAGCCCACGGACCCTGCTTCTCCGACTGTGGCCACCACCCCTGAGCCAGTAGGAGGGGATGCTGGGGACAAGAATGCTACCAAAGCAGCCGACGATGAGCCCGAGTATGAGGATGGCCGGGGCTTTGGCATTGGAGAGCTGGTGTGGGGGAAACTTCGGGGCTTCTCCTGGTGGCCAGGCCGAATTGTGTCTTGGTGGATGACAGGCCGGAGCCGAGCAGCTGAAGGCACTCGCTGGGTCATGTGGTTTGGAGATGGCAAGTTCTCAGTGGTGTGTGTGGAGAAGCTCATGCCGCTGAGCTCCTTCTGCAGTGCATTCCACCAGGCCACCTACAACAAGCAGCCCATGTACCGCAAAGCCATCTATGAAGTCCTCCAGGTGGCCAGCAGCCGTGCCGGGAAGCTGTTTCCAGCTTGCCATGACAGTGATGAAAGTGACAGTGGCAAGGCTGTGGAAGTGCAGAACAAGCAGATGATTGAATGGGCCCTCGGTGGCTTCCAGCCCTCGGGTCCTAAGAGCCTGGAGCCACCAGAAGAAGAGAAGAATCCTTACAAGGAAGTTTACACTGACATGTGGGTGGAGCCTGAGGCAGCTGCTTACGCCCCACCCCCACCGGCCAAGAAACCCAGAAAGAGCACAACAGAGAAACCTAAGGTCAAGGAGATCATTGATGAGCGCACAAGGGAGCGGCTCGTGTATGAGGTTCGCCAGAAGTGCAGAAACATCGAGGACATTTGTATCTCATGTGGGAGCCTCAATGTCACCCTGGAGCACCCACTCTTCATTGGTGGCATGTGCCAGAACTGTAAGAACTGCTTCTTGGAGTGTGCTTACCAGTATGACGACGATGGGTACCAGTCCTATTGCACCATCTGCTGTGGGGGGCGTGAAGTGCTCATGTGTGGGAACAACAACTGCTGCAGGTGCTTTTGTGTCGAGTGTGTGGATCTCTTGGTGGGGCCAGGAGCTGCCCAGGCAGCCATTAAGGAAGACCCCTGGAACTGCTACATGTGTGGGCATAAGGGCACCTATGGGCTGCTGCGAAGACGGGAAGACTGGCCTTCTCGACTCCAGATGTTCTTTGCCAATAACCATGACCAGGAATTTGACCCCCCAAAGGTTTACCCACCTGTGCCAGCTGAGAAGAGGAAGCCCATCCGTGTGCTGTCTCTCTTTGATGGGATTGCTACAGGGCTCCTGGTGCTGAAGGACCTGGGCATCCAAGTGGACCGCTACATTGCCTCTGAGGTGTGTGAGGACTCCATCACGGTGGGCATGGTGCGGCACCAGGGAAAGATCATGTACGTCGGGGACGTCCGCAGCGTCACACAGAAGCATATCCAGGAGTGGGGCCCATTCGACCTGGTGATTGGAGGCAGTCCCTGCAATGACCTCTCCATCGTCAACCCTGCCCGCAAGGGACTTTACGAGGGTACTGGCCGCCTCTTCTTTGAGTTCTACCGCCTCCTGCATGATGCGCGGCCCAAGGAGGGAGATGATCGCCCCTTCTTCTGGCTCTTTGAGAATGTGGTGGCCATGGGCGTTAGTGACAAGAGGGACATCTCGCGATTTCTTGAGTCTAACCCCGTGATGATTGACGCCAAAGAAGTGTCTGCTGCACACAGGGCCCGTTACTTCTGGGGTAACCTTCCTGGCATGAACAGGCCTTTGGCATCCACTGTGAATGATAAGCTGGAGCTGCAAGAGTGTCTGGAGCACGGCAGAATAGCCAAGTTCAGCAAAGTGAGGACCATTACCACCAGGTCAAACTCTATAAAGCAGGGCAAAGACCAGCATTTCCCCGTCTTCATGAACGAGAAGGAGGACATCCTGTGGTGCACTGAAATGGAAAGGGTGTTTGGCTTCCCCGTCCACTACACAGACGTCTCCAACATGAGCCGCTTGGCGAGGCAGAGACTGCTGGGCCGATCGTGGAGCGTGCCGGTCATCCGCCACCTCTTCGCTCCGCTGAAGGAATATTTTGCTTGTGTGTAA

>Phaari3A

ATGCCCTCCAGCGGCCCCGGAGACACCAGCAGCTCCTCTCTGGAGCGGGAGGATGATCGAAAGGAAGGAGAGGAACAGGAGGAGAACCGTGGCAAAGAAGAGCGCCAGGAGCCCAGCGCCACGGCCCGGAAGGTGGGGAGGCCTGGCCGGAAGCGCAAGCACCCACCGGTGGAAAGCAGTGACACCCCCAAGGACCCAGCGGTGACCACCAAGTCTCAGCCCATGGCCCAGGACTCTGGCCCCTCAGATCTGCTACCCAATGGAGACTTGGAGAAGCGGAATGAACCCCAACCTGAGGAGGGTAGCCCAGCTGCAGGGCAGAAGGGTGGGGCCCCAGCTGAGGGAGAGGG---AACTGAGACCCCACCAGAAGCCTCCAGAGCTGTGGAGAATGGCTGCTGTGTGACCAAGGAGGGCCGTGGAGCCTCTGCAGGAGAGGGCAAAGAACAGAAGCAGACCAACATCGAATCCATGAAAATGGAGGGCTCCCGGGGCCGACTGCGCGGTGGCTTGGGCTGGGAGTCCAGCCTCCGTCAGAGGCCCATGCCAAGACTCACCTTCCAGGCGGGGGACCCCTATTACATCAGCAAACGGAAACGGGATGAGTGGCTGGCACGTTGGAAAAGGGAGGCTGAGAAGAAAGCCAAGGTGATTGCAGTAATGAATGCCGTGGAAGAGAACCAGGCCTCTGGAGAGTCTCAGAAGGTGGAGGAGGCCAGCCCTCCTGCTGTGCAGCAGCCCACGGACCCTGCTTCTCCTACTGTGGCCACCACCCCTGAGCCAGTAGGGGGGGATGCTGGGGACAAGAATGCTACCAAAGCAGCTGATGATGAGCCCGAGTATGAGGATGGCCGGGGCTTTGGCATTGGAGAGCTGGTGTGGGGGAAACTTCGGGGCTTCTCCTGGTGGCCAGGCCGAATTGTGTCTTGGTGGATGACAGGCCGGAGCCGAGCAGCTGAAGGCACTCGCTGGGTCATGTGGTTCGGAGATGGCAAGTTCTCAGTGGTGTGTGTGGAGAAGCTCATGCCGCTGAGCTCCTTCTGCAGTGCGTTCCACCAGGCCACCTACAACAAGCAGCCCATGTACCGCAAAGCCATCTACGAAGTCCTCCAGGTGGCCAGCAGCCGCGCCGGGAAGCTGTTTCCAGCTTGCCATGACAGTGATGAAAGTGACAGTGGCAAGGCTGTGGAGGTGCAGAACAAGCAGATGATCGAATGGGCCCTCAGCGGCTTCCAGCCGTCAGGTCCTAAGGGCCTGGAGCCACCAGAAGAGGAGAAGAATCCTTACAAGGAAGTTTACACCGACATGTGGGTTGAGCCTGAGGCAGCTGCTTATGCCCCACCCCCACCAGCCAAGAAACCCAGAAAGAGCACAACAGAGAAGCCAAAGGTCAAGGAGATCATTGATGAACGCACAAGGGAGCGGCTCGTGTATGAGGTGCGTCAGAAGTGCAGAAACATTGAGGACATTTGTATCTCATGTGGGAGCCTCAATGTCACCCTGGAGCACCCACTCTTCATTGGTGGCATGTGCCAGAACTGTAAGAACTGCTTCTTGGAGTGTGCTTACCAGTATGACGACGATGGGTACCAGTCCTATTGTACCATCTGCTGTGGGGGGCGTGAAGTGCTCATGTGTGGGAACAACAACTGCTGCAGGTGCTTTTGTGTCGAGTGCGTGGATCTCTTGGTGGGGCCAGGCGCTGCCCAGGCAGCTATTAAGGAAGACCCCTGGAACTGCTACATGTGTGGGCATAAGGGCACCTATGGGCTGCTGCGAAGACGGGAAGACTGGCCTTCTCGACTTCAGATGTTCTTTGCCAATAACCATGACCAGGAATTTGACCCCCCCAAGGTTTACCCACCTGTGCCAGCTGAGAAAAGGAAACCCATCCGCGTGCTGTCTCTCTTTGATGGGATTGCTACAGGGCTCCTGGTGCTGAAGGACTTGGGCATCCAAGTGGACCGCTACATTGCCTCCGAGGTGTGTGAGGACTCCATCACAGTGGGCATGGTGCGGCACCAGGGAAAGATCATGTACGTCGGGGACGTCCGCAGCGTCACACAGAAGCATATCCAGGAGTGGGGTCCATTTGACCTGGTGATTGGAGGCAGTCCCTGCAATGACCTCTCCATTGTCAACCCTGCCCGCAAGGGACTTTACGAGGGCACTGGCCGCCTCTTTTTTGAGTTCTACCGCCTCCTGCATGATGCGCGGCCCAAGGAGGGAGATGATCGCCCCTTCTTCTGGCTCTTTGAGAATGTGGTGGCCATGGGCGTTAGTGACAAGAGGGACATCTCGCGATTTCTTGAGTCTAACCCCGTGATGATCGACGCCAAAGAAGTGTCTGCAGCACACAGGGCCCGGTACTTCTGGGGTAACCTTCCCGGCATGAACAGGCCTTTGGCATCCACTGTGAATGATAAGCTGGAGCTGCAGGAGTGTCTGGAGCACGGCAGAGTAGCCAAGTTCAGCAAAGTGAGGACCATTACCACCAGGTCAAACTCCATAAAGCAGGGCAAAGACCAGCATTTCCCCGTCTTCATGAACGAGAAGGAGGACATCCTGTGGTGCACTGAAATGGAAAGGGTGTTTGGCTTCCCCGTCCACTATACAGATGTCTCCAACATGAGCCGCTTGGCGAGGCAGAGACTGCTGGGCCGATCGTGGAGCGTGCCAGTCATCCGCCATCTCTTCGCTCCGCTGAAGGAATATTTTGCTTGTGTGTAA

>Rat3A

ATGCCCTCCAGCGGCCCCGGGGACACCAGCATCTCCTCTCTGGAGCGGGAGGATGATCGAAAGGAAGGAGAGGAACAGGAGGAGAACCGTGGTAAGGAAGAGCGTCAGGAGCCCAGCGCCACGGCCCGGAAAGTGGGGAGGCCTGGCCGGAAGCGCAAGCACCCACCGGTGGAAAGCAGTGACACCCCCAAGGACCCAGCGGTGACCACCAAGTCTCAGCCCACAGCCCAGGACTCTGGGCCCTCAGATCTGCTACCCAATGGAGACTTGGAGAAGCGGAGTGAACCCCAACCTGAGGAGGGGAGCCCAGCTGCAGGGCAGAAGGGTGGGGCCCCAGCTGAAGGAGAGGG---AACTGAGACCCCACCAGAAGCCTCCAGAGCAGTGGAGAATGGCTGCTGCGTAACCAAGGAAGGCCGTGGAGCCTCTGCGGGAGAGGGCAAAGAACAGAAGCAGACCAACATCGAATCCATGAAAATGGAGGGCTCCCGGGGCCGACTGCGTGGTGGCTTGGGGTGGGAGTCCAGCCTCCGTCAGCGGCCCATGCCAAGACTCACCTTCCAGGCAGGGGACCCCTACTACATCAGCAAACGGAAGCGGGATGAGTGGCTGGCACGTTGGAAAAGGGAGGCTGAGAAGAAAGCCAAGGTGATTGCAGTAATGAATGCTGTGGAGGAAAGCCAGGCCTCTGGGGAGTCCCAGAAGGTGGAGGAGGCCAGCCCTCCTGCAGTGCAGCAGCCCACGGACCCTGCATCTCCCACTGTGGCCACCACCCCTGAGCCAGTAGGGGCTGATGCTGGGGACAAGAATGCTACCAAGGCAGCTGATGACGAGCCCGAGTATGAGGATGGCCGGGGCTTTGGCATTGGAGAGCTGGTGTGGGGGAAACTCCGGGGCTTCTCCTGGTGGCCAGGCCGAATTGTGTCTTGGTGGATGACAGGCCGGAGCCGAGCAGCCGAAGGCACTCGCTGGGTCATGTGGTTCGGAGATGGCAAATTCTCAGTGGTGTGTGTCGAGAAGCTCATGCCCCTGAGCTCCTTCTGCAGTGCGTTCCACCAGGCCACCTACAACAAGCAGCCCATGTACCGCAAAGCCATCTACGAAGTCCTCCAGGTGGCCAGCAGCCGTGCAGGGAAGCTGTTCCCAGCATGTCATGACAGCGATGAAAGTGACACTGGCAAGGCTGTGGAGGTGCAGAACAAGCAGATGATTGAGTGGGCCCTTGGCGGCTTCCAGCCCTCTGGTCCCAAAGGCCTGGAGCCACCAGAAGAGGAGAAGAATCCTTACAAGGAAGTTTACACCGACATGTGGGTGGAGCCTGAGGCAGCTGCTTATGCCCCACCCCCACCAGCCAAGAAACCCAGAAAGAGCACAACAGAGAAACCCAAGGTCAAGGAGATCATTGATGAACGCACAAGAGAACGGCTCGTGTATGAGGTGCGCCAGAAGTGCCGAAACATCGAGGACATTTGTATCTCATGTGGGAGCCTCAATGTTACCCTGGAGCACCCACTCTTCATTGGTGGAATGTGCCAGAACTGTAAGAACTGCTTCTTGGAGTGTGCTTACCAATACGATGACGATGGGTACCAGTCCTACTGTACCATCTGCTGTGGGGGGCGCGAAGTGCTCATGTGTGGGAACAACAACTGCTGCAGGTGCTTTTGTGTGGAGTGTGTGGATCTCTTGGTGGGGCCAGGGGCTGCCCAAGCAGCCATTAAGGAAGACCCCTGGAACTGCTACATGTGTGGGCACAAGGGCACGTATGGGCTGCTGCGGAGACGGGAGGACTGGCCTTCTCGACTCCAGATGTTCTTCGCCAATAACCACGACCAGGAATTTGACCCCCCGAAGGTTTACCCACCTGTGCCAGCTGAGAAGAGGAAGCCCATCCGGGTGCTATCTCTCTTTGATGGGATTGCTACAGGGCTCCTGGTGCTGAAGGACCTGGGCATCCAAGTGGACCGCTACATCGCCTCCGAGGTGTGCGAGGACTCCATCACAGTGGGCATGGTGCGGCACCAGGGAAAGATCATGTACGTCGGGGACGTCCGCAGCGTCACACAGAAGCATATCCAGGAGTGGGGCCCATTCGATCTGGTGATTGGGGGCAGTCCCTGCAATGACCTCTCCATCGTCAACCCTGCCCGCAAGGGACTTTACGAGGGCACGGGCCGCCTCTTCTTTGAGTTCTACCGCCTCCTGCATGACGCGCGGCCCAAGGAGGGAGATGATCGCCCCTTCTTCTGGCTTTTTGAGAATGTGGTGGCCATGGGCGTTAGTGACAAGAGGGACATCTCGAGATTTCTTGAGTCTAACCCCGTGATGATCGACGCCAAAGAAGTGTCTGCTGCACACAGGGCCCGTTATTTCTGGGGTAACCTTCCTGGCATGAACAGGCCATTGGCATCCACTGTGAATGATAAGCTGGAGTTGCAAGAGTGTCTGGAACACGGCAGAATAGCCAAGTTCAGCAAAGTGAGGACCATTACCACCAGGTCAAACTCCATAAAGCAGGGCAAAGACCAGCATTTCCCTGTCTTCATGAATGAGAAGGAAGACATCCTCTGGTGCACTGAAATGGAAAGGGTGTTTGGCTTCCCTGTCCACTACACAGACGTCTCCAACATGAGCCGCTTGGCGAGGCAGAGACTGCTGGGCCGATCGTGGAGCGTGCCAGTCATCCGCCACCTCTTCGCTCCGCTGAAGGAATATTTTGCTTGTGTGTAA

>DeerMouse3A

ATGCCCTCCAGCGGCCCCGGGGACACCAGCAGCTCCGCTCTGGAGCGGGAGGATGATCGGAAGGAAGGAGAGGAACAGGAGGAGAATCGTGGCAAGGAGGAGCGCCAGGAGCCCAGCACCACGGCCCGGAAGGTGGGGAGGCCTGGCCGGAAGCGCAAGCACCCACCGGTGGAAACCAGTGACACCCCCAAGGACCCCGCAGTGACCAGCAAGTCTCAGCCCATGGCCCAGGACTCTGGCTCCTCAGATCTGTTACCCAATGGAGACTTGGAAAAGCGGAGTGAACCCCAGCCTGAGGAGGGGAGCCCGGCCGCAGGGCAGAAGGGTGGGGCCCCAGCTGAAGGAGAGGG---AACGGAGACCCCGCCGGAATCCTCCCGAGCCGTGGAGAATGGCTGCTGTACCACCAAGGAAGGCCGCGGAGCCTCTGCGGAAGAGGGCAAAGAACAGAAGCAGACCAACATCGAATCCATGAAAATGGAGGGATCCCGTGGACGGCTGCGGGGTGGTCTGGGCTGGGAGTCCAGCCTCCGTCAGCGGCCCATGCCAAGACTCACCTTCCAGGCAGGGGACCCCTACTACATCAGCAAGCGGAAACGGGACGAGTGGCTGGCACGTTGGAAAAGGGAGGCTGAGAAGAAAGCCAAGGTCATTGCAGTAATGAATGCGGTGGAAGAAAACCAGGCCTCTGGAGAGCCTCAGAAGGTAGAGGAGGCTAGCCCTCCTGCTGTGCAGCAGCCCACAGACCCTGCGTCCCCTACTGTGGCCACCACCCCTGAGCCAGTGGGGGCTGATGCTGGGGACAAGAATGCCACCAAAGCAGCCGACGATGAGCCCGAGTATGAGGACGGCCGTGGCTTTGGCATTGGAGAGCTGGTGTGGGGGAAGCTCCGGGGCTTCTCCTGGTGGCCAGGCCGGATCGTGTCTTGGTGGATGACAGGCCGGAGCCGGGCCGCTGAAGGCACTCGCTGGGTCATGTGGTTCGGAGATGGCAAGTTCTCGGTGGTGTGCGTCGAGAAGCTCATGCCCCTGAGCTCCTTCTGCAGCGCCTTCCACCAGGCCACCTACAACAAGCAGCCCATGTACCGCAAAGCCATCTACGAAGTCCTCCAGGTGGCCAGCAGCCGTGCTGGGAAGCTGTTTCCGGCCTGCCACGACAGTGACGAAAGTGACACCGGGAAGGCTGTGGAGGTGCAGAACAAGCAGATGATCGAATGGGCCCTCGGGGGCTTCCAGCCCTCTGGTCCCAAGGGCCTAGAGCCACCGGAAGAGGAGAAGAATCCATACAAGGAGGTCTACACGGACATGTGGGTTGAGCCCGAGGCGGCTGCGTATGCCCCTCCCCCACCAGCCAAGAAACCCAGAAAGAGCACAGCAGAAAAGCCTAAGGTCAAGGAGATCATTGACGAGCGCACAAGAGAGCGGCTGGTGTATGAGGTGCGGCAGAAGTGTCGGAACATCGAGGACATTTGTATCTCATGTGGAAGCCTCAATGTCACCCTGGAGCACCCACTCTTCATTGGTGGAATGTGCCAGAACTGTAAGAACTGCTTCTTGGAGTGCGCCTACCAGTATGACGATGATGGGTACCAATCGTACTGCACCATCTGCTGTGGGGGGCGTGAAGTGCTCATGTGTGGGAACAATAACTGCTGCAGGTGCTTTTGTGTCGAGTGTGTGGATCTCCTGGTGGGGCCAGGAGCTGCCCAGGCGGCCATTAAGGAAGACCCCTGGAACTGCTATATGTGTGGGCACAAGGGCACCTATGGACTGCTGCGGAGACGGGAAGACTGGCCTTCCAGGCTCCAGATGTTCTTTGCCAATAACCATGACCAGGAATTTGACCCCCCAAAGGTTTACCCACCTGTTCCGGCTGAGAAGAGGAAGCCTATCCGGGTGCTATCTCTCTTTGATGGAATTGCTACAGGGCTCCTGGTGCTAAAGGACCTGGGCATCCAAGTGGACCGCTACATCGCTTCAGAGGTGTGTGAGGACTCCATCACTGTGGGCATGGTGCGGCACCAAGGAAAGATCATGTACGTCGGGGACGTCCGCAGCGTCACACAGAAGCATATCCAGGAGTGGGGCCCATTCGATCTGGTGATTGGGGGCAGTCCCTGCAATGACCTCTCCATCGTCAACCCTGCCCGGAAGGGACTTTATGAGGGTACCGGCCGGCTCTTCTTTGAGTTCTACCGCCTCCTCCATGATGCGCGGCCCAAGGAGGGAGATGACCGCCCCTTCTTCTGGCTCTTTGAGAACGTGGTGGCCATGGGCGTGAGCGACAAGAGGGACATCTCACGATTTCTTGAGTCCAACCCCGTGATGATTGACGCCAAAGAAGTGTCTGCTGCACACAGGGCCCGTTACTTCTGGGGTAACCTTCCTGGCATGAACAGGCCATTGGCATCCACTGTGAATGATAAGCTGGAGCTGCAAGAGTGTCTGGAACACGGCAGGATAGCCAAGTTCAGCAAAGTGAGGACCATTACCACCAGGTCAAACTCTATAAAGCAGGGCAAAGACCAGCATTTCCCCGTCTTCATGAATGAGAAGGAAGACATCCTCTGGTGCACTGAAATGGAAAGGGTGTTCGGCTTCCCTGTCCACTACACAGACGTCTCCAACATGAGCCGATTGGCGAGGCAGAGACTGCTGGGCCGGTCGTGGAGCGTACCGGTCATCCGCCATCTCTTCGCTCCGCTGAAGGAATATTTTGCTTGTGTGTAA

>Gebril3A

ATGCCCTCCAGCGGCCCCGGGGACACCAGCAGCTCCTCTCTGGAGCGGGAGGATGATCGAAAGGAAGGAGAGGAACAGGAGGAGAATCGTGGCAAGGAGGAGCGCCAAGAGCCCACCACCACAGCCCGGAAGGTGGGGAGGCCTGGCCGGAAGCGCAAGCACCCACTGGTGGAGAGCAGTGACACCCCCAAGGACCCTGCAGTGACCACCAAGTCTCAGCCCATGGCCCAGGACTCTGGCCCCTCAGATCTGTTACCCAATGGAGACTTGGAGAAGCGGAGTGAACCCCAACCTGAGGAGGGGAGCCCAGCTGCAGGGCAGAAGGGTGGGGCCCCAGCTGAAGGAGAGGG---AACTGAGACCCCACCGGAAGCCTCCAGAGCTGTGGAGAATGGTTGCTGTGCAACCAAGGAGGGTCGTGGAGCCTCTGGGGAAGAGGGCAAAGAACAGAAGCAGACCAACATCGAATCCATGAAAATGGAGGGCTCCCGGGGCCGACTGCGGGGTGGCTTGGGCTGGGAGTCCAGCCTCCGTCAGCGGCCCATGCCAAGACTCACCTTCCAAGCAGGGGACCCCTACTACATCAGCAAACGGAAACGGGACGAGTGGCTGGCACGTTGGAAAAGGGAGGCTGAGAAGAAAGCCAAGGTAATTGCAGTAATGAATGCTGTGGAAGAAAACCAGGCCTCTGGAGAGTCTCAGAAGGTGGAGGAGGCCAGCCCTCCTGCTGTGCAGCAGCCCACGGACCCTGCGTCCCCTACTGTGGCCACCACCCCTGAGCCAGTCGGGGCTGATGCTGGGGACAAGAATGCCACCAAAGCAGCTGATGATGAGCCTGAGTATGAGGATGGCCGGGGCTTCGGCATTGGAGAGCTGGTATGGGGGAAACTTCGGGGCTTCTCCTGGTGGCCAGGCCGAATTGTGTCTTGGTGGATGACGGGCCGGAGCCGAGCCGCTGAAGGCACTCGCTGGGTCATGTGGTTTGGAGACGGCAAGTTCTCAGTGGTGTGCGTGGAGAAGCTCATGCCCCTGAGCTCCTTCTGCAGTGCGTTCCACCAGGCCACCTACAACAAGCAGCCCATGTACCGCAAAGCCATCTACGAAGTCCTCCAGGTGGCTAGCAGCCGTGCTGGGAAGCTCTTTCCAGCCTGCCATGACAGTGACGAAAGTGACACTGGCAAGGCTGTGGAGGTGCAGAACAAGCAGATGATTGAATGGGCCCTCGGAGGGTTCCAGCCATCTGGTCCTAAGGGCCTAGAGCCACCAGAAGAAGAGAAGAACCCATACAAGGAAGTTTACACGGACATGTGGGTTGAGCCAGAGGCCGCTGCTTACGCCCCACCCCCACCAGCCAAGAAACCTAGAAAGAGCACAACAGAGAAGCCCAAGGTCAAGGAGATCATTGATGAGCGCACAAGAGAGCGGCTGGTGTATGAGGTGCGGCAGAAGTGCCGGAACATCGAGGACATCTGCATCTCGTGTGGAAGCCTCAACGTCACCCTGGAGCACCCGCTCTTCATTGGTGGAATGTGCCAGAACTGTAAGAACTGCTTCTTGGAGTGCGCGTACCAGTATGACGATGACGGGTACCAGTCCTACTGCACCATCTGCTGTGGGGGCCGGGAAGTGCTCATGTGTGGGAACAATAACTGCTGCAGGTGCTTTTGTGTGGAGTGTGTGGACCTTTTGGTGGGGCCAGGGGCTGCCCAAGCCGCCATTAAGGAAGACCCCTGGAACTGCTACATGTGTGGGCACAAGGGCACCTACGGGCTGCTGCGGAGACGGGAGGACTGGCCTTCTCGGCTCCAGATGTTCTTTGCCAACAACCACGACCAGGAATTTGATCCCCCGAAGGTTTACCCGCCTGTTCCAGCTGAGAAGAGGAAGCCCATCCGGGTGCTGTCTCTCTTTGATGGAATTGCTACAGGGCTCCTGGTGCTGAAGGACCTGGGCATCCAAGTGGACCGCTACATCGCCTCCGAGGTGTGTGAGGACTCCATCACGGTGGGCATGGTGCGGCACCAGGGAAAGATCATGTACGTCGGGGACGTCCGCAGCGTCACACAGAAGCATATCCAGGAATGGGGCCCATTCGATCTGGTGATTGGGGGCAGTCCCTGCAATGACCTCTCCATCGTCAATCCCGCCCGCAAGGGACTTTACGAGGGTACTGGCCGCCTCTTCTTTGAATTCTACCGCCTCCTGCATGATGCACGGCCCAAGGAAGGAGACGACCGTCCCTTCTTCTGGCTCTTTGAGAATGTGGTGGCCATGGGCGTTAGTGACAAGAGGGACATCTCACGATTCCTTGAGTCTAACCCTGTGATGATTGACGCCAAAGAAGTTTCTGCTGCACACAGGGCCCGTTACTTCTGGGGTAACCTTCCTGGCATGAACAGGCCATTGGCATCCACTGTGAACGATAAGCTGGAGCTGCAGGAGTGTCTGGAACATGGCAGAATAGCCAAGTTCAGCAAAGTGAGGACCATTACCACCAGGTCAAACTCCATAAAGCAGGGCAAAGACCAGCATTTCCCCGTCTTCATGAATGAGAAGGAAGACATCCTCTGGTGCACTGAAATGGAAAGGGTGTTTGGCTTCCCTGTCCACTACACAGACGTCTCCAACATGAGCCGCTTGGCGAGGCAGAGACTGCTGGGCCGGTCGTGGAGCGTGCCAGTCATCCGCCACCTCTTCGCTCCGCTGAAGGAATATTTTGCTTGTGTGTAA

>ChineeseHamster3A

ATGCCCTCCAGCGGTCCCGGGGACACCAGCAGCTCCACTTTGGAGCGGGAGGATGATCGAAAGGAAGGAGAGGAACAGGAGGAGAGTCGTGGCAAGGAAGAGCGCCAGGAACCCAGCACCACGGCCCGGAAGGTGGGGAGGCCTGGCCGGAAGCGCAAACACCCACCGGTGGAAAGTAGTGACACACCCAAGGACTCTGCCGTGACTAGCAAGTCTCAGCCCATGGCCCAGGACTCTGGCTCCTCAGATCTGTTACCCAATGGAGACTTGGAAAAGCGGAGTGAACCCCAACCTGAGGAGGGGAGCCCAGCTGCAGGGCAGAAGGGTGGGGCCCCTGCTGAAGGAGAGGG---AACTGAGACCCCACCAGAATCTTCCCGAGCCGTGGAAAATGGCTGCTGCACAACCAAGGAGGGCCGGGGAGCCTCTGCCGAAGAGGGCAAAGAACAGAAGCAGACCAACATTGAATCCATGAAAATGGAGGGCTCCCGGGGCAGACTGCGGGGTGGCTTGGGCTGGGAGTCCAGTCTCCGTCAGCGGCCAATGCCAAGACTCACCTTCCAGGCAGGGGACCCCTACTACATCAGCAAACGGAAACGGGACGAGTGGCTGGCACGTTGGAAAAGGGAGGCTGAGAAGAAAGCCAAGGTAATTGCAGTAATGAATGCTGTGGAGGAAAACCAGGCCTCTGGAGAGCCTCAGAAGGTAGAGGAGGCCAGCCCTCCTGCTGTGCAGCAGCCCACCGACCCTGCATCCCCTACTGTGGCCACCACTCCTGAGCCAGTGGGGGCTGATGCTGGGGACAAGAATGCCACCAAAGCAGCTGATGATGAGCCTGAGTATGAGGATGGCCGGGGCTTTGGCATCGGAGAGTTGGTGTGGGGGAAACTTCGAGGCTTCTCCTGGTGGCCAGGCCGAATTGTGTCTTGGTGGATGACAGGCCGGAGCCGAGCCGCGGAAGGCACTCGCTGGGTCATGTGGTTTGGAGATGGCAAGTTCTCAGTGGTGTGTGTCGAGAAACTCATGCCACTGAGCTCCTTCTGCAGTGCGTTCCACCAGGCCACCTACAATAAGCAGCCCATGTACCGCAAAGCTATCTACGAAGTTCTCCAGGTGGCCAGCAGCCGTGCTGGGAAGCTGTTTCCAGCTTGCCATGACAGTGACGAAAGTGACACTGGCAAGGCTGTGGAGGTGCAGAACAAGCAGATGATTGAATGGGCCCTTGGAGGGTTCCAGCCCTCTGGTCCCAAGGGCCTGGAGCCACCAGAAGAGGAGAAGAATCCATACAAGGAAGTTTACACAGACATGTGGGTTGAGCCCGAGGCAGCTGCATATGCTCCACCCCCACCAGCCAAGAAACCCAGAAAGAGCACAACAGAAAAGCCTAAGGTCAAGGAGATCATTGATGAACGCACAAGAGAGCGGCTGGTGTATGAGGTGCGTCAGAAGTGCCGGAACATCGAGGACATTTGTATCTCATGTGGAAGCCTCAATGTCACCCTGGAGCACCCACTCTTCATTGGTGGAATGTGCCAGAACTGTAAGAACTGCTTCTTGGAGTGCGCGTACCAGTACGACGATGACGGGTACCAGTCCTACTGCACCATCTGCTGTGGGGGGCGTGAGGTGCTCATGTGTGGCAACAACAACTGTTGCAGGTGCTTTTGTGTCGAGTGTGTGGATCTCTTGGTGGGGCCAGGAGCTGCCCAAGCGGCCATTAAGGAAGACCCCTGGAACTGCTACATGTGTGGCCACAAGGGCACCTATGGGCTGCTGCGGAGACGGGAAGACTGGCCTTCCAGGCTCCAGATGTTCTTTGCCAATAACCATGACCAGGAATTTGACCCCCCGAAGGTTTACCCACCTGTTCCAGCTGAGAAGAGGAAGCCCATCCGGGTGCTGTCTCTCTTTGATGGAATTGCTACAGGGCTCCTGGTGCTGAAGGACCTGGGCATCCAAGTGGACCGCTACATTGCCTCAGAGGTGTGTGAGGACTCCATCACGGTGGGCATGGTGCGGCACCAGGGAAAGATCATGTACGTCGGGGACGTCCGCAGCGTCACACAGAAGCATATCCAGGAGTGGGGCCCATTCGATCTGGTGATTGGGGGCAGTCCCTGCAATGACCTCTCCATCGTCAACCCTGCCCGAAAGGGACTTTACGAGGGTACTGGCCGGCTCTTCTTTGAGTTCTACCGCCTCCTGCATGATGCTCGACCTAAGGAGGGAGATGACCGCCCCTTCTTCTGGCTCTTTGAGAATGTGGTGGCCATGGGCGTTAGTGACAAGAGGGACATCTCACGATTTCTTGAGTCCAACCCCGTGATGATTGACGCCAAAGAAGTGTCTGCTGCACACAGGGCCCGTTACTTCTGGGGTAACCTTCCTGGCATGAACAGGCCATTGGCATCCACTGTGAATGATAAGCTGGAGCTGCAAGAGTGTCTGGAACATGGCAGAATAGCCAAGTTCAGCAAAGTAAGGACCATCACCACCAGGTCAAATTCCATAAAGCAAGGCAAAGACCAGCATTTCCCCGTCTTCATGAACGAGAAGGAGGACATCCTCTGGTGCACTGAAATGGAAAGGGTGTTTGGCTTCCCTGTCCACTACACAGACGTCTCAAACATGAGCCGCTTGGCGAGGCAGAGACTGCTGGGCCGATCGTGGAGCGTGCCAGTCATCCGCCACCTCTTCGCTCCGCTGAAGGAATATTTTGCTTGTGTGTAA

>GoldenHamster3A

ATGCCCTCCAGCGGCCCCGGGGACACCAGCAACTCCGCTCTGGAGCGGGAGGATGATCGCAAGGAAGGAGAGGAACAGGAGGAGAATCGTGGCAAGGAAGAGCGTCAGGAACCCAGCACCACGACCCGGAAGGTGGGGAGGCCTGGCCGGAAGCGCAAGCACCCGCCGGTGGAGAGTAGTGACACACCCAAGGACTCTGCCGTGACCAGCAAGTCTCAGCCGATGGCCCAGGACTCTGGTTCCTCAGATCTGTTACCCAATGGAGACTTGGAAAAGCGGAGTGAACCCCAACCTGAGGAGGGGAGCCCAGCTGCAGGGCAGAAGGGTGGGGCCCCCGCTGAAGGAGAGGG---AACTGAGACCCCACCAGAATCCTCCCGAGCTGTGGAAAATGGCTGCTGCACAACCAAGGAGGGCCGGGGAGCCTCTGCTGAAGAGGGCAAAGAACAGAAGCAGACCAACATCGAGTCCATGAAAATGGAGGGCTCCCGGGGCAGACTGCGGGGTGGCTTGGGCTGGGAGTCCAGCCTCCGTCAGCGGCCCATGCCAAGACTCACCTTCCAGGCAGGGGACCCCTACTACATCAGCAAACGGAAACGGGACGAGTGGCTGGCACGGTGGAAAAGGGAGGCTGAGAAGAAAGCCAAGGTAATTGCAGTAATGAATGCTGTGGAAGAAAACCAGGCCTCTGGAGAGCCTCAGAAGGTAGAGGAGGCCAGCCCTCCTGCAGTGCAACAGCCCACTGACCCTGCATCCCCTACTGTGGCCACCACTCCTGAGCCCGTGGGGGCTGATGCTGGGGACAAGAACGCCACCAAAGCAGCTGACGATGAGCCTGAGTATGAGGATGGCCGGGGCTTTGGCATCGGAGAGCTGGTGTGGGGGAAACTTCGGGGCTTCTCCTGGTGGCCGGGCCGAATTGTGTCTTGGTGGATGACAGGCCGGAGCCGAGCAGCCGAAGGCACTCGCTGGGTCATGTGGTTTGGAGATGGCAAGTTCTCAGTGGTGTGCGTCGAGAAGCTCATGCCACTGAGCTCCTTCTGCAGTGCGTTCCACCAGGCCACCTACAATAAGCAGCCCATGTACCGCAAAGCTATCTACGAAGTCCTCCAGGTGGCCAGCAGCCGTGCCGGGAAGCTGTTTCCAGCTTGCCATGACAGTGACGAAAGTGACACTGGCAAGGCTGTGGAAGTGCAGAACAAGCAGATGATTGAATGGGCCCTTGGGGGTTTCCAGCCCTCTGGTCCCAAGGGCCTGGAGCCGCCAGAAGAGGAGAAGAATCCGTACAAGGAAGTTTACACGGACATGTGGGTTGAGCCTGAGGCAGCTGCGTATGCCCCACCCCCACCAGCCAAAAAACCCAGAAAGAGCACAACAGAGAAGCCTAAGGTCAAGGAGATTATTGATGAACGCACAAGAGAGCGGCTGGTGTACGAGGTCAGGCAGAAGTGCCGGAACATCGAGGACATCTGTATCTCGTGTGGAAGCCTCAATGTCACCCTGGAGCACCCACTCTTCATTGGTGGAATGTGCCAGAACTGTAAGAACTGCTTCTTGGAGTGCGCATACCAGTATGACGATGACGGGTACCAGTCCTACTGCACCATCTGCTGTGGGGGGCGGGAAGTGCTCATGTGTGGCAACAACAACTGCTGCAGGTGCTTTTGTGTTGAGTGTGTGGATCTCTTGGTGGGACCAGGAGCTGCCCAAGCGGCCATCAAGGAGGACCCCTGGAACTGCTACATGTGTGGCCACAAGGGCACCTATGGGCTGCTGCGGAGACGGGAAGACTGGCCTTCCAGGCTCCAGATGTTCTTTGCCAATAACCATGACCAGGAATTTGACCCCCCAAAGGTTTACCCACCTGTTCCAGCTGAGAAGAGGAAACCCATCCGGGTGCTGTCTCTCTTTGATGGGATCGCCACAGGGCTCCTGGTGCTGAAGGACCTGGGCATCCAAGTGGACCGCTACATCGCCTCAGAGGTGTGTGAGGACTCCATCACGGTGGGCATGGTGCGGCACCAGGGAAAGATCATGTACGTCGGGGACGTCCGCAGCGTCACACAGAAGCATATCCAGGAGTGGGGCCCATTCGATTTGGTGATTGGGGGCAGTCCCTGCAATGACCTCTCCATCGTCAACCCTGCCCGAAAGGGACTTTACGAGGGCACTGGCCGGCTCTTCTTTGAGTTCTACCGCCTCCTGCATGATGCGCGACCCAAGGAGGGAGATGACCGCCCCTTCTTCTGGCTCTTTGAGAATGTGGTGGCCATGGGCGTTAGTGACAAGAGGGACATCTCACGATTTCTTGAGTCCAACCCTGTGATGATTGATGCCAAAGAAGTGTCTGCTGCACACAGGGCCCGTTACTTCTGGGGTAACCTTCCTGGCATGAACAGGCCATTGGCATCCACTGTGAATGATAAGCTGGAGCTGCAAGAATGTCTGGAACATGGCAGAATAGCCAAGTTCAGCAAAGTGAGGACCATCACCACCAGGTCAAACTCCATAAAGCAGGGCAAAGACCAGCATTTCCCTGTCTTCATGAACGAGAAGGAGGACATCCTCTGGTGCACTGAAATGGAAAGGGTGTTTGGCTTCCCTGTCCACTACACAGACGTCTCCAACATGAGCCGCTTGGCGAGGCAGAGACTGCTGGGCCGGTCGTGGAGCGTGCCCGTCATCCGCCACCTCTTCGCTCCGCTGAAGGAATATTTTGCTTGTGTGTAA

>MountBlindMole3A

ATGCCCTCCAGCGGCCCCGGGGACAACAGCAGCTCTGCTCCTGAGCGGGAGGAAGACCGGAAGGAAGAAGAG---CAGGAGGAGAATCGTGGCAAAGAGGAACGCCAGGAGCCCAGTGCCACGGCCCGGAAGGTGGGGAGGCCTGGCCGGAAGCGCAAGCACCCACTGGTGGAAAGCAGCGACACACCCAAGGACCCTGCTGTGACCTCCAAGTCCCCACCCATGGCCCAGGACTCAGGCTCTTCAGAGCTGTTACCCAATGGAGACTTGGAGAAGCGGAGTGAGCCCCAGCCTGAGGAGGGGAGTCCTGCTGCAGGGCAAAAAGGTGGGGCCCCAGCTGAGGGAGAGGGTGCAGCTGAGACCCCACCAGAGTCCTCCAGAGCTGTGGAGAATGGCTGCTGTGTACCCAAGGAGGGCCGAGGAGCCTCTGCAGAGGAAGGTAAAGAACAGAAGGAGACCAACATCGAATCCATGAAAATGGAGGGCTCCCGGGGCCGACTCCGTGGTGGCTTGGGTTGGGAGTCCAGCCTCCGCCAGCGGCCCATGCCAAGGCTCACCTTCCAGGCGGGGGATCCCTACTACATCAGCAAACGCAAGCGGGATGAGTGGCTGGCACGCTGGAAAAGGGAGGCTGAGAAGAAAGCCAAGGTAATTGCAGTAATGAATGCTGTGGAAGAGAACCAGGGCTCTGGGGAGCCTCAGAAGGTGGAGGAGGCCAGCCCTCCTACTGTGCAGCAGCCTACAGACCCTGCATCCCCTACTGTGGCCACCACACCTGAGCCTGTGGGGGCTGATGCTGGGGACAAGAATGCCACCAAAGCAGCTGATGATGAGCCAGAGTATGAGGACGGCCGGGGTTTTGGCATTGGGGAGCTGGTGTGGGGGAAACTGCGAGGCTTCTCCTGGTGGCCAGGCCGAATTGTGTCTTGGTGGATGACGGGCCGGAGCCGAGCAGCTGAAGGCACCCGCTGGGTCATGTGGTTTGGCGACGGCAAGTTCTCAGTGGTGTGTGTGGAGAAGCTGATGCCCCTGAGCTCCTTCTGCAGTGCATTCCACCAGGCAACCTACAACAAGCAGCCCATGTACCGCAAAGCCATCTATGAAGTACTCCAGGTGGCCAGCAGCCGTGCAGGGAAGCTGTTCCCGGCCTGTCATGACAGTGATGAAAGTGACACTGGCAAGGCTGTGGAGGTGCAGAACAAGCAGATGATCGAATGGGCCCTTGGGGGGTTCCAGCCCTCTGGTCCCAAGGGCCTGGAGCCACCAGAAGAAGAGAAGAATCCCTATAAGGAAGTTTACACAGACATGTGGGTTGAGCCTGAGGCAGCTGCCTATGCACCACCCCCACCAGCCAAAAAGCCCAGAAAGAGCACAACTGAGAAGCCCAAGGTCAAGGAGATTATCGATGAACGCACAAGAGAGCGGCTGGTGTACGAGGTGCGGCAGAAGTGCCGAAACATCGAAGACATTTGTATCTCATGTGGAAGCCTCAACGTCACCCTGGAGCACCCACTTTTTATTGGTGGAATGTGCCAGAACTGCAAGAACTGCTTCTTGGAGTGTGCCTACCAATATGATGATGATGGGTACCAGTCCTATTGCACCATCTGCTGTGGGGGCCGTGAGGTGCTCATGTGCGGGAACAACAACTGCTGCAGGTGCTTTTGTGTTGAGTGTGTAGATCTCTTGGTGGGGCCAGGAGCTGCCCAAGCAGCCATTAAGGAAGACCCCTGGAACTGCTACATGTGTGGGCACAAGGGCACCTATGGGCTGCTGCGGCGGCGAGAAGACTGGCCTTCTCGGCTCCAGATGTTCTTCGCCAATAATCATGACCAGGAATTTGACCCCCCAAAAGTTTACCCACCTGTCCCTGCCGAGAAAAGGAAGCCTATCCGGGTGCTATCTCTCTTTGATGGAATTGCTACAGGGCTCCTGGTGCTAAAGGACCTGGGCATCCAAGTGGACCGCTACATTGCCTCAGAGGTGTGTGAGGACTCCATCACGGTGGGAATGGTGCGGCACCAGGGGAAGATCATGTACGTCGGGGACGTCCGCAGCGTCACACAGAAGCATATCCAGGAGTGGGGCCCATTTGATCTGGTGATTGGGGGCAGTCCCTGCAATGACCTCTCTATTGTGAACCCTGCCCGCAAGGGACTTTACGAGGGAACTGGACGGCTCTTCTTTGAGTTCTACCGCCTCCTGCATGATGCGCGGCCCAAGGAGGGAGATGATCGCCCCTTCTTCTGGCTCTTTGAGAACGTGGTGGCCATGGGCGTTAGTGACAAGAGGGACATCTCACGATTTCTCGAGTCCAACCCTGTGATGATTGATGCCAAAGAAGTGTCAGCTGCACACAGGGCCCGCTACTTCTGGGGCAACCTTCCCGGTATGAACAGGCCATTGGCATCCACTGTGAATGATAAGCTGGAGCTGCAGGAGTGTCTGGAACATGGCAGAATAGCCAAGTTCAGTAAAGTGAGGACCATTACCACCAGGTCGAACTCCATAAAGCAAGGCAAAGACCAGCATTTCCCTGTCTTCATGAATGAGAAGGAGGACATCCTATGGTGCACTGAAATGGAAAGGGTGTTTGGCTTCCCTGTCCACTACACAGACGTCTCCAACATGAGCCGCTTGGCAAGGCAGAGACTGCTGGGCCGGTCATGGAGTGTGCCAGTCATCCGCCACCTCTTCGCTCCGCTGAAGGAATATTTTGCTTGTGTGTAA
