## Supplementary material for "Dynamic evolution of *de novo* DNA methyltransferases in rodent and primate genomes": Suppl data 4

*SupFile_4: Muroid Dnmt3B sequence alignments*

>Mouse3B

ATGAAGGGAGACAGCAGACATCTGAATGAAGAAGAGGGTGCCAGCGGGTATGAGGAGTGCATTATCGTTAATGGGAACTTCAGTGACCAGTCCTCAGACACGAAGGATGCTCCCTCACCCCCAGTCTTGGAGGCAATCTGCACACCAGAGACCAGAGGCCGCAGGTCAAGCTCCCGGCTGTCTAAGAGGGAGGTCTCCAGCCTTCTGAATTACACGCAGGACGGAGATGGAGACGAAGTAGATGATGGGAATGGCTCTGATATTCTAATGCCAAAGCTCACCCGTGAGACCAGGACGCGCTCTGAAAGCCCGGCTGTCCGAACCCGACATAGCAATGGGACCTCCAGCTTGGAGAGGCAAAGAGCCTCCCCCAGAATCACCCGAGGTCGGCAGGGCCGCCACCATGTGCAGGAGTACCCTGTGGAGTTTCCGGCTACCAGGTCTCGGAGACGTCGAGCATCATCTTCAGCAAGCACGCCATGGTCATCCCCTGCCAGCTTCATGGAAGAAGTGACACCTAAGAGCGTCAGTACCCCATCAGTTGACTTGAGCCAGGATGGAGATCAGGAGGGTATGGATACCACACAGGTGGATGCAGAGAGCAGAGATGGAGACAGCACAGAGTATCAGGATGATAAAGAGTTTGGAATAGGTGACCTCGTGTGGGGAAAGATCAAGGGCTTCTCCTGGTGGCCTGCCATGGTGGTGTCCTGGAAAGCCACCTCCAAGCGACAGGCCATGCCCGGAATGCGCTGGGTACAGTGGTTTGGTGATGGCAAGTTTTCTGAGATCTCTGCTGACAAACTGGTGGCTCTGGGGCTGTTCAGCCAGCACTTTAATCTGGCTACCTTCAATAAGCTGGTTTCTTATAGGAAGGCCATGTACCACACTCTGGAGAAAGCCAGGGTTCGAGCTGGCAAGACCTTCTCCAGCAGTCCTGGAGAGTCACTGGAGGACCAGCTGAAGCCCATGCTGGAGTGGGCCCACGGTGGCTTCAAGCCTACTGGGATCGAGGGCCTCAAACCCAACAAGAAGCAACCAGAGAACAAAAGTCGAAGACGCACAGACTCTGCTGCTTCTGAGTCCCCCCCACCCAAGCGCCTCAAGACAAATAGCTATGGCGGGAAGGACCGAGGGGAGGATGAGGAGAGCCGAGAACGGATGGCTTCTGAAGTCACCAACAACAAGGGCAATCTGGAAGACCGCTGTTTGTCCTGTGGAAAGAAGAACCCTGTGTCCTTCCACCCCCTCTTTGAGGGTGGGCTCTGTCAGAGTTGCCGGGATCGCTTCCTAGAGCTCTTCTACATGTATGATGAGGACGGCTATCAGTCCTACTGCACCGTGTGCTGTGAGGGCCGTGAACTGCTGCTGTGCAGTAACACAAGCTGCTGCAGATGCTTCTGTGTGGAGTGTCTGGAGGTGCTGGTGGGCGCAGGCACAGCTGAGGATGCCAAGCTGCAGGAACCCTGGAGCTGCTATATGTGCCTCCCTCAGCGCTGCCATGGGGTCCTCCGACGCAGGAAAGATTGGAACATGCGCCTGCAAGACTTCTTCACTACTGATCCTGACCTGGAAGAATTTTACTTGGTGCTCAAGGAGTTGGGTATTAAAGTGGAAAAGTACATTGCCTCCGAAGTCTGTGCAGAGTCCATCGCTGTGGGAACTGTTAAGCATGAAGGCCAGATCAAATATGTCAATGACGTCCGGAAAATCACCAAGAAAAATATTGAAGAGTGGGGCCCGTTCGACTTGGTGATTGGTGGAAGCCCATGCAATGATCTCTCTAACGTCAATCCTGCCCGCAAAGGTTTATATGAGGGCACAGGAAGGCTCTTCTTCGAGTTTTACCACTTGCTGAATTATACCCGCCCCAAGGAGGGCGACAACCGTCCATTCTTCTGGATGTTCGAGAATGTTGTGGCCATGAAAGTGAATGACAAGAAAGACATCTCAAGATTCCTGGCATGTAACCCAGTGATGATCGATGCCATCAAGGTGTCTGCTGCTCACAGGGCCCGGTACTTCTGGGGTAACCTACCCGGAATGAACAGG

>Caroli3B

ATGAAGGGAGACAGCAGACATCTGAATGAAGAAGAGGGAGCCAGCGGATTTGAGGAGTGCATAATTGTTAATGGGAACTTCAGTGACCAGTCCTCAGACACGAAGGATGCTCCCTCACCCCCAGTCTTGGAGGCAATCTGCACACCAGAGACCAGAGGCCGCAGGTCAAGCTCCCGGCTGTCTAAGAGGGAGGTCTCCAGCCTTCTGAATTACACGCAGGACGGAGATGGAGATGAAGTGGATGATGGGAATGGCTCTGATATTCTAATGCCAAAGCTCACCCGTGAGACCAGGACGATCTCTGAAAGCCCGGCTGTCCGAACCCGACATAGCAATGGGACCTCCAGCTTGGAGAGGCAAAGAGCCTCCCCCAGAATCACCCGAGGCCGGCAGGGCCGCCACCATGTGCAGGAGTACCCCGTGGAGTTCCCAGCTACCAGGTCTCGGAGACGTCGAGCATCGTCTTCAGCAAGCACGCCGTGGTCATCCCCTGCCAGCTTCATGGAAGAAGCGACACCTAAGAGCGTCAGTACCCCATCAGTTGACTTGAGCCAGGATGGAGATCAGGAGGGCATAGATACCACACAAGTGGATGCAGAGAGCAGAGATGGAGACAGCACAGAGTATCAGGATGATAAAGAGTTTGGAATAGGTGACCTCGTGTGGGGAAAGATCAAGGGCTTCTCCTGGTGGCCTGCCATGGTGGTGTCCTGGAAAGCCACCTCCAAGCGCCAGGCCATGCCCGGAATGCGCTGGGTACAGTGGTTTGGCGATGGCAAGTTTTCTGAGATCTCTGCTGACAAACTGGTGGCTCTGGGGCTGTTCAGCCAGCACTTTAATCTGGCTACCTTCAATAAGCTGGTTTCTTATCGGAAGGCCATGTACCACACTCTGGAGAAAGCCAGGGTACGAGCTGGCAAGACCTTCTCCAGCAGTCCTGGAGAGTCACTGGAGGACCAGCTGAAGCCCATGCTGGAGTGGGCCCACGGCGGCTTCAAGCCCACTGGGATCGAGGGCCTCAAACCCAACAACAAGCAACCAGAGAACAAAAGTCGAAGGCGCACAGACTCTGCTGCTTCTGAGTATCCCCCACCCAAGCGCCTCAAGACCAACAGCTATGGTGGGAAGGACCGAGGGGAGGATGAGGAGAGCCGAGAACGGATGGCTTCTGATGTCACCAACAACAAGGGCAATCTGGAAGACCGCTGTTTGTCCTGTGGAAAGAAGAACCCTGTGTCCTTCCACCCCCTCTTTGAGGGTGGGCTCTGTCAGAGTTGCCGGGATCGCTTCCTAGAGCTCTTCTACATGTATGACGAGGACGGCTATCAGTCCTACTGCACCGTGTGCTGTGAGGGCCGTGAACTGCTGCTGTGCAGTAACACAAGTTGCTGCAGATGCTTCTGTGTGGAGTGTCTGGAGGTGCTGGTGGGCGCAGGCACAGCTGAGGATGCCAAGCTGCAGGAACCCTGGAGCTGCTATATGTGCCTCCCTCAGCGCTGCCATGGGGTCCTCCGACGCAGGAAGGATTGGAACATGCGCCTGCAAGACTTCTTCACTACTGATCCTGACCTGGAAGAATTTTACTTGGTGCTCAAGGAGTTGGGTATTAAAGTGGAGAAGTACATTGCCTCCGAAGTCTGTGCAGAGTCCATCGCTGTGGGAACCATTAAGCATGAAGGCCAAATCAAATATGTCAATGACGTCCGGAAAATCACCAAGAAAAATATTGAAGAGTGGGGCCCGTTCGACTTGGTGATTGGTGGAAGCCCATGCAACGATCTCTCTAATGTCAATCCTGCCCGCAAAGGTTTATATGAGGGCACGGGAAGGCTCTTCTTCGAGTTTTACCACTTGCTGAATTATACCCGCCCCAAGGAGGGCGACAACCGTCCATTCTTCTGGATGTTCGAGAATGTTGTGGCCATGAAAGTGAATGACAAGAAAGACATCTCCCGATTCCTGGCATGTAACCCAGTGATGATTGATGCCATCAAGGTGTCTGCTGCTCACAGGGCCCGGTACTTCTGGGGTAACCTACCGGGAATGAACAGG

>Rat3B

ATGAAGGGAGACAGCAGACATCTTAATGAAGAAGAGGGTGCTAGTGGGTATGAGGACTGTATCATTGTTAATGGGAACTGTAGTGACCAGTCCTCGGACACGAAGGATGCTCCCTCACCCCCAGTCTTGGAGGCAATCTGCACACCAGAGACCAGAGGCCGCAGATCAAGCTCACGGCTGTCTAAGAGGGAGGTCTCCAGCCTTCTGAATTACACGCAGGACGGAGATGGAGATGAAGCGGATGATGGAGATGGCTCTGATATTCTCATGCCAAAGCTCACCCGAGAGACCAGGACTCCCTCTGAAAGCCCAGCCGTCCGAACCCGAAACAGCAACAGTATCTCCAGCCTGGAGAGGCAAAGAACCTCCCCCAGAATCACGCGAGGCCGGCAGGGCCGCTACCACGTTCAGGAGTACCCCGTGGAGTTCCCAGCTACCAAGTCTCGGAGGCGGCGAGCATCGTCTTCAGCAAGCACGCCATGGTCATCCCCTGCCAGCCTCATGGAAGATGTGACACCTAAGAGCAGCAGTACGCCATCGGTTGACTTGAGCCAGGACGGCCCTCAGGAGGGCATGGATGCCACACAGGTGGATGCGGAGAGCAGGGATGGTGACAGCACTGAGTATCAGGATGACAAGGAGTTTGGAATAGGTGACCTTGTGTGGGGAAAGATCAAGGGCTTCTCCTGGTGGCCTGCCATGGTGGTGTCCTGGAAAGCCACCTCCAAGCGCCAGGCCATGCCCGGAATGCGCTGGGTACAGTGGTTTGGCGATGGCAAGTTTTCTGAGATCGCTGCTGACAAGCTGGTGGCTCTGGGTCTGTTCAGCCAGCACTTTAATCTGGCCACCTTCAATAAGCTGGTTTCTTATAGGAAGGCCATGTACCACACTCTGGAGAAAGCCATGGTGCGAGCTGGCAAGACCTTCCCCAGCCGCCCTGGAGACTCACTGGAGGACCAGCTGAAGCCCATGCTGGAGTGGGCCCACGGGGGCTTCAAGCCCACTGGGATCGAGGGCCTCAAACCCAACAACAAGCAACCAGAGAACAAAAGTCGAAGACGCACAGACTTTGCCGCTTCTGAGTACACACCCCCTAAGCGCCTCAAGACAAATAGCTATGGTGGGAAGGACCGAGGGGAAGATGAGGAGAGCCGAGAACGGATGGCTTCTGATGTCACTAACAACAAGGGCAATCTGGAAGACCGCTGTCTGTCCTGCGGTAAGAAGAACCCTGTGTCCTTCCACCCCCTCTTTGAGGGTGGGCTCTGTCAGAGTTGCCGGGATCGCTTCCTGGAGCTCTTCTACATGTACGATGAGGACGGCTATCAGTCCTACTGCACCGTGTGCTGTGAGGGCCGTGAACTGCTTCTGTGCAGTAACACAAGCTGCTGCAGGTGCTTCTGTGTGGAGTGTCTGGAGGTGCTGGTGGGTACAGGCACAGCGGAGGATGCCAAGCTGCAGGAACCCTGGAGCTGCTATATGTGCCTCCCGCAGCGCTGCCATGGGGTCCTGCGGCGCAGGAAGGACTGGAACATGCGCCTGCAAGACTTCTTCACTACTGATCCTGACCTGGAAGAGTTTTACTTGGTGCTCAAGGACTTGGGTATTAAAGTGGAGAAGTACGTTGCCTCTGAAGTCTGTGCAGAGTCCATTGCTGTAGGAACCATTAAGCATGAAGGCCAAATCAAATATGTCAATGACGTCCGGAAAATCACCAAGAAAAATATTGAAGAGTGGGGCCCATTTGACTTGGTGATTGGCGGAAGCCCATGCAATGATCTTTCTAATGTCAATCCTGCCAGGAAAGGCCTGTATGAGGGCACAGGAAGGCTCTTCTTTGAGTTTTACCACTTGCTGAATTATACACGCCCCAAGGAGGGCGACAACCGTCCATTCTTCTGGATGTTCGAGAATGTTGTGGCCATGAAAGTGAATGACAAGAAAGACATCTCCCGATTCCTGGCATGTAACCCAGTGATGATCGACGCCATCAAGGTTTCTGCTGCTCACAGGGCCCGGTACTTCTGGGGTAATCTACCTGGAATGAACAGG

>Gebril3B

ATGAAGGGAGACAGCAGACATCTGAATGAAGAAGAGGGTGCCAGTGGGAGTGAGGACTGTATCATCATTAATGGGAACTGTAGTGACCAGTCCTCAGACACTAAGGATGCTCCCTCACCCCCAGTCCTGGAGGCAATCTGCACACCAGAGAGCAGAGGCCGCAGGTCAAGGTCACGGCTGTCTAAGAGGGAGGTCTCCAGCCTTCTGAGTTACTCTCAGGACACAGATGGAGGCGAGGTGGATGATGATGAAGGTTCTGACATTTTAATGCCAAAGCTCACCCGGGAGACCAGGACATCCTGGGAGAGTCCAGCCGTCCGAACCCGAAATAGCAACAGTAGCACCAGCCTGGAGAGGCAAAGAGCCTCGCCCAGAGTCACCCGCGGCCGGCAAGGCCGCCGCCATGTGCAGGAGTACCCAGTGGAATTTCCAGCTACCAGGTCTCGGAGACGGCGGGCATCTTCTTCAGCAAGCACACCCTGGTCATCCCCTGCAAGCCTCATAGAAGAAGTAACACCTCAGGGCAGCAGTACCCCATCAGTTGACTTGAGCCAGGACGACCACGAGGATGGCATAGATACCACACAGCTGGATGCGGAAAGCCGAGAGGGAGATAGCACTGAATATCAGGATGAGAAGGAGTTTGGAATAGGGGACCTCGTGTGGGGAAAGATCAAGGGCTTCTCCTGGTGGCCTGCCATGGTGGTATCCTGGAAAGCCACCTCCAAGCGCCAGGCCATGCCCGGCATGCGATGGGTACAGTGGTTTGGCGATGGCAAGTTTTCTGAGATCTCTGCTGACAAACTCGTTGCCCTGGGGCTGTTCAGCCAGCACTTTAACCTGGCCACCTTCAATAAGCTGGTTTCTTATAGGAAGGCCATGTACCACACTCTGGAGAAAGCCAGGTTGCGAGCCGGCAAGTCCTTCCCAAGCAGCCCTGGAGACTCACTGGAGGACCAGCTGAAGCCCATGCTGGAGTGGGCCCACGGGGGCTTCAAGCCCACGGGGATCGAGGGCCTCAAACCCAGCAACAAGCAACCAGAGGTCAGAAGTCGGAGACGCACAGACTCTGTCACTTCTGAGTACCCCCCACCCAAGCGCCTCAAGACAAATAGCTATGGCGGGAAAGAGCGAGGGGAAGATGAGGAGAGCCGAGAGCGGATGGCTTCTGATGTCGCCCAGAACAAGGGGAATCTGGAAGACCGCTGTCTGTCCTGCGGTAAGAAGAACCCTGTGTCCTTCCACCCCCTCTTTGAGGGTGGGCTCTGTCAGAATTGCAGAGATCGCTTCCTGGAGCTCTTCTACATGTATGATGAGGATGGCTATCAATCATACTGCACCGTGTGCTGTGAGGGCCGAGAACTGCTTCTGTGCAGCAACACGAGCTGCTGCAGGTGCTTCTGTGTGGAGTGTCTGGAGGTGCTGGTGGGTGCGGGCACAGCCGAGGATGCCAAGCTGCAGGAACCCTGGAGCTGCTATATGTGCCTCCCACAGCGCTGCCACGGGGTCCTCCGGCGCAGGAAGGATTGGAACATGCGCCTGCAAGACTTCTTCACCACTGACCCGGACCTGGAAGAATTTTACTTGGTGCTCAAAGAGTTGGGTATTAAAGTGGAAAAGTACGTTGCCTCTGAAGTCTGTGCAGAGTCCATCGCTGTGGGAACCATTAAGCATGAAGGCCAAATCAAATATGTGGACGACGTTAGGAAAGTCACAAAGGAAAATATTGATGAATGGGGCCCATTCGACCTGGTGATTGGTGGAAGCCCATGCAATGATCTTTCCTGTGTGAATCCTGTCAGGAAAGGCTTGTTTGAGGGTACTGGCCGGCTCTTCTTTGAGTTTTACCGATTGCTAAATTACTCACGCCCTGAAGAGTGTGTTGACCGCCCATTCTTCTGGATGTTTGAGAATGTGGTAGCCATGGAGGTCGGTGATAAGAGAGACATCTCACGATTCCTGGAGTGTAACCCAGTGATGATTGATGCCATCAAGGTTTCTGCTGCTCACAGGGCCCGGTACTTCTGGGGGAACCTACCTGGGATGAACAGG

>DeerMouse3B

ATGAAGGGAGACAGCAGACATCTGAATGAGGAAGAGGGAGCCAGTGGGTGTGAGGACTCTGTCATCGTCAACGGGAACTGCAGTGACCAGTCCTCAGACACTAAGGATGCCCCCTCACCCCCCGTCTTGGAGGCGATCTGCACACCAGAGAGCCGAGGCCGCAGATCAAGCTCACGGCTGTCAAAGAGGGAGGTCTCCAGCCTGCTGTCTTACACTCAGGACGGAGATGGAGATGAGGCAGAGGATGGGGACGGCTCTGACATTGTAATGCCAAAGCTCACGCGTGAGACCAGGTCACCCTCTGAAAGTCCAGCGGTGCGAACCCGAAATAGCAACAGTACCTCCAGCCTGGAGAGGCAAAGAGCCTCCCCTAGAATCACCCGAGGCCGCCAGGGCCGCCACCATGTGCAGGAGTACCCCGTGGAGTTCCCAGCTACCAGGTCTCGGAGACGCCGAGCATCGTCTTCAGCAAGCACACCATGGCCATCCCCTGCCAGCCTCATAGAAGAAGTGACACCTCAGAGCAGCAGCACCCCATCGATTGACCTGAGCCGGGACAGCCCGCAGGAGAGCATGGATGCTACACAGCTGGATGCAGATAGCAAAGACGGAGACAGCACGGAGTATCAGGATGATAAGGAGTTTGGAATAGGTGACCTCGTGTGGGGAAAAATCAAGGGCTTCTCCTGGTGGCCTGCCATGGTGGTATCCTGGAAAGCTACCTCCAAGCGCCAGGCCATGCCCGGCATGCGATGGGTACAGTGGTTTGGTGACGGCAAGTTTTCTGAGGTCTCAGCTGACAAACTCATGGCTCTGGGGCTGTTCAGCCAGCACTTTAACCTGGCCACCTTCAATAAGCTGGTGTCTTATAGGAAGGCCATGTACCACACTCTGGAGAGAGCCAGGGTGCGAGCTGGCAAGACCTTCCCCAGCAGCCCGGGAGACTCGCTGGAGGACCAGCTGAAGCCCATGCTGGAGTGGGCCCATGGGGGCTTCAAGCCCACTGGCATCGAGGGCCTCAAACCAAACAACAAACAACCAGAGAACAAAGGTCGAAGACGCACCGACTCTGCCGCTTCTGAGTACCCCACACCCAAGCGCCTCAAGACAAACAGCTGCGGAGGGAAAGACCGAGGGGAGGATGACGAGAGCCGAGAACGGATGGCTTCTGATGTCACCAACAACAAGGGCAATCTGGAAGACCGCTGTTTGTCCTGCGGTAGGAAGAACCCTGTGTCCTTCCATCCCCTCTTTGAGGGTGGGCTCTGTCAGAGTTGCCGGGATCGCTTCCTGGAGCTCTTCTACATGTATGATGAGGACGGCTATCAGTCCTACTGCACCGTGTGCTGCGAGGGCCGGGAGCTGCTTCTCTGCAGCAACACCAGCTGCTGCAGGTGCTTCTGTGTGGAGTGTCTGGAGGTGCTGGTGGGGACAGGGACAGCCGAGGATGCCAAGCTGAAGGAACCCTGGAGCTGCTATATGTGCATCCCACAGCGCTGCCATGGTGTCCTCAGGCGCAGGAAGGATTGGAACATGCGCCTGCAAGACTTCTTCACGACTGATCCTGACCTCGAAGAATTTTACTTGGTGCTCAAAGAGTTGGGTATCAAAGTGGAAAAGTACGTCGCCTCCGAAGTCTGTTCAGAGTCCATCGCTGTGGGAACCGTTAAGCATGAAGGCCATATCAAATACGTGGATGACGTCAGGAAAATCACAAAGAAAAATATTGAGGAGTGGGGCCCATTTGACTTGGTGATTGGTGGAAGCCCTTGCAATGACCTCTCCAATGTCAATCCCGCCAGGAAAGGCCTATATGAGGGCACCGGCAGGCTCTTCTTTGAGTTTTACCACTTGCTGAATTATACACGCCCCAAGGAGGGCGACAACCGTCCATTCTTCTGGATGTTTGAGAATGTGGTAGCCATGAAGGTCAACGACAAGAAAGACATCTCCCGATTCTTGGCGTGTAACCCAGTGATGATCGATGCCATCAAGGTTTCTGCCGCTCACAGGGCCCGATACTTCTGGGGCAACCTCCCTGGGATGAACAGG

>PraVole3B

ATGAAGGGAGACAGCAGACATCTGAATGAAGAAGAGGGTGCCAGTGGGGGTGAGGAAAGTGTCATTGTCAATGGGAACTGTAGTGACCATTCCTCAGACACTAAGGATGCTCCCTCACCCCCAGTCTTGGAGGCAATCTGCACACTGGAGAGCAGAGGCCGCAGATCAAGATCAAGGCTGTCAAAGAGGGAGGTCTCCGGCCTGCTGAGTTACACTCAGGACGGAAATGGAGATGAGGCAGAGGATGGGGATGGCGATGATGTTCTAATGCCAAAGCTCACACGTGAGACCAGGTCACCCTCGGAAAACCCAGCTATGCGAACCCGAAATAACAACAGTACCTCCAGGCTAGGGAGGCAAAGAGCCTCCCCCAGAATCACCCGAGGCCGCCAGGGCCGACACCATGTGCAGGAATACCCCGTGGAGTTCCCAGCTACCAGGTCTCGGAGGCGTCGGGCATCATCGTCAGCAAGCACGCCATGGTCGTCCCCTGCCAGCTTCATGGAAGAAGGGACACCTCAGAACAGCAGTATCCCATCAATTGACTTGACCCAGGACATCAATCAGGAGAGCATGGACACTACACAGCTGGAGGCAGAAGGCAAAGATGGAGACAGCACTGAGTATCAGGATGACAAGGAGTTTGGAATAGGTGACCTTGTGTGGGGAAAAATCAAGGGATTCTCCTGGTGGCCTGCCATGGTGGTATCCTGGAAAGCCACCTCTAAGCGCCAGGCCATGCCCGGCATGCGATGGGTACAGTGGTTTGGTGACGGCAAGTTTTCAGAGGTCTCTGCTGACAAACTCATGGCCCTGGGGCTATTCAGCCAGCACTTTAACCTGACTACCTTTAATAAGCTGGTTTCCTATAGGAAAGCCATATACCACACTCTGGAGAGAGCCAGGGTGCGAGCTGGCAAGACCTTCCCCAGCAGCCCGGGAGATTCCCTGGAGGACCAGCTGAAGCCCATGCTGGAATGGGCCCATGGGGGCTTCAAGCCCACTGGCATCGAGGGCCTCAAACCCAACAACAAGCAACCAGAGAACAAAAGTCGAAGACGCACAGACTCTGCCTCTGCTGAGTACCCCCCACCCAAGCGCCTCAAGACCAATAGCTATGGCGGGAAAGACCGAGGGGAGGATGATGAGAGCCGAGAACGGATGGCTTCTGATGTTGCCAATAACAAAGTCAGTCTAGAAGACCGCTGTTTGTCCTGTGGTAGGAAGAACCCTGTGTCCTTCCACCCCCTCTTTGAGGGTGGGCTCTGTCAGAGTTGCCGGGACCGCTTCCTGGAGCTCTTCTACATGTACGATGAGGACGGCTATCAGTCCTACTGCACCGTGTGCTGCGAGGGCCGCGAGCTGCTTCTGTGCAGCAACACAAGCTGCTGCAGGTGCTTCTGTGTGGAGTGTCTGGAGGTGCTGGTGGGTACAGGGACAGCTGAGGATGCCAAGCTGCAGGAACCCTGGAGCTGCTATATGTGCCTCCCACAGCGCTGCCATGGGGTCCTCAGGCGCAGGAAGGATTGGAACACGCGCCTGCAAGACTTCTTCACTACTGACCCTGATCTCGAGGAATTTTACTTGGTGCTCAAAGACTTGGGTATCAAAGTGGAAAAGTACGTTGCCTCCGAAGTCTGTGCCGAGTCCATCGCCGTGGGAACTGTTAAGCATGAAGGCCAAATCAAATATGTGAACGACGTCAGGAAAATCACAAAGAAAAACATTGAAGAGTGGGGCCCTTTTGACTTGGTGATTGGTGGAAGCCCTTGCAATGACCTCTCCAATGTCAATCCTGCCAGGAAAGGTCTATATGAGGGTACCGGCCGGCTCTTCTTTGAGTTTTACCACTTGCTGAATTATACACGCCCCAAGGAGGGCGACAACCGTCCGTTCTTCTGGATGTTTGAGAATGTGGTGGCCATGAAGGTTAATGACAAGAAAGACATCTCCAGATTCTTGGCGTGTAACCCAGTGATGATCGATGCCATCAAGGTTTCTGCCGCTCACAGGGCCCGATACTTCTGGGGCAACCTGCCCGGGATGAACAGG

>CHamster3B

ATGAAGGGAGATAGCAGACATCTGAATGAGGAGGAGGGTGCCAGCGGGTGTGAGGAATGTCTCATCGTCAATGGGAACTGTAGTGACCAGGCCTCAGATACTAAGGATGCTCCTTCACCCCCAGTCTTGGAGGCAATGTGCACATCAGAGAGCAGAGGCCGAAGATCAAGCTCACGGCTGTCAAAGAGGGAGGTCTCCAACCTGCTGAGTTACACTCAGGACGGAGATGGAGATGAAGCAGAGGATGGGGATGGCTCAGACATTCTAATGCCAAAGCTCACGCGTGAGACCAGGTCACCCTCGGAAAGTCCAGCTGTTCGAACCCGAAACAGCAACATTACCTCCAGCCTGGAGAGGCAAAGAGCCTCGCCCAGAATCACCAGAGGCCGCCAGGGCCGCCACCATGTGCAGGAATACCCCGTGGAATTCCCAGCTACCAGGTCTCGGAGAAGGCGAGCATCGTCTTCTACAAGCACACCATGGTCGTCCCCTGCCAGCCTCATAGAAGAAGGGACACCTCAGAGGAGCAGTACCCCATCAACTGACTGGAGCCAGGACAGCCAGCAAGAGAGTATGGATGCCACACAACTGTATGCAGAGAGCAAAGATGGAGACAGCACTGAGTACCAGGATGATAAGGAGTTTGGAATAGGCGACCTTGTATGGGGAAAAATCAAGGGCTTCTCCTGGTGGCCTGCCATGGTGGTTTCCTGGAAAGCCACCTCCAAGCGCCAGGCCATGCCTGGCATGCGATGGGTACAGTGGTTTGGTGACGGCAAGTTTTCTGAGGTCTCTGCTGACAAACTCGTGGCCCTTGGGCTGTTCAGCCAGCACTTTAACCTGGCCACCTTCAATAAGCTGGTTTCTTATAGGAAGGCCATGTACCACACTCTGGAGAGAGCCAGGGTGCGAGCTGGCAAGACCTTCCCCAGCAGCCCTGGAGACTCACTGGAGGACAAGCTGAAGCCCATGCTGGAGTGGGCCCACGGGGGCTTCAAGCCCACTGGCATCGAGGGCCTCAAACCGAATAACAAGCAACCAGAGAACAAAAGTCGAAGAGGCACAGACTCTGCAGCTTCTGAGTCCCCCCCTCCCAAGCGCCTCAAGACCAACAGCTACGGTGGGAAAGACCGAGGGCAGGATGATGAGAGCCGAGAACGGATGGCTTCTGATGTCACCAACAACAAGGGCAATCTGGAAGAACGCTGCTTGTCCTGTGGTAGGAAGAACCCTGTATCCTTCCATCCCCTCTTTGAGGGTGGGCTCTGTCAGAGTTGCCGGGATCGCTTCCTGGAGCTGTTCTACATGTACGATGAGGACGGCTATCAGTCCTACTGCACTGTGTGCTGCGAGGGCCGAGAGCTGCTTCTGTGCAGCAACACAAGCTGCTGCAGGTGCTTCTGTGTGGAGTGTCTGGAGGTGCTGGTGGGTACAGGGACAGCTGAGGATGCCAAGCTGCAGGAACCCTGGAGCTGCTATATGTGCCTTCCACAGCGCTGCCATGGGGTCCTCAGGCGCAGGAAGGATTGGAACATGCGCCTGCAAGACTTCTTCACTTCTGATCCTGACCTCGAAGAATTCTACTTGGTGCTCAAAGAGCTGGGTATCAAAGTGGAAAAGTATGTTGCGTCCGAAGTGTGTACAGAGTCCATTGCTGTGGGAACTGTTAAGCATGAAGGTCAAATCAAATATGTGAATGACGTCAGGAAAATCACAAAGAAAAATATTGAAGAGTGGGGCCCTTTTGACTTGGTGATTGGTGGAAGCCCTTGCAATGACCTCTCCAATGTCAATCCAGCCAGGAAAGGCCTATATGAGGGCACCGGCAGGCTCTTCTTTGAGTTTTACCACTTGCTGAATTATACCCGCCCCAAGGAGGGCGACAACCGTCCGTTCTTCTGGATGTTTGAGAATGTGGTGGCCATGAAGGTTAACGACAAGAAAGACATCTCCCGATTCCTGGCGTGTAACCCAGTGATGATCGATGCCATCAAGGTTTCTGCTGCACATAGGGCCCGATATTTCTGGGGCAACCTACCTGGGATGAACAGG

>BMountMole3B

ATGAAGGGAGACAGCAGACACCTCAATGAGGAAGAAGGTGCCAGTGGGTGTGAGGACTCCATCATTGTCAACGGAACCTGCAGTGACCAGTCCTCAGACACCAAGGATGCTCCCTCACCCCCAGTCTTGGAGGCAATCTGCACACCTGAGAGCAGAGGCCGCAGATCAAGCTCACGATTGTCCAAGAGGGAGGTCTCCAGCCTGCTAAGTTACACTCAGGACGGAGATGGAGATGAAGTAGAGGATGGGGATGGCACTGAAATTCCAATGCCAAAACTCACTCGGGAGACCAGGATACCCTCTGAAAGCCCAGCTGTGCGAACCCGAAATAGCAATAGTACCTCCAGCCGGGAGAGGCGCAGAGCCTCCCGGAGAGTCACCCGGGGCCGGCGGGGCCACCACCAAGTGCACGATTCCCCTGTGGAGTTTCCAGCTACCAGGTCCATGAGGCGGAGGGCAAGAACATCAGCAGGCACGCCGTGGCCATCCCCATCCAGCCTCACAGAAGAAGTGACACCTGGGAGCAGCAGGACCCCCTCCGTTGACCTAGGCCAGGACAGCCAGCAGGAGAGCATGGACTCCACACAGGTGGATACAGAAAGCAGGGATGGAGACAGCACTGAGTATCAGGACGGAAAGGAGTTTGGAATAGGCGACCTTGTGTGGGGAAAGATCAAGGGCTTCTCCTGGTGGCCTGCCATGGTGGTTTCCTGGAGGGCCACCTCTAAGCGCCAGGCCATGCCAGGCATGAGATGGGTACAGTGGTTTGGTGATGGCAAGTTCTCTGAGGTCTCTGCAGACAAACTGGTGGCCTTGGGGCTGTTCAGTCAGCACTTTAATCTGGCCACCTTCAATAAACTGGTTTCTTACAGGAAGGCTATGTACCACACTCTGGAGAAAGCCAGGGTTCGAGCTGGCAAGACCTTCCCCACCAGCCCTGCAGACTCACTGGAGGACCAGCTGAAGCCCATGTTGGAGTGGGCCCACGGAGGCTTCAAGCCCACTGGGATTGAGGGCCTCAAACCCAACAACAAGCAACCAGAGAGCAAAAGTCGAAGACGCACAGACTCTGCAACTTCTGACTATCCCCTACCCAAGCGCCTCAAGACAAACAGCTGTGGTGGCAAGGACCGTGGGGAGGATGATCAGAGCCGAGAGCAAATGGCTTCCGATGTTACCAACAACAAGAGCAACCTGGAAGACAGCTGTTTGTCCTGTGGGAGGAAAAACCCTGTGTCCTTCCACCCACTCTTTGAGGGTGGGCTCTGCCAGACATGCCGGGATCGCTTCCTCGAGCTGTTTTACATGTACGATGATGATGGCTATCAGTCATACTGCACCGTGTGCTGTGAGGGCCGTGAGCTACTTCTGTGCAGCAACACAAGCTGCTGCCGGTGCTTCTGTGTTGAGTGTCTGGAGGTGCTGGTGGGCACAGGCACAGCTGAAGAGGCCAAAATGCAGGAACCCTGGAGCTGCTACATGTGTCTCCCTCAGCGCTGCCATGGTGTCCTCCGACGCAGAAAGGACTGGAACACACGCCTGCAGGATTTCTTTACCACTGTTCCCGATCTGGAGGAATTTTACTTGGTCCTCAAAGAATTGGGTATTAAAGTGGACAGGTACGTCGCCTCTGAAGTCTGTGCAGAATCCATCGCTGTGGGAACTGTGAAGCATGAAGGGCAAATCAAATACGTGAATGACGTCAGGAAAATCACAAAGAGAAATATTGAAGAATGGGGCCCCTTCGACTTGGTGATTGGTGGAAGCCCGTGCAATGATCTCTCTAATGTCAATCCTGCCAGGAAAGGACTGTATGAGGGTACTGGCCGACTCTTTTTTGAGTTTTACCACTTGCTGAATTATACCCGTCCCAAAGAGGGTGACAACCGTCCGTTTTTCTGGATGTTTGAGAATGTGGTTGCCATGAAGGTTAATGACAAGAAGGACATCTCACGGTTCCTGGCGTGTAACCCAGTGATGATTGATGCCATCAAGGTTTCTGCGGCTCACAGGGCTCGATACTTCTGGGGCAACCTACCTGGGATGAACAGG
