## Supplementary material for "Dynamic evolution of *de novo* DNA methyltransferases in rodent and primate genomes": Suppl data 5

*SupFile_5: Muroid Dnmt3L sequence alignments*

>Mouse3L

ATGGGTTCCCGGGAGTCTTGCTCTAAGACCACCTTGGACCTGGAGACTTCCGACAGCTCTAGCCCTGATGCTGACAGTCCTCTGGAAGAGCAATGGCTGAAATCCTCCCCAGCCCTGGAGGACAGTGTGGATGTGGTACTGGAAGACTGCAAAGAGCCTCTGTCCCCCTCCTCGCCTCCGACAGGCAGAGAGATGATCAGGTACGAAGTCAAAGTGAACCGACGGAGCATTGAAGACATCTGCCTCTGCTGTGGAACTCTCCAGGTGTACACTCGGCACCCCTTGTTTGAGGGAGGGTTATGTGCCCCATGTAAGGATAAGTTCCTGGAGTCCCTCTTCCTGTATGATGATGATGGACACCAGAGTTACTGCACCATCTGCTGTTCCGGGGGTACCCTGTTCATCTGTGAGAGCCCCGACTGTACCAGATGCTACTGTTTCGAGTGTGTGGACATCCTGGTGGGCCCCGGGACCTCAGAGAGGATCAATGCCATGGCCTGCTGGGTTTGCTTCCTGTGCCTGCCCTTCTCACGGAGTGGACTGCTGCAGAGGCGCAAGAGGTGGCGGCACCAGCTGAAGGCCTTCCATGATCAAGAGGGAGCGGGCCCTATGGAGATATACAAGACAGTGTCTGCATGGAAGAGACAGCCAGTGCGGGTACTGAGCCTTTTTAGAAATATTGATAAAGTACTAAAGAGTTTGGGCTTTTTGGAAAGCGGTTCTGGTTCTGGGGGAGGAACGCTGAAGTACGTGGAAGATGTCACAAATGTCGTGAGGAGAGACGTGGAGAAATGGGGCCCCTTTGACCTGGTGTACGGCTCGACGCAGCCCCTAGGCAGCTCTTGTGATCGCTGTCCCGGCTGGTACATGTTCCAGTTCCACCGGATCCTGCAGTATGCGCTGCCTCGCCAGGAGAGTCAGCGGCCCTTCTTCTGGATATTCATGGACAATCTGCTGCTGACTGAGGATGACCAAGAGACAACTACCCGCTTCCTTCAGACAGAGGCTGTGACCCTCCAGGATGTCCGTGGCAGAGACTACCAGAATGCTATGCGGGTGTGGAGCAACATTCCAGGGCTGAAGAGCAAGCATGCGCCCCTGACCCCAAAGGAAGAAGAGTATCTGCAAGCCCAAGTCAGAAGCAGGAGCAAGCTGGACGCCCCGAAAGTTGACCTCCTGGTGAAGAACTGCCTTCTCCCGCTGAGAGAGTACTTCAAGTATTTTTCT

>Caroli3L

ATGGGTTCCCGGGAGTCTTTCTCTAAGACCACCTTGGACCTGGAGACTTCCGACAGCTCTAGCCCTGACGCTGACAGTCCTCTGGAAGAGCAATGGCTGAAATCCTCCCCAGCCCTGGAGGACAATGTGGATATGGTACTGGAAGACTGCAAAGAGCCTCTGTCCCCCTCCTCACCCCCGACAGGCAGAGAGATGATCAGGTACGAAGTCAAAGTGAACCGACGGAGCATTGAAGACATCTGCCTCTGCTGCGGAACTCTCCAGGTGTACACTCAGCACCCCTTGTTTGAGGGGGGGATATGTGCCCCATGTAAGGATAAGTTCCTGGAGTCCCTCTTCCTGTACGATGATGATGGACACCAGAGTTACTGCACCATCTGCTGTTCCGGGGGTACCCTGTTCATCTGTGAGAGCCCCGACTGTACCAGATGCTACTGTTTCGAGTGTGTGGATATCCTGGTGGGCCCAGGGACCTCAGAGCGGATCAATGCCATGGCCTGCTGGGTTTGCTTCCTGTGCCTGCCCTTCTCACGGAGTGGACTCCTGCAGAGGCGCAAGAGGTGGCGGCACCAGCTGAAGGCCTTCCATGATCAAGAGGGAGCAGGCCCTATGGAGATATACAAGACAGTGTCCACATGGAAGAGACAGCCAGTGCGGGTACTGAGCCTTTTTGGGAATATTGATAAAGTACTAAAGAGTTTGGGCTTTTTGGAAAGCGGTTCTGGTTCTGGGGGAGGAACGCTGAAGTACGTGGAAGATGTCACAAATGTCGTGAGGAGAGACGTGGAGAAATGGGGCCCCTTTGACCTGGTGTACGGCTCGACGCAGCCCCTAGGCAGCTCTTGTGATCGCTGTCCTGGCTGGTACATGTTCCAATTCCACCGGATCCTGCAGTATGCACTGCCTCGCCAGGAGAGTCAGCGGCCCTTCTTCTGGATATTTATGGACAATCTACTGATGACTGAGGATGACCAAGAGACAACTGCCCGCTTCCTTCAGACAGAGGCTGTGACCCTCCAGGATGTCCGTGGTAGAGACTACCAGAATGTTATGCGGGTGTGGAGCAACATTCCAGGGCTGAAGAGCAAGCATGTGCCCTTGACCCCAAAGGAAGAAGAGTATCTGCAAGCCCAAGTCAGGACCAGAAGCAAGCTGGACGCCCAGAAAGTTGACCTCCTGGTGAAGAACTGCCTTCTCCCCCTGAGAGAGTACTTCAAGTATTTTTCT

>Pahari3L

ATGGGTTCCCGGGAGTCTTGCTCTAAGACCACCGTGAACCTGGAGACTTCCGACAGCTCTAATCCCGACGCTGACAGTCCTCTGGAAGAGCAATGGCTGAAATCCTCCCCAGACCTGGAGGACAGTGTGGATGTGGTACTGGAAGACTGCGAGGAGCCTCTGTCCCCCTCCTCACCCCCCGAAGGCAGAGAGATCATCAGGTACGAAGTCAAAGTGAACCAACGGAACATTGAAGACATCTGCCTCGGCTGCGGAACTCTCCAGGTGTACACCCAGCACCCCTTGTTTGAGGGGGGGATATGTGCCCTGTGTAAGGATAAGTTCCTGGAGTCCCTCTTCCTGTATGATGATGATGGACACCAGAGTTACTGCACCATCTGCTGTTCCGGGGGTACCCTGTTCATCTGTGAGAGCCCTGACTGTACCAGATGCTACTGTTTCGAGTGTGTGGACATTCTGGTGGCCCCCGGAACCTCAGAGCGGATCAATGCCATGGCCTGCTGGGTTTGCTTCCTGTGCCTGCCCTTCTCACGGAGCGGACTGCTGCAGAGGCGCAAGAGGTGGCGGCACCAGCTGAAGGCCTTCCATGATCAAGAGGTAGCAGGCCCTATAGAGATATACAAGACAGTGTCCGCGTGGAAGAGACAGCCAGTGCGGGTACTGAGCCTTTTTGGGAATATTGATAAAGTACTAAAGAGTTTGGGCTTTTTGGAAAGCGGTTCTGGTTCTGAGGGAGGAATGCTGAAGTACGTGGAAGATGTCACAAATGTCGTGAGGAGAGACGTGGAGAAATGGGGCCCCTTTGACCTGGTGTATGGCTCGACGCAGCCCCTAGGCAGCTCTTGTGATCGCTGTCCTGGCTGGTACATGTTCCAATTCCACCGGATCCTGCAGTATGCGCTGCCTCGCCAGGAGAGTCAGCGGCCCTTCTTCTGGATATTTATGGACAATCTGTTGCTGACTGAGGATGACCAGGAGACAACTGTCCGCTTCCTTCAGACAGAGACTGTGACCCTCCAGTATGTCCGTGGCAGAGTCCTCCAGAATGCTGTGCGGGTGTGGAGCAACATTCCTGGGCTGAAGAGCAAGCACGCGGTCCTGACCCCACAGGAAGAAGAGACTCTGCAAGCCCAAATCAGAACCAGAAACAAGCCGGACACCCAGAAAGTTGACCCCCTGGTGAAGAGCTGCCTTCTGCCCCTGAGAGAGTACTTCAAGTATTTTTCT

>Rat3L

ATGGGTTCCCGGGAGTCCTGCTCTAAGACCACCTTGAACCTGGAGACTCCGGAGAGCTCTAGCACTGACCCTGACAGTCCCCTGGAAGAGCAATGGCCGAAATCTTCCCCAGATCTGGAAGACAGCGTGGATATGGTACTGGAAGACTCCAAGGAGCCTCTAACCCCTTCCTCACCGCCGACAGGCAGAGAGGTCATCAGGTACGAAGTCAACGTGAACCAGCGGAACATCGAAGACATCTGCCTCTGTTGCGGATCTCTCCAGGTGTACGCTCAGCACCCCTTGTTTGAGGGGGGAATTTGTGCCCCGTGTAAGGACAAGTTCCTGGAGACCCTCTTCCTATACGACGAGGATGGACACCAGAGCTACTGTACCATCTGCTGCTCCGGGCATACCCTGTTCATCTGCGAGAGCCCCGACTGTACCAGATGCTACTGTTTCGAGTGTGTGGATATCCTGGTGGGCCCCGGGACCTCGGAGCGCATCAATGCCATGGCCTGCTGGGTTTGCTTCCTGTGCCTGCCTTTCTCCCGGAGCGGACTGCTGCAGAGGCGCAAGAAGTGGCGTCACCAGCTGAAGGCCTTCCATGACCGAGAGGGGGCAAGCCCTGTGGAGATATACAAGACTGTGTCTGCATGGAAAAGACAGCCAGTGAGGGTGCTGAGTCTTTTTGGGAATATTGATAAAGAACTAAAGAGTTTGGGCTTTTTGGAAAGCAGTTCTGGTTCTGAGGGAGGAACGCTGAAGTACGTGGAAGATGTCACCAATGTCGTGAGGAGAGAAGTGGAGAAATGGGGTCCCTTTGACTTGGTGTATGGCTCAACCCAGCCCCTAGGCTATTCTTGTGACCGCTGTCCTGGCTGGTACATGTTCCAGTTCCACCGGATCCTGCAGTATGCCCGGCCTCGCCAAGACAGCCAGCAGCCCTTCTTCTGGATATTTGTGGACAATCTGCTGCTGACTGAAGATGACCAAGAGACAACTGTCCGCTTCCTTCAGACAGAGGCTGTGACCCTCCAGGATGTCCGCGGCAGAGTCCTCCAGAATGCTATGAGGGTGTGGAGCAACATTCCAGGGCTGAAGAGCAAGCACGCGGACCTGACCCCTAAGGAAGAGCAGTCTCTGCAAACCCAAGTCAGAACCAGAAGCAAGCTGGCCGCCCAGAAAGTTGACTCCCTAGTGAAGTACTGCCTTCTCCCCCTGAGAGAGTACTTCAAGTATTTTTCT

>DeerMouse3L

ATGGGTTCCCAGGAGTCTTGCCCCAAGGCTACCTGGGACCTGGAGACTCTGGACAGTTCTAGCCCGGACTCTGCCAGCCACCTGGAAGAGCAGTGGCCGAAACCCTCCCCGAGCCTGGAGGATAGCATGGATGTTGTACTGGAAGACTCTGGGGAGATACGGTCCTTAGCCTCACCTCCACCAGGCAGAGAGATCATCAGGTATGAAGTCACTGTGAACCGACGGAACATCGAAGACATCTGCCTCTGTTGTGGAAGTCTCCAGGTGCATACGCAGCACCCCTTATTTGAGGGGGGCATATGTGCCTCGTGTAAGGACACGTTCCTGGAGACCATCTTCCTTTATGATGAGGATGGGCATCAGAGCTACTGTTCCATCTGCTGCTCAGGGCACACCTTGTTCATCTGTGAGAGCCCCGACTGTACCAGATGTTACTGTTTTGAGTGTGTGGATATCCTGGTGGGCCCCGGGACTTCAGAGCGAATCAACGCCATGGCGTGCTGGGTTTGCTTCCTGTGCCTGCCTTTCACTCGGAGCGGGCTGCTGCAGAGGCGAAAGAGGTGGCGTCACCAGCTGAAGGCCTTCCATGATCTAGAGGGGGCAAGCCCTCTGGAGATGTATAAGCCAGTGTCTGCTTGGAAGAGACAGCCAGTGCAGGTACTGAGCCTCTTTGGGAATATTGAGAAAGAGCTAGAGAGTTTGGGCTTTCTGGAAAGAGGTTCTGGTTCTGAAGGAGGAAGGCTGAAGTACTTGGAAGATGTCACGAATGTTGTGAGGAGAGATGTGGAGAGATGGGGCCCCTTTGACCTTGTATATGGCTCAACACAGCCTCTAGGTCGCTCTTGTGACCGCTGCCCTGGCTGGTACATGTTCCAGTTCCACCGGATCCTGCAGTATGCTCGCCCACACCCAGGGAGTCAGCGGCCTTTCTTCTGGCTGTTTGTGGACAGTCTGCTGCTGACTGAGGATGACCAAGTCACCACAACCCGCTTCCTCCAGATGGAGCCTGTGACCCTCCAGAATATTCGCGGCAGAGTCCTCCAGAATGCTGTGCGGGTGTGGAGCAACATCCCAGGGTTGAAGAGCAAACACTTGGCCCTGACACCAAAGGAGCAACAGTCCTTGGAAGGCCAAGTCAGAACCAGAGCCAAGATGGCCTCCCAGAAAGACGACCCCCTGGTGAAGAACTGCCTTCTCCCCCTGAGAGAGTATTTCAAGTATTTTTCT

>PrairieVole3L

ATGGGTTCCCGGAAATCTTGCTCCAAGACAACATGGGACCTGGAGACTCTGGACAGCTCTAGCCCGGACTCTGCCAATCACCTGGAAGAGCAATGGCTGAAATCCACCCGTAGCCTCGAGCACAGCGTGGAGGTCGTACTGGAAGATTCCCGGGAGCTGCTGTCCTTAACCTCACCCCCGCCAGGCAGAGAGATCATCAGGTATGAAGTCACTGTGAACCAACGGAACATCGAAGACATCTGCCTCTGTTGTGGAAGTCTCCAGGTGTATACGCAGCACCCCTTATTTGAGGGGGGGATGTGTGCCCCATGTAAGGACATATTCCTGGAGACCATCTTCCTTTACGATATGGATGGGCACCAGAGTTACTGCTCCATATGCTGCTTCGGGAAAACCCTGTTTATCTGTGAGAGCCCCGACTGTACCAGATGCTACTGCTTTGAGTGTGTGGATATCCTGGTGGGTCCCGGGACCTCAGAGCGAATCAGCACCATGGCCTGCTGGGTTTGCTTCCTGTGCCTGCCTTTCACTCGGAGCGGACTGCTGCAGAGGCGAAGGAAGTGGCGTCACCAGCTGAAGGCCTTCCACGATCTCGAGGGGTCAAATCCTGTGAAGATGTACAAGATAGTGTCTGCCTGGAAGAGACAGCCCGTGAGGGTGCTTAACCTTTTTGGGAAGATTGATAAAGAGCTCGAGAGTTTGGGCTTTCTGGAAAGTGGTTCTGGTTCGGAAGAGGGAAGACTGAAGTACTTGGAAGACGTCACCAATGTTGTGAGAAGAGATGTGGAGAGATGGGGCCCCTTTGACCTCGTGTATGGCTCGACACAGCCCCTAGGCTACTCTTGTAACCGCTGCCCTGGCTGGTACATGTTCCAATTCCACCGGATCCTGCAGTATGCTCAACCACCCTCAGGGAGTCAGCAGCCTTTCTTCTGGCTCTTTATGGACAATCTACTGCTGACAGAGGATGACCAGGCCACGGCAACCCGCTTCTTCCAGGTGGAGGCTGTGACCCTCCAGGATGTCCGCAGCAGAGTCCTCCAGAATGCTGTGCGGGTGTGGAGCAACATCCCAGGGTTGAAGAGCAAACACTCGGCCCTAACACCTAAGGAGCTGCAGTCCCTGGAAACCCAGACCAGAACCAGAGGCAAGATGGCTGCCCAGAAAGTTGACCTCCTGGTGAAGACCTGCCTTCTCCCCCTGAGAGAGTATTTCAAGTATTTTTCT

>Gebril3L

ATGGGTTCCCAGGAGACCCGCGCCAAGACCACCTGGAACCTGGAGAGTACCGACAGCTCTAGCCCCGAATCTCTCGGCCACCTGGAAGAGCAATGGGCCAACTCCTCCCCGGACCTGGAGCATAGCAAGGACGTGGAGCCGGAGGACTCCAAGGAGCTGATTTCCTCAGCTTCACCCCCATCAGGCAGAGAAATCATCAGGTATGAAATCTCTGTGAACCAACGTAACATCGAAGACATCTGCCTCTGTTGTGGAACTCTCCAGGTGTATAAGCAGCACCCCTTGTTTGAGGGTGGAATATGCGCCCCATGTAAGGACAAGTTCCTGGAAACCTTCTTCCTTTACGATGAGGATGGGCACCAGAGTTACTGCTCCATCTGCTGCTCGGGGGGTACCCTGTTCATCTGTGAGAGTCCAGACTGTACCAGATGCTACTGTTTTGAGTGTGTGGATATCCTGGTGGGCCCGGGGACCTCAGAGCGGATCAATGCCATGCCCTGCTGGGTCTGCTTCCTGTGCCTGCCTTTCACCCGGAGTGGACTGCTGCAGAGGCGGAGGAAGTGGCGCCACCAGCTGAAGGCCTTCTTTGATGAAGGGGGGGCAAGCCCTCTGGAGATGTACAAGACAGTGTCTGCGTGGAAGAGAAAACCAATGAGGGTGCTGAGCCTTTTTAAGAATATTGATAAAGAGCTAAAGAATTTAGGCTTTTTGGAAAGTGGTTCTGGTTCTGAAGAAGAAAGACTGAAGTACTTGGAAGATGTCACAAATGTCGTGAGAAGAGATGTGGAGAAATGGGGTCCCTTTGACCTCGTGTACGGCTCGACACGGCCCCGAGGCTCCTCTTGTGACCACTGCCCTGCCTGGTACATGTTCCAGTTCCACCGGATCCTGCAGTATGCGCGGCCACCCTCGGGGAGTGAGCAGCCCTTCTTCTGGGTGTTTGTGGACAATCTGCTGATGACCGAGGACGACCAAATCACAGCAGACCGCTTCCTTCAGATGAAAGCTGTGACCCTCCAGGACGTCCGAGGCAGAGTCCTTCAGAATGCTGTGCGGGTGTGGAGCAACATCCCAGGGGTGAAAAGCAAGCACATGGCCCTGACAGAAAAGGAGGAGCAATCTCTGGAAGCTCAGGCTGGAACCAGAACCAAGCTATCAGCCCAGAAAGTTGACCCTCTGGTGAAAAACTGCCTTCTTCCCCTGAGAGAGTACTTCAAGTTTTTTTCT

>ChHams3L

ATGGGTTCCCGGGAGTCTTGCTCCAAGGCCACTTGGGACCTGGAGACGCTGGACAGCTCTAGCCGGGACTCTGCCAGCCACCTGGAAGAGCAATGGCGGAAATTCTCCCCCGTCCTCGAGGACAGCACGGATATTGTACTGGAAGACTCCCGGGAGCTGCTGTCCTTAACTTCACCCCCGCCAGGCAGAGAGATCATCAGGTATGAAGTCAGTGTGAACCGACGGAACATCGAAGACATCTGCCTAGGTTGTGGAAGTCTCCAGGTGTACACGCAGCACCCCTTATTTGAGGGGGGGATATGTGCCTCGTGTAAGGACAATTTCCTGGAAACCCTCTTCCTTTATGATGAGGATGGGCACCAGAGCTACTGTACCATCTGCTGCTCTGGGGACACCCTGTTCATCTGTGAGAGCCCCGACTGTACCAGATGCTACTGTTTCGAGTGTGTGGATATCCTGGTAGGCCCTGGGACCTCAGAACGAATCAATGCCATGGCCTGCTGGGTTTGCTTCCTGTGCCTGCCTTTCGCTCGTAGCGGACTGCTGCAGAGGCGAAGACAGTGGCGTCACCAGCTGAAGGCCTTCTATGATCTAGAAGGGGCAAGCCCTCTGGAGATGTACAAGACAGTGTCTGCGTGGAAGAGACAGCCCATGCGGGTGCTGAGCCTTTTTGGGAATATTGATAAAGAGCTAAAGAGTTTGGGCTTTTTGGAAAGTGGCTCTCGCACTGAGGAAGGAAGACTGAAGTACTTGGATGATGTCACAAATGTTGTGAGGAGAGACGTGGAGAGATGGGGTCCCTTTGACCTTGTGTATGGCTCAACACAGCCCCTAGGCTACTCTTGTGACCACTGCCCTGGATGGTACATGTTCCAATTTCACCGGATACTGCAATACGCCCGGCCTCACCCAGGGAGTCAGCAGCCTTTCTTCTGGATATTTGTGGACAATCTGCTGCTGTCTGAGGATGACCAAGTCACAGCAGCCCGCTTCTTTCAGACGGAGGCTGTGACCCTCCAGGATGTCCGCGGCAGAGTCCTCCAGAATGCTGTGCGGGTATGGAGCAACATCCCAGGCTTGAAGAGCAAACACTCGGCCCTGACACCAAAGGAGGAGCAGTCCCTGGAAGGACACGTTAGAACCAGAGCCAAGGTGGCCGCCCAGAAAACTGACGCCCTGGTGAAGAACTGCCTTCTCCCTCTGAGAGAGTATTTCAAGTATTTTTCT

>GalilMole3L

ATGGGTTCCTGGGAGTCTTGTTCCAAGGCCACATGGAACTTGGAGACTCCAGACAACTCGAGCCCTGACCCAGCCGGCCACCTGGAAGAACAGAAGCC---CTCACCTCCACCTCTGGAGAACAACATGGATGTAGTACTGGTGGACTCCAGGGAGCTGCTGTGCCCCACTTCACCCCCTCCAGGTAGAGATCGCATCGCCTATGAAGTGACCGTCAACCACCGGAACATAGAAGAGCTCTGCCTGTGCTGTGGAAGTCTCCAGGTGTACACACAGCACCCCCTGTTCCAGGGAGGAATGTGTGCCCCATGTAAGGACAAATTCTTGGAGACCCTCTTCCTGTATGACGATGACGGGTATCAGTGTTCCTGCTCCATCTGCTGCTCGGGGGACACGCTGTTCATCTGCGAGAGTCCAGACTGTACCCGGTGCTACTGTTTTGAGTGTGTGGATGTCCTGGTGGGCCCTGGGACCTCAGAGCGAATCCACGCCATGAACTGCTGGGTGTGCTTCCTGTGCCTGCCTTTCTCTCAAAGCGGGCTGCTCCTGAGGCGGAGGAAGTGGCGCCACCGGCTGAAGGCCTTCTATGACCTCGAGGCGGCAAGCCCTCTGGAAATGTACAAAACAGTGCCTGTGTGGAAGAGAGAGGCAGTGCGGGTGCTGACCCTTTTTGAAGACATTCAGAAAGAACTAAGGAGCTTAGGCTTTTTGGAAAGTGGTTCCGGTTCTGAGAGAGGAAGACTGAAACACTTGGATGATGTCACAAACGTAGTGCGAAGAGATGTGGAGGGATGGGGCCCCTTTGACCTTGTGTATGGCTCAACACAGCCCCTGGGCTATTCCTGCAACCGTTGTCCTGGCTGGTATCTGTTCCAGTTTCACCGGATCCTGCAGTATGCACGGTCACACCGGGGCAGCCAGCAACCCTTTTTCTGGATGTTTGTGGACAACTTGCTGCTGACTGAGGCTGACCAGGCCACAGCAGCCCGCTTCCTTGAGATGGAGCCCATGACCCTCCAGGACATCCGAGGCAGAGTCCTGCAGAACGCGGTTCACGTGTGGAGCAACATCCCAGGGGTGAAGAGCAAGCACATGGCCCTGACAGCCGAGGAGGAACAGTTTCTGCAGGCCCAAGGTGATGCCCGAGCCAAGCTGGTTGCCAAGGGACTGGCCCCGCTGGTGAAGAAGTGCTTTCTCCCCCTTAGAGAGTATTTCAAGTATTTTTCT
