## Supplementary material for "Dynamic evolution of *de novo* DNA methyltransferases in rodent and primate genomes": Suppl data 6

*SupFile_6: Primate* DNMT3A *sequence alignments*

*>Human_Dnmt3A*

*ATGCCCTCCAGCGGCCCCGGGGACACCAGCAGCTCTGCTGCGGAGCGGGAGGAGGACCGAAAGGACGGAGAGGAGCAGGAGGAGCCGCGTGGCAAGGAGGAGCGCCAAGAGCCCAGCACCACGGCACGGAAGGTGGGGCGGCCTGGGAGGAAGCGCAAGCACCCCCCGGTGGAAAGCGGTGACACGCCAAAGGACCCTGCGGTGATCTCCAAGTCCCCATCCATGGCCCAGGACTCAGGCGCCTCAGAGCTATTACCCAATGGGGACTTGGAGAAGCGGAGTGAGCCCCAGCCAGAGGAGGGGAGCCCTGCTGGGGGGCAGAAGGGCGGGGCCCCAGCAGAGGGAGAGGGTGCAGCTGAGACCCTGCCTGAAGCCTCAAGAGCAGTGGAAAATGGCTGCTGCACCCCCAAGGAGGGCCGAGGAGCCCCTGCAGAAGCGGGCAAAGAACAGAAGGAGACCAACATCGAATCCATGAAAATGGAGGGCTCCCGGGGCCGGCTGCGGGGTGGCTTGGGCTGGGAGTCCAGCCTCCGTCAGCGGCCCATGCCGAGGCTCACCTTCCAGGCGGGGGACCCCTACTACATCAGCAAGCGCAAGCGGGACGAGTGGCTGGCACGCTGGAAAAGGGAGGCTGAGAAGAAAGCCAAGGTCATTGCAGGAATGAATGCTGTGGAAGAAAACCAGGGGCCCGGGGAGTCTCAGAAGGTGGAGGAGGCCAGCCCTCCTGCTGTGCAGCAGCCCACTGACCCCGCATCCCCCACTGTGGCTACCACGCCTGAGCCCGTGGGGTCCGATGCTGGGGACAAGAATGCCACCAAAGCAGGCGATGACGAGCCAGAGTACGAGGACGGCCGGGGCTTTGGCATTGGGGAGCTGGTGTGGGGGAAACTGCGGGGCTTCTCCTGGTGGCCAGGCCGCATTGTGTCTTGGTGGATGACGGGCCGGAGCCGAGCAGCTGAAGGCACCCGCTGGGTCATGTGGTTCGGAGACGGCAAATTCTCAGTGGTGTGTGTTGAGAAGCTGATGCCGCTGAGCTCGTTTTGCAGTGCGTTCCACCAGGCCACGTACAACAAGCAGCCCATGTACCGCAAAGCCATCTACGAGGTCCTGCAGGTGGCCAGCAGCCGCGCGGGGAAGCTGTTCCCGGTGTGCCACGACAGCGATGAGAGTGACACTGCCAAGGCCGTGGAGGTGCAGAACAAGCCCATGATTGAATGGGCCCTGGGGGGCTTCCAGCCTTCTGGCCCTAAGGGCCTGGAGCCACCAGAAGAAGAGAAGAATCCCTACAAAGAAGTGTACACGGACATGTGGGTGGAACCTGAGGCAGCTGCCTACGCACCACCTCCACCAGCCAAAAAGCCCCGGAAGAGCACAGCGGAGAAGCCCAAGGTCAAGGAGATTATTGATGAGCGCACAAGAGAGCGGCTGGTGTACGAGGTGCGGCAGAAGTGCCGGAACATTGAGGACATCTGCATCTCCTGTGGGAGCCTCAATGTTACCCTGGAACACCCCCTCTTCGTTGGAGGAATGTGCCAAAACTGCAAGAACTGCTTTCTGGAGTGTGCGTACCAGTACGACGACGACGGCTACCAGTCCTACTGCACCATCTGCTGTGGGGGCCGTGAGGTGCTCATGTGCGGAAACAACAACTGCTGCAGGTGCTTTTGCGTGGAGTGTGTGGACCTCTTGGTGGGGCCGGGGGCTGCCCAGGCAGCCATTAAGGAAGACCCCTGGAACTGCTACATGTGCGGGCACAAGGGTACCTACGGGCTGCTGCGGCGGCGAGAGGACTGGCCCTCCCGGCTCCAGATGTTCTTCGCTAATAACCACGACCAGGAATTTGACCCTCCAAAGGTTTACCCACCTGTCCCAGCTGAGAAGAGGAAGCCCATCCGGGTGCTGTCTCTCTTTGATGGAATCGCTACAGGGCTCCTGGTGCTGAAGGACTTGGGCATTCAGGTGGACCGCTACATTGCCTCGGAGGTGTGTGAGGACTCCATCACGGTGGGCATGGTGCGGCACCAGGGGAAGATCATGTACGTCGGGGACGTCCGCAGCGTCACACAGAAGCATATCCAGGAGTGGGGCCCATTCGATCTGGTGATTGGGGGCAGTCCCTGCAATGACCTCTCCATCGTCAACCCTGCTCGCAAGGGCCTCTACGAGGGCACTGGCCGGCTCTTCTTTGAGTTCTACCGCCTCCTGCATGATGCGCGGCCCAAGGAGGGAGATGATCGCCCCTTCTTCTGGCTCTTTGAGAATGTGGTGGCCATGGGCGTTAGTGACAAGAGGGACATCTCGCGATTTCTCGAGTCCAACCCTGTGATGATTGATGCCAAAGAAGTGTCAGCTGCACACAGGGCCCGCTACTTCTGGGGTAACCTTCCCGGTATGAACAGGCCGTTGGCATCCACTGTGAATGATAAGCTGGAGCTGCAGGAGTGTCTGGAGCATGGCAGGATAGCCAAGTTCAGCAAAGTGAGGACCATTACTACGAGGTCAAACTCCATAAAGCAGGGCAAAGACCAGCATTTTCCTGTCTTCATGAATGAGAAAGAGGACATCTTATGGTGCACTGAAATGGAAAGGGTATTTGGTTTCCCAGTCCACTATACTGACGTCTCCAACATGAGCCGCTTGGCGAGGCAGAGACTGCTGGGCCGGTCATGGAGCGTGCCAGTCATCCGCCACCTCTTCGCTCCGCTGAAGGAGTATTTTGCGTGTGTGTAA*

*>Gorilla3A*

*ATGCCCTCCAGCGGCCCCGGGGACACCAGCAGCTCTGCTGCGGAGCGGGAGGAGGACCGAAAGGACGGAGAGGAGCAGGAGGAGCCGCGTGGCAAGGAGGAGCGCCAAGAGCCCAGCACCACGGCACGGAAGGTGGGGCGGCCTGGGAGGAAGCGCAAGCACCCCCCGGTGGAAAGCGGTGACACGCCAAAGGACCCTGCGGTGATCTCCAAGTCCCCATCCATGGCCCAGGACTCAGGCGCCTCAGAGCTATTACCCAATGGGGACTTGGAGAAGCGGAGTGAGCCCCAGCCAGAGGAGGGGAGCCCTGCTGGGGGGCAGAAGGGCGGGGCCCCAGCAGAGGGAGAGGGTGCAGCTGAGACCCTGCCTGAAGCCTCAAGAGCAGTGGAAAATGGCTGCTGCACCCCCAAGGAGGGCCGAGGAGCCCCTGCAGAAGCGGGCAAAGAACAGAAGGAGACCAACATCGAATCCATGAAAATGGAGGGCTCCCGGGGCCGGCTGCGGGGTGGCTTGGGCTGGGAGTCCAGCCTCCGTCAGCGGCCCATGCCGAGGCTCACCTTCCAGGCAGGGGACCCCTACTACATCAGCAAGCGCAAGCGGGACGAGTGGCTGGCACGCTGGAAAAGGGAGGCTGAGAAGAAAGCCAAGGTCATTGCAGGAATGAATGCTGTGGAAGAAAACCAGGGGCCCGGGGAGTCTCAGAAGGTGGAGGAGGCCAGCCCTCCTGCTGTGCAGCAGCCCACTGACCCCGCATCCCCCACTGTGGCTACCACGCCTGAGCCCGTGGGGTCCGATGCTGGGGACAAGAATGCCACCAAAGCAGGCGATGACGAGCCAGAGTACGAGGACGGCCGGGGCTTTGGCATTGGGGAGCTGGTGTGGGGGAAACTGCGGGGCTTCTCCTGGTGGCCAGGCCGCATTGTGTCTTGGTGGATGACGGGCCGGAGCCGAGCAGCTGAAGGCACCCGCTGGGTCATGTGGTTCGGAGACGGCAAATTCTCAGTGGTGTGTGTTGAGAAGCTGATGCCGCTGAGCTCGTTTTGCAGTGCGTTCCACCAGGCCACGTACAACAAGCAGCCCATGTACCGCAAAGCCATCTACGAGGTCCTGCAGGTGGCCAGCAGCCGCGCGGGGAAGCTGTTCCCGGTGTGCCACGACAGCGATGAGAGTGACACTGCCAAGGCCGTGGAGGTGCAGAACAAGCCCATGATTGAATGGGCCCTGGGGGGCTTCCAGCCCTCTGGCCCTAAGGGCCTAGAGCCACCAGAAGAAGAGAAGAATCCCTACAAAGAAGTGTACACGGACATGTGGGTGGAACCTGAGGCAGCTGCCTACGCACCACCTCCACCAGCCAAAAAGCCCCGGAAGAGCACAGCAGAGAAGCCCAAGGTCAAGGAGATTATTGATGAGCGCACAAGAGAGCGGCTGGTGTACGAGGTGCGGCAGAAGTGCCGGAACATTGAGGACATCTGCATCTCCTGTGGGAGCCTCAATGTTACCCTGGAACACCCCCTCTTCGTTGGAGGAATGTGCCAAAACTGCAAGAACTGCTTTCTGGAGTGTGCGTACCAGTACGACGACGACGGCTACCAGTCCTACTGCACCATCTGCTGTGGGGGCCGTGAGGTGCTCATGTGCGGAAACAACAACTGCTGCAGGTGCTTTTGCGTGGAGTGTGTGGACCTCTTGGTGGGGCCGGGGGCTGCCCAGGCAGCCATTAAGGAAGACCCCTGGAACTGCTACATGTGCGGGCACAAGGGTACCTACGGGCTGCTGCGGCGGCGAGAGGACTGGCCCTCCCGGCTCCAGATGTTCTTCGCTAATAACCACGACCAGGAATTTGACCCTCCAAAGGTTTACCCACCTGTCCCAGCTGAGAAGAGGAAGCCCATCCGGGTGCTGTCTCTCTTTGATGGAATCGCTACAGGGCTCCTGGTGCTGAAGGACTTGGGCATTCAGGTGGACCGCTACATTGCCTCGGAGGTGTGTGAGGACTCCATCACGGTGGGCATGGTGCGGCACCAGGGGAAGATCATGTACGTCGGGGACGTCCGCAGCGTCACACAGAAGCATATCCAGGAGTGGGGCCCATTCGATCTGGTGATTGGGGGCAGTCCCTGCAATGACCTCTCCATCGTCAACCCTGCTCGCAAGGGCCTCTACGAGGGCACTGGCCGGCTCTTCTTTGAGTTCTACCGCCTCCTGCATGATGCGCGGCCCAAGGAGGGAGATGATCGCCCCTTCTTCTGGCTCTTTGAGAATGTGGTGGCCATGGGCGTTAGTGACAAGAGGGACATCTCGCGATTTCTCGAGTCCAACCCTGTGATGATTGATGCCAAAGAAGTGTCAGCTGCACACAGGGCCCGCTACTTCTGGGGTAACCTTCCCGGTATGAACAGGCCGTTGGCATCCACTGTGAATGATAAGCTGGAGCTGCAGGAGTGTCTGGAGCATGGCAGGATAGCCAAGTTCAGCAAAGTGAGGACCATTACTACGAGGTCAAACTCCATAAAGCAGGGCAAAGACCAGCATTTTCCTGTCTTCATGAATGAGAAAGAGGACATCTTATGGTGCACTGAAATGGAAAGGGTATTTGGTTTCCCAGTCCACTATACTGACGTCTCCAACATGAGCCGCTTGGCGAGGCAGAGACTGCTGGGCCGGTCATGGAGCGTGCCAGTCATCCGCCACCTCTTCGCTCCGCTGAAGGAGTATTTTGCGTGTGTGTAA*

*>Chimp3A*

*ATGCCCTCCAGCGGCCCCGGGGACACCAGCAGCTCTGCTGCGGAGCGGGAGGAGGACCGAAAGGACGGAGAGGAGCAGGAGGAGCCGCGTGGCAAGGAGGAGCGCCAAGAGCCCAGCACCACAGCACGGAAGGTGGGGCGGCCTGGGAGGAAGCGCAAGCACCCCCCGGTGGAAAGTGGTGACACGCCAAAGGACCCTGCGGTGATCTCCAAGTCCCCATCCATGGCCCAGGACCCAGGCGCCTCAGAGCTATTACCCAATGGGGACTTGGAGAAGCGGAGTGAGCCCCAGCCAGAGGAGGGGAGCCCTGCTGGGGGGCAGAAGGGCGGGGCCCCAGCAGAGGGAGAGGGTGCAGCTGAGACCCTGCCTGAAGCCTCAAGAGCAGTGGAAAATGGCTGCTGTACCCCCAAGGAGGGCCGAGGAGCCCCTGCAGAAGTGGGCAAAGAACAGAAGGAGACCAACATCGAATCCATGAAAATGGAGGGCTCCCGGGGCCGGCTGCGGGGTGGCTTGGGCTGGGAGTCCAGCCTCCGTCAGCGGCCCATGCCGAGGCTCACCTTCCAGGCGGGGGACCCCTACTACATCAGCAAGCGCAAGCGGGACGAGTGGCTGGCACGCTGGAAAAGGGAGGCTGAGAAGAAAGCCAAGGTCATTGCAGGAATGAATGCTGTGGAAGAAAACCAGGGGCCCGGGGAGTCTCAGAAGGTGGAGGAGGCCAGCCCTCCTGCTGTGCAGCAGCCCACTGACCCCGCATCCCCCACTGTGGCTACCACGCCTGAGCCCGTGGGGTCCGATGCTGGGGACAAGAATGCCACCAAAGCAGGCGATGACGAGCCAGAGTACGAGGACGGCCGGGGCTTTGGCATTGGGGAGCTGGTGTGGGGGAAACTGCGGGGCTTCTCCTGGTGGCCAGGCCGCATTGTGTCTTGGTGGATGACGGGCCGGAGCCGAGCAGCTGAAGGCACCCGCTGGGTCATGTGGTTCGGAGACGGCAAATTCTCAGTGGTGTGTGTTGAGAAGCTGATGCCACTGAGCTCGTTTTGCAGTGCGTTCCACCAGGCCACGTACAACAAGCAGCCCATGTACCGCAAAGCCATCTACGAGGTCCTGCAGGTGGCCAGCAGCCGCGCGGGGAAGCTGTTCCCGGTGTGCCACGACAGCGATGAGAGTGACACTGCCAAGGCCGTGGAGGTGCAGAACAAGCCCATGATTGAATGGGCCCTGGGGGGCTTCCAGCCCTCTGGCCCTAAGGGCCTAGAGCCACCAGAAGAAGAGAAGAATCCCTACAAAGAAGTGTACACGGACATGTGGGTGGAACCTGAGGCAGCTGCCTACGCACCACCTCCACCAGCCAAAAAGCCCCGGAAGAGCACAGCGGAGAAGCCCAAGGTCAAGGAGATTATTGATGAGCGCACAAGAGAGCGGCTGGTGTACGAGGTGCGGCAGAAGTGCCGGAACATTGAGGACATCTGCATCTCCTGTGGGAGCCTCAATGTTACCCTGGAACACCCCCTCTTCGTTGGAGGAATGTGCCAAAACTGCAAGAACTGCTTTCTGGAGTGTGCGTACCAGTACGACGACGACGGCTACCAGTCCTACTGCACCATCTGCTGTGGGGGCCGTGAGGTGCTCATGTGCGGAAACAACAACTGCTGCAGGTGCTTTTGCGTGGAGTGTGTGGACCTCTTGGTGGGGCCGGGGGCTGCCCAGGCAGCCATTAAGGAAGACCCCTGGAACTGCTACATGTGCGGGCACAAGGGTACCTACGGGCTGCTGCGGCGGCGAGAGGACTGGCCCTCCCGGCTCCAGATGTTCTTCGCTAATAACCACGACCAGGAATTTGACCCTCCAAAGGTTTACCCACCTGTCCCAGCTGAGAAGAGGAAGCCCATCCGGGTGCTGTCTCTCTTTGATGGAATCGCTACAGGGCTCCTGGTGCTGAAGGACTTGGGCATTCAGGTGGACCGCTACATTGCCTCGGAGGTGTGTGAGGACTCCATCACGGTGGGCATGGTGCGGCACCAGGGGAAGATCATGTACGTCGGGGACGTCCGCAGCGTCACACAGAAGCATATCCAGGAGTGGGGCCCATTCGATCTGGTGATTGGGGGCAGTCCCTGCAATGACCTCTCCATCGTCAACCCTGCTCGCAAGGGCCTCTACGAGGGCACTGGCCGGCTCTTCTTTGAGTTCTACCGCCTCCTGCATGATGCGCGGCCCAAGGAGGGAGATGATCGCCCCTTCTTCTGGCTCTTTGAGAATGTGGTGGCCATGGGCGTTAGTGACAAGAGGGACATCTCGCGATTTCTCGAGTCCAACCCTGTGATGATTGATGCCAAAGAAGTGTCAGCTGCACACAGGGCCCGCTACTTCTGGGGTAACCTTCCCGGTATGAACAGGCCGTTGGCATCCACTGTGAATGATAAGCTGGAGCTGCAGGAGTGTCTGGAGCATGGCAGGATAGCCAAGTTCAGCAAAGTGAGGACCATTACTACGAGGTCAAACTCCATAAAGCAGGGCAAAGACCAGCATTTTCCTGTCTTCATGAATGAGAAAGAGGACATCTTATGGTGCACTGAAATGGAAAGGGTATTTGGTTTCCCAGTCCACTATACTGACGTCTCCAACATGAGCCGCTTGGCGAGGCAGAGACTGCTGGGCCGGTCATGGAGCGTGCCAGTCATCCGCCACCTCTTCGCTCCGCTGAAGGAGTATTTTGCGTGTGTGTAA*

*>Bonobo3A*

*ATGCCCTCCAGCGGCCCCGGGGACACCAGCAGCTCTGCTGCGGAGCGGGAGGAGGACCGAAAGGACGGAGAGGAGCAGGAGGAGCCGCGTGGCAAGGAGGAGCGCCAAGAGCCCAGCACCACGGCACGGAAGGTGGGGCGGCCTGGGAGGAAGCGCAAGCACCCCCCGGTGGAAAGTGGTGACACGCCAAAGGACCCTGCGGTGATCTCCAAGTCCCCATCCATGGCCCAGGACCCAGGCGCCTCAGAGCTATTACCCAATGGGGACTTGGAGAAGCGGAGTGAGCCCCAGCCAGAGGAGGGGAGCCCTGCTGGGGGGCAGAAGGGCGGGGCCCCAGCAGAGGGAGAGGGTGCAGCTGAGACCCTGCCTGAAGCCTCAAGAGCAGTGGAAAATGGCTGCTGCACCCCCAAGGAGGGCCGAGGAGCCCCTGCAGAAGTGGGCAAAGAACAGAAGGAGACCAACATCGAATCCATGAAAATGGAGGGCTCCCGGGGCCGGCTGCGGGGTGGCTTGGGCTGGGAGTCCAGCCTCCGTCAGCGGCCCATGCCGAGGCTCACCTTCCAGGCGGGGGACCCCTACTACATCAGCAAGCGCAAGCGGGACGAGTGGCTGGCACGCTGGAAAAGGGAGGCTGAGAAGAAAGCCAAGGTCATTGCAGGAATGAATGCTGTGGAAGAAAACCAGGGGCCCGGGGAGTCTCAGAAGGTGGAGGAGGCCAGCCCTCCTGCTGTGCAGCAGCCCACTGACCCCGCATCCCCCACTGTGGCTACCACGCCTGAGCCCGTGGGGTCCGATGCTGGGGACAAGAATGCCACCAAAGCAGGCGATGACGAGCCAGAGTACGAGGACGGCCGGGGCTTTGGCATTGGGGAGCTGGTGTGGGGGAAACTGCGGGGCTTCTCCTGGTGGCCAGGCCGCATTGTGTCTTGGTGGATGACGGGCCGGAGCCGAGCAGCTGAAGGCACCCGCTGGGTCATGTGGTTCGGAGACGGCAAATTCTCAGTGGTGTGTGTTGAGAAGCTGATGCCACTGAGCTCGTTTTGCAGTGCGTTCCACCAGGCCACGTACAACAAGCAGCCCATGTACCGCAAAGCCATCTACGAGGTCCTGCAGGTGGCCAGCAGCCGCGCGGGGAAGCTGTTCCCGGTGTGCCACGACAGCGATGAGAGTGACACTGCCAAGGCCGTGGAGGTGCAGAACAAGCCCATGATTGAATGGGCCCTGGGGGGCTTCCAGCCCTCTGGCCCTAAGGGCCTAGAGCCACCAGAAGAAGAGAAGAATCCCTACAAAGAAGTGTACACGGACATGTGGGTGGAACCTGAGGCAGCTGCCTACGCACCACCTCCACCAGCCAAAAAGCCCCGGAAGAGCACAGCGGAGAAGCCCAAGGTCAAGGAGATTATTGATGAGCGCACAAGAGAGCGGCTGGTGTACGAGGTGCGGCAGAAGTGCCGGAACATTGAGGACATCTGCATCTCCTGTGGGAGCCTCAATGTTACCCTGGAACACCCCCTCTTCGTTGGAGGAATGTGCCAAAACTGCAAGAACTGCTTTCTGGAGTGTGCGTACCAGTATGACGACGACGGCTACCAGTCCTACTGCACCATCTGCTGTGGGGGCCGTGAGGTGCTCATGTGCGGAAACAACAACTGCTGCAGGTGCTTTTGCGTGGAGTGTGTGGACCTCTTGGTGGGGCCGGGGGCTGCCCAGGCAGCCATTAAGGAAGACCCCTGGAACTGCTACATGTGCGGGCACAAGGGTACCTACGGGCTGCTGCGGCGGCGAGAGGACTGGCCCTCCCGGCTCCAGATGTTCTTCGCTAATAACCACGACCAGGAATTTGACCCTCCAAAGGTTTACCCACCTGTCCCAGCTGAGAAGAGGAAGCCCATCCGGGTGCTGTCTCTCTTTGATGGAATCGCTACAGGGCTCCTGGTGCTGAAGGACTTGGGCATTCAGGTGGACCGCTACATTGCCTCGGAGGTGTGTGAGGACTCCATCACGGTGGGCATGGTGCGGCACCAGGGGAAGATCATGTACGTCGGGGACGTCCGCAGCGTCACACAGAAGCATATCCAGGAGTGGGGCCCATTCGATCTGGTGATTGGGGGCAGTCCCTGCAATGACCTCTCCATCGTCAACCCTGCTCGCAAGGGCCTCTACGAGGGCACTGGCCGGCTCTTCTTTGAGTTCTACCGCCTCCTGCATGATGCGCGGCCCAAGGAGGGAGATGATCGCCCCTTCTTCTGGCTCTTTGAGAATGTGGTGGCCATGGGCGTTAGTGACAAGAGGGACATCTCGCGATTTCTCGAGTCCAACCCTGTGATGATTGATGCCAAAGAAGTGTCAGCTGCACACAGGGCCCGCTACTTCTGGGGTAACCTTCCCGGTATGAACAGGCCGTTGGCATCCACTGTGAATGATAAGCTGGAGCTGCAGGAGTGTCTGGAGCATGGCAGGATAGCCAAGTTCAGCAAAGTGAGGACCATTACTACGAGGTCAAACTCCATAAAGCAGGGCAAAGACCAGCATTTTCCTGTCTTCATGAATGAGAAAGAGGACATCTTATGGTGCACTGAAATGGAAAGGGTATTTGGTTTCCCAGTCCACTATACTGACGTCTCCAACATGAGCCGCTTGGCGAGGCAGAGACTGCTGGGCCGGTCATGGAGCGTGCCAGTCATCCGCCACCTCTTCGCTCCGCTGAAGGAGTATTTTGCGTGTGTGTAA*

*>Orangoutan3A*

*ATGCCCTCCAGCGGCCCCGGGGACACCAGCAGCTCTGCTGCGGAGCGGGAGGAGGACCGAAAGGACGGAGAGGAGCAGGAGGAGCCGCGTGGCAAGGAGGAGCGCCAAGAGCCCAGCACCACGGCACGGAAGGTGGGGCGGCCTGGGAGGAAGCGCAAGCACCCCCCGGTGGAAAGCAGTGACACGCCAAAGGACCCTGCGGTGATCTCCAAGTCCCCATCCATGGCCCAGGACTCAGGCTCCTCAGAGCTATTACCCAATGGGGACTTGGAGAAGCGGAGTGAGCCCCAGCCAGAGGAGGGGAGCCCTGCTGGGGGGCAGAAGGGCGGGGCCCCAGCAGAGGGAGAGGGTGCAGCTGAGACCCTGCCCGAAGCCTCAAGAGCAGTGGAAAATGGCTGCTGCACCCCCAAGGAGGGCCGAGGAGCCCCTGCAGAAGCGGGCAAAGAACAGAAGGAGACCAGCATCGAATCCATGAAAATGGAGGGCTCCCGGGGCCGGCTGCGGGGTGGCTTGGGCTGGGAGTCCAGCCTCCGTCAGCGGCCCATGCCGAGGCACACCTTCCAGGCGGGGGACCCCTACTACATCAGCAAGCGCAAGCGGGACGAGTGGCTGGCACGCTGGAAAAGGGAGGCTGAGAAGAAAGCCAAGGTCATTGCAGGAATGAATGCTGTGGAAGAAAACCAGGGGCCCGGGGAGTCTCAGAAGGTGGAGGAGGCCAGCCCTCCTGCTGTGCAGCAGCCCACTGACCCCGCATCCCCCACTGTGGCTACCACGCCTGAGCCCGTGGGGTCCGATGCCGGGGACAAGAATGCCACCAAAGCAGGCGATGACGAGCCAGAGTACGAGGACGGCCGGGGCTTTGGCATTGGGGAGCTGGTGTGGGGGAAACTGCGGGGCTTCTCCTGGTGGCCAGGCCGCATTGTGTCTTGGTGGATGACGGGCCGGAGCCGAGCAGCTGAAGGCACCCGCTGGGTCATGTGGTTCGGAGACGGCAAATTCTCAGTGGTGTGTGTTGAGAAGCTGATGCCGCTGAGCTCGTTTTGCAGTGCGTTCCACCAGGCCACGTACAACAAGCAGCCCATGTACCGCAAAGCCATCTACGAGGTCCTGCAGGTGGCCAGCAGCCGCGCAGGGAAGCTGTTCCCCGTGTGCCACGACAGCGATGAGAGTGACACTGCCAAGGCCGTGGAGGTGCAGAACAAGCCTATGATTGAATGGGCCCTGGGGGGCTTCCAGCCCTCTGGCCCTAAGGGCCTGGAACCACCAGAAGAAGAGAAGAATCCCTACAAAGAAGTGTACACGGACATGTGGGTGGAACCTGAAGCAGCTGCCTACGCACCACCTCCACCAGCCAAAAAGCCCCGGAAGAGCACAGCGGAGAAGCCCAAGGTCAAGGAGATTATTGATGAGCGCACAAGAGAGCGGCTAGTGTACGAGGTGCGGCAGAAGTGCCGGAACATTGAGGACATCTGCATCTCCTGTGGGAGCCTCAATGTCACCCTGGAACACCCCCTCTTCGTTGGAGGAATGTGCCAAAACTGCAAGAACTGCTTTCTGGAGTGTGCGTACCAGTACGACGACGACGGCTACCAGTCCTACTGCACCATCTGCTGCGGGGGCCGTGAGGTGCTCATGTGCGGAAACAACAACTGCTGCAGGTGCTTTTGCGTGGAGTGTGTGGACCTCTTGGTGGGGCCGGGGGCTGCCCAGGCAGCCATTAAGGAAGACCCCTGGAACTGCTACATGTGTGGGCACAAGGGTACCTACGGGCTGCTGCGGCGGCGAGAGGACTGGCCCTCCCGGCTCCAGATGTTCTTCGCTAATAACCACGACCAGGAATTTGACCCTCCAAAGGTTTACCCACCTGTCCCAGCTGAGAAGAGGAAGCCCATCCGGGTGCTGTCTCTCTTTGATGGAATTGCTACAGGGCTCCTGGTGCTGAAGGACTTGGGCATTCAGGTGGACCGCTACATTGCCTCGGAGGTGTGTGAGGACTCCATCACGGTGGGCATGGTGCGGCACCAGGGGAAGATCATGTACGTCGGGGACGTCCGCAGCGTCACACAGAAGCATATCCAGGAGTGGGGCCCATTCGATCTGGTGATTGGGGGCAGTCCCTGCAATGACCTCTCCATCGTCAACCCTGCTCGCAAGGGCCTCTACGAGGGCACTGGCCGGCTCTTCTTTGAGTTCTACCGCCTCCTGCATGATGCGCGGCCCAAGGAGGGAGATGATCGCCCCTTCTTCTGGCTCTTTGAGAATGTGGTGGCCATGGGCGTTAGTGACAAGAGGGACATCTCGCGATTTCTCGAGTCCAACCCTGTGATGATTGATGCCAAAGAAGTGTCAGCTGCACACAGGGCCCGCTACTTCTGGGGTAACCTTCCCGGTATGAACAGGCCGTTGGCATCCACTGTGAATGATAAGCTGGAGCTGCAGGAGTGTCTGGAGCATGGCAGGATAGCCAAGTTCAGCAAAGTGAGGACCATTACTACGAGGTCAAACTCCATAAAGCAGGGCAAAGACCAGCATTTTCCTGTCTTCATGAATGAGAAAGAGGACATCTTATGGTGCACTGAAATGGAAAGGGTATTTGGTTTCCCAGTCCACTATACTGACGTCTCCAACATGAGCCGCTTGGCGAGGCAGAGACTGCTGGGCCGGTCATGGAGCGTGCCAGTCATCCGCCACCTCTTCGCTCCGCTGAAGGAGTATTTTGCGTGTGTGTAA*

*>CrabMac3A*

*ATGCCCTCCAGCGGCCCCGGGGACACCAGCAGCTCTGCTGCAGAGCGGGAGGAGGACCGAAAGGACGGAGAGGAGCAGGAGGAGCCGCGTGGCAAGGAGGAGCGCCAAGAGCCCAGCACCACGGCACGGAAGGTGGGGAGGCCTGGGAGGAAGCGCAAGCACCCCCCGGTGGAAAGCAGTGACACGCCAAAGGACCCTGCGGTGACCTCCAAGTCCCCATCCATGGCCCAGGACTCAGGCTCCTCAGAGCTGTTACCCAACGGGGACTTGGAGAAGCGGAGTGAGCCCCAGCCAGAGGAGGGGAGCCCTGCTGGGGGGCAGAAGGGCGGGGCCCCAGCAGAGGGAGAGGGTGCAGCTGAGACCCCGCCTGAAGCCTCAAGAGCAGTGGAAAATGGCTGCTGCACCCCCAAGGAGGGCCGAGGAGCACCTGCAGAAGCGGGCAAAGAACAGAAGGAGACCAACATCGAATCCATGAAAATGGAGGGCTCCCGGGGCCGGCTGCGGGGTGGCTTGGGCTGGGAGTCCAGCCTCCGTCAGCGGCCCATGCCGAGGCACACCTTCCAGGCGGGGGACCCCTACTACATCAGCAAGCGCAAGCGGGACGAGTGGCTGGCACGCTGGAAAAGGGAGGCTGAGAAGAAAGCCAAGGTCATTGCAGGAATGAATGCTGTGGAAGAAAACCAGGGGCCCGGGGAGTCTCAGAAGGTGGAGGAGGCCAGCCCTCCTGCTGTGCAGCAGCCCACTGACCCCGCATCCCCCACTGTGGCTACCACACCTGAGCCTGTGGGGGCCGATGCTGGGGACAAGAACGCCACCAAAGCAGGCGATGATGAGCCAGAGTACGAGGACGGCCGGGGCTTTGGCATTGGGGAGCTGGTGTGGGGGAAACTGCGGGGCTTCTCCTGGTGGCCAGGCCGCATTGTGTCTTGGTGGATGACGGGCCGGAGCCGAGCAGCTGAAGGCACCCGCTGGGTCATGTGGTTCGGAGACGGCAAATTCTCAGTGGTGTGTGTTGAGAAGCTGATGCCGCTGAGCTCGTTTTGCAGTGCGTTTCACCAGGCCACGTACAACAAGCAGCCCATGTACCGCAAAGCCATCTACGAGGTCCTGCAGGTGGCCAGCAGCCGTGCAGGGAAGCTGTTCCCGGTGTGCCATGACAGCGATGAGAGTGACACTGCCAAGGCCGTGGAGGTGCAGAACAAACCCATGATTGAATGGGCGCTGGGGGGCTTCCAGCCCTCTGGCCCCAAGGGCCTGGAGCCACCAGAAGAAGAGAAGAATCCCTACAAAGAAGTGTACACAGACATGTGGGTGGAACCTGAGGCAGCTGCCTATGCACCACCTCCACCAGCCAAAAAGCCCCGGAAGAGCACAACGGAGAAGCCCAAGGTCAAGGAGATTATTGATGAGCGCACAAGAGAGCGGCTAGTGTATGAGGTGCGGCAGAAGTGCCGGAACATTGAGGACATCTGCATCTCCTGTGGGAGCCTCAACGTCACCCTGGAACACCCCCTCTTCGTTGGAGGAATGTGCCAAAACTGCAAGAACTGCTTTCTGGAGTGTGCGTATCAGTACGACGACGACGGCTACCAGTCCTACTGCACCATCTGCTGCGGGGGCCGCGAGGTGCTCATGTGCGGAAACAACAACTGCTGCAGGTGCTTTTGCGTGGAGTGTGTAGACCTCTTGGTGGGGCCGGGGGCTGCCCAGGCGGCCATTAAGGAAGACCCCTGGAACTGCTACATGTGCGGGCACAAGGGTACCTATGGGCTGCTGCGGCGGCGAGAGGACTGGCCCTCCCGGCTCCAGATGTTCTTCGCTAATAACCACGACCAGGAATTTGACCCTCCAAAGGTTTACCCACCTGTCCCAGCTGAGAAGAGGAAGCCCATCCGGGTGCTGTCTCTCTTTGATGGAATTGCTACAGGGCTCCTGGTGCTGAAGGACTTGGGCATTCAGGTGGACCGCTACATTGCCTCGGAGGTGTGTGAGGACTCCATCACGGTGGGCATGGTGCGGCACCAGGGGAAGATCATGTACGTCGGGGACGTCCGCAGCGTCACACAGAAGCATATCCAGGAGTGGGGCCCATTCGATCTGGTGATTGGGGGCAGTCCCTGCAATGACCTCTCCATCGTCAACCCTGCTCGCAAGGGCCTTTACGAGGGCACTGGCCGGCTCTTCTTTGAGTTCTACCGCCTCCTGCATGATGCGCGGCCCAAGGAGGGAGATGATCGCCCCTTCTTCTGGCTCTTTGAGAATGTGGTGGCCATGGGCGTTAGTGACAAGAGGGACATCTCACGATTTCTCGAGTCCAACCCTGTGATGATTGATGCCAAAGAAGTGTCAGCTGCACACAGGGCCCGCTACTTCTGGGGTAACCTTCCCGGTATGAACAGGCCGTTGGCATCCACTGTGAATGATAAGCTGGAGCTGCAGGAGTGTCTGGAGCATGGCAGGATAGCCAAGTTCAGCAAAGTGAGGACCATTACTACGAGGTCAAACTCCATAAAGCAGGGCAAAGACCAGCATTTCCCCGTCTTCATGAATGAGAAAGAGGACATCTTATGGTGCACTGAAATGGAAAGGGTATTTGGTTTCCCTGTCCACTATACTGACGTCTCCAACATGAGCCGCTTGGCGAGGCAGAGACTGCTGGGCCGGTCGTGGAGCGTGCCAGTCATCCGCCACCTCTTCGCTCCGCTGAAGGAGTATTTTGCGTGTGTGTAA*

*>RhesusMac3A*

*ATGCCCTCCAGCGGCCCCGGGGACACCAGCAGCTCTGCTGCAGAGCGGGAGGAGGACCGAAAGGACGGAGAGGAGCAGGAGGAGCCGCGTGGCAAGGAGGAGCGCCAAGAGCCCAGCACCACGGCACGGAAGGTGGGGAGGCCTGGGAGGAAGCGCAAGCACCCCCCGGTGGAAAGCAGTGACACGCCAAAGGACCCTGCGGTGACCTCCAAGTCCCCATCCATGGCCCAGGACTCAGGCTCCTCAGAGCTGTTACCCAACGGGGACTTGGAGAAGCGGAGTGAGCCCCAGCCAGAGGAGGGGAGCCCTGCTGGGGGGCAGAAGGGCGGGGCCCCAGCAGAGGGAGAGGGTGCAGCTGAGACCCCGCCTGAAGCCTCAAGAGCAGTGGAAAATGGCTGCTGCACCCCCAAGGAGGGCCGAGGAGCACCTGCAGAAGCGGGCAAAGAACAGAAGGAGACCAACATCGAATCCATGAAAATGGAGGGCTCCCGGGGCCGGCTGCGGGGTGGCTTGGGCTGGGAGTCCAGCCTCCGTCAGCGGCCCATGCCGAGGCACACCTTCCAGGCGGGGGACCCCTACTACATCAGCAAGCGCAAGCGGGACGAGTGGCTGGCACGCTGGAAAAGGGAGGCTGAGAAGAAAGCCAAGGTCATTGCAGGAATGAATGCTGTGGAAGAAAACCAGGGGCCCGGGGAGTCTCAGAAGGTGGAGGAGGCCAGCCCTCCTGCTGTGCAGCAGCCCACTGACCCCGCATCCCCCACTGTGGCTACCACACCTGAGCCTGTGGGGGCCGATGCTGGGGACAAGAACGCCACCAAAGCAGGCGATGATGAGCCAGAGTACGAGGACGGCCGGGGCTTTGGCATTGGGGAGCTGGTGTGGGGGAAACTGCGGGGCTTCTCCTGGTGGCCAGGCCGCATTGTGTCTTGGTGGATGACGGGCCGGAGCCGAGCAGCTGAAGGCACCCGCTGGGTCATGTGGTTCGGAGACGGCAAATTCTCAGTGGTGTGTGTTGAGAAGCTGATGCCGCTGAGCTCGTTTTGCAGTGCGTTTCACCAGGCCACGTACAACAAGCAGCCCATGTACCGCAAAGCCATCTACGAGGTCCTGCAGGTGGCCAGCAGCCGTGCAGGGAAGCTGTTCCCGGTGTGCCATGACAGCGATGAGAGTGACACTGCCAAGGCCGTGGAGGTGCAGAACAAACCCATGATTGAATGGGCGCTGGGGGGCTTCCAGCCCTCTGGCCCCAAGGGCCTGGAGCCACCAGAAGAAGAGAAGAATCCCTACAAAGAAGTGTACACAGACATGTGGGTGGAACCTGAGGCAGCTGCCTATGCACCACCTCCACCAGCCAAAAAGCCCCGGAAGAGCACAACGGAGAAGCCCAAGGTCAAGGAGATTATTGATGAGCGCACAAGAGAGCGGCTAGTGTATGAGGTGCGGCAGAAGTGCCGGAACATTGAGGACATCTGCATCTCCTGTGGGAGCCTCAACGTCACCCTGGAACACCCCCTCTTCGTTGGAGGAATGTGCCAAAACTGCAAGAACTGCTTTCTGGAGTGTGCGTATCAGTACGACGACGACGGCTACCAGTCCTACTGCACCATCTGCTGCGGGGGCCGCGAGGTGCTCATGTGCGGAAACAACAACTGCTGCAGGTGCTTTTGCGTGGAGTGTGTAGACCTCTTGGTGGGGCCGGGGGCTGCCCAGGCGGCCATTAAGGAAGACCCCTGGAACTGCTACATGTGCGGGCACAAGGGTACCTATGGGCTGCTGCGGCGGCGAGAGGACTGGCCCTCCCGGCTCCAGATGTTCTTCGCTAATAACCACGACCAGGAATTTGACCCTCCAAAGGTTTACCCACCTGTCCCAGCTGAGAAGAGGAAGCCCATCCGGGTGCTGTCTCTCTTTGATGGAATTGCTACAGGGCTCCTGGTGCTGAAGGACTTGGGCATTCAGGTGGACCGCTACATTGCCTCGGAGGTGTGTGAGGACTCCATCACGGTGGGCATGGTGCGGCACCAGGGGAAGATCATGTACGTCGGGGACGTCCGCAGCGTCACACAGAAGCATATCCAGGAGTGGGGCCCATTCGATCTGGTGATTGGGGGCAGTCCCTGCAATGACCTCTCCATCGTCAACCCTGCTCGCAAGGGCCTTTACGAGGGCACTGGCCGGCTCTTCTTTGAGTTCTACCGCCTCCTGCATGATGCGCGGCCCAAGGAGGGAGATGATCGCCCCTTCTTCTGGCTCTTTGAGAATGTGGTGGCCATGGGCGTTAGTGACAAGAGGGACATCTCACGATTTCTCGAGTCCAACCCTGTGATGATTGATGCCAAAGAAGTGTCAGCTGCACACAGGGCCCGCTACTTCTGGGGTAACCTTCCCGGTATGAACAGGCCGTTGGCATCCACTGTGAATGATAAGCTGGAGCTGCAGGAGTGTCTGGAGCATGGCAGGATAGCCAAGTTCAGCAAAGTGAGGACCATTACTACGAGGTCAAACTCCATAAAGCAGGGCAAAGACCAGCATTTCCCCGTCTTCATGAATGAGAAAGAGGACATCTTATGGTGCACTGAAATGGAAAGGGTATTTGGTTTCCCTGTCCACTATACTGACGTCTCCAACATGAGCCGCTTGGCGAGGCAGAGACTGCTGGGCCGGTCGTGGAGCGTGCCAGTCATCCGCCACCTCTTCGCTCCGCTGAAGGAGTATTTTGCGTGTGTGTAA*

*>PigTailMac3A*

*ATGCCCTCCAGCGGCCCCGGGGACACCAGCAGCTCTGCTGCAGAGCGGGAGGAGGACCGAAAGGACGGAGAGGAGCAGGAGGAGCCGCGTGGCAAGGAGGAGCGCCAAGAGCCCAGCACCACGGCACGGAAGGTGGGGAGGCCTGGGAGGAAGCGCAAGCACCCCCCGGTGGAAAGCAGTGACACGCCAAAGGACCCTGCGGTGACCTCCAAGTCCCCATCCATGGCCCAGGACTCAGGCTCCTCAGAGCTGTTACCCAACGGGGACTTGGAGAAGCGGAGTGAGCCCCAGCCAGAGGAGGGGAGCCCTGCTGGGGGGCAGAAGGGCGGGACCCCAGCAGAGGGAGAGGGTGCAGCTGAGACCCCGCCTGAAGCCTCAAGAGCAGTGGAAAATGGCTGCTGTACCCCCAAGGAGGGCCGAGGAGCACCTGCAGAAGCGGGCAAAGAACAGAAGGAGACCAACATCGAATCCATGAAAATGGAGGGCTCCCGGGGCCGGCTGCGGGGTGGCTTGGGCTGGGAGTCCAGCCTCCGTCAGCGGCCCATGCCGAGGCACACCTTCCAGGCGGGGGACCCCTACTACATCAGCAAGCGCAAGCGGGACGAGTGGCTGGCACGCTGGAAAAGGGAGGCTGAGAAGAAAGCCAAGGTCATTGCAGGAATGAATGCTGTGGAAGAAAACCAGGGGCCCGGGGAGTCTCAGAAGGTGGAGGAAGCCAGCCCTCCTGCTGTGCAGCAGCCCACTGACCCCGCATCCCCCACTGTGGCTACCACACCTGAGCCTGTGGGGGCCGATGCTGGGGACAAGAACGCCACCAAAGCAGGCGATGATGAGCCAGAGTACGAGGACGGCCGGGGCTTTGGCATTGGGGAGCTGGTGTGGGGGAAACTGCGGGGCTTCTCCTGGTGGCCAGGCCGCATTGTGTCTTGGTGGATGACGGGCCGGAGCCGAGCAGCTGAAGGCACCCGCTGGGTCATGTGGTTCGGAGACGGCAAATTCTCAGTGGTGTGTGTTGAGAAGCTGATGCCGCTGAGCTCGTTTTGCAGTGCGTTTCACCAGGCCACGTACAACAAGCAGCCCATGTACCGCAAAGCCATCTACGAGGTCCTGCAGGTGGCCAGCAGCCGTGCAGGGAAGCTGTTCCCGGTGTGCCATGACAGCGATGAGAGTGACACTGCCAAGGCTGTGGAGGTGCAGAACAAACCCATGATTGAATGGGCGCTGGGGGGCTTCCAGCCCTCTGGCCCCAAGGGCCTGGAGCCACCAGAAGAAGAGAAGAATCCCTACAAAGAAGTGTACACAGACATGTGGGTGGAACCTGAGGCAGCTGCCTATGCACCACCTCCACCAGCCAAAAAGCCCCGGAAGAGCACAACGGAGAAGCCCAAGGTCAAGGAGATTATTGATGAGCGCACAAGAGAGCGGCTGGTGTATGAGGTGCGGCAGAAGTGCCGGAACATTGAGGACATCTGCATCTCCTGTGGGAGCCTCAACGTCACCCTGGAACACCCCCTCTTCGTTGGAGGAATGTGCCAAAACTGCAAGAACTGCTTTCTGGAGTGTGCGTATCAGTACGACGACGACGGCTACCAGTCCTACTGCACCATCTGCTGCGGGGGCCGCGAGGTGCTCATGTGCGGAAACAACAACTGCTGCAGGTGCTTTTGCGTGGAGTGTGTAGACCTCTTGGTGGGGCCGGGGGCTGCCCAGGCGGCCATTAAGGAAGACCCCTGGAACTGCTACATGTGCGGGCACAAGGGTACCTATGGGCTGCTGCGGCGGCGAGAGGACTGGCCCTCCCGGCTCCAGATGTTCTTCGCTAATAACCACGACCAGGAATTTGACCCTCCAAAGGTTTACCCACCTGTCCCAGCTGAGAAGAGGAAGCCCATCCGAGTGCTGTCTCTCTTTGATGGAATTGCTACAGGGCTCCTGGTGCTGAAGGACTTGGGCATTCAGGTGGACCGCTACATTGCCTCGGAGGTGTGTGAGGACTCCATCACGGTGGGCATGGTGCGGCACCAGGGGAAGATCATGTACGTCGGGGACGTCCGCAGCGTCACACAGAAGCATATCCAGGAGTGGGGCCCATTCGATCTGGTGATTGGGGGCAGTCCCTGCAATGACCTCTCCATCGTCAACCCTGCTCGCAAGGGCCTTTACGAGGGCACTGGCCGGCTCTTCTTTGAGTTCTACCGCCTCCTGCATGATGCGCGGCCCAAGGAGGGAGATGATCGCCCCTTCTTCTGGCTCTTTGAGAATGTGGTGGCCATGGGCGTTAGTGACAAGAGGGACATCTCACGATTTCTCGAGTCCAACCCTGTGATGATTGATGCCAAAGAAGTGTCAGCTGCACACAGGGCCCGCTACTTCTGGGGTAACCTTCCCGGTATGAACAGGCCGTTGGCATCCACTGTGAATGATAAGCTGGAGCTGCAGGAGTGTCTGGAGCATGGCAGGATAGCCAAGTTCAGCAAAGTGAGGACCATTACTACGAGGTCAAACTCCATAAAGCAGGGCAAAGACCAGCATTTCCCCGTCTTCATGAATGAGAAAGAGGACATCTTATGGTGCACTGAAATGGAAAGGGTATTTGGTTTCCCTGTCCACTATACTGACGTCTCCAACATGAGCCGCTTGGCGAGGCAGAGACTGCTGGGCCGGTCGTGGAGCGTGCCAGTCATCCGCCACCTCTTCGCTCCGCTGAAGGAGTATTTTGCGTGTGTGTAA*

*>Mangabey3A*

*ATGCCCTCCAGCGGCCCCGGGGACACCAGCAGCTCTGCTGCAGAGCGGGAGGAGGACCGAAAGGACGGAGAGGAGCAGGAGGAGCCGCGTGGCAAGGAGGAGCGCCAAGAGCCCAGCACCACGGCACGGAAGGTGGGGAGGCCTGGGAGGAAGCGCAAGCACCCCCCGGTGGAAAGCAGTGACACGCCAAAGGACCCTGCGGTGACCTCCAAGTCCCCATCCATGGCCCAGGACTCAGGCTCCTCAGAGCTGTTACCCAATGGGGACTTGGAGAAGCGGAGTGAGCCCCAGCCAGAGGAGGGGAGCCCTGCTGGGGGGCAGAAGGGCGGGACCCCAGCAGAGGGAGAGGGTGCAGCTGAGACCCCGCCTGAAGCCTCAAGAGCAGTGGAAAATGGCTGCTGCACCCCCAAGGAGGGCCGAGGAGCACCTGCAGAAGCGGGCAAAGAACAGAAGGAGACCAACATCGAATCCATGAAAATGGAGGGCTCCCGGGGCCGGCTGCGGGGTGGCTTGGGCTGGGAGTCCAGCCTCCGTCAGCGGCCCATGCCGAGGCACACCTTCCAGGCGGGGGACCCCTACTACATCAGCAAGCGCAAGCGGGACGAGTGGCTGGCACGCTGGAAAAGGGAGGCTGAGAAGAAAGCCAAGGTCATTGCAGGAATGAATGCTGTGGAAGAAAACCAGGGGCCCGGGGAGTCTCAGAAGGTGGAGGAGGCCAGCCCTCCTGCTGTGCAGCAGCCCACTGACCCCGCATCCCCCACTGTGGCTACCACACCTGAGCCTGTGGGGGCCGATGCTGGGGACAAGAATGCCACCAAAGCAGGCGATGATGAGCCAGAGTACGAGGACGGCCGGGGCTTTGGCATTGGGGAGCTGGTGTGGGGGAAACTGCGGGGCTTCTCCTGGTGGCCAGGCCGCATTGTGTCTTGGTGGATGACGGGCCGGAGCCGAGCAGCTGAAGGCACCCGCTGGGTCATGTGGTTCGGAGACGGCAAATTCTCAGTGGTGTGTGTTGAGAAGCTGATGCCGCTGAGCTCGTTTTGCAGTGCGTTCCACCAGGCCACGTACAACAAGCAGCCCATGTACCGCAAAGCCATCTACGAGGTCCTGCAGGTGGCCAGCAGCCGTGCAGGGAAGCTGTTCCCGGTGTGCCATGACAGCGACGAGAGTGACACTGCCAAGGCCGTGGAGGTGCAAAACAAACCCATGATTGAATGGGCGCTGGGGGGCTTCCAGCCCTCTGGCCCCAAGGGCCTGGAGCCACCAGAAGAAGAGAAGAATCCCTACAAAGAAGTGTACACAGACATGTGGGTGGAACCTGAGGCAGCTGCCTATGCACCACCTCCACCAGCCAAAAAGCCCCGGAAGAGCACAACGGAGAAGCCCAAGGTCAAGGAGATTATTGATGAGCGCACAAGAGAGCGGCTGGTGTATGAGGTGCGGCAGAAGTGCCGGAACATTGAGGACATCTGCATCTCCTGTGGGAGCCTCAACGTCACCCTGGAACACCCCCTCTTCGTTGGAGGAATGTGCCAAAACTGCAAGAACTGCTTTCTGGAGTGTGCGTATCAGTACGACGACGACGGCTACCAGTCCTACTGCACCATCTGCTGCGGGGGCCGCGAGGTGCTCATGTGCGGAAACAACAACTGCTGCAGGTGCTTTTGCGTGGAGTGTGTAGACCTCTTGGTGGGGCCGGGGGCTGCCCAGGCGGCCATTAAGGAAGACCCCTGGAACTGCTACATGTGCGGGCACAAGGGTACCTATGGGCTGCTGCGGCGGCGAGAGGACTGGCCCTCCCGGCTCCAGATGTTCTTCGCTAATAACCACGACCAGGAATTTGACCCTCCAAAGGTTTACCCACCTGTCCCAGCTGAGAAGAGGAAGCCCATCCGGGTGCTGTCTCTCTTTGATGGAATTGCTACAGGGCTCCTGGTGCTGAAGGACTTGGGCATTCAGGTGGACCGCTACATTGCCTCGGAGGTGTGTGAGGACTCCATCACGGTGGGCATGGTGCGGCACCAGGGGAAGATCATGTACGTCGGGGACGTCCGCAGCGTCACACAGAAGCATATCCAGGAGTGGGGCCCATTCGATCTGGTGATTGGGGGCAGTCCCTGCAATGACCTCTCCATCGTCAACCCTGCTCGCAAGGGCCTTTACGAGGGCACTGGCCGGCTCTTCTTTGAGTTCTACCGCCTCCTGCATGATGCGCGGCCCAAGGAGGGAGATGATCGCCCCTTCTTCTGGCTCTTTGAGAATGTGGTGGCCATGGGCGTTAGTGACAAGAGGGACATCTCACGATTTCTCGAGTCCAACCCTGTGATGATTGATGCCAAAGAAGTGTCAGCTGCACACAGGGCCCGCTACTTCTGGGGTAACCTTCCCGGTATGAACAGGCCGTTGGCATCCACTGTGAATGATAAGCTGGAGCTGCAGGAGTGTCTGGAGCATGGCAGGATAGCCAAGTTCAGCAAAGTGAGGACCATTACTACGAGGTCAAACTCCATAAAGCAGGGCAAAGACCAGCATTTCCCCGTCTTCATGAATGAGAAAGAGGACATCTTATGGTGCACTGAAATGGAAAGGGTATTTGGTTTCCCTGTCCACTATACTGACGTCTCCAACATGAGCCGCTTGGCGAGGCAGAGACTGCTGGGCCGGTCGTGGAGCGTGCCAGTCATCCGCCACCTCTTCGCTCCGCTGAAGGAGTATTTTGCGTGTGTGTAA*

*>Drill3A*

*ATGCCCTCCAGCGGCCCCGGGGACACCAGCAGCTCTGCTGCAGAGCGGGAGGAGGACCGAAAGGACGGAGAGGAGCAGGAGGAGCCGCGTGGCAAGGAGGAGCGCCAAGAGCCCAGCACCACGGCACGGAAGGTGGGGAGGCCTGGGAGGAAGCGCAAGCACCCCCCGGTGGAAAGCAGTGACACGCCAAAGGACCCTGCGGTGACCTCCAAGTCCCCATCCATGGCCCAGGACTCAGGCTCCTCAGAGCTGTTACCCAATGGGGACTTGGAGAAGCGGAGTGAGCCCCAGCCAGAGGAGGGGAGCCCTGCTGGGGGGCAGAAGGGCGGGACCCCAGCAGAGGGAGAGGGTGCAGCTGAGACCCCGCCTGAAGCCTCAAGAGCAGTGGAAAATGGCTGCTGCACCCCCAAGGAGGGCCGAGGAGCACCTGCAGAAGCGGGCAAAGAACAGAAGGAGACCAACATCGAATCCATGAAAATGGAGGGCTCCCGGGGCCGGCTGCGGGGTGGCTTGGGCTGGGAGTCCAGCCTCCGTCAGCGGCCCATGCCGAGGCACACCTTCCAGGCGGGGGACCCCTACTACATCAGCAAGCGCAAGCGGGACGAGTGGCTGGCACGCTGGAAAAGGGAGGCTGAGAAGAAAGCCAAGGTCATTGCAGGAATGAATGCTGTGGAAGAAAACCAGGGGCCCGGGGAGTCTCAGAAGGTGGAGGAGGCCAGCCCTCCTGCTGTGCAGCAGCCCACTGACCCCGCATCCCCCACTGTGGCTACCACACCTGAGCCTGTGGGGGCCGATGCTGGGGACAAGAATGCCACCAAAGCAGGCGATGATGAGCCAGAGTACGAGGACGGCCGGGGCTTTGGCATTGGGGAGCTGGTGTGGGGGAAACTGCGGGGCTTCTCCTGGTGGCCAGGCCGCATTGTGTCTTGGTGGATGACGGGCCGGAGCCGAGCAGCTGAAGGCACCCGCTGGGTCATGTGGTTCGGAGACGGCAAATTCTCAGTGGTGTGTGTTGAGAAGCTGATGCCGCTGAGCTCGTTTTGCAGTGCGTTCCACCAGGCCACGTACAACAAGCAGCCCATGTACCGCAAAGCCATCTACGAGGTCCTGCAGGTGGCCAGCAGCCGTGCAGGGAAGCTGTTCCCGGTGTGCCATGACAGCGACGAGAGTGACACTGCCAAGGCCGTGGAGGTGCAGAACAAACCCATGATTGAATGGGCGCTGGGGGGCTTCCAGCCCTCTGGCCCCAAGGGCCTGGAGCCACCAGAAGAAGAGAAGAATCCCTACAAAGAAGTGTACACAGACATGTGGGTGGAACCTGAGGCAGCTGCCTATGCACCACCTCCACCAGCCAAAAAGCCCCGGAAGAGCACAACGGAGAAACCCAAGGTCAAGGAGATTATTGATGAGCGCACAAGAGAGCGGCTGGTGTATGAGGTGCGGCAGAAGTGCCGGAACATTGAGGACATCTGCATCTCCTGTGGGAGCCTCAACGTCACCCTGGAACACCCCCTCTTCGTTGGAGGAATGTGCCAAAACTGCAAGAACTGCTTTCTGGAGTGTGCGTATCAGTACGACGACGACGGCTACCAGTCCTACTGCACCATCTGCTGCGGGGGCCGCGAGGTGCTCATGTGCGGAAACAACAACTGCTGCAGGTGCTTTTGCGTGGAGTGTGTAGACCTCTTGGTGGGGCCGGGGGCTGCCCAGGCGGCCATTAAGGAAGACCCCTGGAACTGCTACATGTGCGGGCACAAGGGTACCTATGGGCTGCTGCGGCGGCGAGAGGACTGGCCCTCCCGGCTCCAGATGTTCTTCGCTAATAACCACGACCAGGAATTTGACCCTCCAAAGGTTTACCCACCTGTCCCAGCTGAGAAGAGGAAGCCCATCCGGGTGCTGTCTCTCTTTGATGGAATTGCTACAGGGCTCCTGGTGCTGAAGGACTTGGGCATTCAGGTGGACCGCTACATTGCCTCGGAGGTGTGTGAGGACTCCATCACGGTGGGCATGGTGCGGCACCAGGGGAAGATCATGTACGTCGGGGACGTCCGCAGCGTCACACAGAAGCATATCCAGGAGTGGGGCCCATTCGATCTGGTGATTGGGGGCAGTCCCTGCAATGACCTCTCCATCGTCAACCCTGCTCGCAAGGGCCTTTACGAGGGCACTGGCCGGCTCTTCTTTGAGTTCTACCGCCTCCTGCATGATGCGCGGCCCAAGGAGGGAGATGATCGCCCCTTCTTCTGGCTCTTTGAGAATGTGGTGGCCATGGGCGTTAGTGACAAGAGGGACATCTCACGATTTCTCGAGTCCAACCCTGTGATGATTGATGCCAAAGAAGTGTCAGCTGCACACAGGGCCCGCTACTTCTGGGGTAACCTTCCCGGTATGAACAGGCCGTTGGCATCCACTGTGAATGATAAGCTGGAGCTGCAGGAGTGTCTGGAGCATGGCAGGATAGCCAAGTTCAGCAAAGTGAGGACCATTACTACGAGGTCAAACTCCATAAAGCAGGGCAAAGACCAGCATTTCCCCGTCTTCATGAATGAGAAAGAGGACATCTTATGGTGCACTGAAATGGAAAGGGTATTTGGTTTCCCTGTCCACTATACTGACGTCTCCAACATGAGCCGCTTGGCGAGGCAGAGACTGCTGGGCCGGTCGTGGAGCGTGCCAGTCATCCGCCACCTCTTCGCTCCGCTGAAGGAGTATTTTGCGTGTGTGTAA*

*>RedColobus3A*

*ATGCCCTCCAGCGGCCCCGGGGACACCAGCAGCTCTGCTGCAGAGCGGGAGGAGGACCGAAAGGACGGAGAGGAGCAAGAGGAGCCGCGTGGCAAGGAGGAGCGCCAAGAGCCCAGCACCACGGCACGGAAGGTGGGGAGGCCTGGGAGGAAGCGCAAGCACCCCCCGGTGGAAAGCAGTGATACACCAAAGGACCCTGCGGTGACCTCCAAGTCCCCATCCATGGCCCAGGACTCAGGCCCCTCAGAGCTGTTACCCAACGGGGACTTGGAGAAGCGGAGTGAGCCCCAGCCAGAGGAGGGGAGCCCTGCTGGGGGGCAGAAGGGCGGGGCCCCAGCAGAGGGAGAGGGTGCAGCTGAGACCCCGCCTGAAGCCTCAAGAGCAGTGGAAAATGGCTGCTGCACCCCCAAGGAGGGCCGAGGAGCACCTGCAGAAGCGGGCAAAGAACAGAAGGAGACCAACATCGAATCCATGAAAATGGAGGGCTCCCGGGGCCGGCTGCGGGGTGGCTTGGGCTGGGAGTCCAGCCTCCGTCAGCGGCCCATGCCGAGGCACACCTTCCAGGCGGGGGACCCCTACTACATCAGCAAGCGCAAGCGGGACGAGTGGCTGGCACGCTGGAAAAGGGAGGCTGAGAAGAAAGCCAAGGTCATTGCAGGAATGAATGCTGTGGAAGAAAACCAGGGGCCCGGGGAGTCTCAGAAGGTGGAGGAGGCCAGCCCTCCTGCTGTGCAGCAGCCCACTGACCCCGCATCCCCCACTGTGGCTACCACACCTGAGCCTGTGGGGGCTGATGCTGGGGACAAGAACGCCACCAAAGCAGGCGATGATGAACCAGAGTACGAGGACGGCCGGGGCTTTGGCATTGGGGAGCTGGTGTGGGGGAAACTGCGGGGCTTCTCCTGGTGGCCAGGCCGCATTGTGTCTTGGTGGATGACGGGCCGGAGCCGAGCAGCTGAAGGCACCCGCTGGGTCATGTGGTTTGGAGACGGCAAATTCTCAGTGGTGTGTGTTGAGAAGCTGATGCCGCTGAGCTCGTTTTGCAGTGCGTTCCACCAGGCCACGTACAACAAGCAGCCCATGTACCGCAAAGCCATCTACGAGGTCCTGCAGGTGGCCAGCAGCCGCGCAGGGAAGCTGTTCCCGGTGTGCCATGACAGCGACGAGAGTGACACTGCCAAGGCCGTGGAGGTGCAGAACAAGCCCATGATTGAATGGGCCCTGGGGGGATTCCAGCCCTCTGGCCCCAAGGGCCTGGAGCCACCAGAAGAAGAGAAGAATCCCTACAAAGAAGTGTACACAGACATGTGGGTGGAACCTGAGGCAGCTGCCTATGCACCACCTCCACCAGCCAAAAAGCCCCGGAAGAGCACAACAGAGAAGCCCAAGGTCAAGGAGATTATTGATGAGCGCACAAGAGAGCGGCTGGTGTACGAGGTGCGGCAGAAGTGCCGGAACATTGAGGACATCTGCATCTCCTGTGGGAGCCTCAACGTCACCCTGGAACACCCCCTCTTCGTTGGAGGAATGTGCCAAAACTGCAAGAACTGCTTTCTGGAGTGTGCGTATCAGTACGACGACGACGGCTACCAGTCCTACTGCACCATCTGCTGCGGGGGCCGCGAGGTGCTCATGTGCGGAAACAACAACTGCTGCAGGTGCTTTTGCGTGGAGTGTGTGGACCTCTTGGTGGGGCCGGGGGCTGCCCAGGCGGCCATTAAGGAAGACCCCTGGAACTGCTACATGTGCGGGCACAAGGGTACCTATGGGCTGCTGCGGCGGCGAGAGGACTGGCCCTCCCGGCTCCAGATGTTCTTCGCTAATAACCACGACCAGGAATTTGATCCTCCAAAGGTTTACCCACCTGTCCCAGCTGAGAAGAGGAAGCCCATCCGGGTGCTGTCTCTCTTTGATGGAATTGCTACAGGGCTCCTGGTGCTGAAGGACTTGGGCATTCAGGTGGACCGCTACATTGCCTCGGAGGTGTGTGAGGACTCCATCACGGTGGGCATGGTGCGGCACCAGGGGAAGATCATGTACGTCGGGGACGTCCGCAGCGTCACACAGAAGCATATCCAGGAGTGGGGCCCATTCGATCTGGTGATTGGGGGCAGTCCCTGCAATGACCTCTCCATCGTCAACCCTGCTCGCAAGGGCCTCTACGAGGGCACTGGCCGGCTCTTCTTTGAGTTCTACCGCCTCCTGCATGATGCGCGGCCCAAGGAGGGAGATGATCGCCCCTTCTTCTGGCTCTTTGAGAATGTGGTGGCCATGGGCGTTAGTGACAAGAGGGACATCTCACGATTTCTCGAGTCCAACCCTGTGATGATTGATGCCAAAGAAGTGTCAGCTGCACACAGGGCCCGCTACTTCTGGGGTAACCTTCCCGGTATGAACAGGCCGTTGGCATCCACTGTGAATGATAAGCTGGAGCTGCAGGAGTGTCTGGAGCATGGCAGGATAGCCAAGTTCAGCAAAGTGAGGACCATTACTACGAGGTCAAACTCCATAAAGCAGGGCAAAGACCAGCATTTCCCCGTCTTCATGAATGAGAAAGAGGACATCTTATGGTGCACTGAAATGGAAAGGGTATTTGGTTTCCCTGTCCACTATACTGACGTCTCCAACATGAGCCGCTTGGCGAGGCAGAGACTGCTGGGCCGGTCGTGGAGCGTGCCAGTCATCCGCCACCTCTTCGCTCCGCTGAAGGAGTATTTTGCGTGTGTGTAA*

*>AfrGreenMonkey3A*

*ATGCCCTCCAGCGGCCCCGGGGACACCAGCAGCTCTGCTGCAGAGCGGGAGGAGGACCGAAAGGACGGAGAGGAGCAGGAGGAGCCGCGTGGCAAGGAGGAGCGCCAAGAGCCCAGCACCACAGCACGGAAGGTGGGGAGGCCTGGGAGGAAGCGCAAGCACCCCCCGGTGGAAAGCAGTGACACGCCAAAGGACCCTGCGGTGACCTCCAAGTCCCCATCCATGGCCCAGGACTCAGGCCCCTCAGAGCTGTTACCCAACGGGGACTTGGAGAAGCGGAGTGAGCCCCAGCCAGAGGAGGGGAGCCCTGCTGGGGGGCAGAAGGGCGGGGCCCCAGCAGAGGGAGAGGGTGCAGCTGAGACCCCGCCTGAAGCCTCGAGAGCAGTGGAAAATGGCTGCTGCACCCCCAAGGAGGGCCGAGGAGCACCTGCAGAAGCGGGCAAAGAACAGAAGGAGCCCAACATCGAATCCATGAAAATGGAGGGCTCCCGGGGCCGGCTGCGGGGTGGCCTGGGCTGGGAGTCCAGCCTCCGTCAGCGGCCCATGCCGAGGCACACCTTCCAGGCGGGGGACCCCTACTACATCAGCAAGCGCAAGCGGGACGAGTGGCTGGCACGCTGGAAAAGGGAGGCTGAGAAGAAAGCCAAGGTCATTGCAGGAATGAATGCTGTGGAAGAAAACCAGGGGCCCGGGGAGTCTCAGAAGGTGGAGGAGGCCAGCCCTCCTGCTGTGCAGCAGCCCACTGACCCCGCATCCCCCACTGTGGCTACCACACCTGAGCCTGTGGGGGCCGATGCTGGGGACAAGAACGCCACCAAAGCAGGCGATGATGAGCCAGAGTACGAGGACGGCCGGGGCTTTGGCATTGGGGAGCTGGTGTGGGGGAAACTGCGGGGCTTCTCCTGGTGGCCAGGCCGCATTGTGTCTTGGTGGATGACGGGCCGGAGCCGAGCAGCTGAAGGCACCCGCTGGGTCATGTGGTTCGGAGACGGCAAATTCTCAGTGGTGTGTGTTGAGAAGCTGATGCCGCTGAGCTCGTTTTGCAGTGCGTTCCACCAGGCCACGTACAACAAGCAGCCCATGTACCGCAAAGCCATCTACGAGGTCCTGCAGGTGGCCAGCAGCCGTGCAGGGAAGCTGTTCCCGGTGTGCCATGACAGCGACGAGAGTGACACTGCCAAGGCCGTGGAGGTGCAGAACAAACCCATGATTGAATGGGCACTGGGGGGCTTCCAGCCCTCTGGCCCCAAGGGCCTGGAGCCACCAGAAGAAGAGAAGAATCCCTACAAAGAAGTGTACACAGACATGTGGGTGGAACCTGAGGCAGCTGCCTATGCACCACCTCCACCAGCCAAAAAGCCCCGGAAGAGCACAACGGAGAAGCCCAAGGTCAAGGAGATTATTGATGAGCGCACAAGAGAGCGGCTGGTGTATGAGGTGCGGCAGAAGTGCCGGAACATTGAGGACATCTGCATCTCCTGTGGGAGCCTCAACGTCACCCTGGAACACCCCCTCTTCGTTGGAGGAATGTGCCAAAACTGCAAGAACTGCTTTCTGGAGTGTGCGTATCAGTACGACGACGACGGCTACCAGTCCTACTGCACCATCTGCTGCGGGGGCCGCGAGGTGCTCATGTGCGGAAACAACAACTGCTGCAGGTGCTTTTGCGTGGAGTGTGTGGACCTCTTGGTGGGGCCGGGGGCTGCCCAGGCGGCCATTAAGGAAGACCCCTGGAACTGCTACATGTGCGGGCACAAGGGTACCTATGGGCTGCTGCGGCGGCGAGAGGACTGGCCCTCCCGGCTCCAGATGTTCTTCGCTAATAACCACGACCAGGAATTTGACCCTCCAAAGGTTTACCCACCTGTCCCAGCTGAGAAGAGGAAGCCCATCCGGGTGCTGTCTCTCTTTGATGGAATTGCTACAGGGCTCCTGGTGCTGAAGGACTTGGGCATTCAGGTGGACCGCTACATTGCCTCGGAGGTGTGTGAGGACTCCATCACGGTGGGCATGGTGCGGCACCAGGGGAAGATCATGTACGTCGGGGACGTCCGCAGCGTCACACAGAAGCATATCCAGGAGTGGGGCCCATTCGATCTGGTGATTGGGGGCAGTCCCTGCAATGACCTCTCCATCGTCAACCCTGCTCGCAAGGGCCTTTACGAGGGCACTGGCCGGCTCTTCTTTGAGTTCTACCGCCTCCTGCATGATGCGCGGCCCAAGGAGGGAGATGATCGCCCCTTCTTCTGGCTCTTTGAGAATGTGGTGGCCATGGGCGTTAGTGACAAGAGGGACATCTCACGATTTCTCGAGTCCAACCCTGTGATGATTGATGCCAAAGAAGTGTCAGCTGCACACAGGGCCCGCTACTTCTGGGGTAACCTTCCCGGTATGAACAGGCCGTTGGCATCCACTGTGAATGATAAGCTGGAGCTGCAGGAGTGTCTGGAGCATGGCAGGATAGCCAAGTTCAGCAAAGTGAGGACCATTACTACGAGGTCAAACTCCATAAAGCAGGGCAAAGACCAGCATTTCCCCGTCTTCATGAATGAGAAAGAGGACATCTTATGGTGCACTGAAATGGAAAGGGTATTTGGTTTCCCTGTCCACTATACTGACGTCTCCAACATGAGCCGCTTGGCGAGGCAGAGACTGCTGGGCCGGTCGTGGAGCGTGCCAGTCATCCGCCACCTCTTCGCTCCGCTGAAGGAGTATTTTGCGTGTGTGTAA*

*>BlackSnubNose3A*

*ATGCCCTCCAGCGGCCCCGGGGACACCAGCAGCTCTGCTGCAGAGCGGGAGGAGGACCGAAAGGACGGAGAGGAGCAAGAGGAGCCGCGTGGCAAGGAGGAGCGCCAAGAGCCCAGCACCACGGCACGGAAGGTGGGGAGGCCTGGGAGGAAGCGCAAGCACCCCCCGGTGGAAAGCAGTGACACGCCAAAGGACCCTGCGGTGACCTCCAAGTCCCCATCCATGGCCCAGGACTCAGGCCCCTCAGAGCTGTTACCCAACGGGGACTTGGAGAAGCGGAGTGAGCCCCAGCCAGAGGAGGGGAGCCCTGCTGGGGGGCAGAAGGGCGGGGCCCCAGCAGAGGGAGAGGGTGCAGCTGAGACCCCGCCTGAAGCCTCGAGAGCAGTGGAAAATGGCTGCTGCACCCCCAAGGAGGGCCGAGGAGCACCTGCAGAAGTGGGCAAAGAACAGAAGGAGACCAACATCGAATCCATGAAAATGGAGGGCTCCCGGGGCCGGCTGCGGGGTGGCTTGGGCTGGGAGTCCAGCCTCCGTCAGCGGCCCATGCCGAGGCACACCTTCCAGGCGGGGGACCCCTACTACATCAGCAAGCGCAAGCGGGACGAGTGGCTGGCACGCTGGAAAAGGGAGGCTGAGAAGAAAGCCAAGGTCATTGCAGGAATGAATGCTGTGGAAGAAAACCAGGGGCCTGGGGAGTCTCAGAAGGTGGAGGAGGCCAGCCCTCCTGCTGTGCAGCAGCCCACTGACCCCGCATCCCCCACTGTGGCTACCACACCTGAGCCTGTGGGGGCCGATGCTGGGGACAAGAACGCCACCAAAGCAGGCGATGATGAGCCAGAGTACGAGGACGGCCGGGGCTTTGGCATTGGGGAGCTGGTGTGGGGGAAACTGCGGGGCTTCTCCTGGTGGCCAGGCCGCATTGTGTCTTGGTGGATGACGGGCCGGAGCCGAGCAGCTGAAGGCACCCGCTGGGTCATGTGGTTTGGAGACGGCAAATTCTCAGTGGTGTGTGTTGAGAAGCTGATGCCGCTGAGCTCGTTTTGCAGTGCGTTCCACCAGGCCACGTACAACAAGCAGCCCATGTACCGCAAAGCCATCTACGAGGTCCTTCAGGTGGCCAGCAGCCGTGCAGGGAAGCTGTTCCCGGTGTGCCATGACAGCGACGAGAGTGACACTGCCAAGGCCGTGGAGGTGCAGAACAAGCCCATGATTGAATGGGCCCTGGGGGGCTTCCAGCCCTCTGGCCCCAAGGGCCTGGAGCCACCAGAAGAAGAGAAGAATCCCTACAAAGAAGTGTACACAGACATGTGGGTGGAACCTGAGGCAGCTGCCTATGCACCACCTCCACCAGCCAAAAAGCCCCGGAAGAGCACAACAGAGAAGCCCAAGGTCAAGGAGATTATTGATGAGCGCACAAGAGAGCGGCTGGTGTATGAGGTGCGGCAGAAGTGCCGGAACATTGAGGACATCTGCATCTCCTGTGGGAGCCTCAACGTCACCCTGGAACACCCCCTCTTCGTTGGAGGAATGTGCCAAAACTGCAAGAACTGCTTTCTGGAGTGTGCGTATCAGTACGACGACGACGGCTACCAGTCCTACTGCACCATCTGCTGCGGGGGCCGCGAGGTGCTCATGTGCGGAAACAACAACTGCTGCAGGTGCTTTTGCGTGGAGTGTGTGGACCTCTTAGTGGGGCCGGGGGCTGCCCAGGCGGCCATTAAGGAAGACCCCTGGAACTGCTACATGTGCGGGCACAAGGGTACCTATGGGCTGCTGCGGCGGCGAGAGGACTGGCCCTCCCGGCTTCAGATGTTCTTCGCTAATAACCACGACCAGGAATTTGATCCTCCAAAGGTTTACCCACCTGTCCCAGCTGAGAAGAGGAAGCCCATCCGGGTGCTGTCTCTCTTTGATGGAATTGCTACAGGGCTCCTGGTGCTGAAGGACTTGGGCATTCAGGTGGACCGCTACATTGCCTCGGAGGTGTGTGAGGACTCCATCACGGTGGGCATGGTGCGGCACCAGGGGAAGATCATGTACGTCGGGGACGTCCGCAGCGTCACACAGAAGCATATCCAGGAGTGGGGTCCATTTGATCTGGTGATTGGGGGCAGTCCCTGCAATGACCTCTCCATCGTCAACCCTGCTCGCAAGGGCCTCTACGAGGGCACTGGCCGGCTCTTCTTTGAGTTCTACCGCCTCCTGCATGATGCGCGGCCCAAGGAGGGAGATGATCGCCCCTTCTTCTGGCTCTTTGAGAATGTGGTGGCCATGGGCGTTAGTGACAAGAGGGACATCTCACGATTTCTCGAGTCCAACCCTGTGATGATTGATGCCAAAGAAGTGTCAGCTGCACACAGGGCCCGCTACTTCTGGGGTAACCTTCCCGGTATGAACAGGCCGTTGGCATCCACTGTGAATGATAAGCTGGAGCTGCAGGAGTGTCTGGAGCATGGCAGGATAGCCAAGTTCAGCAAAGTGAGGACCATTACTACGAGGTCAAACTCCATAAAGCAGGGCAAAGACCAGCATTTCCCCGTCTTCATGAATGAGAAAGAGGACATCTTATGGTGCACTGAAATGGAAAGGGTATTTGGTTTCCCTGTCCACTATACTGACGTCTCCAACATGAGCCGCTTGGCGAGGCAGAGACTGCTGGGCCGGTCGTGGAGCGTGCCAGTCATCCGCCACCTCTTCGCTCCGCTGAAGGAGTATTTTGCGTGTGTGTAA*

*>Colobus3A*

*ATGCCCTCCAGCGGCCCCGGGGACACCAGCAGCTCTGCTGCAGAGCGGGAGGAGGACCGAAAGGACGGAGAGGAGCAAGAGGAGCCGCGTGGCAAGGAGGAGCGCCAAGAGCCCAGCACCACGGCACGGAAGGTGGGGAGGCCTGGGAGGAAGCGCAAGCACCCCCCGGTGGAAAGCAGTGACACGCCAAAGGACTCTGTGGTGACCTCCAAGTCCCCATCCATGGCCCAGGACTCAGGCCCCTCAGAGCTGTTACCCAACGGGGACTTGGAGAAGCGGAGTGAGCCCCAGCCAGAGGAGGGGAGCCCTGCTGGGGGGCAGAAGGGCGGGGCCCCAGCAGAGGGAGAGGGTGCAGCTGAGACCCCGCCTGAAGCCTCAAGAGCAGTGGAAAATGGCTGCTGCACCCCCAAGGAGGGCCGAGGAGCACCTGCAGAAGCGGGCAAAGAACAGAAGGAGACCAACATCGAATCCATGAAAATGGAGGGCTCCCGGGGCCGGCTGCGGGGTGGCTTGGGCTGGGAGTCCAGCCTCCGCCAGCGGCCCATGCCGAGGCACACCTTCCAGGCGGGGGACCCCTACTACATCAGCAAGCGCAAGCGGGACGAGTGGCTGGCACGCTGGAAAAGGGAGGCTGAGAAGAAAGCCAAGGTCATTGCAGGAATGAATGCTGTGGAAGAAAACCAGGGGCCCGGGGAGTCTCAGAAGGTGGAGGAGGCCAGCCCTCCTGCTGTGCAGCAGCCCACTGACCCCGCATCCCCCACTGTGGCTACCACACCTGAGCCTGTGGGGGCTGATGCTGGGGACAAGAACGCCACCAAAGCAGGCGATGATGAGCCAGAGTACGAGGACGGCCGGGGCTTTGGCATTGGGGAGCTGGTGTGGGGGAAACTGCGGGGCTTCTCCTGGTGGCCAGGCCGCATTGTGTCTTGGTGGATGACGGGCCGGAGCCGAGCAGCTGAAGGCACCCGCTGGGTCATGTGGTTTGGAGACGGCAAATTCTCAGTGGTGTGTGTTGAGAAGCTGATGCCGCTGAGCTCGTTTTGCAGTGCGTTCCACCAGGCCACGTACAACAAGCAGCCCATGTACCGCAAAGCCATCTACGAGGTCCTGCAGGTGGCCAGCAGCCGCGCAGGGAAGCTGTTCCCGGTGTGCCATGACAGCGACGAGAGTGACACTGCCAAGGCCGTGGAGGTGCAGAACAAGCCCATGATTGAATGGGCCCTGGGGGGATTCCAGCCCTCTGGCCCCAAGGGCCTGGAGCCACCAGAAGAAGAGAAGAATCCCTACAAAGAAGTGTACACAGACATGTGGGTGGAACCTGAGGCAGCTGCCTATGCACCACCTCCACCAGCCAAAAAGCCCCGGAAGAGCACAACAGAGAAGCCCAAGGTCAAGGAGATTATTGATGAGCGCACAAGAGAGCGGCTGGTGTATGAGGTGCGGCAGAAGTGCCGGAACATTGAGGACATCTGCATCTCCTGTGGGAGCCTCAACGTCACCCTGGAACACCCCCTCTTCGTTGGAGGAATGTGCCAAAACTGCAAGAACTGCTTTCTGGAGTGTGCGTATCAGTACGACGACGACGGCTACCAGTCCTACTGCACCATCTGCTGCGGGGGCCGCGAGGTGCTCATGTGCGGAAACAACAACTGCTGCAGGTGCTTTTGCGTGGAGTGTGTGGACCTCTTGGTGGGGCCGGGGGCTGCCCAGGCGGCCATTAAGGAAGACCCCTGGAACTGCTACATGTGCGGGCACAAGGGTACCTATGGGCTGCTGCGGCGGCGAGAGGACTGGCCCTCCCGGCTCCAGATGTTCTTCGCTAATAACCACGACCAGGAATTTGATCCTCCAAAGGTTTACCCACCTGTCCCAGCTGAGAAGAGGAAGCCCATCCGGGTGCTGTCTCTCTTTGATGGAATTGCTACAGGGCTCCTGGTGCTGAAGGACTTGGGCATTCAGGTGGACCGCTACATTGCCTCGGAGGTGTGTGAGGACTCCATCACGGTGGGCATGGTGCGGCACCAGGGGAAGATCATGTACGTCGGGGACGTCCGCAGCGTCACACAGAAGCATATCCAGGAGTGGGGCCCATTCGATCTGGTGATTGGGGGCAGTCCCTGCAATGACCTCTCCATCGTCAACCCTGCTCGCAAGGGCCTCTACGAGGGCACTGGCCGGCTCTTCTTTGAGTTCTACCGCCTCCTGCATGATGCGCGGCCCAAGGAGGGAGATGATCGCCCCTTCTTCTGGCTCTTTGAGAATGTGGTGGCCATGGGCGTTAGTGACAAGAGGGACATCTCACGATTTCTCGAGTCCAACCCTGTGATGATTGATGCCAAAGAAGTGTCAGCTGCACACAGGGCCCGCTACTTCTGGGGTAACCTTCCCGGTATGAACAGGCCGTTGGCATCCACTGTGAATGATAAGCTGGAGCTGCAGGAGTGTCTGGAGCATGGCAGGATAGCCAAGTTCAGCAAAGTGAGGACCATTACTACGAGGTCAAACTCCATAAAGCAGGGCAAAGACCAGCATTTCCCCGTCTTCATGAATGAGAAAGAGGACATCTTATGGTGCACTGAAATGGAAAGGGTATTTGGTTTCCCTGTCCACTATACTGACGTCTCCAACATGAGCCGCTTGGCGAGGCAGAGACTGCTGGGCCGGTCGTGGAGCGTGCCAGTCATCCGCCACCTCTTCGCTCCACTGAAGGAGTATTTTGCGTGTGTGTAA*

*>Squirell3A*

*ATGCCCTCCAGCGGCCCCGGGGACACCAGCAGCTCTGCTGCGGAGCGGGAGGAGGACCGAAAGGATGGAGAGGAGCAGGAGGAGCCTCGTGGCAAGGAGGAGCGCCAAGAGCCCAGCACCACGGCCCGGAAGGTGGGGCGGCCTGGGAGGAAGCGCAAGCACCCCCCGGTGGAAAGCAGTGACACGCCAAAGGACCCTGCGGTGACCTCCAAGTCCCCATCCATGGCCCAGGACTCAGGCCCCTCAGAGCTGTTACCCAACGGGGACTTGGAGAAGCGGAGTGAGCCCCAGCCAGAGGAGGGGAGCCCTGCTGGGGGGCAGAAGGGCGGGTCCCCAGCAGAGGGAGAGGGTGCAGCTGAGACCCCGTCAGAAGCGTCAAGAGCAGTGGAGAATGGCTGCTGCACCCCCAAGGAGGGCCGAGGAGCTGCTGCAGAAGAGGGCAAAGAACAGAAGGAGACCAACATCGAAACCATGAAAATGGAGGGCTCCCGGGGCCGGCTGCGGGGTGGCTTGGGCTGGGAGTCCAGCCTCCGTCAGCGGCCCATGCCACGGCTCACCTTCCAGGCGGGGGATCCGTACTACATCAGCAAGCGCAAGCGGGATGAGTGGCTGGCACGCTGGAAAAGGGAGGCTGAGAAGAAAGCCAAGGTTATTGCAGTAATGAATGCTGTGGAAGAAAATCAGGGTTCCGGGGAGTCTCAGAAGGTGGAGGAGGCCAGCCCTCCTGCTGTGCAGCAGCCCACTGACCCCGCATCCCCCACTGTGGCCACCACGCCAGAGCCCGTGGGGGCCGATGCTGGGGACAAGAATGCCACCAAAGCAGGCGATGATGAGCCGGAGTACGAGGACGGCCGGGGCTTTGGCATTGGGGAGCTGGTGTGGGGGAAACTGCGGGGCTTCTCCTGGTGGCCAGGCCGCATTGTGTCTTGGTGGATGACGGGCCGGAGCCGAGCAGCTGAAGGCACCCGCTGGGTCATGTGGTTCGGAGACGGCAAGTTCTCAGTGGTGTGTGTTGAGAAGCTGATGCCGCTGAGCTCGTTTTGCAGTGCGTTCCACCAGGCCACGTACAACAAGCAGCCCATGTACCGCAAAGCCATCTATGAGGTCCTGCAGGTGGCCAGCAGCCGTGCGGGGAAGCTGTTCCCAGTGTGCCACGACAGCGACGAGAGTGACACTGCCAAGGCCGTGGAGGTGCAGAACAAGCAGATGATTGAATGGGCCCTGGGGGGCTTCCAGCCCTCTGGTCCCAAGGGCCTGGAGCCACCAGAAGAGGAGAAGAATCCCTATAAAGAAGTGTACACGGACATGTGGGTGGAACCTGAGGCAGCTGCCTACGCACCACCTCCACCAGCCAAAAAGCCCCGGAAGAGCACAACAGAGAAGCCCAAGGTCAAGGAGATCATCGATGAGCGCACAAGAGAGCGGCTGGTGTACGAGGTGCGGCAGAAGTGCCGGAACATCGAGGACATCTGCATCTCCTGTGGAAGTCTAAACGTCACCCTGGAACACCCCCTCTTCATTGGAGGAATGTGCCAAAACTGCAAGAACTGCTTTCTGGAGTGTGCATACCAGTATGACGACGACGGCTATCAGTCCTACTGCACCATCTGCTGCGGGGGCCGAGAGGTGCTCATGTGCGGGAACAACAACTGCTGCAGGTGCTTTTGCGTGGAATGTGTGGACCTCTTGGTGGGGCCGGGGGCTGCCCAGGCAGCCATTAAGGAAGACCCCTGGAACTGCTACATGTGTGGGCACAAGGGCACCTACGGGCTGCTGCGGCGGCGAGACGACTGGCCCTCCCGGCTCCAGATGTTCTTCGCTAATAACCACGACCAGGAATTTGACCCCCCAAAGGTTTACCCACCTGTCCCAGCTGAGAAGAGGAAGCCCATCCGGGTGCTGTCTCTCTTTGACGGAATTGCTACAGGGCTCCTGGTGCTGAAGGACTTGGGCATTCAGGTGGACCGCTACATTGCCTCGGAGGTGTGTGAGGACTCCATCACGGTGGGCATGGTGCGGCACCAGGGGAAGATCATGTACGTCGGGGACGTCCGCAGCGTCACACAGAAGCATATCCAGGAGTGGGGCCCATTCGATCTTGTGATTGGGGGCAGTCCCTGCAATGACCTCTCCATTGTCAACCCTGCTCGCAAGGGCCTCTACGAGGGCACTGGCCGGCTCTTCTTTGAGTTCTACCGCCTCCTGCATGATGCGCGGCCCAAGGAGGGAGATGACCGCCCCTTCTTCTGGCTCTTTGAGAATGTGGTGGCCATGGGCGTTAGTGACAAGAGGGACATCTCACGATTTCTCGAGTCCAACCCTGTGATGATTGATGCCAAAGAAGTGTCAGCTGCACACAGGGCCCGCTACTTCTGGGGTAACCTTCCCGGTATGAACAGGCCGTTGGCATCCACTGTGAATGATAAGCTGGAGCTGCAGGAGTGTCTGGAGCATGGAAGGATAGCCAAGTTCAGCAAAGTGAGGACCATTACTACGAGGTCAAACTCCATAAAGCAAGGCAAAGACCAGCATTTCCCCGTCTTCATGAATGAGAAAGAGGACATCCTATGGTGCACTGAAATGGAAAGGGTATTTGGTTTCCCTGTCCACTATACTGACGTCTCCAACATGAGCCGCTTGGCGAGGCAGAGACTGCTGGGCCGGTCATGGAGCGTGCCAGTCATCCGCCACCTCTTCGCTCCGCTGAAGGAGTATTTTGCGTGTGTGTAA*

*>Sapajou3A*

*ATGCCCTCCAGCGGCCCCGGGGACACCAGCAGCTCTGCTGCGGAGCGGGAGGAGGACCGAAAGGATGGAGAGGAGCAGGAGGAGCCTCGTGGCAAGGAGGAGCGCCAAGAGCCCAGCACCACGGCCCGGAAGGTGGGGCGGCCTGGGAGGAAGCGCAAGCACCCCCCGGTGGAAAGTAGTGACACGCCAAAGGACGCTGCGGTAACCTCCAAGTCCCCATCCATGGCCCAGGACTCAGGCCCCTCAGAGCTGTTACCCAACGGGGACTTGGAGAAGCGAAGTGAGCCCCAGCCAGAGGAGGGGAGCCCTGCTGGGGGGCAGAAGGGCGGGTCCCCAGCAGAGGGAGAGGGTGCAGCTGAGACCCCGCCGGAAGCCTCTAGAGCAGTGGAGAATGGCTGCTGCACCCCCAAGGAGGGCCGAGGAGCCGCTGCAGAAGAGGGCAAAGAACAGAAGGAGACCAACATCGAAACCATGAAAATGGAGGGCTCCCGGGGCCGGCTGCGGGGTGGCTTGGGCTGGGAGTCCAGCCTCCGTCAGCGGCCCATGCCACGGCTCACCTTCCAGGCGGGGGATCCGTACTACATCAGCAAGCGTAAGCGGGACGAGTGGCTGGCACGCTGGAAAAGAGAGGCTGAGAAGAAAGCCAAGGTTATTGCAGTAATGAATGCTGTGGAAGAAAACCAGGGTTCCGGGGAGTCTCAGAAGGTGGAGGAGGCCAGCCCTCCTGCTGTGCAGCAGCCCACTGACCCCGCATCCCCCACTGTGGCCACCACGCCAGAGCCCGTGGGAGCCGATGCTGGGGACAAGAATGCCACCAAAGCAGGCGATGATGAGCCGGAGTACGAGGACGGCCGGGGCTTTGGCATTGGGGAGCTGGTGTGGGGGAAACTGCGGGGCTTCTCCTGGTGGCCAGGCCGCATTGTGTCTTGGTGGATGACGGGCCGGAGCCGAGCAGCTGAAGGCACCCGCTGGGTCATGTGGTTCGGAGACGGCAAGTTCTCAGTGGTGTGTGTTGAGAAGCTGATGCCGCTGAGCTCGTTTTGCAGTGCGTTCCACCAGGCCACGTACAACAAGCAGCCCATGTATCGCAAAGCAATCTATGAGGTCCTGCAGGTGGCCAGCAGCCGTGCGGGGAAGCTGTTCCCAGTGTGCCACGACAGCGACGAGAGTGACACTGCCAAGGCCGTGGAGGTGCAGAACAAGCAGATGATTGAATGGGCCCTGGGGGGCTTCCAGCCCTCTGGTCCCAAGGGCCTGGAGCCACCAGAAGAGGAGAAGAATCCCTACAAAGAAGTGTACACGGACATGTGGGTGGAACCTGAGGCAGCTGCCTACGCACCACCTCCACCAGCCAAAAAGCCCCGGAAGAGCACAACGGAGAAGCCCAAGGTCAAGGAGATCATCGATGAGCGCACAAGAGAGCGGCTGGTGTACGAGGTGCGGCAGAAGTGCCGGAACATTGAGGACATCTGCATCTCCTGTGGGAGTCTCAACGTTACCCTGGAACACCCCCTCTTCATTGGAGGAATGTGCCAAAACTGCAAGAACTGCTTTCTAGAGTGTGCATACCAGTATGACGACGACGGCTATCAGTCCTACTGCACCATCTGCTGCGGGGGCCGAGAGGTGCTCATGTGTGGGAACAACAATTGCTGCAGGTGCTTTTGTGTGGAATGTGTGGACCTCTTGGTGGGGCCGGGGGCTGCCCAGGCAGCCATTAAGGAAGACCCCTGGAACTGCTACATGTGTGGGCACAAGGGCACCTACGGGCTGCTGCGGCGGCGAGACGACTGGCCCTCCCGGCTCCAGATGTTCTTCGCTAATAACCACGACCAGGAATTTGACCCTCCAAAGGTTTACCCACCTGTCCCAGCTGAGAAGAGGAAGCCCATCCGGGTGCTGTCTCTCTTTGACGGAATTGCTACAGGGCTCCTGGTGCTGAAGGACTTGGGCATTCAGGTGGACCGCTACATTGCCTCGGAGGTGTGTGAGGACTCCATCACGGTGGGCATGGTGCGGCACCAGGGGAAGATCATGTACGTCGGGGACGTCCGCAGCGTCACACAGAAGCATATCCAGGAGTGGGGCCCATTCGATCTTGTGATTGGGGGCAGTCCCTGCAATGACCTCTCCATTGTCAACCCTGCTCGCAAGGGCCTCTACGAGGGCACTGGCCGGCTCTTCTTTGAGTTCTACCGCCTCCTGCATGATGCGCGGCCCAAGGAGGGAGATGACCGCCCCTTCTTCTGGCTCTTTGAGAATGTGGTGGCCATGGGCGTTAGTGACAAGAGGGACATCTCGCGATTTCTCGAGTCCAACCCTGTGATGATTGATGCCAAAGAAGTGTCAGCTGCACACAGGGCCCGCTACTTCTGGGGTAACCTTCCCGGTATGAACAGGCCGTTGGCATCCACTGTGAATGATAAGCTGGAGCTGCAGGAGTGTCTGGAGCATGGAAGGATAGCCAAGTTCAGCAAAGTGAGGACCATTACTACGAGGTCAAACTCCATAAAGCAAGGCAAAGACCAGCATTTCCCCGTCTTCATGAATGAGAAAGAGGACATCCTATGGTGCACTGAAATGGAAAGGGTATTTGGTTTCCCTGTCCACTATACTGACGTCTCCAACATGAGCCGCTTGGCAAGGCAGAGACTGCTGGGCCGGTCGTGGAGCGTGCCAGTCATCCGCCACCTCTTCGCTCCGCTGAAGGAGTATTTTGCGTGTGTGTAA*

*>Marmoset3A*

*ATGCCCTCCAGCGGCCCCGGGGACATCAGCAGCTCTGCTGTGGAGCGGGAGGAGGACCGAAAGGATGGAGAGGAGCAGGAGGAGCCTCGTGGCAAAGAGGAGCGCCAAGAGCCCAGCACCACGGCCCGGAAGGTGGGGCGGCCTGGGAGGAAGCGCAAGCACCCCCCGGTGGAAAGCAGTGACACACCAAAGGACCCTGCGGTGACCTCCAAGACCCCATCCATGGCCCAGGACTCAGGCCCCTCAGAGCTGTTACCCAACGGGGACTTGGAGAAGCGGAGTGAGCCACAGCCAGAGGAGGGGACCCCTGCTGGGGGGCAGAAGGGCGGGTCCCCAGCAGAGGGAGAGGGTGCAGCTGAGACCCCGCCAGAAGCCTCCAGAGCAGTGGAGAATGGCTGCTGCACCCCCAAGGAGGGACGAGGAGCCCCTGCAGAAGAGGGCAAAGAACAGAAGGAGACCAACATCGAAACCATGAAAATGGAGGGCTCCCGGGGCCGGCTGCGGGGTGGCTTGGGCTGGGAGTCCAGCCTCCGTCAGCGGCCCATGCCACGGCTCACCTTCCAGGCAGGGGATCCATACTACATCAGCAAGCGCAAGCGGGACGAGTGGCTGGCACGCTGGAAAAGGGAGGCTGAGAAGAAAGCCAAGGTCATTGCAGTAATGAATGCTGTGGAAGAAAACCAGGGTTCTGGGGAGTCTCAGAAGGTGGAGGAGGCCAGCCCTCCTGCTGTGCAGCAGCCCACTGACCCTGCATCCCCCACTGTGGCCACCACACCAGAGCCCGTGGGGGCTGATGCTGGGGACAAGAATGCCACCAAAGCAGGCGATGATGAGCCAGAGTACGAGGACGGCCGGGGCTTTGGCATTGGGGAGCTGGTGTGGGGGAAACTGCGGGGCTTCTCCTGGTGGCCAGGCCGCATTGTGTCTTGGTGGATGACGGGCCGGAGCCGAGCAGCTGAAGGCACCCGCTGGGTCATGTGGTTCGGAGACGGCAAGTTCTCAGTGGTGTGTGTTGAGAAGCTGATGCCGCTGAGCTCGTTTTGCAGTGCGTTCCACCAGGCCACATACAACAAGCAGCCCATGTACCGCAAAGCCATCTATGAGGTCCTGCAGGTGGCCAGCAGCCGTGCGGGGAAGCTGTTCCCAGTGTGCCACGACAGCGACGAGAGTGACACTGCCAAGGCTGTGGAGGTGCAGAACAAGCAGATGATTGAATGGGCCCTGGGGGGCTTCCAGCCCTCTGGTCCCAAGGGCCTGGAGCCACCAGAAGAGGAAAAGAATCCCTACAAAGAAGTGTACACGGACATGTGGGTAGAACCTGAGGCAGCTGCCTACGCACCACCTCCACCAGCCAAAAAGCCCCGGAAGAGCACAACAGAGAAGCCCAAGGTCAAGGAGATCATCGATGAGCGCACAAGAGAGCGGCTGGTGTACGAGGTGCGGCAGAAGTGCCGGAACATCGAGGACATCTGCATCTCCTGTGGGAGCCTCAACGTCACCCTGGAACACCCCCTCTTCATTGGAGGAATGTGCCAAAACTGCAAGAACTGCTTTCTGGAGTGTGCATACCAGTACGACGACGATGGCTATCAGTCCTACTGCACCATCTGCTGCGGGGGCCGAGAGGTGCTCATGTGTGGGAACAACAACTGCTGCAGGTGCTTTTGTGTGGAATGTGTGGACCTCTTGGTGGGGCCGGGGGCTGCCCAGGCAGCCATTAAGGAAGACCCCTGGAACTGCTACATGTGTGGGCACAAGGGCACCTACGGGCTGCTGCGGCGGCGAGACGACTGGCCCTCCCGGCTCCAGATGTTCTTTGCTAATAACCACGACCAGGAATTTGACCCTCCAAAGGTTTACCCACCTGTCCCAGCTGAGAAGAGGAAGCCCATCCGGGTGCTGTCTCTCTTTGATGGAATTGCTACAGGGCTCCTGGTGCTGAAGGACTTGGGCATTCAGGTGGACCGCTACATTGCCTCGGAGGTGTGTGAGGACTCCATCACGGTGGGCATGGTGCGGCACCAGGGGAAGATCATGTACGTCGGGGACGTCCGCAGCGTCACACAGAAGCATATCCAGGAGTGGGGCCCATTCGATCTTGTGATTGGGGGCAGTCCCTGCAATGACCTCTCCATTGTCAACCCTGCTCGCAAGGGCCTCTACGAGGGCACTGGCCGGCTCTTCTTTGAGTTCTACCGCCTCCTGCATGATGCGCGGCCCAAGGAGGGAGATGACCGCCCCTTCTTCTGGCTCTTTGAGAATGTGGTGGCCATGGGCGTTAGTGACAAGAGGGACATCTCGCGATTTCTCGAGTCCAACCCTGTGATGATTGATGCCAAAGAAGTGTCAGCTGCACACAGGGCCCGCTACTTCTGGGGTAACCTTCCCGGTATGAACAGGCCGTTGGCATCCACTGTGAATGATAAGCTGGAGCTGCAGGAGTGTCTGGAGCATGGAAGGATAGCCAAGTTCAGCAAAGTGAGGACCATTACTACAAGGTCAAACTCCATAAAGCAAGGCAAAGACCAGCATTTCCCCGTCTTCATGAATGAGAAAGAGGACATCCTATGGTGCACTGAAATGGAAAGGGTATTTGGTTTCCCTGTCCACTATACTGACGTCTCTAACATGAGCCGCTTGGCGAGGCAGAGACTGCTGGGCCGGTCGTGGAGCGTGCCAGTCATCCGCCACCTCTTCGCTCCGCTGAAGGAGTATTTTGCGTGTGTGTAA*

*>OwlMonkey3A*

*ATGCCCTCCAGCGGCCCCGGGGACACCAGCAGCTCTGCTGCGGAGCGGGAGGAGGACCGAAAGGATGGAGAGGAGCAGGAGGAGCCTCGTGGCAAGGAGGAGCGCCAAGAGCCCAGCACCACGGCCCGGAAGGTGGGGCGGCCTGGGAGGAAGCGCAAGCACCCCCCGGTGGAAAGCAGTGACACGCCAAAGGACCCTGCGGTGACCTCCAAGTCCCCATCCATGGCCCAGGACTCAGGCCTCTCAGAGCTGTTACCCAACGGGGACTTGGAGAAGCGGAGTGAGCCCCAACCAGAGGAGGGGAGCCCTGCTGGGGGGCAGAAGGGCGGGTCCCCAGCAGAGGGAGAGGGTGCAGCTGAGACTCCGCCGGAAGCCTCCAGAGCAGTGGAGAATGGCTGCTGCACCCCCAAGGAGGGCCGAGGAGCTGCTGCAGAA---GGCAAAGAACAGAAGGAGACCAACATCGAAACCATGAAAATGGAGGGCTCCCGGGGCCGGTTGCGGGGTGGCTTGGGCTGGGAGTCCAGCCTCCGTCAGCGGCCCATGCCACGGCTCACCTTCCAGGCGGGGGATCCGTACTACATCAGCAAGCGCAAGCGGGACGAGTGGCTGGCACGCTGGAAAAGGGAGGCTGAGAAGAAAGCCAAGGTCATTGCAGTAATGAATGCTGTGGAAGAAAACCAGGGTTCCGGGGAGTCTCAGAAGGTGGAGGAGGCCAGCCCTCCTGCTGTGCAGCAGCCCACTGACCCCGCATCCCCCACTGTGGCCACCACGCCAGAGCCCGTGGGGGCCGATACTGGGGACAAGAATGCCACCAAAGCAGGCGATGATGAGCCGGAGTACGAGGACGGCCGGGGCTTTGGCATTGGGGAGCTGGTGTGGGGGAAACTGCGGGGCTTCTCCTGGTGGCCAGGCCGCATTGTGTCTTGGTGGATGACGGGCCGGAGCCGAGCAGCTGAAGGCACCCGCTGGGTCATGTGGTTCGGAGACGGCAAGTTCTCAGTGGTGTGTGTTGAGAAGCTGATGCCGCTGAGCTCGTTTTGCAGTGCGTTCCACCAGGCCACGTACAACAAGCAGCCCATGTACCGCAAAGCCATCTATGAGGTCCTGCAGGTGGCCAGCAGCCGTGCAGGGAAGCTGTTCCCAGTGTGCCATGACAGCGACGAGAGTGACACTGCCAAGGCCGTGGAGGTGCAGAACAAGCAGATGATTGAATGGGCCCTGGGGGGCTTCCAGCCTTCTGGTCCCAAGGGCCTGGAGCCACCAGAAGAGGAGAAGAATCCCTACAAAGAAGTGTACACGGACATGTGGGTGGAACCTGAGGCAGCTGCCTACGCACCACCTCCACCAGCCAAAAAGCCCCGGAAGAGCACAACGGAGAAGCCCAAGGTCAAGGAGATCATCGATGAGCGCACGAGAGAGCGGCTGGTATACGAGGTGCGGCAGAAGTGCCGGAACATCGAGGACATCTGCATCTCCTGTGGGAGCCTCAACGTCACCCTGGAACACCCCCTCTTCATTGGAGGAATGTGCCAAAACTGCAAGAACTGCTTTCTGGAGTGTGCATACCAGTACGACGACGACGGCTATCAGTCCTACTGCACCATCTGCTGCGGGGGCCGAGAGGTGCTCATGTGCGGGAACAACAACTGCTGCAGGTGCTTTTGCGTGGAATGTGTGGACCTCTTGGTGGGGCCAGGGGCTGCCCAGGCAGCCATTAAGGAAGACCCCTGGAACTGCTATATGTGTGGGCACAAGGGCACCTACGGGCTGCTGCGGCGGCGAGACGACTGGCCCTCCCGGCTCCAGATGTTCTTCGCTAATAACCATGACCAGGAATTTGACCCTCCAAAGGTTTACCCACCTGTCCCAGCTGAGAAGAGGAAGCCCATCCGGGTGCTATCTCTCTTTGACGGAATTGCTACAGGGCTCCTGGTGCTGAAGGACTTGGGCATTCAGGTGGACCGCTACATTGCCTCGGAGGTGTGTGAGGACTCCATCACAGTGGGCATGGTGCGGCACCAAGGGAAGATCATGTACGTCGGGGACGTCCGCAGCGTCACACAGAAGCATATCCAGGAGTGGGGCCCATTCGATCTTGTGATTGGGGGCAGTCCCTGCAATGACCTCTCCATTGTCAACCCTGCTCGCAAGGGCCTCTACGAGGGCACTGGCCGGCTCTTCTTTGAGTTCTACCGCCTCCTGCATGATGCGCGGCCCAAGGAGGGAGATGACCGCCCCTTCTTCTGGCTCTTTGAGAATGTGGTGGCCATGGGCGTTAGTGACAAGAGGGACATCTCGCGATTTCTCGAGTCCAACCCTGTGATGATTGATGCCAAAGAAGTGTCAGCTGCACACAGGGCCCGCTACTTCTGGGGTAACCTTCCCGGTATGAACAGGCCGTTGGCATCCACTGTGAATGATAAGCTGGAGCTGCAGGAGTGTCTGGAGCATGGAAGGATAGCCAAGTTCAGCAAAGTGAGGACCATTACTACGAGGTCAAACTCCATAAAGCAGGGCAAAGACCAGCATTTCCCCGTCTTCATGAATGAGAAAGAGGACATCCTATGGTGCACTGAAATGGAAAGGGTATTTGGTTTCCCTGTCCACTATACTGACGTCTCCAACATGAGCCGCTTGGCGAGGCAGAGACTGCTGGGCCGGTCGTGGAGCGTGCCAGTCATCCGCCACCTCTTCGCTCCGCTGAAGGAGTATTTTGCGTGTGTGTAA*

*>GrayMouse3A*

*ATGCCCGCCAGCGGCCCCGGGGACACCAGCAGCTCTGCTGCGGACCGGGAGGAGGACCGAAAGGACGGAGAGGAGCAGGAGGAGCCTCGTGGCAAGGAGGAGCGCCAGGAGCCCAGCACCACGGCTCGGAAGGTGGGCAGGCCTGGGCGGAAGCGCAAGCACCCCCCGGTGGAAAGCAGCGACACGCCAAAGGACACTGCAGTGACCTCCAAGTCCCCATCCACGGCCCAGGACTCAGGCCCCTCCGAACTGTTACCCAATGGGGACTTGGAGAAGCGGAGTGAGCCCCAGCCGGAAGAGGGGAGCCCTGCTGGGGGGCAGAAGGGCGGGGCCCCAGCAGAGGGAGAGGGTGCAGCTGAGACCCCGCCTGAAGCCTCCAGAGCAGTGGAAAATGGCTGCTGCACCCCCAAGGAGGGCCGAGGAGCCCCTGCAGAAGAGGGCAAAGAACAGAAGGAGACCAACATTGAATCCATGAAAATGGAGGGCTCCCGGGGCCGGCTGCGGGGTGGCTTGGGCTGGGAGTCCAGCCTCCGCCAGCGGCCCATGCCGCGGCTCACCTTCCAGGCGGGGGACCCCTACTACATCAGCAAGCGCAAGCGGGACGAGTGGCTGGCACGCTGGAAAAGGGAGGCTGAGAAGAAAGCCAAGGTAATTGCAGTAATGAATGCTGTGGAAGAGAACCAGGGGTCTGGGGAGTCTCAGAAGGTGGAGGAGGCCAGCCCTCCCGCTGTGCAGCAGCCCACTGACCCCGCGTCCCCCACGGTGGCCACCACGCCTGAACCCGTGGGGGCTGATGCCGGCGACAAGAATGCCACCAAAGCAGCCGATGACGAACCGGAGTACGAGGACGGCCGGGGCTTTGGCATTGGGGAGCTGGTGTGGGGGAAACTGCGGGGCTTCTCCTGGTGGCCAGGCCGCATTGTGTCTTGGTGGATGACGGGCCGGAGCCGAGCAGCTGAAGGCACCCGCTGGGTCATGTGGTTCGGAGACGGCAAGTTCTCAGTGGTGTGTGTGGAGAAGCTGATGCCACTGAGCTCGTTTTGCAGCGCGTTCCACCAGGCCACCTACAACAAGCAGCCCATGTACCGCAAAGCCATCTACGAAGTCCTGCAGGTGGCCAGCAGCCGCTCCGGGAAGCTGTTCCCAGCGTGCCATGACAGCGACGAGAGTGACACTGCCAAGGCCGTGGAGGTGCAGAACAAGCAAATGATTGAATGGGCCCTTGGGGGGTTCCAGCCCTCCGGCCCCAAGGGCCTGGAGCCACCAGAAGAGGAAAAGAATCCCTACAAAGAAGTTTACACAGACATGTGGGTTGAACCTGAGGCAGCTGCCTATGCACCACCCCCACCAGCCAAAAAGCCCCGAAAGAGCACAACAGAGAAGCCCAAGGTCAAGGAGATTATCGATGAACGCACAAGAGAGCGGCTGGTGTACGAGGTGCGGCAGAAGTGCCGGAATATTGAAGATATTTGCATTTCTTGTGGGAGCCTCAACGTCACCCTGGAACACCCTCTCTTCATTGGAGGAATGTGCCAAAACTGCAAGAACTGCTTCCTGGAGTGTGCATACCAGTATGATGACGATGGCTATCAGTCCTACTGCACCATCTGCTGTGGGGGGCGTGAGGTGCTCATGTGCGGGAACAACAACTGCTGCAGGTGCTTTTGCGTGGAGTGTGTGGACCTCTTGGTGGGGCCGGGGGCTGCCCAGGCGGCCATTAAGGAAGACCCCTGGAATTGCTACATGTGTGGGCACAAGGGTACCTATGGGCTGCTGCGGCGGCGGGACGACTGGCCCTCTAGGCTCCAGATGTTCTTCGCCAATAACCACGACCAGGAATTTGACCCTCCGAAGGTTTATCCACCTGTCCCAGCCGAGAAGAGGAAGCCTATCCGGGTGCTGTCTCTCTTTGACGGAATTGCTACAGGGCTCCTGGTGCTGAAGGACCTGGGCATCCAGGTGGACCGCTACATCGCCTCGGAGGTGTGCGAGGACTCCATCACGGTGGGCATGGTGCGGCACCAGGGGAAGATCATGTACGTCGGGGACGTCCGCAGCGTCACACAGAAGCATATCCAGGAGTGGGGCCCATTCGATCTGGTGATTGGGGGCAGTCCCTGCAACGATCTCTCCATCGTCAACCCTGCCCGCAAGGGACTCTACGAGGGCACTGGCCGGCTCTTCTTTGAGTTCTACCGCCTCCTGCATGATGCGCGGCCCAAGGAGGGAGATGATCGCCCCTTCTTCTGGCTCTTTGAGAATGTGGTGGCCATGGGCGTTAGTGACAAGAGGGACATCTCGCGATTTCTCGAGTCCAACCCTGTGATGATTGATGCCAAAGAAGTGTCAGCTGCACACAGGGCCCGCTACTTTTGGGGTAACCTTCCCGGTATGAACAGGCCATTGGCGTCCACTGTGAATGATAAGCTGGAGCTGCAGGAGTGTCTGGAGCACGGCAGGATAGCCAAGTTCAGCAAAGTAAGGACCATTACTACTAGGTCGAACTCCATAAAGCAGGGCAAAGACCAGCATTTTCCCGTCTTCATGAACGAGAAAGAGGACATCTTATGGTGCACTGAAATGGAAAGGGTGTTTGGCTTCCCCGTCCACTATACCGACGTCTCCAACATGAGCCGCTTGGCCAGGCAGAGACTTCTGGGCCGGTCGTGGAGCGTGCCAGTCATCCGCCACCTCTTCGCTCCGCTGAAGGAATATTTTGCTTGTGTGTAA*

*>Sifaka3A*

*ATGCCCTCCAGTGGCCCCGGGGACACCAGCAGCTCTGCTGCAGAGCGGGAGGAGGACCGAAAGGACGCAGAGGAGCAGGAGGAGCCTCGTGGCAAGGAGGAGCGCCAGGAGCCCAGCACCACAGCACGGAAGGTGGGCAGGCCTGGGCGGAAGCGCAAGCACCCCCCGGTGGAAAGCAGCGACACGCCGAAGGACACTGCAGTGACCTCCAAGTCCCCATCCACAGCCCAGGACTCAGGCTGCTCCGAACTGTTGCCCAATGGGGACTTGGAGAAGCGGAGTGAGCCCCAGCCAGAAGAGGGGAGCCCTGCTGGGGGGCAGAAGGGCGGGGCCCCAGCAGAGGGAGAGGGTGCAGCTGAGACCCCGCCTGAAGCCTCCAGAGCAGTGGAAAATGGCTGCTGCACCCCCAAGGAGGGCCGAGGAGCCCCTGCAGAAGAGGGCAAAGAACAGAAGGAGACCAACATTGAATCCATGAAAATGGAGGGCTCCCGGGGCCGGCTGCGGGGTGGCTTGGGCTGGGAGTCCAGCCTCCGCCAGCGGCCCATGCCGCGGCTCACCTTCCAGGCGGGGGACCCCTACTACATCAGCAAGCGCAAGCGGGACGAGTGGCTGGCACGCTGGAAAAGGGAGGCTGAGAAGAAAGCCAAGGTAATTGCAGTAATGAATGCTGTGGAAGAGAACCAGGGGTCTGGGGAGTCTCAGAAGGTGGAGGAGGCCAGCCCCCCTGCTGTGCAGCAGCCCACTGACCCCGCATCGCCCACCGTGGCCACCACCCCCGAACCTCTGGGGGCTGATGCCGGGGACAAGAATGCCACCAAAGCAGCCGACGATGAACCGGAGTACGAGGACGGCCGGGGCTTTGGCATTGGGGAGCTGGTGTGGGGGAAACTGCGGGGCTTCTCCTGGTGGCCAGGCCGCATTGTGTCTTGGTGGATGACGGGCCGGAGCCGAGCAGCCGAAGGCACCCGCTGGGTCATGTGGTTCGGAGACGGCAAGTTCTCAGTGGTGTGCGTGGAGAAGCTGATGCCGCTGAGCTCGTTTTGCAGCGCGTTCCACCAGGCCACCTACAACAAGCAGCCCATGTACCGCAAAGCCATCTACGAAGTCCTGCAGGTGGCCAGCAGCCGCTCCGGGAAACTGTTCCCAGCGTGCCATGACAGCGACGAGAGTGACACTGCCAAGGCCGTGGAGGTGCAGAACAAGCAAATGATTGAATGGGCCCTTGGGGGGTTCCAGCCCTCTGGCCCCAAGGGCCTGGAGCCACCAGAAGAGGAGAAGAATCCTTACAAAGAAGTTTACACAGACATGTGGGTTGAACCTGAGGCAGCTGCCTATGCACCACCCCCACCAGCCAAAAAGCCCCGAAAGAGCACAACAGAGAAGCCCAAGGTCAAGGAGATTATCGATGAACGCACAAGAGAGCGGCTGGTGTACGAAGTGCGGCAGAAGTGCCGGAATATTGAAGACATTTGCATTTCTTGTGGGAGCCTCAACGTCACCCTGGAACACCCTCTCTTCACTGGAGGAATGTGCCAAAACTGCAAGAACTGCTTCCTGGAGTGCGCATACCAGTACGATGACGATGGCTATCAGTCCTACTGCACCATCTGCTGTGGGGGGCGTGAGGTGCTCATGTGCGGGAACAACAACTGCTGCAGGTGCTTTTGCGTGGAGTGTGTGGACCTCTTGGTGGGGCCGGGGGCTGCCCAGGCGGCCATTAAGGAAGACCCCTGGAATTGCTACATGTGTGGGCACAAGGGTACCTATGGGCTGCTGCGGCGGCGGGACGACTGGCCCTCTAGGCTCCAGATGTTCTTCGCCAATAACCACGACCAGGAATTTGACCCTCCGAAGGTTTATCCACCTGTCCCAGCTGAGAAGAGGAAGCCCATCCGGGTGCTGTCTCTCTTTGACGGAATTGCTACAGGGCTCCTGGTGCTGAAGGACCTGGGCATCCAGGTGGACCGCTACATTGCCTCGGAGGTGTGTGAGGACTCCATCACAGTGGGCATGGTGCGGCATCAGGGGAAGATCATGTACGTCGGGGACGTCCGCAGCGTCACACAGAAGCATATCCAGGAATGGGGCCCATTCGATCTGGTGATTGGGGGCAGTCCCTGCAATGATCTCTCCATCGTCAACCCTGCCCGCAAGGGACTCTACGAGGGCACTGGCCGGCTCTTCTTTGAGTTCTACCGCCTCCTGCATGATGCGCGGCCCAAGGAGGGAGATGATCGCCCCTTCTTCTGGCTCTTTGAGAATGTGGTGGCCATGGGCGTTAGTGACAAGAGGGACATCTCGCGATTTCTCGAGTCCAACCCTGTGATGATTGATGCCAAAGAAGTGTCAGCTGCACACAGGGCCCGCTACTTCTGGGGTAACCTTCCCGGTATGAACAGGCCGTTGGCGTCCACTGTGAATGATAAGCTGGAGCTGCAGGAGTGTCTGGAGCACGGCAGGATAGCCAAGTTCAGCAAAGTGAGGACCATTACTACTAGGTCGAACTCCATAAAGCAGGGCAAAGACCAGCATTTCCCCGTCTTCATGAACGAGAAAGAGGACATCTTATGGTGCACTGAAATGGAAAGGGTGTTTGGCTTCCCCGTCCACTATACCGACGTCTCCAACATGAGCCGCTTGGCCAGGCAGAGACTTCTGGGCCGGTCGTGGAGCGTGCCAGTCATCCGCCACCTCTTCGCTCCACTGAAGGAATATTTTGCTTGTGTGTAA*

*>BushBaby3A*

*ATGCCCTCCAGCGGCCCCGGGGACACCAGCAGCTCTGCTGTGGAGCGGGAGGAGGACCGAAAGGATGGAGAGGAGCAGGAGGAGTCTCGATCCAAGGAGGAGCGACAGGAGCCCAGCACCACGGCCCGAAAAGTGGGGAGGCCTGGGCGGAAGCGCAAGCACCCTCCGGTGGAAAGTAGCGACACACCGAAAGACCCTGCAGTGACCTCCAAGACCCCATCCACGGCCCAGGACTCAGGCCCCTCAGAGCTGTTACCCAATGGGGACTTGGAGAAGCGGAGTGAGCCCCAGCCAGAAGAGGGGAGCCCTGCTGGGGGACAGAAGGGCGGGGCCCCAGCAGAAGGAGAAAGTGCAGCTAACACCCCGTCCGAAGCCTCCAGAGCAGTGGAAAATGGCTGTTGCACCCCCAAGGAGGGCCGAGGAGCCTCT---GAAGAGGGCAAAGAACAGAAGGAGACCAACATTGAAACCATGAAAATGGAGGGCGCTCGGGGCCGGCTGCGGGGTGGCTTGGGCTGGGAGTCCAGCCTCCGCCAGCGGCCCATGCCGCGGCTCACCTTCCAGGCGGGGGACCCCTACTACATCAGCAAGCGCAAGCGGGACGAATGGCTGGCACGCTGGAAAAGGGAGGCTGAGAAGAAAGCCAAGGTAATTGCAGTAATGAACGCTGTGGAAGAGAACCAGGGGGCTGGGGAGTCTCAGAAGGTGGAGGAGGCTAGCCCTCCCGCTGTGCAGCAGCCCACTGACCCCGCATCCCCCACTGTGGCCACCACGCCTGAACCCGTGGGTGCTGATGCAGGGGACAAGAACACCACCAAAGCAGCCGATGATGAACCAGAGTACGAGGACGGCCGGGGCTTTGGCATTGGGGAGCTGGTGTGGGGGAAACTGCGGGGCTTCTCCTGGTGGCCAGGCCGCATTGTGTCTTGGTGGATGACGGGTCGGAGCCGAGCAGCTGAAGGCACCCGCTGGGTCATGTGGTTCGGAGACGGCAAGTTCTCAGTGGTGTGTGTGGAGAAGCTGATGCCACTGAGTTCGTTCTGTAGTGCGTTCCACCAGGCCACCTACAACAAGCAGCCCATGTACCGCAAAGCCATCTACGAAGTCCTGCAGGTGGCCAGCAGCCGCTCAGGGAAACTGTTCCCAGCGTGTCATGACAGTGATGAGAGTGACACTGCCAAGGCTGTGGAGGTGCAGAACAAGCAAATGATTGAATGGGCCCTCGGGGGGTTCCAGCCCTCTGGCCCCAAGGGCCTGGAGCCACCAGAAGAGGAAAAGAATCCCTACAAAGAAGTTTACACAGACATGTGGGTTGAACCTGAGACAGCTGCCTATGCGCCACCCCCACCAGCCAAAAAGCCCCGAAAGAGCACAACAGAGAAGCCCAAGGTCAAGGAGATTATTGATGAACGCACAAGAGAGCGGCTGGTGTATGAGGTGCGGCAGAAGTGCCGGAACATCGAAGACATCTGCATTTCTTGTGGAAGCCTCAATGTCACCTTGGAACACCCTCTCTTCATTGGAGGAATGTGCCAAAACTGCAAGAATTGCTTCCTGGAGTGTGCGTACCAGTACGACGACGACGGCTACCAGTCCTACTGCACCATCTGCTGTGGGGGGCGCGAGGTGCTCATGTGCGGGAACAACAACTGCTGTAGGTGCTTTTGTGTGGAGTGTGTGGACCTCTTGGTGGGGCCAGGGGCTGCCCAGGCGGCCATTAAGGAAGACCCCTGGAATTGCTACATGTGTGGGCACAAGGGTACCTATGGGCTGCTGCGGCGGCGGGACGACTGGCCCTCTAGGCTCCAGATGTTCTTCGCCAATAACCACGACCAGGAATTTGACCCTCCGAAGGTTTACCCACCTGTCCCAGCTGAGAAGAGGAAGCCCATCCGGGTGCTATCTCTCTTTGACGGAATTGCTACAGGGCTCCTGGTGCTGAAGGACTTGGGGATCCAGGTGGACCGCTACATTGCATCGGAGGTGTGTGAAGACTCCATCACAGTGGGCATGGTGCGGCACCAGGGGAAGATCATGTACGTCGGGGACGTCCGCAGCGTCACACAGAAGCATATCCAGGAGTGGGGCCCATTCGATCTGGTGATTGGGGGCAGTCCCTGCAATGATCTCTCCATCGTCAACCCTGCCCGAAAGGGACTCTATGAGGGCACTGGCCGGCTCTTCTTTGAGTTCTACCGCCTCCTGCATGATGCGCGGCCCAAGGAGGGAGATGATCGCCCCTTCTTCTGGCTCTTTGAGAATGTGGTGGCCATGGGCGTTAGTGACAAGAGGGACATCTCGCGATTTCTCGAGTCCAACCCTGTGATGATTGATGCCAAAGAAGTGTCAGCTGCACACAGGGCTCGCTACTTCTGGGGTAACCTTCCTGGTATGAACAGGCCATTGGCATCCACTGTGAATGATAAGCTGGAGCTGCAGGAGTGTCTGGAGCATGGCAGGATAGCCAAGTTCAGCAAAGTGCGGACCATTACTACTAGGTCAAACTCCATAAAGCAGGGCAAAGACCAGCATTTCCCCGTCTTCATGAATGAGAAAGAGGACATCTTATGGTGCACTGAAATGGAAAGAGTGTTTGGCTTCCCCGTCCACTATACCGACGTCTCCAACATGAGCCGCTTGGCGAGGCAGAGACTTCTGGGCCGGTCATGGAGCGTGCCAGTCATCCGCCACCTCTTCGCTCCACTGAAGGAATATTTTGCTTGTGTGTAA*
