## Supplementary material for "Dynamic evolution of *de novo* DNA methyltransferases in rodent and primate genomes": Suppl data 7

*SupFile_7: Primate* DNMT3B *sequence alignments*

>Human3B

ATGAAGGGAGACACCAGGCATCTCAATGGAGAGGAGGACGCCGGCGGGAGGGAAGACTCGATCCTCGTCAACGGGGCCTGCAGCGACCAGTCCTCCGACTCGCCCCCAATCCTGGAGGCTATCCGCACCCCGGAGATCAGAGGCCGAAGATCAAGCTCGCGACTCTCCAAGAGGGAGGTGTCCAGTCTGCTAAGCTACACACAGGACTTGACAGGCGATGG------CGACGGGGAAGATGGGGATGGCTCTGACACCCCAGTCATGCCAAAGCTCTTCCGGGAAACCAGGACTCGTTCAGAAAGCCCAGCTGTCCGAACTCGAAATAACAACAGTGTCTCCAGCCGGGAGAGGCACAGGCCTTCCCCACGTTCCACCCGAGGCCGGCAGGGCCGCAACCATGTGGACGAGTCCCCCGTGGAGTTCCCGGCTACCAGGTCCCTGAGACGGCGGGCAACAGCATCGGCAGGAACGCCATGGCCGTCCCCTCCCAGCTCTTACCTTACCATCGACCTCACAGACGACACAGAGGACACACATGGGACGCCCCAGAGCAGCAGTACCCCCTACGCCCGCCTAGCCCAGGACAGCCAGCAGGGGGGCATGGAGTC---CCCGCAGGTGGAGGCAGACAGTGGAGATGGAGACAGTTCAGAGTATCAGGATGGGAAGGAGTTTGGAATAGGGGACCTCGTGTGGGGAAAGATCAAGGGCTTCTCCTGGTGGCCCGCCATGGTGGTGTCTTGGAAGGCCACCTCCAAGCGACAGGCTATGTCTGGCATGCGGTGGGTCCAGTGGTTTGGCGATGGCAAGTTCTCCGAGGTCTCTGCAGACAAACTGGTGGCACTGGGGCTGTTCAGCCAGCACTTTAATTTGGCCACCTTCAATAAGCTCGTCTCCTATCGAAAAGCCATGTACCATGCTCTGGAGAAAGCTAGGGTGCGAGCTGGCAAGACCTTCCCCAGCAGCCCTGGAGACTCATTGGAGGACCAGCTGAAGCCCATGTTGGAGTGGGCCCACGGGGGCTTCAAGCCCACTGGGATCGAGGGCCTCAAACCCAACAACACGCAACCAGTGGTTAATAAGTCGAAGGTGCGTCGTGCAGGCAGTAGGAAATTAGAATCAAGGAAATACGAGAACAAGACTCGAAGACGCACAGCTGACGACTCAGCCACCTCTGACTACTGCCCCGCACCCAAGCGCCTCAAGACAAATTGCTATAACAACGGCAAAGACCGAGGGGATGAAGATCAGAGCCGAGAACAAATGGCTTCAGATGTTGCCAACAACAAGAGCAGCCTGGAAGATGGCTGTTTGTCTTGTGGCAGGAAAAACCCCGTGTCCTTCCACCCTCTCTTTGAGGGGGGGCTCTGTCAGACATGCCGGGATCGCTTCCTTGAGCTGTTTTACATGTATGATGACGATGGCTATCAGTCTTACTGCACTGTGTGCTGCGAGGGCCGAGAGCTGCTGCTTTGCAGCAACACGAGCTGCTGCCGGTGTTTCTGTGTGGAGTGCCTGGAGGTGCTGGTGGGCACAGGCACAGCGGCCGAGGCCAAGCTTCAGGAGCCCTGGAGCTGTTACATGTGTCTCCCGCAGCGCTGTCATGGCGTCCTGCGGCGCCGGAAGGACTGGAACGTGCGCCTGCAGGCCTTCTTCACCAGTGACACGGGGCTTGAATATGAAGCCCCCAAGCTGTACCCTGCCATTCCCGCAGCCCGAAGGCGGCCCATTCGAGTCCTGTCATTGTTTGATGGCATCGCGACAGGCTACCTAGTCCTCAAAGAGTTGGGCATAAAGGTAGGAAAGTACGTCGCTTCTGAAGTGTGTGAGGAGTCCATTGCTGTTGGAACCGTGAAGCACGAGGGGAATATCAAATACGTGAACGACGTGAGGAACATCACAAAGAAAAATATTGAAGAATGGGGCCCATTTGACTTGGTGATTGGCGGAAGCCCATGCAACGATCTCTCAAATGTGAATCCAGCCAGGAAAGGCCTGTATGAGGGTACAGGCCGGCTCTTCTTCGAATTTTACCACCTGCTGAATTACTCACGCCCCAAGGAGGGTGATGACCGGCCGTTCTTCTGGATGTTTGAGAATGTTGTAGCCATGAAGGTTGGCGACAAGAGGGACATCTCACGGTTCCTGGAGTGTAATCCAGTGATGATTGATGCCATCAAAGTTTCTGCTGCTCACAGGGCCCGATACTTCTGGGGCAACCTACCCGGGATGAACAGGCCCGTGATAGCATCAAAGAATGATAAACTCGAGCTGCAGGACTGCTTGGAATACAATAGGATAGCCAAGTTAAAGAAAGTACAGACAATAACCACCAAGTCGAACTCGATCAAACAGGGGAAAAACCAACTTTTCCCTGTTGTCATGAATGGCAAAGAAGATGTTTTGTGGTGCACTGAGCTCGAAAGGATCTTTGGCTTTCCTGTGCACTACACAGACGTGTCCAACATGGGCCGTGGTGCCCGCCAGAAGCTGCTGGGAAGGTCCTGGAGCGTGCCTGTCATCCGACACCTCTTCGCCCCTCTGAAGGACTACTTTGCATGTGAATAG

>Chimp3B

ATGAAGGGAGACACCAGGCATCTCAATGGAGAGGAGGACGCCGGCGGGAGGGAAGACTCGATCCTCGTCAACGGGGCCTGCAGCGACCAGTCCTCCGACTCGCCCCCAATCCTGGAGGCTATCCGCACCCCGGAGATCAGAGGCCGAAGATCGAGCTCGCGACTCTCCAAGAGGGAGGTGTCCAGTCTGCTAAGCTACACGCAGGACTTGACAGGCGATGG------CGACGGGGAAGATGGGGATGGCTCTGACACCCCAGTGATGCCAAAGCTCTTCCGGGAAACCAGGACTCGTTCAGAAAGCCCAGCTGTCCGAACTCGAAATAACAACAGTGTCTCCAGCCGGGAGAGGCACAGGCCTTCCCCACGTTCCACCCGAGGCCGGCAGGGCCGCAACCATGTGGACGAGTCCCCCGTGGAGTTCCCGGCTACCAGGTCCCTGAGACGGCGGGCAACAGCATCGGCAGGAACGCCATGGCCGTCCCCTCCCAGCTCTTACCTTACCATCGACCTCACAGACGACACAGAGGACACACATGGGACGCCCCAGAGCAGCAGTACCCCCTACGCCCGCCTAGCCCAGGACAGCCAGCAGGGGGGCATGGAGTC---CCCGCAGGTGGAGGCAGACAGTGGAGATGGAGACAGTTCAGAGTATCAGGATGGGAAGGAGTTTGGAATAGGGGACCTCGTGTGGGGAAAGATCAAGGGCTTCTCCTGGTGGCCCGCCATGGTGGTGTCTTGGAAGGCCACCTCCAAGCGACAGGCTATGTCTGGCATGCGGTGGGTCCAGTGGTTTGGCGATGGCAAGTTCTCCGAGGTCTCTGCAGACAAACTGGTGGCACTGGGGCTGTTCAGCCAGCACTTTAATTTGGCCACCTTCAATAAGCTCGTCTCCTATCGAAAAGCCATGTACCATGCTCTGGAGAAAGCTAGGGTGCGAGCTGGCAAGACCTTCCCCAGCAGCCCTGGAGACTCATTGGAGGACCAGCTGAAGCCCATGTTGGAGTGGGCCCACGGGGGCTTCAAGCCCACTGGGATCGAGGGCCTCAAACCCAACAACACGCAACCAGTGGTTAATAAGTCGAAGGTGCGTCGTGCAGGCAGTAGGAAATTAGAATCAAGGAAATACGAGAACAAGACTCGAAGACGCACAGCTGATGACTCAGCCACCTCTGACTACTGCCCCGCACCCAAGCGCCTCAAGACAAATTGCTATAACAATGGCAAAGACCGAGGGGATGAAGATCAGAGCCGAGAACAAATGGCTTCAGATGTTGCCAACAACAAGAGCAGCCTGGAAGATGGCTGTTTGTCTTGTGGCAGGAAAAACCCTGTGTCCTTCCACCCTCTCTTTGAGGGGGGGCTCTGTCAGACATGCCGGGATCGCTTCCTTGAGCTGTTTTACATGTATGATGACGATGGCTATCAGTCTTACTGCACTGTGTGCTGCGAGGGCCGAGAGCTGCTGCTTTGCAGCAACACGAGCTGCTGCCGGTGCTTCTGTGTGGAGTGCCTGGAGGTGCTGGTGGGCACAGGCACAGCGGCCGAGGCCAAGCTTCAGGAGCCCTGGAGCTGCTACATGTGTCTCCCGCAGCGCTGTCATGGCGTCCTGCGGCGCCGGAAGGACTGGAACGTGCGCCTGCAGGCCTTCTTCACCAGTGACACGGGGCTTGAATACGAAGCCCCCAAGCTGTACCCTGCCATTCCCGCAGCCCGAAGGCGGCCCATTCGAGTCCTGTCATTGTTTGATGGCATCGCGACAGGCTACCTAGTCCTCAAAGAGTTGGGCATAAAGGTAGGAAAGTACGTCGCCTCTGAAGTGTGTGAGGAGTCCATTGCTGTTGGAACCGTGAAGCACGAGGGGAATATCAAATACGTGAACGACGTGAGGAACATCACAAAGAAAAATATTGAAGAATGGGGCCCATTTGACTTGGTGATTGGCGGAAGCCCATGCAACGATCTCTCAAATGTGAATCCAGCCAGGAAAGGCCTGTATGAGGGTACAGGCCGGCTCTTCTTCGAATTTTACCACCTGCTGAATTACTCACGCCCCAAGGAGGGTGATGACCGGCCATTCTTCTGGATGTTTGAGAATGTTGTAGCCATGAAGGTTGGCGACAAGAGGGACATCTCACGGTTCCTGGAGTGTAATCCAGTGATGATTGATGCCATCAAAGTTTCTGCTGCTCACAGGGCCCGATACTTCTGGGGCAACCTACCCGGGATGAACAGGCCCGTGATAGCATCAAAGAATGATAAACTCGAGCTGCAGGACTGCTTGGAATACAATAGGATAGCCAAGTTAAAGAAAGTACAGACAATAACCACCAAGTCGAACTCGATCAAACAGGGGAAAAACCAACTTTTCCCTGTTGTCATGAATGGCAAAGAAGATGTTTTGTGGTGCACTGAGCTCGAAAGGATCTTTGGCTTTCCTGTGCACTACACAGACGTGTCCAACATGGGCCGGGGTGCCCGCCAGAAGCTGCTGGGAAGGTCCTGGAGCGTGCCTGTCATCCGACACCTCTTCGCCCCTCTGAAGGACTACTTTGCATGTGAATAG

>Bonobo3B

ATGAAGGGAGACACCAGGCATCTCAATGGAGAGGAGGACGCCGGCGGGAGGGAAGACTCGATCCTCGTCAACGGGGCCTGCAGCGACCAGTCCTCCGACTCGCCCCCAATCCTGGAGGCTATCCGCACCCCGGAGATCAGAGGCCGAAGATCGAGCTCGCGACTCTCCAAGAGGGAGGTGTCCAGTCTGCTAAGCTACACGCAGGACTTGACAGGCGATGG------CGACGGGGAAGATGGGGATGGCTCTGACACCCCAGTGATGCCAAAGCTCTTCCGGGAAACCAGGACTCGTTCAGAAAGCCCAGCTGTCCGAACTCGAAATAACAACAGTGTCTCCAGCCGGGAGAGGCACAGGCCTTCCCCACGTTCCACCCGAGGCCGGCAGGGCCGCAACCATGTGGACGAGTCCCCCGTGGAGTTCCCGGCTACCAGGTCCCTGAGACGGCGGGCAACAGCATCGGCGGGAACGCCATGGCCGTCCCCTCCCAGCTCTTACCTTACCATCGACCTCACAGACGACACAGAGGACACACATGGGACGCCCCAGAGCAGCAGTACCCCCTACGCCCGCCTAGCCCAGGACAGCCAGCAGGGGGGCATGGAGTC---CCCGCAGGTGGAGGCAGACAGTGGAGATGGAGACAGTTCAGAGTATCAGGATGGGAAGGAGTTTGGAATAGGGGACCTCGTGTGGGGAAAGATCAAGGGCTTCTCCTGGTGGCCCGCCATGGTGGTGTCTTGGAAGGCCACCTCCAAGCGACAGGCTATGTCTGGCATGCGGTGGGTCCAGTGGTTTGGCGATGGCAAGTTCTCCGAGGTCTCTGCAGACAAACTGGTGGCACTGGGGCTGTTCAGCCAGCACTTTAATTTGGCCACCTTCAATAAGCTCGTCTCCTATCGAAAAGCCATGTACCATGCTCTGGAGAAAGCTAGGGTGCGAGCTGGCAAGACCTTCCCCAGCAGCCCTGGAGACTCATTGGAGGACCAGCTGAAGCCCATGTTGGAGTGGGCCCACGGGGGCTTCAAGCCCACTGGGATCGAGGGCCTCAAACCCAACAACACGCAACCAGTGGTTAATAAGTCGAAGGTGCGTCGTGCAGGCAGTAGGAAATTAGAATCAAGGAAATACGAGAACAAGACTCGAAGACGCACAGCTGACGACTCAGCCACCTCTGACTACTGCCCCGCACCCAAGCGCCTCAAGACAAATTGCTATAACAACGGCAAAGACCGAGGGGATGAAGATCAGAGCCGAGAACAAATGGCTTCAGATGTTGCCAACAACAAGAGCAGCCTGGAAGATGGCTGTTTGTCTTGTGGCAGGAAAAACCCTGTGTCCTTCCACCCTCTCTTTGAGGGGGGGCTCTGTCAGACATGCCGGGATCGCTTCCTTGAGCTGTTTTACATGTATGATGACGATGGCTATCAGTCTTACTGCACTGTGTGCTGCGAGGGCCGAGAGCTGCTGCTTTGCAGCAACACGAGCTGCTGCCGGTGCTTCTGTGTGGAGTGCCTGGAGGTGCTGGTGGGCACAGGCACAGCGGCCGAGGCCAAGCTTCAGGAGCCCTGGAGCTGCTACATGTGTCTCCCGCAGCGCTGTCATGGCGTCCTGCGGCGCCGGAAGGACTGGAACGTGCGCCTGCAGGCCTTCTTCACCAGTGACACGGGGCTTGAATACGAAGCCCCCAAGCTGTACCCTGCCATTCCCGCAGCCCGAAGGCGGCCCATTCGAGTCCTGTCATTGTTTGATGGCATCGCGACAGGCTACCTAGTCCTCAAAGAGTTGGGCATAAAGGTAGGAAAGTACGTCGCCTCTGAAGTGTGTGAGGAGTCCATTGCTGTTGGAACCGTGAAGCACGAGGGGAATATCAAATACGTGAACGACGTGAGGAACATCACAAAGAAAAATATTGAAGAATGGGGCCCATTTGACTTGGTAATTGGCGGAAGCCCATGCAACGATCTCTCAAATGTGAATCCAGCCAGGAAAGGCCTGTATGAGGGTACAGGCCGGCTCTTCTTCGAATTTTACCACCTGCTGAATTACTCACGCCCCAAGGAGGGTGATGACCGGCCATTCTTCTGGATGTTTGAGAATGTTGTAGCCATGAAGGTTGGCGACAAGAGGGACATCTCACGGTTCCTGGAGTGTAATCCAGTGATGATTGATGCCATCAAAGTTTCTGCTGCTCACAGGGCCCGATACTTCTGGGGCAACCTACCCGGGATGAACAGGCCCGTGATAGCATCAAAGAATGATAAACTCGAGCTGCAGGACTGCTTGGAATACAATAGGATAGCCAAGTTAAAGAAAGTACAGACAATAACCACCAAGTCGAACTCGATCAAACAGGGGAAAAACCAACTTTTCCCTGTTGTCATGAATGGCAAAGAAGATGTTTTGTGGTGCACTGAGCTCGAAAGGATCTTTGGCTTTCCTGTGCACTACACAGACGTGTCCAACATGGGCCGGGGTGCCCGCCAGAAGCTGCTGGGAAGGTCCTGGAGCGTGCCTGTCATCCGACACCTCTTCGCCCCTCTGAAGGACTACTTTGCCTGTGAATAG

>Gorilla3B

ATGAAGGGAGACACCAGGCATCTCAATGGAGAGGAGGACGCCGGCGGGAGGGAAGACTCGATCCTCGTCAACGGGGCCTGCAGCGACCAGTCCTCCGACTCGCCCCCAATCCTGGAGGCTATCCGCACCCCGGAGATCAGAGGCCGAAGATCAAGCTCGCGACTCTCCAAGAGGGAGGTGTCCAGTCTGCTAAGCTACACGCAGGACTTGACAGGCGATGG------CGACGGGGAAGATGGGGATGGCTCTGACACCCCAGTGATGCCAAAGCTCTTCCGGGAAACCAGGACTCGTTCAGAAAGCCCAGCTGTCCGAACTCGAAATAACAACAGTGTCTCCAGCCGGGAGAGGCACAGGCCTTCCCCACGTTCCACCCGAGGCCGGCAGGGCCGCAACCATGTGGACGAGTCCCCCGTGGAGTTCCCGGCTACCAGGTCCCTGAGACGGCGGGCAACAGCATCGGCAGGAACGCCATGGCCGTCCCCTCCCAGCTCTTACCTTACCATCGACCTCACAGACGACACAGAGGACACACATGGGACGCCCCAGAGCAGCAGTACCCCCTACGCCCGCCTAGCCCAGGACAGCCAGCAGGGGGGCATGGAGTC---CCCGCAGGTGGAGGCAGACAGTGGAGATGGAGACAGTTCAGAGTATCAGGATGGGAAGGAGTTTGGAATAGGGGACCTCGTGTGGGGAAAGATCAAGGGCTTCTCCTGGTGGCCCGCCATGGTGGTGTCTTGGAAGGCCACCTCCAAGCGACAGGCTATGTCTGGCATGCGGTGGGTCCAGTGGTTTGGCGATGGCAAGTTCTCCGAGGTCTCTGCAGACAAACTGGTGGCACTGGGGCTGTTCAGCCAGCACTTTAATTTGGCCACCTTCAATAAGCTCGTCTCCTATCGAAAAGCCATGTACCATGCTCTGGAGAAAGCTAGGGTGCGAGCTGGCAAGACCTTCCCCAGCAGCCCTGGAGACTCATTGGAGGACCAGCTGAAGCCCATGTTGGAGTGGGCCCACGGGGGCTTCAAGCCCACTGGGATCGAGGGCCTCAAACCCAACAACACGCAACCAGTGGTTAATAAGTCGAAGGTGCGTCGTGCAGGCAGTAGGAAATTAGAATCAAGGAAATACGAGAACAAGACTCGAAGACGCACAGCTGACGACTCAGCCACCTCTGACTACTGCCCCGCACCCAAGCGCCTCAAGACAAATTGCTATAACAACGGCAAAGACCGAGGGGATGAAGATCAGAGCCGAGAACAAATGGCTTCAGATGTTGCCAACAACAAGAGCAGCCTGGAAGGTGGCTGTTTGTCTTGTGGCAGGAAAAACCCCGTGTCCTTCCACCCTCTCTTTGAAGGGGGGCTCTGTCAGACATGCCGGGATCGCTTCCTTGAGCTGTTTTACATGTATGACGACGATGGCTATCAGTCTTACTGCACTGTGTGCTGCGAGGGCCGAGAGCTGCTGCTTTGCAGCAACACGAGCTGCTGCCGGTGCTTCTGTGTGGAGTGCCTGGAGGTGCTGGTGGGCACAGGCACAGCGGCCGAGGCCAAGCTTCAGGAGCCCTGGAGCTGCTACATGTGTCTCCCGCAGCGCTGTCATGGCGTCCTGCGGCGCCGGAAGGACTGGAACGTGCGCCTGCAGGCCTTCTTCACCAGTGACACGGGGCTTGAATACGAAGCCCCCAAGCTGTACCCTGCCATTCCCGCAGCCCGAAGGCGGCCCATTCGAGTCCTGTCATTGTTTGATGGCATCGCGACAGGCTACCTAGTCCTCAAAGAGTTGGGCATAAAGGTAGGAAAGTACGTCGCCTCTGAAGTGTGTGAGGAGTCCATTGCTGTTGGAACCGTGAAGCACGAGGGGAATATCAAATACGTAAACGACGTGAGGAACATCACAAAGAAAAATATTGAAGAATGGGGCCCATTTGACTTGGTGATTGGCGGAAGCCCATGCAACGATCTCTCAAATGTGAATCCAGCCAGGAAAGGCCTGTATGAGGGTACAGGCCGGCTCTTCTTCGAATTTTACCACCTGCTGAATTATTCACGCCCCAAGGAGGGTGATGACCGGCCGTTCTTCTGGATGTTTGAGAATGTTGTAGCCATGAAGGTTGGCGACAAGAGGGACATCTCGCGGTTCCTGGAGTGTAATCCAGTGATGATTGATGCCATCAAAGTTTCTGCTGCTCACAGGGCCCGATACTTCTGGGGCAACCTACCCGGGATGAACAGGCCCGTGATAGCATCAAAGAATGATAAACTCGAGCTGCAGGACTGCTTGGAATACAATAGGATAGCCAAGTTAAAGAAAGTACAGACAATAACCACCAAGTCGAACTCGATCAAACAGGGGAAAAACCAACTTTTCCCTGTTGTCATGAATGGCAAAGAAGATGTTTTGTGGTGCACTGAGCTCGAAAGGATCTTTGGCTTTCCTGTGCACTACACAGATGTGTCCAACATGGGCCGCGGTGCCCGCCAGAAGCTGCTGGGAAGGTCCTGGAGCGTGCCTGTCATCCGACACCTCTTCGCCCCTCTGAAGGACTACTTTGCATGTGAATAG

>Orangoutan3B

ATGAAGGGAGACACCAGGCATCTCAATGGAGAGGAGGACGCCGGCGGGAGGGAAGACTCGATCTTCGTCAACGGGGCCTGCAGCGACCAGTCCTCTGACTCGCCCCCGATCCTGGAGGCTATCCGCACCCCGGAGATCAGAGGCCGAAGATCAAGCTCGCGACTCTCCAAGAGGGAGGTGTCCAGTCTGCTAAGCTACACGCAGGACTTGACAGGCGATGG------TGACGGGGAAGATGGGGATGGCTCTGACACCCCAGTGATGCCAAAGCTCTTCCGGGAAACCAGGACTCGGTCAGAAAGCCCAGCTGTCCGAACTCGAAATAACAACAGTGTCTCCAGCCGGGAGAGGCACAGGCCTTCCCCACGTTCCACCCGAGGCCGGCAGGGCCGCAACCATGTGGACGAGTCCCCCGTGGAGTTCCCGGCTACCAGGTCCCTGAGACGGCGGGCAACAGCATCGGCAGGAGCGCCATGGCCGTCCCCTCCCAGCTCTTACCTTACCATCGACCTCACAGACGACACAGAGGACACACATGGGACACCCCAGAGCAGCAGTACACCCTACGCCCGCCTAGCCCAGGACAGCCAGCAGGGGGGCATGGAGTC---CCCGCAGGTGGAGGCAGACAGTGGAGATGGAGACAGTTCAGAGTATCAGGATGGGAAGGAGTTTGGAATAGGGGACCTCGTGTGGGGAAAGATCAAGGGCTTCTCCTGGTGGCCCGCCATGGTGGTGTCTTGGAAGGCCACCTCCAAGCGACAGGCTATGTCTGGCATGCGGTGGGTCCAGTGGTTTGGCGATGGCAAGTTCTCCGAGGTCTCTGCAGACAAACTGGTGGCACTGGGGCTGTTCAGCCAGCACTTTAATTTGGCCACCTTCAATAAGCTCGTCTCCTATCGAAAAGCCATGTACCATGCTCTGGAGAAAGCTAGGGTGCGAGCTGGCAAGACCTTCCCCAGCAGCCCTGGAGACTCATTGGAGGACCAGCTGAAGCCCATGTTGGAGTGGGCCCACGGGGGCTTCAAGCCCACTGGGATCGAGGGCCTCAAACCCAACAACACGCAACCAGTGGTTAATAAGTCGAAGGTGCGTCGTGCAGGCAGTAGGAAATTAGAATCAAGGAAATACGAGAACAAGACTCGAAGACGCACAGCTGACGACTCAGCCACCTCTGACTACTGCCCCACACCCAAGCGCCTCAAGACAAATTGCTATAACAACGGCAAAGACCGAGGGGATGAAGATCAGAGCCGAGAACAAATGGCTTCAGATGTTGCCAACAACAAGAGCAGCCTGGAAGATGGCTGTTTGTCTTGTGGCAGGAAAAACCCCGTGTCCTTCCACCCTCTCTTTGAGGGGGGGCTCTGTCAGACATGCCGGGATCGCTTCCTTGAGCTGTTTTACATGTATGACGACGATGGCTATCAGTCTTACTGCACCGTGTGCTGCGAGGGCCGAGAGCTGCTGCTTTGCAGCAACACGAGCTGCTGCCGGTGCTTCTGTGTGGAGTGCCTGGAGGTGCTGGTGGGCACAGGCACAGCGGCCGAGGCCAAGCTTCAGGAGCCCTGGAGCTGCTACATGTGTCTCCCGCAGCGCTGTCATGGTGTCCTGCGGCGCCGGAAGGACTGGAACGTGCGCCTGCAGGCCTTCTTCACCAGTGACACAGGGCTTGAATACGAGGCCCCCAAGCTGTACCCTGCCATTCCCGCAGCCCGAAGGCGGCCCATTCGAGTCCTGTCATTGTTTGATGGCATTGCGACAGGCTACCTAGTCCTCAAAGAGTTGGGCATAAAGGTAGGAAAGTACGTCGCCTCTGAAGTGTGTGAGGAGTCCATTGCTGTTGGAACCGTGAAGCACGAGGGGAATATCAAATACGTGAATGACGTGAGGAACATCACAAAGAAAAATATTGAAGAATGGGGCCCATTTGACTTGGTGATTGGCGGAAGCCCATGCAACGATCTCTCAAATGTGAATCCAGCCAGGAAAGGCCTGTATGAGGGTACAGGCCGGCTCTTCTTCGAATTTTACCACCTGCTGAATTACTCACGCCCCAAGGAGGGTGATGACCGGCCGTTCTTCTGGATGTTTGAGAATGTTGTAGCCATGAAGGTTGGCGACAAGAGGGACATCTCACGGTTCCTGGAGTGTAATCCAGTGATGATTGATGCCATCAAAGTTTCTGCTGCTCACAGGGCCCGATACTTTTGGGGCAACCTACCCGGGATGAACAGGCCCGTGATAGCATCAAAGAATGATAAACTCGAGCTGCAGGACTGCTTGGAATACAATAGGATAGCCAAGTTAAAGAAAGTACAGACAATAACCACCAAGTCGAACTCGATCAAACAGGGGAAAAACCAACTTTTCCCTGTTGTCATGAATGGCAAAGAAGATGTTTTGTGGTGCACTGAGCTCGAAAGGATCTTTGGCTTTCCTGTGCACTACACAGACGTGTCCAACATGGGCCGCGGTGCCCGCCAGAAGCTGCTGGGAAGGTCCTGGAGCGTGCCTGTCATCCGACACCTCTTCGCCCCTCTGAAGGACTACTTTGCATGTGAATAG

>RhesusMac3B

ATGAAGGGAGACACCAGGCATCTCAACGGAGAGGAGGACGCCGGCGGGAGGGAAGACTCGATCCTCGTCAATGGGGCCTGCAGCGACCAGTCCTCCGACTCGCCCCCGATCCTGGAGGCCATCCGCACCCCGGAGATCAGAGGCCGAAGATCAAGCTCGCGACTCTCCAAGAGGGAGGTGTCCAGTCTGCTAAGCTACACGCAGGACTTGACAGGCGATGG------CGACGGGGAAGATGGGGATGGCTCTGACACCCCAGTGATGCCAAAGCTCTTCCGGGAAACCAGGACTCGGTCAGAAAGCCCAGCTGTCCGAACTCGAAATAACAACAGTGTCTCCAGCCGGGAGAGGCACAGGCCTTCCCCACGTTCCACCCGAGGCCGGCAGGGCCGCAACCATGTGGACGAGTCTCCCGTGGAGTTCCCGGCTACCAGGTCCCTGAGACGGCGGGCAACAGCATCGGCAGGAACGCCATGTCCGTCCCCTGCCAGCTCTTACCTTACCATCGACCTCACAGATGACACAGAGGACACACGTGTGACGCCCCAGAGCAGCAGTACCCCCTATGCCCGCCTAGCCCATGACAGCCAGCAGGAGGGCATGGAGTC---CCCGCAGGTGGAGGCAGACAGTGGAGATGGAGACAGTTCAGAGTATCAGGATGGGAAGGAGTTTGGAATAGGGGACCTCGTGTGGGGAAAGATCAAGGGCTTCTCCTGGTGGCCTGCCATGGTGGTGTCTTGGAAGGCCACCTCTAAGCGACAGGCTATGTCTGGCATGCGGTGGGTCCAGTGGTTTGGCGATGGCAAGTTTTCCGAGGTCTCTGCAGACAAACTGGTGGCACTGGGGCTGTTCAGCCAGCACTTTAATTTGGCCACCTTCAATAAGCTCGTCTCCTATCGAAAAGCCATGTACCATGCTCTGGAGAAAGCTAGGGTGCGAGCTGGCAAGACCTTCCCCAGCAGCCCTGGAGACTCATTGGAGGACCAGCTGAAGCCCATGTTGGAGTGGGCCCACGGGGGCTTCAAGCCCACTGGGATCGAGGGCCTCAAACCCAACAACACGCAACCA------------------------------------------------------------GAGAACAAGACTCGAAGACGCACAGCTGACGACTCAGCCACCTCTGACTACTGCCCCGCACCCAAGCGCCTCAAGACAAATTGCTATAACAACGGCAAAGACCGAGGGGATGAAGATCAGAGCCGAGAACAAATGGCTTCAGATGTTGCCAACAACAAGAGCAGCCTGGAAGACGGCTGTTTGTCTTGTGGCAGGAAAAACCCCGTGTCCTTCCACCCTCTGTTTGAGGGGGGCCTCTGTCAGACATGCCGGGATCGCTTCCTTGAGCTGTTTTACATGTATGACGACGATGGCTATCAGTCTTACTGCACCGTGTGCTGCGAGGGCCGAGAGCTGCTGCTTTGTAGCAACACGAGCTGCTGCCGGTGCTTCTGTGTGGAGTGCCTGGAGGTGCTGGTGGGCACAGGCACAGCGGCCGAGGCCAAGCTTCAGGAGCCCTGGAGCTGCTACATGTGTCTCCCGCAGCGCTGTCACGGCGTCCTGCGGCGCCGGAAGGACTGGAACGTGCGCCTGCAGGCCTTCTTCACCAGTGACACGGGGCTTGAATACGAAGCCCCCAAGCTGTACCCTGCCATTCCCGCAGCCCGAAGGCGGCCCATTCGAGTCCTGTCATTGTTTGATGGCATCGCAACAGGCTACCTAGTCCTCAAAGAGCTGGGCATAAAGGTAGGAAAGTACGTCGCCTCTGAAGTGTGTGAAGAGTCCATCGCTGTTGGAACCGTGAAGCACGAGGGGAATATCAAATACGTGAACGACGTGAGGAACATCACAAAGAAAAATATTGAAGAATGGGGCCCATTTGACTTGGTGATTGGCGGAAGCCCATGCAACGATCTCTCAAATGTGAATCCAGCCAGGAAAGGCCTGTATGAGGGTACAGGCCGGCTCTTCTTTGAATTTTACCACCTGCTGAATTACTCACGCCCCAAGGAGGGTGATGACCGGCCGTTCTTCTGGATGTTTGAGAATGTTGTAGCCATGAAGGTTGGCGACAAGAGGGACATCTCACGGTTCCTGGAGTGTAATCCAGTGATGATTGATGCCATCAAAGTTTCTGCTGCTCACAGGGCCCGATACTTTTGGGGCAACCTACCTGGGATGAACAGGCCCGTGATAGCATCAAAGAATGATAAACTCGAGCTGCAGGACTGCTTGGAATACAATAGGATAGCCAAGTTAAAGAAAGTACAGACAATAACCACCAAGTCGAACTCGATCAAACAGGGGAAAAACCAACTTTTCCCTGTTGTCATGAATGGCAAAGAAGATGTTTTGTGGTGCACTGAGCTCGAAAGGATCTTTGGCTTTCCTGTGCACTACACAGACGTGTCCAACATGGGCCGCGGTGCCCGCCAGAAGCTGCTGGGAAGGTCCTGGAGCGTGCCTGTCATCCGACACCTCTTCGCCCCTCTGAAGGACTACTTTGCATGTGAATAG

>CrabMac3B

ATGAAGGGAGACACCAGGCATCTCAACGGAGAGGAGGACGCCGGCGGGAGGGAAGACTCGATCCTCGTCAATGGGGCCTGCAGCGACCAGTCCTCCGACTCGCCCCCGATCCTGGAGGCCATCCGCACCCCGGAGATCAGAGGCCGAAGATCAAGCTCGCGACTCTCCAAGAGGGAGGTGTCCAGTCTGCTAAGCTACACGCAGGACTTGACAGGCGATGG------CGACGGGGAAGATGGGGATGGCTCTGACACCCCAGTGATGCCAAAGCTCTTCCGGGAAACCAGGACTCGGTCAGAAAGCCCAGCTGTCCGAACTCGAAATAACAACAGTGTCTCCAGCCGGGAGAGGCACAGGCCTTCCCCACGTTCCACCCGAGGCCGGCAGGGCCGCAACCATGTGGACGAGTCTCCCGTGGAGTTCCCGGCTACCAGGTCCCTGAGACGGCGGGCAACAGCATCGGCAGGAACGCCATGGCCGTCCCCTGCCAGCTCTTACCTTACCATCGACCTCACAGATGACACAGAGGACACACGTGTGACGCCCCAGAGCAGCAGTACCCCCTATGCCCGCCTAGCCCATGACAGCCAGCAGGAGGGCATGGAGTC---CCCGCAGGTGGAGGCAGACAGTGGAGATGGAGACAGTTCAGAGTATCAGGATGGGAAGGAGTTTGGAATAGGGGACCTCGTGTGGGGAAAGATCAAGGGCTTCTCCTGGTGGCCTGCCATGGTGGTGTCTTGGAAGGCCACCTCTAAGCGACAGGCTATGTCTGGCATGCGGTGGGTCCAGTGGTTTGGCGATGGCAAGTTTTCCGAGGTCTCTGCAGACAAACTGGTGGCACTGGGGCTGTTCAGCCAGCACTTTAATTTGGCCACCTTCAATAAGCTCGTCTCCTATCGAAAAGCCATGTACCATGCTCTGGAGAAAGCTAGGGTGCGAGCTGGCAAGACCTTCCCCAGCAGCCCTGGAGACTCATTGGAGGACCAGCTGAAGCCCATGTTGGAGTGGGCCCACGGGGGCTTCAAGCCCACTGGGATCGAGGGCCTCAAACCCAACAACACGCAACCAGTGGTTAATAAGTCGAAGGTGCGTCGTGCAGGCAGTAGGAAATTAGAATCAAGGAAATACGAGAACAAGACTCGAAGACGCACAGCTGACGACTCAGCCACCTCTGACTACTGCCCCGCACCCAAGCGCCTCAAGACAAATTGCTATAACAACGGCAAAGACCGAGGGGATGAAGATCAGAGCCGAGAACAAATGGCTTCAGATGTTGCCAACAACAAGAGCAGCCTGGAAGACGGCTGTTTGTCTTGTGGCAGGAAAAACCCCGTGTCCTTCCACCCTCTGTTTGAGGGGGGCCTCTGTCAGACATGCCGGGATCGCTTCCTTGAGCTGTTTTACATGTATGACGACGATGGCTATCAGTCTTACTGCACCGTGTGCTGCGAGGGCCGAGAGCTGCTGCTTTGTAGCAACACGAGCTGCTGCCGGTGCTTCTGTGTGGAGTGCCTGGAGGTGCTGGTGGGCACAGGCACAGCGGCCGAGGCCAAGCTTCAGGAGCCCTGGAGCTGCTACATGTGTCTCCCGCAGCGCTGTCACGGCGTCCTGCGGCGCCGGAAGGACTGGAACGTGCGCCTGCAGGCCTTCTTCACCAGTGACACGGGGCTTGAATACGAAGCCCCCAAGCTGTACCCTGCCATTCCCGCAGCCCGAAGGCGGCCCATTCGAGTCCTGTCATTGTTTGATGGCATCGCAACAGGCTACCTAGTCCTCAAAGAGCTGGGCATAAAGGTAGGAAAGTACGTCGCCTCTGAAGTGTGTGAAGAGTCCATCGCTGTTGGAACCGTGAAGCACGAGGGGAATATCAAATACGTGAACGACGTGAGGAACATCACAAAGAAAAATATTGAAGAATGGGGCCCATTTGACTTGGTGATTGGCGGAAGCCCATGCAACGATCTCTCAAATGTGAATCCAGCCAGGAAAGGCCTGTATGAGGGTACCGGCCGGCTCTTCTTTGAATTTTACCACCTGCTGAATTACTCACGCCCCAAGGAGGGTGATGACCGGCCGTTCTTCTGGATGTTTGAGAATGTTGTAGCCATGAAGGTTGGCGACAAGAGGGACATCTCACGGTTCCTGGAGTGTAATCCAGTGATGATTGATGCCATCAAAGTTTCTGCTGCTCACAGGGCCCGATACTTTTGGGGCAACCTACCTGGGATGAACAGGCCCGTGATAGCATCAAAGAATGATAAACTCGAGCTGCAGGACTGCTTGGAATACAATAGGATAGCCAAGTTAAAGAAAGTACAGACAATAACCACCAAGTCGAACTCGATCAAACAGGGGAAAAACCAACTTTTCCCTGTTGTCATGAATGGCAAAGAAGATGTTTTGTGGTGCACTGAGCTCGAAAGGATCTTTGGCTTTCCTGTGCACTACACAGACGTGTCCAACATGGGCCGCGGTGCCCGCCAGAAGCTGCTGGGAAGGTCCTGGAGCGTGCCTGTCATCCGACACCTCTTCGCCCCTCTGAAGGACTACTTTGCATGTGAATAG

>PigTailedMac3B

ATGAAGGGAGACACCAGGCATCTCAACGGAGAGGAGGACGCCGGCGGGAGGGAAGACTCGATCCTCGTCAATGGGGCCTGCAGCGACCAGTCCTCCGACTCGCCCCCGATCCTGGAGGCCATCCGCACCCCGGAGATCAGAGGCCGAAGATCAAGCTCGCGACTCTCCAAGAGGGAGGTGTCCAGTCTGCTAAGCTACACGCAGGACTTGACAGGCGATGG------CGACGGGGAAGATGGGGATGGCTCTGACACCCCAGTGATGCCAAAGCTCTTCCGGGAAACCAGGACTCGGTCAGAAAGCCCAGCTGTCCGAACTCGAAATAACAACAGTGTCTCCAGCCGGGAGAGGCACAGGCCTTCCCCACGTTCCACCCGAGGCCGGCAGGGCCGCAACCATGTGGACGAGTCTCCCGTGGAGTTCCCGGCTACCAGGTCCCTGAGACGGCGGGCAACAGCATCGGCAGGAACGCCATGGCCGTCCCCTGCCAGCTCTTACCTTACCATCGACCTCACAGATGACACAGAGGACACACGTGTGACGCCCCAGAGCAGCAGTACCCCCTATGCCCGCCTAGCCCATGACAGCCAGCAGGAGGGCATGGAGTC---CCCGCAGGTGGAGGCAGACAGTGGAGATGGAGACAGTTCAGAGTATCAGGATGGGAAGGAGTTTGGAATAGGGGACCTCGTGTGGGGAAAGATCAAGGGCTTCTCCTGGTGGCCTGCCATGGTGGTGTCTTGGAAGGCCACCTCTAAGCGACAGGCTATGTCTGGCATGCGGTGGGTCCAGTGGTTTGGCGATGGCAAGTTTTCCGAGGTCTCTGCAGACAAACTGGTGGCACTGGGGCTGTTCAGCCAGCACTTTAATTTGGCCACCTTCAATAAGCTCGTCTCCTATCGAAAAGCCATGTACCATGCTCTGGAGAAAGCTAGGGTGCGAGCTGGCAAGACCTTCCCCAGCAGCCCTGGAGACTCATTGGAGGACCAGCTGAAGCCCATGTTGGAGTGGGCCCACGGGGGCTTCAAGCCCACTGGGATCGAGGGCCTCAAACCCAACAACACGCAACCA------------------------------------------------------------GAGAACAAGACTCGAAGACGCACAGCTGACGACTCAGCCACCTCTGACTACTGCCCCGCACCCAAGCGCCTCAAGACAAATTGCTATAACAACGGCAAAGACCGAGGGGATGAAGATCAGAGCCGAGAACAAATGGCTTCAGATGTTGCCAACAACAAAAGCAGCCTGGAAGACGGCTGTTTGTCTTGTGGCAGGAAAAACCCCGTGTCCTTCCACCCTCTGTTTGAGGGGGGCCTCTGTCAGACATGCCGGGATCGCTTCCTTGAGCTGTTTTACATGTATGACGACGATGGCTATCAGTCTTACTGCACCGTGTGCTGCGAGGGCCGAGAGCTGCTGCTTTGTAGCAACACGAGCTGCTGCCGGTGCTTCTGTGTGGAGTGCCTGGAGGTGCTGGTGGGCACAGGCACAGCGGCCGAGGCCAAGCTTCAGGAGCCCTGGAGCTGCTACATGTGTCTCCCGCAGCGCTGTCACGGCGTCCTGCGGCGCCGGAAGGACTGGAACGTGCGCCTGCAGGCCTTCTTCACCAGTGACACGGGGCTTGAATACGAAGCCCCCAAGCTGTACCCTGCCATTCCCGCAGCCCGAAGGCGGCCCATTCGAGTCCTGTCATTGTTTGATGGCATCGCAACAGGCTACCTAGTCCTCAAAGAGCTGGGCATAAAGGTAGGAAAGTACGTCGCCTCTGAGGTGTGTGAAGAGTCCATCGCTGTTGGAACCGTAAAGCACGAGGGGAATATCAAATACGTGAACGACGTGAGGAACATCACAAAGAAAAATATTGAAGAATGGGGCCCATTTGACTTGGTGATTGGCGGAAGCCCATGCAACGATCTCTCAAATGTGAATCCAGCCAGGAAAGGCCTGTATGAGGGTACAGGCCGGCTCTTCTTTGAATTTTACCACCTGCTGAATTACTCACGCCCCAAGGAGGGTGATGACCGGCCGTTCTTCTGGATGTTTGAGAATGTTGTAGCCATGAAGGTTGGCGACAAGAGGGACATCTCACGGTTCCTGGAGTGTAATCCAGTGATGATTGATGCCATCAAAGTTTCTGCTGCTCACAGGGCCCGATACTTTTGGGGCAACCTACCTGGGATGAACAGGCCCGTGATAGCATCAAAGAATGATAAACTCGAGCTGCAGGACTGCTTGGAATACAATAGGATAGCCAAGTTAAAGAAAGTACAGACAATAACCACCAAGTCGAACTCGATCAAACAGGGGAAAAACCAACTTTTCCCTGTTGTCATGAATGGCAAAGAAGATGTTTTGTGGTGCACTGAGCTCGAAAGGATCTTTGGCTTTCCTGTGCACTACACAGACGTGTCCAACATGGGCCGCGGTGCCCGCCAGAAGCTGCTGGGAAGGTCCTGGAGCGTGCCTGTCATCCGACACCTCTTCGCCCCTCTGAAGGACTACTTTGCATGTGAATAG

>Baboon

ATGAAGGGAGACACCAGGCATCTCAACGGAGAGGAGGACGCCGGCGGGAGGGAAGACTCAATCTTCGTCAATGGGGCCTGCAGCGACCAGTCCTCCGACTCGCCCCCGATCCTGGAGGCCATCCGCACCCCGGAGATCAGAGGCCGAAGATCAAGCTCGCGACTCTCCAAGAGGGAGGTGTCCAGTCTGCTAAGCTACACGCAGGACTTGACAGGCGATGG------CGATGGGGAAGATGGGGATGGCTCTGACACCCCAGTGATGCCAAAGCTTTTCCGGGAAACCAGGACTCGGTCAGAAAGCCCAGCTGTCCGAACTCGAAATAACAACAGTGTCTCCAGCCGGGAGAGGCACAGGCCTTCCCCACGTTCCACCCGAGGCCGGCAGGGCCGCAACCATGTGGACGAGTCTCCCGTGGAGTTCCCGGCTACCAGGTCCCTGAGACGGCGGGCAACAGCATCGGTAGGAACGCCATGGCCGTCCCCTGCCAGCTCTTACCTTACCATCGACCTCACAGATGACACAGAGGACACACGTGTGACGCCCCAGAGCAGCAGTACCCCCTATGCCCGCCTAGCCCAGGACAGCCAGCAGGAGGGCATGGAGTC---CCCGCAGGTGGAGGCAGACAGTGGAGATGGAGACAGTTCAGAGTATCAGGATGGGAAGGAGTTTGGAATAGGGGACCTCGTGTGGGGAAAGATCAAGGGCTTCTCCTGGTGGCCTGCCATGGTGGTGTCTTGGAAGGCCACCTCTAAGCGACAGGCTATGTCTGGCATGCGGTGGGTCCAGTGGTTTGGCGATGGCAAGTTTTCCGAGGTCTCTGCAGACAAACTGGTGGCACTGGGGCTGTTCAGCCAGCACTTTAATTTGGCCACCTTCAATAAGCTCGTCTCCTATCGAAAAGCCATGTACCATGCTCTGGAGAAAGCTAGGGTGCGAGCTGGCAAGACCTTCCCCAGCAGCCCTGGAGACTCATTGGAGGACCAGCTGAAGCCCATGTTGGAGTGGGCCCACGGGGGCTTCAAGCCCACTGGGATCGAGGGCCTCAAACCCAACAACACGCAACCA------------------------------------------------------------GAGAACAAGACTCGAAGACGCACAGCTGACGACTCAGCCACCTCTGACTACTGCCCCGCACCCAAGCGCCTCAAGACAAATTGCTATAACAACGGCAAAGACCGAGGGGATGAAGATCAGAGCCGAGAACAAATGGCTTCAGATGTTGCCAACAACAAGAGCAGCCTGGAAGACGGCTGTTTGTCTTGTGGCAGGAAAAACCCCGTGTCCTTCCACCCTCTGTTTGAGGGGGGCCTCTGTCAGACATGCCGGGATCGCTTCCTTGAGCTGTTTTACATGTATGACGACGATGGCTATCAGTCTTACTGCACCGTGTGCTGCGAGGGCCGAGAGCTGCTGCTTTGTAGCAACACAAGCTGCTGCCGGTGCTTCTGTGTGGAGTGCCTGGAGGTGCTGGTGGGCACAGGCACAGCGGCCGAGGCCAAGCTTCAGGAGCCCTGGAGCTGCTACATGTGTCTCCCGCAGCGCTGTCACGGCGTCCTGCGGCGCCGGAAGGACTGGAACGTGCGCCTGCAGGCCTTCTTCACCAGTGACACGGGGCTTGAATACGAAGCCCCCAAGCTGTACCCTGCCATTCCCGCAGCCCGAAGGCGGCCCATTCGAGTCCTGTCATTGTTTGATGGCATTGCGACAGGCTACCTAGTCCTCAAAGAGCTGGGCATAAAGGTAGGAAAGTACGTCGCCTCTGAAGTGTGTGAAGAGTCCATCGCTGTTGGAACTGTGAAGCACGAGGGGAATATCAAATATGTGAACGACGTGAGGAACATCACAAAGAAAAATATTGAAGAATGGGGCCCATTTGACTTGGTGATTGGCGGAAGCCCATGCAATGATCTCTCAAATGTGAATCCAGCCAGGAAAGGCCTGTATGAGGGTACAGGCCGGCTCTTCTTTGAATTTTACCACCTGCTGAATTACTCACGCCCCAAGGAGGGTGATGACCGGCCGTTCTTCTGGATGTTTGAGAATGTTGTAGCCATGAAGGTTGGCGACAAGAGGGACATCTCGCGGTTCCTGGAGTGTAATCCAGTGATGATTGATGCCATCAAAGTTTCTGCTGCTCACAGGGCCCGATACTTTTGGGGCAACCTACCTGGGATGAACAGGCCCGTGATAGCATCAAAGAATGATAAACTCGAGCTGCAGGACTGCTTGGAATACAATAGGATAGCCAAGTTAAAGAAAGTACAGACAATAACCACCAAGTCGAACTCGATCAAACAGGGGAAAAACCAACTTTTCCCTGTTGTCATGAATGGCAAAGAAGATGTTTTGTGGTGCACTGAGCTCGAAAGGATCTTTGGCTTTCCTGTGCACTACACAGACGTGTCCAACATGGGCCGCGGTGCCCGCCAGAAGCTGCTGGGAAGGTCCTGGAGCGTGCCTGTCATCCGACACCTCTTCGCCCCTCTGAAGGACTACTTTGCATGTGAATAG

>Mangabey

ATGAAGGGAGACACCAGGCATCTCAACGGAGAGGAGGACGCCGGCGGGAGGGAAGACTCGATCCTCGTCAATGGGGCCTGCAGCGACCAGTCCTCCGACTCGCCCCCGATCCTGGAGGCCATCCGCACCCCGGAGATCAGAGGCCGAAGATCAAGCTCGCGACTCTCCAAGAGGGAGGTGTCCAGTCTGCTAAGCTACACGCAGGACTTGACAGGCGATGG------CGATGGGGAAGATGGGGATGGCTCTGACACCCCAGTGATGCCAAAGCTCTTCCGGGAAACCAGGACTCGGTCAGAAAGCCCAGCTGTCCGAACTCGAAATAACAACAGTGTCTCCAGCCGGGAGAGGCACAGGCCTTCCCCACGTTCCACCCGAGGCCGGCAGGGCCGCAACCATGTGGACGAGTCTCCCGTGGAGTTCCCGGCTACCAGGTCCCTGAGACGGCGGGCAACAGCATCGGCAGGAACGCCATGGCCGTCCCCTGCCAGCTCTTACCTTACCATCGACCTCACAGATGACACAGAGGACACACGTGTGACGCCCCAGAGCAGCAGTACCCCCTATGCCCGCCTAGCCCAGGACAGCCAGCAGGAGGGCATGGAGTC---CCCGCAGGTGGAGGCAGACAGTGGAGATGGAGACAGTTCAGAGTATCAGGATGGGAAGGAGTTTGGAATAGGGGACCTCGTGTGGGGAAAGATCAAGGGCTTCTCCTGGTGGCCTGCCATGGTGGTGTCTTGGAAGGCCACCTCTAAGCGACAGGCTATGTCTGGCATGCGGTGGGTCCAGTGGTTTGGCGATGGCAAGTTTTCCGAGGTCTCTGCAGACAAACTGGTGGCACTGGGGCTGTTCAGCCAGCACTTTAATTTGGCCACCTTCAATAAGCTCGTCTCCTATCGAAAAGCCATGTACCATGCTCTGGAGAAAGCTAGGGTGCGAGCTGGCAAGACCTTCCCCAGCAGCCCTGGAGACTCATTGGAGGACCAGCTGAAGCCCATGTTGGAGTGGGCCCACGGGGGCTTCAAGCCCACTGGGATCGAGGGCCTCAAACCCAACAACACGCAACCAGTGGTTAATAAGTCGAAGGTGCGTCGTGCAGGCAGTAGGAAATTAGAATCAAGGAAATACGAGAACAAGACTCGAAGACGCACAGCTGACGACTCAGCCACCTCTGACTACTGCCCCGCACCCAAGCGCCTCAAGACAAATTGCTATAACAACGGCAAAGACCGAGGGGATGAAGATCAGAGCCGAGAACAAATGGCTTCAGATGTTGCCAACAACAAGAGCAGCCTGGAAGACGGCTGTTTGTCTTGTGGCAGGAAAAACCCCGTGTCCTTCCACCCTCTGTTTGAGGGGGGCCTCTGTCAGACATGCCGGGATCGCTTCCTTGAGCTGTTTTACATGTATGACGACGATGGCTATCAGTCTTACTGCACCGTGTGCTGCGAGGGCCGAGAGCTGCTGCTTTGTAGCAACACGAGCTGCTGCCGGTGCTTCTGTGTGGAGTGCCTGGAGGTGCTGGTGGGCACAGGCACAGCGGCCGAGGCCAAGCTTCAGGAGCCCTGGAGCTGCTACATGTGTCTCCCGCAGCGCTGTCACGGCGTCCTGCGGCGCCGGAAGGACTGGAACGTGCGCCTGCAGGCCTTCTTCACCAGTGACACGGGGCTTGAATACGAAGCCCCCAAGCTGTACCCTGCCATTCCCGCAGCCCGAAGGCGGCCCATTCGAGTCCTGTCATTGTTTGATGGCATTGCGACAGGCTACCTAGTCCTCAAAGAGCTGGGCATAAAGGTAGGAAAGTACGTCGCCTCTGAAGTGTGTGAAGAGTCCATCGCTGTTGGAACCGTGAAGCACGAGGGGAATATCAAATACGTGAACGACGTGAGGAACATCACAAAGAAAAATATTGAAGAATGGGGCCCATTTGACTTGGTGATTGGCGGAAGCCCATGCAACGATCTCTCAAATGTGAATCCAGCCAGGAAAGGCCTGTATGAGGGTACGGGCCGGCTCTTCTTTGAATTTTACCACCTGCTGAATTACTCACGCCCCAAGGAGGGTGATGACCGGCCGTTCTTCTGGATGTTTGAGAATGTTGTAGCCATGAAGGTTGGCGACAAGAGGGACATCTCGCGGTTCCTGGAGTGTAATCCAGTGATGATTGATGCCATCAAAGTTTCTGCTGCTCACAGGGCCCGATACTTTTGGGGCAACCTACCTGGGATGAACAGGCCCGTGATAGCATCAAAGAATGATAAACTCGAGCTGCAGGACTGCTTGGAATACAATAGGATAGCCAAGTTAAAGAAAGTACAGACAATAACCACCAAGTCGAACTCGATCAAACAGGGGAAAAACCAACTTTTCCCTGTTGTCATGAATGGCAAAGAAGATGTTTTGTGGTGCACTGAGCTCGAAAGGATCTTTGGCTTCCCTGTGCACTACACAGACGTGTCCAACATGGGCCGCGGTGCCCGCCAGAAGCTGCTGGGAAGGTCCTGGAGCGTGCCTGTCATCCGACACCTCTTCGCCCCTCTGAAGGACTACTTTGCATGTGAATAG

>Drill

ATGAAGGGAGACACCAGGCATCTCAACGGAGAGGAGGACGCCGGCGGGAGGGAAGACTCGATCCTCGTCAATGGGGCCTGCAGCGACCAGTCCTCCGACTCGCCCCCGATCCTGGAGGCCATCCGCACCCCGGAGATCAGAGGCCGAAGATCAAGCTCGCGACTCTCCAAGAGGGAGGTGTCCAGTCTGCTAAGCTACACGCAGGACTTGACAGGCGATGG------TGACGGGGAAGATGGGGATGGCTCTGACACCCCAGTGATGCCAAAGCTCTTCCGGGAAACCAGGACTCGGTCAGAAAGCCCAGCTGTCCGAACTCGAAATAACAACAGCGTCTCCAGCCGGGAGAGGCACAGGCCTTCCCCACGTTCCACCCGAGGCCGGCAGGGCCGCAACCATGTGAACGAGTCTCCCGTGGAGTTCCCGGCTACGAGGTCCCTGAGACGGCGGGCAACAGCATCGGCAGGAACGCCATGGCCGTCCCCTGCCAGCTCTTACCTTACCATCGACCTCACAGATGACACAGAGGACACACGTGTGACGCCCCAGAGCAGCAGTACCCCCTATGCCCGCCTAGCCCAGGACAGCCAGCAGGAGGGCATGGAGTC---CCCGCAGGTGGAGGCAGACAGTGGAGATGGAGACAGTTCAGAGTATCAGGATGGGAAGGAGTTTGGAATAGGGGACCTCGTGTGGGGAAAGATCAAGGGCTTCTCCTGGTGGCCTGCCATGGTGGTGTCTTGGAAGGCCACCTCTAAGCGACAGGCTATGTCTGGCATGCGGTGGGTCCAGTGGTTTGGCGATGGCAAGTTTTCCGAGGTCTCTGCAGACAAACTGGTGGCACTGGGGCTGTTCAGCCAGCACTTTAATTTGGCCACCTTCAATAAGCTCGTCTCCTATCGAAAAGCCATGTACCATGCTCTGGAGAAAGCTAGGGTGCGAGCTGGCAAGACCTTCCCCAGCAGCCCTGGAGACTCATTGGAGGACCAGCTGAAGCCCATGTTGGAGTGGGCCCACGGGGGCTTCAAGCCCACTGGGATCGAGGGCCTCAAACCCAACAACACGCAACCAGTGGTTAATAAGTCGAAGGTGCGTCGTGCAGGCAGTAGGAAATTAGAATCAAGGAAATACGAGAACAAGACTCGAAGACGCACAGCTGACGACTCAGCCACCTCTGACTACTGCCCCGCACCCAAGCGCCTCAAGACAAATTGCTATAACAACGGCAAAGACCGAGGGGATGAAGATCAGAGCCGAGAACAAATGGCTTCAGATGTTGCCAACAACAAGAGCAGCCTGGAAGACGGCTGTTTGTCTTGTGGCAGGAAAAACCCCGTGTCCTTCCACCCTCTGTTTGAGGGGGGCCTCTGTCAGACATGCCGGGACCGCTTCCTTGAGCTGTTTTACATGTATGACGACGATGGCTATCAGTCTTACTGCACCGTGTGCTGCGAGGGCCGAGAGCTGCTGCTTTGCAGCAACACGAGCTGCTGCCGGTGCTTCTGTGTGGAGTGCCTGGAGGTGCTGGTGGGCACAGGCACAGCGGCCGAGGCCAAGCTTCAGGAGCCCTGGAGCTGCTACATGTGTCTCCCGCAGCGCTGTCACGGCGTCCTGCGGCGCCGGAAGGACTGGAACGTGCGCCTGCAGGCCTTCTTCACCAGTGACACGGGGCTTGAATACGAAGCCCCCAAGCTGTACCCTGCCATTCCCGCAGCCCGAAGGCGGCCCATTCGAGTCCTGTCATTGTTTGATGGCATTGCGACAGGCTACCTAGTCCTCAAAGAGCTGGGCATAAAGGTAGGAAAGTACGTCGCCTCTGAAGTGTGTGAAGAGTCCATCGCTGTTGGAACCGTGAAGCACGAGGGGAATATCAAATACGTGAACGACGTGAGGAACATCACAAAGAAAAATATTGAAGAATGGGGCCCATTTGACTTGGTGATTGGCGGAAGCCCATGCAACGATCTCTCAAATGTGAATCCAGCCAGGAAAGGCCTGTATGAGGGTACAGGCCGGCTCTTCTTTGAATTTTACCACCTGCTGAATTACTCACGCCCCAAGGAGGGTGATGACCGGCCGTTCTTCTGGATGTTTGAGAATGTTGTAGCCATGAAGGTTGGCGACAAGAGGGACATCTCACGGTTCCTGGAGTGTAATCCAGTGATGATTGATGCCATCAAAGTTTCTGCTGCTCACAGGGCCCGATACTTTTGGGGCAACCTACCTGGGATGAACAGGCCCGTGATAGCATCAAAGAATGATAAACTCGAGCTGCAGGACTGCTTGGAATACAATAGGATAGCCAAGTTAAAGAAAGTACAGACAATAACCACCAAGTCGAACTCGATCAAACAGGGGAAAAACCAACTTTTCCCTGTTGTCATGAATGGCAAAGAAGATGTTTTGTGGTGCACTGAGCTCGAAAGGATCTTTGGCTTTCCTGTGCACTACACAGACGTGTCCAACATGGGCCGCGGTGCCCGCCAGAAGCTGCTGGGAAGGTCCTGGAGCGTGCCTGTCATCCGACACCTCTTCGCCCCTCTGAAGGACTACTTTGCATGTGAATAG

>AfrGreenMonkey

ATGAAGGGAGACACCAGGCATCTCAATGGAGAGGAGGACGCCGGCGGGAGGGAAGACTCGATCCTCGTCAATGGGGCCTGCAGCGACCAGTCCTCCGACTCGCCCCCGATCCTGGAGGCCATCCGCACCCCGGAGATCAGAGGCCGAAGATCAAGCTCGCGACTCTCCAAGAGGGAGGTGTCCAGTCTGCTAAGCTACACGCAGGACTTGACAGGTGATGG------CGACGGGGAAGATGGGGATGGCTCTGACACCCCAGTGATGCCAAAGCTCTTCCGGGAAACCAGGACTCGGTCAGAAAGCCCAGCTGTCCGAACTCGAAATAACAACAGTGTCTCCAGCCGGGAGAGGCACAGGCCTTCCCCACGTTCCACCCGAGGCCAGCAGGGCCGCAACCATGTGGACGAGTCTCCCGTGGAGTTCCCGGCTACCAGGTCCCTGAGACGGCGGGCAACAGCATCGGCAGGAACGCCATGGCCGTCCCCTGCCAGCTCTTACCTTACCATCGACCTCACAGATGACACAGAGGACACATGTGTGACGCCCCAGAGCAGCAGTACCCCCTATGCCCGCCTAGCCCAGGACAGCCGGCAGGAGGGCATGGAGTC---CCCGCAGGTGGAGGCAGACAGTGGAGATGGAGACAGTTCAGAGTATCAGGATGGGAAGGAGTTTGGAATAGGGGACCTCGTGTGGGGAAAGATCAAGGGCTTCTCCTGGTGGCCTGCCATGGTGGTGTCTTGGAAGGCCACCTCTAAGCGACAGGCTATGTCTGGCATGCGGTGGGTCCAGTGGTTTGGTGATGGCAAGTTTTCCGAGGTCTCTGCAGACAAACTGGTGGCACTGGGGCTGTTCAGCCAGCACTTTAATTTGGCCACCTTCAATAAGCTCGTCTCCTATCGAAAAGCCATGTATCATGCTCTGGAGAAAGCTAGGGTGCGAGCTGGCAAGACCTTCCCCAGCAGCCCTGGAGACTCATTGGAGGACCAGCTGAAGCCCATGTTGGAGTGGGCCCACGGGGGCTTCAAGCCCACTGGGATCGAGGGCCTCAAACCCAACAACACGCAACCA------------------------------------------------------------GAGAGCAAGACTCGAAGACGCACAGCTGACGACTCAGCCACCTCTGACTACTGCCCCGCACCCAAGCGCCTCAAGACAAATTGCTATAACAACGGCAAAGACCGAGGGGATGAAGATCAGAGCCGAGAACAAATGGCTTCAGATGTTGCCAACAACAAGAGCATCCTGGAAGACGGCTGTTTGTCTTGTGGCAGAAAAAACCCCGTGTCCTTCCACCCTCTCTTTGAGGGGGGCCTCTGCCAGACATGCCGGGATCGCTTCCTTGAGCTGTTTTACATGTATGACGACGATGGCTATCAGTCTTACTGCACCGTGTGCTGCGAGGGCCGAGAGCTGCTGCTCTGCAGCAACATGAGCTGCTGCCGGTGCTTCTGTGTGGAGTGCCTGGAGGTGCTGGTGGGCACAGGCACAGCGGCCAAGGCCAAGCTTCAGGAGCCCTGGAGCTGCTACATGTGTCTCCCGCAGCGCTGTCACGGCGTCCTGCGGCGCCGGAAGGACTGGAACGTGCGCCTGCAGGCCTTCTTCACCAGTGACACGGGGCTTGAATACGAAGCCCCCAAGCTGTACCCTGCCATTCCCGCAGCCCGAAGGCGGCCCATTCGAGTCCTGTCATTGTTTGATGGCATCGCGACAGGCTACCTAGTCCTCAAAGAGCTGGGCATAAAGGTAGGAAAGTACATCGCCTCTGAAGTGTGTGAAGAGTCCATCGCTGTTGGAACCGTGAAGCACGAGGGGAATATCAAATACGTGAACGACGTGAGGAACATCACAAAGAAAAATATTGAAGAATGGGGCCCATTTGACTTGGTGATTGGCGGAAGCCCGTGCAACGATCTCTCAAATGTGAATCCAGCCAGGAAAGGCCTGTATGAGGGTACAGGCCGGCTCTTCTTTGAATTTTACCACCTGCTGAATTACTCACGCCCCAAGGAGGGTGATGACCGGCCGTTCTTCTGGATGTTTGAGAATGTTGTAGCCATGAAGGTTGGCGACAAGAGGGACATCTCGCGGTTCCTGGAGTGTAATCCAGTGATGATTGATGCCATCAAAGTTTCTGCTGCTCACAGGGCCCGATACTTTTGGGGCAACCTACCTGGGATGAACAGGCCCGTGATAGCATCAAAGAATGATAAACTCGAGCTGCAGGACTGCTTGGAATACAATAGGATAGCCAAGTTAAAGAAAGTACAGACAATAACCACCAAGTCGAACTCGATCAAACAGGGGAAAAACCAACTTTTCCCTGTTGTCATGAATGGCAAAGAAGATGTTTTGTGGTGCACTGAGCTCGAAAGGATCTTTGGCTTTCCTGTGCACTACACAGACGTGTCCAACATGGGCCGCGGTGCCCGCCAGAAGCTGCTGGGAAGGTCCTGGAGCGTGCCTGTCATCCGACACCTCTTCGCCCCTCTGAAGGAATACTTTGCATGTGAATAG

>SnubNoseGolden

ATGAAGGGAGACACCAGGCATCTCAATGGAGAGGAGGACGCCGGCGGGAGGGAAGACTCGATCCTCGTCAATGGGGCCTGCAGCGACCAGTCCTCCGACTCGCCCCCGATCCTGGAGGCCATCCGCACCCCGGAGATCAGAGGCCGAAGATCAAGCTCACGACTCTCCAAGAGGGAGGTGTCCAGTCTGCTAAGCTACACGCAGGACTTGACAGGCGATGG------CGACGGGGAAGATGGGGATGGCTCTGACACCCCAGTGATGCCAAAGCTCTTCCGGGAAACCAGGACTCGGTCAGAAAGCCCAGCTGTCCGAACTCGAAATAACAACAGTGTCTCCAGCCGGGAGAGGCATAGGCCTTCCCCACGTTCCACCCGAGGCCGGCAGGGCCGCAACCATGTGGACGAGTCTCCCGTGGAGTTCCCGGCTACCAGGTCCCTGAGGCGGCGGGCAACGGCATCGGCAGGAACGCCATGGCCGTCCCCTCCCAGCTCTTACCTTACCATCGACCTCACAGATGACACAGAGGACACACGTGTGACGCCCCAGAGCAGCAGTACCCCCTATGCCCGCCTAGCCCAGGATAGCCAGCAGGAGGGCATGGAGTC---CCCGCAGGTGGAGGCAGACAGTGGAGATGGAGACAGTTCAGAGTATCAGGATGGGAAGGAGTTTGGAATAGGGGACCTCGTGTGGGGAAAGATCAAGGGCTTCTCCTGGTGGCCTGCCATGGTGGTGTCTTGGAAGGCCACCTCCAAGCGACAGGCTATGTCTGGCATGCGGTGGGTCCAGTGGTTTGGCGATGGCAAGTTTTCCGAGGTCTCTGCAGACAAACTGGTGGCACTGGGGCTGTTCAGCCAGCACTTTAATTTGGCCACCTTCAATAAGCTCGTCTCCTATCGAAAAGCCATGTACCATGCTCTGGAGAAAGCTAGGGTGCGAGCTGGCAAGACCTTCCCCAGCAGCCCTGGAGACTCATTGGAGGACCAGCTGAAGCCCATGTTGGAGTGGGCCCACGGGGGCTTCAAGCCCACTGGGATCGAGGGCCTCAAACCCAACAACACGCAACCAGTGGTTAATAAGTCGAAGGTGCGTCGTGCAGGCAGTAGGAAATTAGAATCAAGGAAATACGAGAACAAGACTCGAAGACGCACAGCTGACGACTCAGCCACCTCTGACTACTGCCCTGCACCCAAACGCCTCAAGACAAATTGCTATAACAACGGCAAAGACCGAGGGGATGAAGATCAGAGCCGAGAACAAATGGCTTCAGATGTTGCCAACAACAAGAGCAGCCTGGAAGATGGCTGTTTGTCTTGTGGCAGGAAAAACCCTGTGTCCTTCCACCCTCTCTTTGAGGGGGGCCTCTGTCAGACATGCCGGGATCGCTTCCTTGAGCTGTTTTACATGTATGACGACGATGGCTATCAGTCTTACTGCACCGTGTGCTGTGAGGGCCGAGAGCTGCTGCTTTGCAGCAACACGAGCTGCTGCCGGTGCTTCTGTGTGGAGTGCCTGGAGGTGCTGGTGGGCACAGGCACAGCGGCCGAGGCCAAGCTTCAGGAGCCCTGGAGCTGCTACATGTGTCTCCCGCAGCGCTGTCATGGCGTCCTGCGGCGCCGGAAGGACTGGAACGTGCGCCTGCAGGCCTTCTTCACCAGTGACACGGGGCTTGAATACGAAGCCCCCAAGCTGTACCCTGCCATTCCTGCAGCCCGAAGGCGGCCCATTCGAGTCCTGTCATTGTTTGATGGCATCGCGACAGGCTACCTAGTCCTCAAAGAGTTGGGCATAAAGGTAGGAAAGTACGTCGCCTCTGAAGTGTGTGAGGAGTCCATCGCTGTTGGAACCGTGAAGCACGAGGGGAATATCAAATACGTGAACGACGTGAGGAACATCACAAAGAAAAATATTGAAGAATGGGGCCCATTTGACTTGGTGATTGGCGGAAGCCCATGCAACGATCTCTCAAATGTGAATCCAGCCAGGAAAGGCCTGTATGAGGGTACAGGCCGGCTCTTCTTCGAATTTTACCACCTGCTGAATTACTCCCGCCCCAAGGAGGGTGATGACCGGCCGTTCTTCTGGATGTTTGAGAATGTTGTAGCCATGAAGGTTGGCGACAAGAGGGACATCTCGCGGTTCCTGGAGTGTAATCCAGTGATGATTGATGCCATCAAAGTTTCTGCTGCTCACAGGGCCCGATACTTTTGGGGCAACCTACCTGGGATGAACAGGCCCGTGATAGCATCAAAGAATGATAAACTCGAGCTGCAGGACTGCTTGGAATACAATAGGATAGCCAAGTTAAAGAAAGTACAGACAATAACCACCAAGTCGAACTCGATCAAACAGGGGAAAAACCAACTTTTCCCTGTTGTCATGAATGGCAAAGAAGATGTTTTGTGGTGCACTGAGCTCGAAAGGATCTTTGGCTTTCCTGTGCACTACACAGACGTGTCCAACATGGGCCGCGGTGCCCGCCAGAAGCTGCTGGGAAGGTCCTGGAGCGTGCCTGTCATCCGACACCTCTTCGCCCCTCTGAAGGACTACTTTGCATGTGAATAG

>Colobus

ATGAAGGGAGACACCAGGCATCTCAATGGAGAGGAGGACGCCGGCGGGAGGGAAGACTCGATCCTCGTCAATGGGGCCTGCAGCGACCAGTCCTCCGACTCGCCCCCGATCCTGGAGGCCATCCGCACCCCGGAGATCAGAGGCCGAAGATCAAGCTCACGACTCTCCAAGAGGGAGGTGTCCAGTCTGCTAAGCTACACGCAGGACTTGACAGGTGATGG------CGACGGGGAAGATGGGGATGGCTCTGACACCCCAGTGATGCCAAAGCTCTTCCGGGAAACCAGGACTCGGTCAGAAAGCCCAGCTGTCCGAACTCGAAATAACAACAGTGTCTCCAGCCGGGAGAGGCACAGGCCTTCCCCACGTTCCACCCGAGGCCGGCAGGGCCGCAACCATGTGGATGAGTCTCCCGTGGAGTTCCCGGCTACCAGGTCCCTGAGGCGGCGGGCAACAGCATCGGCAGGAACACCATGGTCGTCCCCTCCCAGCTCTTACCTTACCATCGACCTCACAGACGACACAGAGGACACACGTGTGACTCCCCAGAGCAGCAGTACCCCCTATGCCCGCCTAGCCCAGGACAGCCAGCAGGAGGGCATGGAGTC---CCCGCAGGTGGAGGCAGACAGTGGAGATGGAGACAGTTCAGAGTATCAGGATGGGAAGGAGTTTGGAATAGGGGACCTCGTGTGGGGAAAGATCAAGGGCTTCTCCTGGTGGCCTGCCATGGTGGTGTCTTGGAAGGCCACCTCCAAGCGACAGGCTATGTCTGGCATGCGGTGGGTCCAGTGGTTTGGCGATGGCAAGTTTTCCGAGGTCTCTGCAGACAAACTGGTGGCACTGGGGCTGTTCAGCCAGCACTTTAATTTGGCCACCTTCAATAAGCTCGTCTCGTATCGAAAAGCCATGTACCATGCTCTGGAGAAAGCTAGGGTGCGAGCTGGCAAGACCTTCCCCAGCAGCCCTGGAGACTCATTGGAGGACCAGCTGAAGCCCATGTTGGAGTGGGCCCACGGGGGCTTCAAGCCCACTGGGATCGAGGGCCTCAAACCCAACAACACGCAACCAGTGGTTAATAAATCGAAGGTGCGTCGTGCAGGCAGTAGGAAATTAGAATCAAGGAAATACGAGAACAAGACTCGAAGACGCACAGCTGACGACTCAGCCACCTCTGACTACTGCCCTGCACCCAAGCGCCTCAAGACAAATTGCTATAACAACGGCAAAGACCGAGGGGATGAAGATCAGAGCCGAGAACAAATGGCTTCAGATGTTGCCAACAACAAGAGCAGCCTGGAAGATGGCTGTTTGTCTTGTGGCAGGAAAAACCCTGTGTCCTTCCACCCTCTCTTTGAGGGGGGCCTCTGTCAGACATGCCGGGATCGCTTCCTTGAGCTGTTTTACATGTATGACGACGATGGCTATCAGTCTTACTGCACCGTGTGCTGTGAGGGCCGAGAGCTGCTACTTTGCAGCAACACGAGCTGCTGCCGGTGCTTCTGTGTGGAGTGCCTGGAGGTGCTGGTGGGCACAGGCACAGCGGCCGAGGCCAAGCTTCAGGAGCCCTGGAGCTGCTACATGTGTCTCCCGCAGCGCTGTCATGGCGTCCTGCGGCGCCGGAAGGACTGGAACGTGCGCCTGCAGGCCTTCTTCACCAGTGACACGGGGCTTGAATACGAAGCCCCCAAGCTGTACCCTGCCATTCCCGCAGCCCGAAGGCGGCCCATTCGAGTCCTGTCATTGTTTGATGGCATCGCAACAGGCTACCTAGTCCTCAAAGAGTTGGGCATAAAGGTAGGAAAGTACGTCGCCTCTGAAGTGTGTGAGGAGTCCATCGCTGTTGGAACCGTGAAGCACGAGGGGAATATCAAATACGTGAATGACGTGAGGAACATCACAAAGAAAAATATTGAAGAATGGGGCCCATTTGACTTGGTGATTGGCGGAAGCCCATGCAACGATCTCTCAAATGTGAATCCAGCCAGGAAAGGCCTATATGAGGGTACAGGCCGGCTCTTCTTCGAATTTTACCACCTGCTGAATTACTCACGCCCCAAGGAGGGTGATGACCGGCCGTTCTTCTGGATGTTTGAGAATGTTGTAGCCATGAAGGTTGGCGACAAGAGGGACATCTCACGGTTCCTGGAGTGTAATCCAGTGATGATTGATGCCATCAAAGTTTCTGCTGCTCACAGGGCCCGATACTTTTGGGGCAACCTACCTGGGATGAACAGGCCCGTGATAGCATCAAAGAATGATAAACTCGAGCTGCAGGACTGCTTGGAATACAATAGGATAGCCAAGTTAAAGAAAGTACAGACAATAACCACCAAGTCGAACTCGATCAAACAGGGGAAAAACCAACTTTTCCCTGTTGTCATGAATGGCAAAGAAGATGTTTTGTGGTGCACTGAGCTCGAAAGGATCTTTGGCTTTCCTGTGCACTACACAGACGTGTCCAACATGGGCCGTGGTGCCCGCCAGAAGCTGCTGGGAAGGTCCTGGAGCGTGCCTGTCATCCGACACCTCTTCGCCCCTCTGAAGGACTACTTTGCATGTGAATAG

>RedColobus

ATGAAGGGAGACACCAGGCATCTCAATGGAGAGGAGGACGCCAGCGGGAGGGAAGACTCGATCCTCGTCAATGGGGCCTGCAGCGACCAGTCCTCCGACTCGCCCCCGATCCTGGAGGCCATCCGCACCCCGGAGATCAGAGGCCGAAGATCCAGCTCACGACTCTCCAAGAGGGAGGTGTCCAGTCTGCTAAGCTACACGCAGGACTTGACAGGTGATGG------TGACGGGGAAGATGGGGATGGCTCTGACACCCCAGTGATGCCAAAGCTCTTCCGGGAAACCAGGACTCGGTCAGAAAGCCCAGCTGTCCGAACTCGAAATAACAACAGTGTCTCCAGCCGGGAGAGGCACAGGCCTTCCCCACGTTCCACCCGAGGCCGGCAGGGCCGTAACCATGTGGATGAGTCTCCCGTGGAGTTCCCGGCTACCAGGTCCCTGAGGCGGCGGGCAACAGCATCGGCAGGAACACCATGGTCGTCCCCTCCCAGCTCTTACCTTACCATCGACCTCACAGACGACACAGAGGACACGCGTGTGACGCCCCAGAGGAGCAGTACCCCCTATGCCCGCCTAGCCCAGGACAGCCAGCAGGAGGGCATGGAGTCCCTCCTGCAGGTGGAGGCAGATGGTGGAGATGGAGACAGTTCGGAGTATCAGGATGGGAAGGAGTTTGGAATAGGGGACCTCGTGTGGGGAAAGATCAAGGGCTTCTCCTGGTGGCCTGCCATGGTGGTGTCTTGGAAGGCCACCTCCAAGCGACAGGCTATGTCTGGCATGCGGTGGGTCCAGTGGTTTGGCGATGGCAAGTTTTCCGAGGTCTCTGCAGACAAACTGGTGGCACTGGGGCTGTTCAGCCAGCACTTTAATTTGGCCACCTTCAATAAGCTCGTCTCCTATCGAAAAGCCATGTACCATGCTCTGGAGAAAGCTAGGGTGCGAGCTGGCAAGACCTTCCCCAGCAGCCCTGGAGACTCATTGGAGGACCAGCTGAAGCCCATGTTGGAGTGGGCCCACGGGGGCTTCAAGCCCACTGGGATCGAGGGCCTCAAACCCAACAACACGCAACCA------------------------------------------------------------GAGAACAAGACTCGAAGACGCACAGCTGATGACTCAGCCACCTCTGACTACTGCCCCGCACCCAAGCGCCTCAAGACAAATTGCTATAACAACGGCAAAGACCGAGGGGATGAAGATCAGAGCCGAGAACAAATGGCTTCAGATGTTGCCAACAACAAGAGCAGCCTGGAAGATGGCTGTTTGTCTTGTGGCAAGAAAAACCCGGTGTCCTTCCACCCTCTCTTTGAGGGGGGCCTCTGTCAGACATGCCGGGATCGCTTCCTTGAGCTGTTTTACATGTATGACGACGATGGCTATCAGTCTTACTGCACCGTGTGCTGTGAGGGCCGAGAGCTGCTACTTTGCAGCAACACAAGCTGCTGCCGGTGCTTCTGTGTGGAGTGCCTGGAGGTGCTGGTGGGCACAGGCACAGCGGCCGAGGCCAAGCTTCAGGAGCCCTGGAGCTGCTACATGTGTCTCCCGCAGCGCTGTCATGGCGTCCTGCGGCGCCGGAAGGACTGGAACGTGCGCCTTCAGGCCTTCTTCACCAGTGACACGGGGCTTGAATACGAAGCCCCCAAGCTGTACCCTGCCATTCCCGCAGCCCGAAGGCGGCCCATTCGAGTCCTGTCATTGTTTGATGGCATCGCGACAGGCTACCTAGTCCTCAAAGAGTTGGGCATAAAGGTAGGAAAGTACGTCGCCTCTGAAGTGTGTGAGGAGTCCATCGCTGTTGGAACCGTGAAGCACGAGGGGAATATCAAATACGTGAACGATGTGAGGAACATCACAAAGAAAAATATTGAAGAATGGGGCCCATTTGACTTGGTGATTGGCGGAAGCCCATGCAACGATCTCTCAAATGTGAATCCAGCCAGGAAAGGCCTGTATGAGGGTACAGGCCGGCTCTTCTTCGAATTTTACCACCTGCTGAATTACTCACGCCCCAAGGAGGGTGATGACCGGCCATTCTTCTGGATGTTTGAGAATGTTGTAGCCATGAAGGTTGGCGACAAGAGGGACATCTCGCGGTTCCTGGAGTGTAATCCAGTGATGATTGATGCCATCAAAGTTTCTGCTGCTCACAGGGCCCGATACTTTTGGGGCAACCTACCTGGGATGAACAGGCCCGTGATAGCATCAAAGAATGATAAACTCGAGCTGCAGGACTGCTTGGAATACAATAGGATAGCCAAGTTAAAGAAAGTACAGACAATAACCACCAAGTCGAACTCGATCAAACAGGGGAAAAACCAACTTTTCCCTGTTGTCATGAATGGCAAAGAAGATGTTTTGTGGTGCACTGAGCTCGAAAGGATCTTTGGCTTTCCTGTGCACTACACAGACGTGTCCAACATGGGCCGCGGTGCCCGCCAGAAGCTGCTGGGAAGGTCCTGGAGCGTGCCTGTCATCCGACACCTCTTCGCCCCTCTGAAGGACTACTTTGCATGTGAATAG

>Marmoset

ATGAAGGGAGACACCAGGCATCTCAATGGAGAGGAGGACGCCAGCGGGAGGGAAGACTCTATCCTCGTCAATGGGGCCTGCAGTGACCAGTCCTCCGACTCGCCCCCAATCCTGGAGGCCATCCGCACCCCCGAGATCAGAGGCCGAAGATCAAGCTTGAGACTCTCCAAGAGGGAGGTGTCCAGTCTGCTAAGCTATACGCAGGACTTGACAGGCAATGG------CGACGGGGAAGATGGGGATGGCTCTGACACCCCAGTGATGCCGAAGCTCTTCCGGGAAACCAGGACTCGGTCAGAAAGCCCAGCTGTCCGAACCCGAAATAACAACAGTGCCTCCAGCAGGGAGAGGCACAGGCCTTCCCCACGTTCCACCCGAGGCCGGCAGGCCCGCAGCCATGTGGAGGAGTCCCCCGTGGAGTTCCCGGCTACCAGGTCCCTGAGACGGCGGGCGACAGCATCGGCAGGCATGCCATGGTCGTCCCCTCCCAGCCCTTACCTCACCATCGACCTCACGGATGATACAGAGGACACAGATGTGATGCCCCAGAGCAGCAGTACCCCCTACGCCCGCCTAGCCCAGGACAACCATCAGGAGGGCATGGAGTC---CCCGCAGGTGGAGGCAGACAGTAGAGATGGAGACAGTTCAGAGTATCAGGATGGGAAGGAGTTTGGAATAGGGGACCTCGTGTGGGGGAAGATCAAGGGCTTCTCCTGGTGGCCTGCCATGGTGGTATCTTGGAAGGCCACCTCCAAGCGACAGGCCATGTCAGGCATGCGGTGGGTCCAGTGGTTTGGCGATGGCAAGTTCTCCGAGGTCTCTGCAGACAAACTGGTGGCACTGGGGCTGTTCAGCCAGCACTTTAATTTGGCCACCTTCAATAAGCTCGTCTCCTATCGAAAAGCCATGTACCATGCTCTGGAGAAAGCTAGGGTGCGAGCTGGCAAGACCTTCCCCAGCAGCCCTGGAGACTCATTGGAGGACCAGCTGAAGCCCATGCTGGAGTGGGCCCACGGGGGCTTCAAGCCCACTGGGATCGAGGGCCTCAAACCCAACAACACGCAACCAGTGGTTAATAAGTCGAAGGTGCGTCGTGCAGGCAGTAGGAAATTAGAATCAAAGAAATACGAGAACAAGACTCGAAGACGCACAGCTGACGACTCGGCCGCCTCTGACTACTGCCCCCCACCCAAGCGCCTCAAGACAAATTGCTATAACAATGGCAAAGACCGAGGGGATGAAGATCAGACCCGAGAACAAATGGCTTCAGACGTTGCCAACAACAAGAGCAGCCTGGAAGATGGCTGTTTGTCTTGTGGCAGGAAAAATCCCGTGTCCTTCCACCCTCTCTTTGAGGGGGGGCTCTGTCAGACATGCCGGGATCGGTTCCTTGAGCTGTTTTACATGTATGATGACGATGGCTATCAGTCTTACTGCACCGTGTGCTGTGAGGGCCGAGAGCTGCTGCTTTGCAGCAACACGAGCTGCTGCCGGTGCTTCTGTGTGGAGTGCTTGGAGGTGCTGGTGGGCACAGGCACAGCGGCTGAGGCCAAACTGCAGGAGCCCTGGAGCTGCTACATGTGTCTCCCGCAGCGCTGTCACGGCATCCTGCGGCGCCGAAAGGACTGGAACGTGCGCCTACAGGCCTTCTTCACCAGTGACACGGGGCTTGAATATGAAGCCCCCAAGCTGTACCCTGCCATTCCCGCAGCCCGAAGGCGGCCCATTCGAGTCCTGTCTCTGTTTGATGGAATCGCAACAGGCTACCTAGTCCTCAAAGAGTTGGGCATAAAGGTGGGAAAGTACGTCGCCTCCGAAGTGTGTGAGGAGTCCATCGCTGTTGGAACCGTGAAGCACGAGGGGAATATCAAATATGTGAACGACGTGAGGAACATCACAAAGAAAAATATTGAAGAATGGGGCCCATTTGACTTGGTGATTGGTGGAAGCCCATGCAATGATCTCTCAAATGTGAATCCAGCCAGGAAAGGCCTATATGAGGGCACAGGCCGGCTCTTCTTCGAATTTTACCACCTGCTGAATTACTCACGCCCCAAGGAGGGTGATGACCGGCCGTTCTTCTGGATGTTTGAGAATGTTGTAGCCATGAAGGTTGGCGACAAGAGGGACATCTCGCGGTTCCTGGAGTGTAATCCAGTGATGATTGATGCCATCAAAGTTTCTGCTGCTCACAGGGCCCGATATTTTTGGGGCAACCTTCCTGGGATGAACAGGCCGGTGATAGCATCAAAGAATGATAAACTCGAGCTGCAGGACTGCTTGGAATACAATAGGATAGCCAAGTTAAAGAAAGTACAGACAATAACCACCAAGTCGAACTCAATCAAACAGGGGAAAAACCAACTTTTCCCTGTTGTCATGAATGGCAAAGAAGATGTTTTGTGGTGCACTGAGCTCGAAAGGATCTTTGGCTTTCCTGTGCACTACACAGATGTTTCCAACATGGGCCGTGGCGCCCGCCAGAAGCTGCTGGGAAGGTCCTGGAGTGTGCCTGTCATCCGACACCTCTTCGCCCCTCTGAAGGACTACTTTGCATGTGAATAG

>OwlMonkey

ATGAAGGGAGACACCAGGCATCTCAATGGAGAGGAGGACGCCGGCGGGAGGGAAGACTCGATCCTCGTCAATGGGGCCTGCAGTGACCAGTCCTCCGACTCGCCCCCAATCCTGGAGGCCATCCGCACCCCCGAGATCAGAGGCCGAAGATCAAGCTTGAGACTCTCCAAGAGGGAGGTGTCCAGTCTGCTAAGCTATACGCAGGACTTGACAGGCAATGG------CGACGGGGAAGATGGCGATGGCTCTGACACCCCAGTGATGCCGAAGCTCTTCCGGGAAACCAGGACTCGGTCAGAAAGCCCAGCTGTCCGAACCCGAAATAACAACAGTGCCTCCAGCAGGGAGAGGCACAGGCCTTCCCCACGTTCCACCCGAGGCCGGCAGGGCCGCAGCCATGTGGAGGAGTCCCCCGTGGAGTTCCCGGCTACCAGGTCCCTGAGACGACGGGCGACAGCATCGGCAGGCACGCCATGGTCGTCCCCTCCCAGCCCTTACCTCACCATCGACCTCACAGACGATACAGAGGACACAGATGTGATGCCCCAGAGCAGCAGTACCCCCTACGCCCGCCTAGCCCAGGACAGCCAGCAGGAGGGCATGGAGTC---CCCGCAGGTGGAGGCAGACAGTAGAGATGGAGACAGTTCAGAGTATCAGGATGGGAAGGAGTTTGGAATAGGGGACCTTGTGTGGGGGAAGATCAAGGGCTTCTCCTGGTGGCCCGCCATGGTGGTATCTTGGAAGGCCACCTCCAAGCGACAGGCCATGTCGGGCATGCGGTGGGTCCAGTGGTTTGGCGATGGCAAGTTCTCCGAGGTCTCTGCAGACAAACTGGTGGCACTGGGGCTGTTCAGCCAGCACTTTAATTTGGCCACCTTCAATAAGCTCGTCTCCTATCGAAAAGCCATGTACCATGCTCTGGAGAAAGCTAGGGTTCGAGCTGGCAAGACCTTCCCCAGCAGCCCTGGAGACTCATTGGAGGACCAGCTGAAGCCCATGCTGGAGTGGGCCCACGGGGGCTTCAAGCCCACTGGGATCGAGGGCCTCAAACCCAACAACACGCAACCAGTGGTTAATAAGTCGAAGGTGCGTCGTGCAGGCAGTAGGAAATTAGAATCAAAGAAATACGAGAACAAGACTCGAAGACGCACAGCTGACGACTCAACTGCCTCTGACTACTGTCCCCCACCCAAGCGCCTCAAGACAAATTGCTATAACAACGGCAAAGACCGAGGGGATGAAGATCAGACCCGAGAACAAATGGCTTCAGATGTTGCCAGCAACAAGAGCAGCCTGGAAGATGGCTGTTTGTCTTGTGGCAGGAAAAATCCTGCGTCCTTCCACCCTCTCTTTGAGGGGGGGCTCTGTCAGACATGCCGGGATCGGTTCCTCGAGCTGTTTTACATGTATGATGACGATGGCTATCAGTCTTACTGCACCGTGTGCTGCGAGGGCCGAGAACTGCTGCTTTGCAGCAACACGAGCTGCTGCCGGTGCTTCTGTGTGGAGTGCTTGGAGGTGCTGGTGGGTACAGGCACAGCGGCCGAGGCCAAGCTGCAGGAGCCCTGGAGCTGCTACATGTGTCTCCCGCAGCGCTGTCACGGCATCCTGCGGCGCCGGAAGGACTGGAACATGCGCCTACAGGCCTTCTTCACCAGTGACACGGGGCTTGAATACGAAGCCCCCAAGCTGTACCCTGCCATTCCCGCAGCCCGAAGGCGGCCCATTCGAGTCCTGTCTCTGTTTGATGGAATTGCAACAGGCTACCTAGTCCTCAAAGAGTTGGGCATAAAGGTGGGAAAGTACGTCGCCTCTGAAGTGTGTGAGGAGTCCATTGCTGTTGGAACCGTGAAGCACGAGGGAAATATCAAATATGTGAATGACGTGAGGAACATCACAAAGAAAAATATTGAAGAATGGGGCCCATTTGACTTGGTGATTGGTGGAAGCCCATGCAACGATCTCTCAAATGTGAATCCAGCCAGGAAAGGCCTATATGAGGGCACAGGCCGGCTCTTCTTCGAATTTTACCACCTGCTGAATTACTCACGCCCCAAGGAGGGTGATGACCGGCCGTTCTTCTGGATGTTTGAGAATGTTGTAGCCATGAAGGTTGGCGACAAGAGGGACATCTCACGGTTCCTGGAGTGTAATCCAGTGATGATTGATGCCATCAAAGTTTCTGCTGCTCACAGGGCCCGATATTTTTGGGGCAACCTTCCTGGGATGAACAGGCCCGTGATAGCATCAAAGAATGATAAACTCGAGCTGCAGGACTGCTTGGAATACAATAGGATAGCCAAGTTAAAGAAAGTACAGACAATAACCACCAAGTCGAACTCAATCAAACAGGGGAAAAACCAACTTTTCCCTGTTGTCATGAATGGCAAAGAAGATGTTTTGTGGTGCACTGAGCTCGAAAGGATCTTTGGCTTTCCTGTGCACTACACAGATGTTTCCAACATGGGCCGTGGCGCCCGCCAGAAGCTGCTGGGAAGGTCCTGGAGTGTGCCTGTCATCCGACACCTCTTCGCCCCTCTGAAGGACTACTTTGCATGTGAATAG

>Sapajou

ATGAAGGGAGACACCAGGCATCTCAGTGGAGAGGAGGACGCCGGCGGGAGGGAAGACTCCATCCTCGTCAATGGGGCCTGCAGTGACCAGTCCTCCGACTCGCCCCCAATCCTGGAGGCCATCCGCACCCCCGAGATCAGAGGCCGAAGATCAAGCTTGAGACTCTCCAAGAGGGAGGTGTCCAGTCTGCTAAGCTATACGCAGGACTTGACAGGCAATGG------CGACGGGGAAGATGGGGATGGCTCTGACACCCCAGTGATGCCGAAGCTCTTCCGGGAAACCAGGACTCGGTCAGAAAGCCCAGCTGTCCGAACCCGAAATAACAACAGTGCCTCCAGCAGGGAGAGGCACAGGCCTTCCCTACGTTCCACCCGAGGCCGGCAGGGCCGCAGCCATGTGGAGGAGTCCCCCGTGGAGTTCCCGGCTACCAGGTCCCTGAGACGGCGGGCGACAGCATCGGCAGGCATTCCATGGTCGTCCCCTCCCAGCCCTTACCTCACCATCGACCTCACAGACGATACAGAGGACACAGATGTGATGCCCCAGAGCAGCAGTACCCCCTACACCCGCCTAGCCCAGGACAGCCAGCAGGAGGGCATGGAGTC---CCCGCAGGTGGAGGCAGACAGTAGAGACGGAGACAGTTCAGAGTATCAGGATGGGAAGGAGTTTGGAATAGGGGACCTCGTGTGGGGGAAGATCAAGGGCTTCTCCTGGTGGCCCGCCATGGTGGTATCTTGGAAGGCCACCTCCAAGCGACAGGCCATGTCGGGCATGCGGTGGGTCCAGTGGTTTGGCGATGGCAAGTTCTCCGAGGTCTCTGCAGACAAACTGGTGGCACTGGGGCTGTTCAGCCAGCACTTTAATTTGGCCACCTTCAATAAGCTCGTCTCCTATCGAAAAGCCATGTACCATGCTCTGGAGAAAGCTAGGGTGCGAGCTGGCAAGACCTTCCCCAGCAGCCCTGGAGACTCATTGGAGGACCAGCTGAAGCCCATGCTGGAGTGGGCCCACGGGGGCTTCAAGCCCACTGGGATCGAGGGCCTCAAACCCAACAACACGCAACCAGTGGTTAATAAGTCGAAGGTGCGTCGTGCAGGCAGTAGGAAATTAGAATCAAAGAAATACGAGAACAAGACTCGAAGACGCACAGCTGACGACTCAGCCGCCTCTGACTACTGCCCCCAACCCAAGCGCCTCAAGACAAATTGCTATAACAACGGCAAAGACCGAGGGGATGAAGATCAGACCCGAGAACAAATGGCTTCAGATGTTGCCAACAACAAGAGCAGCCTGGAAGATGGCTGTTTGTCTTGTGGCAGGAAAAATCCCGTGTCCTTTCACCCTCTCTTTGAGGGGGGCCTCTGTCAGACATGCCGGGATCGGTTCCTCGAGCTGTTTTACATGTATGATGATGATGGCTATCAGTCTTACTGTACTGTGTGCTGTGAGGGCCGAGAGCTGCTGCTCTGCAGCAACACAAGCTGCTGCCGGTGCTTCTGTGTGGAGTGCTTGGAGGTGCTGGTGGGCACAGGCACCGCGGCCGAGGCCAAGCTGCAGGAGCCCTGGAGCTGCTACATGTGTCTCCCGCAGCGCTGTCACGGCATCCTGCGGCGCCGGAAGGACTGGAACGTGCGCCTACAGGCCTTCTTCACCAGTGACACGGGGCTTGAATATGAAGCCCCCAAGCTGTACCCTGCCATTCCCGCAGCCCGAAGGCGGCCCATTCGAGTCCTGTCTCTGTTTGATGGAATCGCAACAGGCTACCTAGTCCTCAAAGAGTTGGGCATAAAGGTGGGAAAGTACGTCGCCTCCGAAGTGTGTGAGGAGTCCATCGCTGTTGGAACCGTGAAGCACGAGGGGAATATCAAATACGTGAACGATGTGAGGAACATCACAAAGAAAAATATTGAAGAATGGGGCCCATTTGACTTGGTGATTGGTGGAAGCCCATGCAATGATCTCTCAAATGTGAATCCAGCCAGGAAAGGCCTATATGAGGGCACAGGCCGGCTCTTCTTTGAATTTTACCACCTGCTGAATTACTCACGCCCCAAGGAGGGTGATGACCGGCCGTTCTTCTGGATGTTTGAGAATGTTGTAGCCATGAAGGTTGGCGACAAGAGGGACATCTCGCGGTTCCTGGAGTGTAATCCAGTGATGATTGATGCCATCAAAGTTTCTGCTGCTCACAGGGCCCGATATTTTTGGGGCAACCTTCCTGGGATGAACAGGCCCGTGATAGCATCAAAGAATGATAAACTCGAGCTGCAGGACTGCTTGGAATACAATAGGATAGCCAAGTTAAAGAAAGTACAGACAATAACCACCAAGTCGAACTCAATCAAACAGGGGAAAAACCAACTTTTCCCTGTTGTCATGAATGGCAAAGAAGATGTTTTGTGGTGCACTGAGCTCGAAAGGATCTTTGGCTTTCCTGTGCACTACACAGATGTTTCCAACATGGGCCGTGGCGCCCGCCAGAAGCTGCTGGGAAGGTCCTGGAGTGTGCCTGTCATCCGACACCTCTTCGCCCCTCTGAAGGACTACTTTGCATGCGAATAG

>Sifaka

ATGAAGGGAGACACCAGACATCTCAATGGAGAGGAGGATGCCAGCGGGAGGGAGGACTCAATTATCATCAACGGGGCCTGCAGCGACCATTCCTCAGACTCGCCCCCCATCCTGGAGGCTATCCACACCCCGGAGATCAGAGGCCGCAGGTCAAGCTCACGACTGTCCAAGAGGGAGGTCTCCAGTCTGCTAAGCTACACTCAGGATCTGACAGGAGATGGAGATGCCGAAGGGGAGGATGGGGATGGCTCCGACACTCCGGTGATGCCAAAACTCTTCCGTGAAACCAGGACTCGGTCTGAAAGCCCAGCTGTCCGAACCCGAAATAACAACAGTGCCTCCAGCCGGGAGAGGCACAGGCCCTCCCCACGTGCCACCCGAGGCCGGCAGGGCCGCAGTCATGTGGACGAGTCCCCCGTGGAGTTCCCAGCTACCAGGTCCCTGAGGCGGCGGGCAGCAGCATCGGCAGGCACGCCGTGGCCGTCCCCTGCCAGCCCTTACCTCACCATCGACCTCACAGATGACACCGATGACACAGATGTGACACCCCAGAGCAGCAGTACCCCCTACACCCGCCTAGCCCAGGACAGCCAGCAGGAGAGCTTGGAGTC---TCCACAGGTGGATGCAGAAAGGAGAGACGCAGACAGTGCAGAGTATCAGGATGGGAAGGAGTTTGGAATAGGGGACCTCGTGTGGGGAAAGATCAAGGGCTTCTCCTGGTGGCCTGCCATGGTAGTGTCCTGGAAGGCCACCTCCAAGCGACAGGCCATGTCCGGCATGCGATGGGTCCAGTGGTTTGGCGATGGCAAGTTCTCAGAGGTCTCTGCGGACAAACTGGTGGCCCTGGGGCTGTTCAGCCAGCACTTTAACCTGGCCACCTTCAATAAGCTGGTTTCTTACAGGAAGGCCATGTACCATGCCCTGGAGAAAGCCAGGGTGCGAGCTGGCAAGACCTTCCCCACCAGTCCTGGAGACTCATTGGAGGACCAGCTGAAGCCCATGTTGGAGTGGGCCCACGGGGGCTTCAAGCCCACTGGGATCGAGGGCCTCAGACCCAACAACAAGCAACCAGTGGTTAATAAGTCGAAGGTGCGTCGTGCAGGCAGTAGGAACTTAGAATCAAGGAAATACGAGAACAAGACAAGAAGACGCACAGCTGACGACTCGGCCACTTCTGACTACTGCCCTCCACCCAAGCGCCTCAAGACAAATTGCTACAACAATGGAAAAGACCGCGGGGACGAGGACCAGAGCCGAGAACAAATGGCATCTGATGTCACCAACAACAAGGGTAACCTGGAAGACAGCTGCCTATCTTGTGGCAAGAAGAACTCTGTGTCGTTCCACCCTCTCTTTGAGGGGGGCCTCTGCCAGACATGCCGGGATCGGTTCTTGGAGTTGTTCTACATGTACGACGACGATGGCTACCAGTCTTACTGCACCGTGTGCTGTGAGGGTCGGGAGCTGCTTCTCTGCAGCAACACGAGCTGCTGCCGGTGCTTTTGTGTGGAGTGCCTGGAGGTGCTGGTTGGCTCAGGCACTGCGGCAGAGGCCAAGCTTCAGGAGCCATGGAGTTGCTATATGTGTCTGCCACAGCGCTGTCACGGTGTCCTGCGGCGCAGGAAGGACTGGAATGTGCGCCTGCAGGCCTTCTTCACCAGTGATATGGGGCGAGAATACGAAGCCCCCAAGGTGTACCCTGCGATTCCTGCAGCCCGAAGGAGGCCCATTCGAGTTCTGTCTCTGTTTGATGGTATTGCAACAGGGTACTTGGTCCTCAAAGAATTGGGCATAAAGGTGGAAAAGTATGTCGCCTCTGAAGTGTGTGAAGAGTCCATCGCCGTTGGAACTGTCAAGCATGAGGGGAATATCAAATATGTGAACGACGTGAGAAACATCACAAAGAAAAATATTGAAGAATGGGGCCCCTTTGACTTGGTGATCGGTGGAAGCCCATGCAATGATCTCTCAAATGTGAATCCTGCCAGAAAAGGCTTGTACGAGGGGACAGGCCGGCTCTTCTTTGAGTTCTACCACCTGCTGAATTACTCGCGCCCCAAGGAGGGGGATGACCGGCCGTTCTTCTGGATGTTTGAGAACGTCGTAGCCATGAAGGTCGGCGACAAGAGGGACATCTCAAGGTTCCTGGAGTGCAATCCAGTGATGATCGATGCCATAAAAGTATCTGCTGCTCACAGGGCACGATATTTCTGGGGCAACCTACCCGGGATGAACAGGCCCGTGATAGCATCAAAGAATGATAAACTCGAGCTTCAAGACTGCTTGGAATACAATAGGACAGCAAAGTTAAAGAAAGTACAGACAATAACCACCAAGTCGAACTCGATCAGACAGGGGAAAAACCAACTTTTCCCTGTTGTCATGAATGGCAAAGAAGATGTTTTGTGGTGCACTGAGCTCGAAAGGATCTTCGGCTTTCCTGTACACTACACAGATGTGTCCAACATGGGCCGAGGCGCCCGCCAGAAGCTGCTCGGGAGGTCCTGGAGTGTGCCAGTCATCCGACACCTCTTCGCCCCCCTGAAGGACTACTTTGCATGTGAATAG

>grayMouseLemur

ATGAAGGGAGACACCAGACATCTCAATGGAGAGGAGGACGCCAGCGGGAGGGAGGACTCAATTGTCATCAACGGGGCCTGCAGCGACCATTCCTCAGACTCGCCCCCCATCCTGGAGGCTATCCGCACCCCGGAGATCAGAGGTCGCAGGTCAAGCTCACGGCTGTCCAAGAGGGAGGTCTCCAGTCTGCTAAGCTACACTCAGGATCTGACTGGAGATGGAGATGGCGACGGGGAGGATGGGTACGGCTCCGACACTCCGGTGATGCCAAAACTCTTCCGTGAAACCAGGACTCGGTCTGAAAGCCCCGCTGTCCGAACCCGAAATAACACCAGTGCCTACAGCCGGGAGAGGCTCAGGCCCTCCATACGTGCCACCCGAGGCCGGCAGGGCCGCAGTCATATGGACGAGTCCCCCGTGGAGTTCCCGGCTACCAGGTCCCTGAGGCGGCGGGCAGCAGCATCAGCAGGCACGCCGTGGCCGTCCCCTGCCAGCCCTTACCTCACCATCGACCTCACAGATGACACCGATGACACAGATGTGACACCCCAGAGCAGCAGTACCCCCTACACCCGCCTAGCCCAGGACAGCCAGCAGGAGGGCTTGGAGTC---CCCACAGGTGGATGCAGAAA---GAGACGTAGACAGTGCCGAGTATCAGGATGGGAAGGAGTTTGGAATAGGGGACCTCGTGTGGGGAAAGATCAAGGGCTTCTCCTGGTGGCCTGCCATGGTAGTGTCCTGGAAGGCCACCTCCAAGCGACAGGCCATGTCCGGCATGCGATGGGTCCAGTGGTTCGGTGATGGCAAATTCTCAGAGGTCTCTGCGGACAAACTGGTGGCCCTGGGGCTGTTCAGTGAGCACTTTAACCTGGCCACCTTCAATAAGCTGGTTTCTTACAGGAAGGCCATGTACCATGCCCTGGAGAAAGCCAGGGTACGAGCTGGCAAGACCTTCCCCACCAGCCCTGGAGACTCACTGGAGGACCAGCTGAAGCCCATGTTGGAGTGGGCCCACGGGGGCTTCAAGCCCACTGGGATCGAGGGCCTAAGACCCAGCAACAAGCAACCAGTGGTTAATAAGTCGAAGGTGCGTCGTGCAGGCAGTAGGAAATTAGAATCAAGGAAATACGAGAACAAGACAAGAAGACGCACAGCTGACGACTCGGCCGCTTCTGACTACTGCCCTCCATCCAAGCGCCTCAAGACAAATTGCTACAAC---GGAAGAGACCGGGGGGACGAGGACCAGAGCCGAGAACAAATGGCATCTGATGTCACCAACAACAAGGGTAACCTGGAAGACAGCTGCTTGTCTTGCGGTAGGAAGAACTCTGTGTCCTTCCACCCTCTCTTTGAGGGGGGCCTCTGCCAGACATGCCGGGATCGGTTCTTGGAGTTGTTCTACATGTACGACGACGACGGCTACCAGTCCTACTGCACCGTGTGCTGTGAGGGTCGGGAGCTGCTTCTCTGCAGCAACACGAGCTGCTGCCGGTGCTTTTGTGTGGAGTGCCTGGAGGTGCTGGTTGGCTCAGGCACTGCAGCAGAGGCCAAGCTTCAGGAGCCGTGGAGTTGCTACATGTGTCTGCCACAGCGCTGCCACGGAGTACTGCGGCGCAGGAAGGACTGGAATGTGCGCCTGCAGGCCTTCTTCACCAGCGACATGGGGCGAGAATACGAAGCCCCCAAGGTGTACCCTGCGATCCCTGCAGCCCGAAGGAGGCCCATTCGAGTTCTGTCTCTGTTTGATGGAATCGCAACAGGGTACTTGGTCCTCAAGGAATTGGGCATAAAGGTTGAGAAGTACGTCGCCTCTGAAGTGTGTGAAGAGTCCATCGCTGTCGGAACTGTTAAGCATGAGGGGAATATCAAATATGTGAACGACGTGAGAAACATCACAAAGAAAAATATTGAAGAATGGGGCCCATTTGACTTGGTGATCGGTGGAAGCCCATGCAACGATCTCTCAAATGTGAATCCTGCCAGAAAAGGCTTGTACGAGGGGACAGGCCGGCTCTTCTTTGAGTTCTACCACCTGCTGAATTACTCACGTCCCAAAGAGGGGGATGACCGACCCTTCTTCTGGATGTTTGAGAACGTTGTAGCCATGAAGGTCGGCGACAAGAGGGACATCTCGAGGTTCCTGGAGTGCAATCCAGTGATGATTGATGCCATAAAAGTATCTGCTGCTCACAGGGCACGATATTTCTGGGGCAACCTACCCGGGATGAACAGGCCCGTGATAGCATCAAAGAATGATAAACTCGAGCTTCAAGACTGCTTGGAATACAATAGGACAGCAAAGTTAAAGAAAGTACAGACAATAACCACCAAGTCGAACTCGATCAGACAGGGGAAAAACCAACTTTTCCCCGTTGTCATGAATGGCAAAGAAGATGTTTTGTGGTGCACTGAGCTCGAAAGGATCTTCGGCTTTCCTGTACACTACACAGATGTGTCCAACATGGGCCGAGGCGCCCGCCAGAAGCTGCTCGGGAGGTCCTGGAGTGTGCCAGTCATCAGACACCTCTTCGCCCCCCTGAAGGACTACTTTGCATGTGAATAG

>Otolemur

ATGAAGGGAGACACCAGACATCTTAATGGAGAGGAGGACGTCAGCGGGAGGGAGGACTCTATCATCATCAACGGGGCCTGCAGTGACCATTCCTCGGACTCGCCCCCAATCCTAGAGGCTATCCGCACCCCGGAGATCAGAGGCCGCAGGTCCAGTGCACGGCTGTCCAAGAGGGAAGTCTCCAGTCTGCTAAGCTACACTCAGGATCTGACGGGCGATGGAGATGGAGAAGGGGAGGACGGGGATGGCTCCGACACACCAGTGATGCCAAAGCTCTTCCGTGAAACCAGGACTCGTTCTGAAAGCCCAGCTGTCCGAACCCGAAATAGCAACAGTGCCTCCAGGTGGGAGAGGCACAGGCCCTCCCCACGTACTACCCGAGGCCGACAGGGCCGCAATCATGTGGACGAGTCCCCTGTGGAGTTCCCGGCTACCAGGTCCCTGAGGCGGCGGGCAACAGCATCGGTTGGCACACCGTGGTTATCCCCTGCCAGCCCTTACCTCACCATTGACCTCACAGATGACACGAATGACGCTGATGTGACACCCCAGAGCAGCAGTACCCCCTACTCCCACCTAGGCCAGGATAGCCAGCAGGATAGCTTCGAGTC---CCCACAGGTGGATGCAGAAAGAGGAGACACAGACAGTGCTGAGTATCAGGATGGAAAGGAATTTGGAATAGGGGACCTTGTGTGGGGAAAGATCAAGGGCTTCTCCTGGTGGCCTGCCATGGTAGTGTCATGGAAGGCCACCTCCAAGCGACAGGCCATGTCCGGCATGCGATGGGTCCAGTGGTTTGGTGATGGCAAGTTCTCTGAGGTCTCTGCAGATAAACTGGTGGCCCTGGGGCTGTTCAGCCAGCACTTTAACCTGGCTACCTTCAATAAGCTGGTTTCTTACAGGAAGGCCATGTATCATGCCCTGGAGAAAGCCAGGGTGAGAGCTGGCAAGACATTCCCTAGCAGCCCTGGAGACTCATTGGAGGACCAGCTGAAGCCCATGCTGGAGTGGGCCCACGGGGGCTTCAAGCCCACTGGCATCGAGGGCCTCAAACCCAACAACAACCAACCA------------------------------------------------------------GAGAACAAGACAAGAAGACGAACAGCTGATGACTCAGCTACCTCTGACTATTGTCTTCCACCCAAGCGCCTCAAGACAAATTGCTACAACAATGGCAAAGACCGTGGGGAGGAAGACCAGAGCCGAGAACAAATGGCATCAGATGTCACCAACAACCAGAGCAATCTAGAAGACAGCTGCCTGTCTTGTGGCAAGAAGAACTCCGTGTCATTCCACCCTCTCTTTGAGGGGGGGCTCTGTAAGACATGCAGGGATCGGTTCTTGGAGTTGTTCTACATGTACGACGACGATGGCTATCAGTCTTACTGCACCGTGTGCTGTGAGGGCTACGAGCTGCTGCTCTGCAGTAACACAAGCTGCTGTCGGTGCTTCTGCGTGGAGTGCTTGGAGGTGCTGGTAGGCCCCGGCACAGCAGCGGAGGCCAAGCTCCAGGAGCCCTGGAGCTGCTACATGTGTCTGCCACAGCGCTGTCACGGCATCTTGCGGCGCCGGAAGGACTGGAACATGCGCCTGCAGGCTTTCTTCACCAGCGACATGGGGCATGAATATGAAGCCCCCAAGCTGTACCCTGCGATTCCTGCAGCCCGAAGGCGGCCTATTCGAGTTCTGTCTCTGTTTGATGGAATCGCAACAGGCTACTTGGTCCTCAAAGAATTGGGCATAAAGGTGGAAAAGTATGTTGCCTCTGAAGTGTGTGAAGAGTCCATTGCTGTTGGAACCGTTAAGCATGAGGGGAATATCAAATATGTGAACGACGTGAGAAACATCACTAAGAAAAATATTGAAGAATGGGGCCCATTTGACTTGGTGATTGGTGGAAGCCCATGCAATGATCTCTCAAATGTGAATCCTGCTAGGAAAGGCTTGTATGAGGGGACAGGTCGGCTCTTCTTTGAGTTCTACCACCTGCTGAATTACTCGCGCCCCAAGGAGGGTGATGACCGACCATTCTTCTGGATGTTTGAGAATGTAGTAGCCATGAAGGTCGGAGACAAGAGGGACATCTCTCGGTTCTTGGAGTGCAATCCAGTGATGATTGATGCCATAAAAGTATCTGCTGCTCACAGGGCACGATATTTCTGGGGCAACCTTCCTGGGATGAACAGGCCGGTGATAGCATCAAAGAATGATAAACTCGAGCTTCAAGACTGCTTGGAATACAATAGGACAGCAAAGTTAAAGAAAGTACAGACAATAACCACCAAGTCGAACTCGATCAGACAGGGGAAAAACCAACTTTTCCCTGTTGTCATGAATGGCAAAGAAGATGTTTTGTGGTGCACTGAGCTCGAAAGGATCTTCGGCTTTCCTGTACACTACACAGATGTGTCCAACATGGGCCGAGGTGCCCGCCAGAAGCTGCTTGGGAGGTCCTGGAGTGTGCCTGTGATCCGACACCTCTTCGCCCCCCTGAAGGACTACTTTGCATGTGAATAG
