## Supplementary material for "Dynamic evolution of *de novo* DNA methyltransferases in rodent and primate genomes": Suppl data 8

*SupFile_8: Primate* DNMT3L *sequence alignments*

>Human3L

GCGGCCATCCCAGCCCTGGACCCAGAGGCCGAGCCCAGCATGGACGTGATTTTGGTGGGATCCAGTGAGCTCTCAAGCTCCGTTTCACCCGGGACAGGCAGAGATCTTATTGCATATGAAGTCAAGGCTAACCAGCGAAATATAGAAGACATCTGCATCTGCTGCGGAAGTCTCCAGGTTCACACACAGCACCCTCTGTTTGAGGGAGGGATCTGCGCCCCATGTAAGGACAAGTTCCTGGATGCCCTCTTCCTGTACGACGATGACGGGTACCAATCCTACTGCTCCATCTGCTGCTCCGGAGAGACGCTGCTCATCTGCGGAAACCCTGATTGCACCCGATGCTACTGCTTCGAGTGTGTGGATAGCCTGGTCGGCCCCGGGACCTCGGGGAAGGTGCACGCCATGAGCAACTGGGTGTGCTACCTGTGCCTGCCGTCCTCCCGAAGCGGGCTGCTGCAGCGTCGGAGGAAGTGGCGCAGCCAGCTCAAGGCCTTCTACGACCGAGAGTCGGAGAATCCCCTTGAGATGTTCGAAACCGTGCCTGTGTGGAGGAGACAGCCAGTCCGGGTGCTGTCCCTTTTTGAAGACATCAAGAAAGAGCTGACGAGTTTGGGCTTTTTGGAAAGTGGTTCTGACCCGGGACAACTGAAGCATGTGGTTGATGTCACAGACACAGTGAGGAAGGATGTGGAGGAGTGGGGACCCTTCGATCTTGTGTACGGCGCCACACCTCCCCTGGGCCACACCTGTGACCGTCCTCCCAGCTGGTACCTGTTCCAGTTCCACCGGCTCCTGCAGTACGCACGGCCCAAGCCAGGCAGCCCCAGGCCCTTCTTCTGGATGTTCGTGGACAATCTGGTGCTGAACAAGGAAGACCTGGACGTCGCATCTCGCTTCCTGGAGATGGAGCCAGTCACCATCCCAGATGTCCACGGCGGATCCTTGCAGAATGCTGTCCGCGTGTGGAGCAACATCCCAGCCATAAGG---AGCAGGCACTGGGCTCTGGTTTCGGAAGAAGAATTGTCCCTGCTGGCCCAGAACAAGCAGAGCTCGAAGCTCGCGGCCAAGTGGCCCACCAAGCTGGTGAAGAACTGCTTTCTCCCCCTAAGAGAATATTTCAAGTATTTTTCAACAGAACTCACTTCCTCTTTA

>Gorilla3L

GCGGCCATCCCAGCCCTGGACCCAGAGGCCGAGCCCAGCATGGACGTGATTTTGGTGGGATCCAGTGAGCTCTCAAGCTCCGTTTCACCCGGGACAGGCAGAGATCTTATTGCATATGAAGTCAAGGCTAACCAGCGAAATATAGAAGACATCTGCATCTGCTGCGGAAGTCTCCAGGTTCACACACAGCACCCTCTGTTTGAGGGAGGGATCTGCGCCCCGTGTAAGGACAAGTTCCTGGATGCCCTCTTCCTGTACGACGATGACGGGTACCAATCCTACTGCTCCATCTGCTGCTCTGGAGAGACGCTGCTCATCTGCGGAAACCCTGATTGCACCCGATGCTACTGCTTCGAGTGTGTGGATAGCCTGGTCGGCCCTGGGACCTCGGGGAAGGTGCACGCCATGAGCAACTGGGTGTGCTACCTGTGCCTGCCGTCCTCCCGAAGCGGCCTGCTGCAGCGTCGGAGGAAGTGGCGCAGCCAGCTCAAGGCCTTCTACGACCGAGAGTCGGAGAATCCCCTTGAGATGTTCGAAACCGTGCCTGTGTGGAGGAGACAGCCAGTCCGGGTGCTGTCCCTTTTTGAAGACATCAAGAAAGAGCTGATGAGTTTGGGCTTTTTGGAAAGTGGTTCTGACCCGGGACAACTGAAGCATGTGGTTGATGTCACAGACACAGTGAGGAAGGATGTGGAGGAGTGGGGACCCTTCGATCTTGTGTACGGCGCCACACCTCCCCTGGGCCACACCTGTGACCGTCCTCCCAGCTGGTACCTGTTCCAGTTCCACCGGCTCCTGCAGTACGCACGGCCCAAGCCAGGCAGCCCCAGGCCCTTCTTCTGGATGTTCGTGGACAATCTGGTGCTGAACAAGGAAGACCTGGACGTCGCATCTCGCTTCCTGGAGATGGAGCCAGTCACCATCCCAGATGTCCACGGCGGATCCCTGCAGAATGCTGTCCGCGTGTGGAGCAACATCCCAGCCATAAGG---AGCAGGCACTGGGCTCTGGTTTTGGAAGAAGAATTGTCCCTGCTGGCCCAGAACAAGCAGAGCTCGAAGCTCGCGGCCAAGTGGCCCACCAAGCTGGTGAAGAACTGCTTTCTCCCCCTAAGAGAATATTTCAAGTATTTTTCAACAGAACTCACTTCCTCTTTA

>Bonobo3L

GCGGCCATCCCAGCCCTGGACCCAGAGGCCGAGCCCAGCATGGACGTGATTTTGGTGGGATCCAGTGAGCTCTCAAGCTCCGTTTCACCCAGGACAGGCAGAGATCTTATTGCATATGAAGTCAAGGCTAACCAGCGAAATATAGAAGACATCTGCATCTGCTGCGGAAGTCTCCAGGTTCACACACAGCACCCTCTGTTTGAGGGAGGGATCTGCGCCCCGTGTAAGGACAAGTCCCTGGATGCCCTCTTCCTGTACGACGATGACGGGTACCAATCCTACTGCTCCATCTGCTGCTCCGGAGAGACGCTGCTCATCTGCGGAAACCCTGATTGCACCCGATGCTACTGCTTCGAGTGTGTGGATAGCCTGGTCGGCCCCGGGACCTCGGGGAAGGTGCACGCCATGAGCAACTGGGTGTGCTACCTGTGCCTGCCGTCCTCCCGAAGTGGGCTGCTACAGCGTCGGAGGAAGTGGCGCAGCCAGCTCAAGGCCTTCTACGACCGAGAGTCGGAGAATCCCCTTGAGATGTTCGAAACCGTGCCTGTGTGGAGGAGACAGCCAGTCCGGGTGCTGTCCCTTTTTGAAGACATCAAGAAAGAGCTGACGAGTTTGGGCTTTTTGGAAAGTGGTTCTGACCCGGGACAACTGAAGCATGTGGTTGATGTCACAGACACAGTGAGGAAGGATGTGGAGGAGTGGGGACCCTTCGATCTTGTGTACGGCGCCACACCTCCCCTGGGCCACACCTGTGACCGTCCTCCCAGCTGGTACCTGTTCCAGTTCCACCGGCTCCTGCAGTACGCACGGCCCAAGCCAGGCAGCCCCAGGCCCTTCTTCTGGATGTTCGTGGACAATCTGGTGCTGAACAAGGAAGACCTGGACGTTGCATCTCGCTTCCTGGAGATGGAGCCAGTCACCATCCCAGATGTCCACGGCGGATCCCTGCAGAATGCTGTCCGCGTGTGGAGCAACATCCCAGCCATAAGG---AGCAGGCACTGGGCTCTGGTTTCGGAAGAAGAATTGTCCCTGCTGGCCCAGAACAAGCAGAGCTCGAAGCTCGCGGCCAAGTGGCCCACCAAGCTAGTGAAGAACTGCTTTCTCCCCCTAAAAGAATATTTCAAGTATTTTTCAACAGAACTCACTTCCTCTTTA

>Chimp3L

GCGGCCATCCCAGCCCTGGACCCAGAGGCCGAGCCCAGCATGGACGTGATTTTGGTGGGATCCAGTGAGCTCTCAAGCTCCATTTCACCCAGGACAGGCAGAGATCTTATTGCATATGAAGTCAAGGCTAACCAGCGAAATATAGAAGACATCTGCATCTGCTGCGGAAGTCTCCAGGTTCACACACAGCACCCTCTGTTTGAGGGAGGGATCTGCGCCCCCTGTAAGGACAAGTCCCTGGATGCCCTCTTCCTGTACGACGATGACGGGTACCAATCCTACTGCTCCATCTGCTGCTCCGGAGAGACGCTGCTCATCTGCGGAAACCCTGATTGCACCCGATGCTACTGCTTCGAGTGTGTGGATAGCCTGGTCGGCCCCGGGACCTCGGGGAAGGTGCACGCCATGAGCAACTGGGTGTGCTACCTGTGCCTGCCGTCCTCCCGAAGCGGGCTGCTACAGCGTCGGAGGAAGTGGCGCAGCCAGCTCAAGGCCTTCTACGACCGAGAGTCGGAGAATCCCCTTGAGATGTTCGAAACCGTGCCTGTGTGGAGGAGACAGCCAGTCCGGGTGCTGTCCCTTTTTGAAGACATCAAGAAAGAGCTGACGAGTTTGGGCTTTTTGGAAAGTGGTTCTGACCCGGGACAACTGAAGCATGTGGTTGATGTCACAGACACAGTGAGGAAGGATGTGGAGGAGTGGGGACCCTTCGATCTTGTGTACGGCGCCACACCTCCCCTGGGCCACACCTGTGACCGTCCTCCCAGCTGGTACCTGTTCCAGTTCCACCGGCTCCTGCAGTACGCACGGCCCAAGCCAGGCAGCCCCAGGCCCTTCTTCTGGATGTTCGTGGACAATCTGGTGCTGAACAAGGAAGACCTGGACGTTGCATCTCGCTTCCTGGAGATGGAGCCAGTCACCATCCCAGATGTCCACGGCGGATCCCTGCAGAATGCTGTCCGCGTGTGGAGCAACATCCCAGCCATAAGGAGCAGCAGGCACTGGGCTCTGGTTTCGGAAGAAGAATTGTCCCTGCTGGCCCAGAACAAGCAGAGCTCGAAGCTCGCGGCCAAGTGGCCCACCAAGCTGGTGAAGAACTGCTTTCTCCCCCTAAGAGAATATTTCAAGTATTTTTCAACAGAACTCACTTCCTCTTTA

>Orangoutan3L

GCAGCCATCCCAGCCCTGGACCCGGAGGCCGAGCCCAGCATGGATGTGATTTTGGTGGGATCCAGTGAGCTCTCAAGCTCCGTTTCACCCGGGACAGGCAGAGATCTTATTGCATATGAAGTCAAGGCTAACCAGCGAAATATAGAAGACATCTGCATCTGCTGCGGAAGTCTCCAGGTTCACACACAGCACCCTCTGTTTGAGGGAGGGATCTGTGCCCCGTGTAAGGACAAGTTCCTGGATGCCCTCTTCCTGTACGACGATGACGGGTACCAATCCTACTGCTCCATCTGCTGCTCTGGAGAGACACTGCTCATCTGCGGAAACCCTGATTGCACCCGATGCTACTGCTTCGAGTGTGTGGATAGCCTGGTCAGCCCCGGGACCTCAGGGAAGGTGCATGCCATGAGCAACTGGGTGTGCTACCTGTGCCTGCCCTCCTCCCGAAGCGGGCTGCTGCAGCGTCGGAGGAAGTGGCGCAGCCAGCTCAAGGCCTTCTACGACCGAGAGTCGGAGAATCCCCTTGAGATGTTCGAAACTGTGCCTGTGTGGAGGAGACAGCCGGTCCGGGTGCTGTCCCTTTTTGAAGACATCAAGAAAGAGCTGACAAGTTTGGGCTTTTTGGAAAGTGGTTCTGACGCGGGACAACTGAAGCACGTGGATGATGTCACAGACACGGTGAGGAAGGATGTGGAGGAGTGGGGACCCTTCGATCTTGTGTACGGCGCCACACCTCTCCTGGGCCACACCAGTGACCGTCCTCCCAGCTGGTACCTGTTCCAGTTCCACCGGCTCCTGCAGTACGCACGGCCCAAGCTAGGCAGCCCCAGACCCTTCTTCTGGATGTTTGTGGACAATCTGGTGCTCAACAAAGAAGACGAGGACGTCGCATGTCGCTTCCTGGAGATGGAGCCAGTCACCATCCCAGATGTCCACGGCGGATCCCTGCAGAATGCTGTCCGTGTGTGGAGCAACATCCCAGCCATAAGG---AGCAGGCACTTGGCTCTGGTTTCGGAAGAAGAATTGTCCCTGCTGGCCCAGAACAGGCAGAGCTCGAAGCTCGCGGCCAAGTGGCCCACCAAACTGGTGAAGAACTGCTTTCTCCCCCTAAGAGAATATTTCAAGTATTTTTCAACAGAACTCACTTCCTCTTTA

>Mangabey3L

GCGGCCATCCCAGCCCTGGACCCGGAGGCCGAACCCAGCATGGACGTGATCTTGGTGGGATCCAGTGAGCTCTCAAGCCCCGTTTCGCCCGGGGAAGGCAGAGATCTGATTGCATATGAAGTCAAGGTTAACCAGCGAAATATAGAAGACATCTGTCTCTGCTGCGGAAGTCTCCAGATTCACACGCAGCACCCTCTGTTTGAGGGAGGGATGTGCGCCCCGTGTAAGGACAAGTTCCTGGATGCCCTCTTCCTGTACGACGATGACGGGTACCAATCCTACTGCTCCATCTGCTGCTCCGGAGAGACGCTGCTCATCTGCGGAAACCCCGACTGCACCCGATGCTACTGCTTTGAGTGTGTGGATAGCCTGGTCGGCCCCGGGACCTCGGGGAAGGTGCATGCCATGAGCAACTGGGTGTGCTTCCTGTGCCTGCCCTTCTCCCGAAGCGGGCTGCTGCAACGCCGGAGGAAGTGGCGCAGTCAGCTCAAGGCCTTCTACGACCGAGAGTCGGAGAGTCCCCTTGAGATGTTTGAAACCGTGCCTGTGTGGAGGAGAGAGCCGGTCCGGGTGCTGTCTCTTTTTGAAGACATCAAGAAAGAGCTGACGAGTTTGGGCTTTTTGGAAAGCGGTTCTGACCCGGGACAGCTGAAGCATCTGGATGATGTCACAGACACAGTGAGGAAGGATGTGGAGGAGTGGGGGCCTTTCGATCTCCTGTACGGCGCCACACCTCCCCTGGGCCACACCTGCGACCATCCTCCCAGCTGGTACCTGTTCCAGTTCCACCGGCTCCTGCAGTACGCGCGGCCCGGGCCAGGCAGCCCCAGACCCTTCTTCTGGATGTTCGTGGACAATCTGGTACTGAACAAAGAAGACCAGGACGTCGCGTCTCGCTTCCTGGAGATGGAGCCCACCACCATCCCAGATGTCCGCGGCGGATCCCTACAGAATGCTGTCCGCGTGTGGAGCAACATTCCGGCCATAAGG---AGCAGGCAGTGGGCTCTGGTTTCGGAAGAAGAATTGTCCCTGCTGACTCAGAACAGGCAGAGCTCGAAGCTCGCGGCCAAGTGGCCCACCAAGCTGGTGAAGAACTGCTTTCTCCCCCTAAGAGAATATTTCAAGTATTTTTCAACCGAACTCACTTCCTCTTTA

>Gibbon3L

GCGGACATCCCAGCCCTGGACCCGGAGGCTGAGCCCAGCATGGATGTGATTTTGGTGGGATCCAGTGAGCTCTCAAGCTCTGTTTCACCCGGGACAGGCAGAGATCTTATTGCATATGAAGTCAAGGCTAACCAGCGAAATATAGAAGACATCTGCATCTGCTGCGGAAGTCTCCAGGTTCACACGCAGCACCCTCTGTTTGAGGGAGGGATCTGCACCCCGTGTAAGGACAAGTTCCTGGATGCCCTCTTCCTGTACGACGATGACGGGTACCAATCTTACTGCTCCATCTGCTGCTCCGGAGAGACGCTGCTCATCTGCGGAAACCCTGATTGCACCCGATGCTACTGCTTCGAATGTGTGGATAGCCTGGTCGGCCCCGGGACGTCGGGGAAGGTGCATGCTATGAGCAACTGGGTGTGCTTCCTGTGCCTGCCCTTCTCCCGAAACGGGCTGCTGCAGCGTCGGAGGAAGTGGCGCAGCCAGCTCAAGGCCTTCTACGACCGAGAGTCGGAGAATCCCCTTGAGATGTTCGAAACCGTGCCTGTGTGGAGGAGACAGCCGGTCCGGGTGCTGTCCCTTTTTGAAAACATCAAGAAAGAGCTGACGAGTTTGGGCTTTCTGGAAAGTGGTTCTAAGCCGGGACAGCTGAAGCATGTGGATGATGTCACAGACACAGTGAGGAAGGACGTGGAGGAGTGGGGACCCTTTGATCTTCTGTATGGCGCCACACCTCCCCTGGGCCACACCTGTGACCGTCCTCCCAGCTGGTACCTGTTCCAGTTCCACCGGCTCCTGCAGTACGCACGGCCCAAGCCAGGCAGCCCCAGGCCCTTCTTCTGGATGTTCGTGGACAATCTGGTGCTGAACAAAGAAGACCAGGATGTCGCGTCTCGCTTCCTGGAGATGGAGCCAGTCACCATCCCAGATGTCCACGGCGGATCCCTACAGAATGCTGTCCACGTGTGGAGCAACATCCCAGCCATAAGGAGCGGCAGGCACTCGGCTCTGGTTTCGGAAGTGGAATTGTCCCTGCTGGCTCAGAACAGGCAGAGTTCGAAGCTCATGGCCAAGTGGCCCACCAAGCTGGTGAAGAACTGCTTTCTCCCCCTAAGAGAATATTTCAAGTATTTTTCAACAGAACTCACTTCCTCTTTA

>Mandrill3L

GCGGCCATCCCAGCCCTGGACCCGGAGGCCGAGCCCAGCATGGACGTGATCTTGGTGGGATCCAGTGAGCTCTCAAGCCCCGTTTCGCCCGGGGAAGGCAGAGATCTGATTGCATATGAAGTCAAGGTTAACCAGCGAAATATAGAAGACATCTGCCTCTGCTGCGGAAGTCTCCAGGTTCACACGCAGCACCCTCTGTTTGAGGGAGGGATGTGCGCCCCGTGTAAGGACAAGTTCCTGGATGCCCTCTTCCTGTACGACGATGATGGGTACCAATCCTACTGCTCCATCTGCTGCTCTGGAGAGACGCTGCTCATCTGCGGAAACCCCGACTGCACCCGATGCTACTGCTTTGAGTGTGTGGATAGCCTGGTCGGCCCCGGGACCTCGGGGAAGGTGCATGCCATGAGCAACTGGGTGTGCTTCCTGTGCCTGCCCTTCTCCCGAAGCGGGCTGCTGCAACGCCGGAGGAAGTGGCGCGGTCAGCTCAAGGCCTTCTACGACCGAGAGTCGGAGAGTCCCCTTGAGATGTTTGAAACCGTGCCTGTGTGGAGGAGAGAGCCGGTCCGGGTGCTGTCTCTTTTTGAAGACATCAAGAAAGAGCTGACGAGTTTGGGCTTTTTGGAAAGCGGTTCTGACCCAGGACAGCTGAAGCATCTGGATGATGTCACAGACACAGTGAGGAAGGATGTGGAGGAGTGGGGGCCCTTCGATCTCCTGTACGGCGCCACACCTCCCCTGGGCCACACCTGCGACCATCCTCCCAGCTGGTACCTGTTCCAGTTCCACCGGCTCCTGCAGTACGCGCGGCCCGGGCCAGGCAGCCCCAGACCCTTCTTCTGGATGTTCGTGGACAATCTGGTACTGAACAAAGAAGACCAGGACGTCGCGTCTCGCTTCCTGGAGATGGAGCCCACCACCATCCCAGATGTCCGCGGCGGATCCCTACAGAATGCTGTCCGCGTGTGGAGCAACATTCCGGCCATAAGG---AGCAGGCAGTGGGCTCTGGTTTCGGAAGAAGAATTGTCCCTGCTGACTCAGAACAGGCAGAGCTCGAAGCTCGCGGCCAAGTGGCCCACCAAGCTGGTGAAGAACTGCTTTCTCCCCCTAAGAGAATATTTCAAGTATTTTTCAACCGAACTCACTTCCTCTTTA

>GoldenSnubNosed3L

GCAGCCATCCCAGCCCTGGACCCGGAGGCCGAGCCCAGTATGGACGTGATCTTGGTGGGATCCAGTGAGCTCTCAAGCCCCGTTTCGCCCGGGGCAGGCAGAGATCTTATTGCATATGAAGTCAAGGTTAACCAGCGAAATATAGAAGACATCTGTCTCTGCTGCGGGAGTCTCCAGGTTCACACGCAGCACCCTCTGTTTGAGGGAGGGATGTGCGCCCCATGTAAGGACAAGTTCCTGGATGCCCTCTTCCTGTACGACGATGACGGGTACCAATCCTACTGCTCTATCTGCTGCTCCGGAGAGACGCTGCTTATCTGCGGAAACCCCGATTGCACCCGATGCTACTGCTTTGAGTGTGTGGACAGCCTGGTCGGCCCCGGGACCTCAGGGAAGGTGCATGCCATGAGCAACTGGGTGTGCTTCCTGTGCCTGCCCTTCTCCCGAAGCGGGCTGCTGCAACGCCGGAGGAAGTGGCGCGGTCAGCTCAAGGCCTTCTATGACCGAGAGTCGGAGAGTCCCCTTGAGATGTTTGAAACAGTGCCTGTGTGGAGGAGAGAGCCAGTCCGGGTGCTGTCTCTTTTTGAAGACATCAAGAAGGAGCTGACGAGTTTGGGCTTTTTGGAAAGCGGTTCTGACCCGGGACAGCTGAAGCATTTGGATGATGTCACAGACACAGTGAGGAAGGATGTGGAGGAGTGGGGGCCCTTCGATCTCCTGTACGGCGCCACACCTCCCTTGGGCCACACCTGCGACCATCCTCCCAGCTGGTACCTGTTCCAGTTCCACCGGCTCCTGCAGTACGCGCGGCCCGGGCCGGGCAGCCCCAGGCCCTTCTTCTGGATGTTCGTGGACAATCTGGTACTGAACAAAGAAGACCAGGACGTTGCCTCTCGCTTCCTGGAGATGGAGCCAACCACCATCCCAGATGTCCGCGGCGGATCCCTACAGAATGCTGTCCGCGTGTGGAGCAACATTCCAGCCATAAGG---AGCAGGCAGTGGGCTCTGGTTTCGGAAGAAGAATTGTCCCTGCTGACACAGAACAGGCAGAGCTCGAAGCTCGCGGCCAAGCAGCCCACCAAGCTGGTGAAGAACTGCTTTCTCCCCCTAAGAGAATATTTCAAGTATTTTTCAACCGAACTCACTTCCTCTTTA

>Baboon3L

GCGGCCATCCCAGCCCTGGACCCGGAGGCCGAGCCCAGCATGGACGTGATCTTGGTGGGATCCAGTGAGCTCTCAAGCCCCGTTTCGCACGGGGAAGGCAGAGATCTGATTGCATATGAAGTCAAGGTCAACCAGCGAAATATAGAAGACATCTGTCTCTGCTGCGGAAGTCTCCAGGTTCACACGAAGCACCCTCTGTTTGAGGGAGGGATATGCGCCCCGTGTAAGGACAAGTTCCTGGATGCCCTCTTCCTGTACGACGATGACGGGTACCAATCCTACTGCTCCATCTGCTGCTCCGGAGAGACGCTGCTCATCTGCGGAAACCCCGATTGCACCCGATGCTACTGCTTTGAGTGTGTGGATAGCCTGGTCGGCCCCGGGACCTCGGGGAAGGTGCATGCCATGAGCAACTGGGTGTGCTTCCTGTGCCTGCCCTTCTCCCGAAGCGGGCTGCTGCAACGCCGGAGGAAGTGGCGCAGTCAGCTCAAGGCCTTCTACGACCGAGAGTCGGAGAGTCCCCTTGAGATGTTTGAAACTGTGCCTGTGTGGAGGAGAGAGCCGGTCCGGGTGCTGACTCTTTTTGAAGACATCAAGAAAGAGCTGACGAGTTTGGGCTTTTTGGAAAGCGGTTCTGACCCGGGACAGCTGAAGCATCTGGATGATGTCACAGACACAGTGAGGAAGGATGTGGAGGAGTGGGGGCCCTTCGATCTCCTGTACGGCGCCACACCTCCCCTGGGCCACACCTGCGACCATCCTCCCAGCTGGTACCTGTTCCAGTTCCACCGGCTCCTGCAGTACGCGCGGCCCGGGGCAGGCAGCCCCAGACCCTTCTTCTGGATGTTCGTGGACAATCTGGTACTGAACAAAGAAGACCAGGACGTCGCGTCTCGCTTCCTGGAGATGGAGCCCACCACCATCCCAGATGTCCGCGGCGGATCCCTACAGAATGCTGTCCGCGTGTGGAGCAACATTCCGGCCATAAGG---AGCAGGCAGTGGGCTCTGGTTTCGGAAGAAGAATTGTCCCTGCTGACTCAGAACAGGCAGAGCTCGAAGCTCGCGGCCAAGTGGCCCACCAAGCTGGTGAAGAACTGCTTTCTCCCCCTAAGAGAATATTTCAAGTATTTTTCAACCGAACTCACTTCCTCTTTA

>PigTailedMacaque

GCGGCCATCCCAGCCCTGGACCCAGAGGCCGAGCCCAGCATGGACGTGATCTTGGTGGGATCCAGTGAGCTCTCAAGCCCCGTTTCGCCCGGGGAAGGCAGAGATCTGATTGCATATGAAGTCAAGGTTAACCAGCGAAATATAGAAGACATCTGTCTCTGCTGCGGAAGTCTCCAGGTTCACACGCAGCACCCTCTGTTTGAGGGAGGGATATGCGCCCCGTGTAAGGACAAGTTCCTGGATGCCCTCTTCCTGTACGACGATGATGGGTACCAATCCTACTGCTCCATCTGCTGCTCCGGAGAGACGCTGCTCATCTGCGGAAACCCCGATTGCACCCGATGCTACTGCTTTGAGTGTGTGGATAGCCTGGTCGGCCCCGGGACCTCGGGGAAGGTGCATGCCATGAGCAACTGGGTGTGCTTCCTGTGCCTGCCCTTCTCCCGAAGCGGGCTGCTGCAACGCCGGAGGAAGTGGCGCGGTCAGCTCAAGGCCTTCTACGACCGAGAGTCGGAGAGTCCCCTTGAGATGTTTGAAACCGTGCCTGTGTGGAGGAGAGAGCCGGTCCGCGTGCTGTCTCTTTTTGAAGACATCAAGAAAGAGCTGACGAGTTTGGGCTTTTTGGAAAGCGGTTCTGACCCGGGACAGCTGAAGCATCTGGATGATGTCACAGACACAGTGAGGAAGGATGTGGAGGAGTGGGGGCCCTTTGATCTCCTGTACGGCGCCACACCTCCCCTGGGCCACACCTGCGACCATCCTCCCAGCTGGTACCTGTTCCAGTTCCACCGGCTCCTGCAGTACGTGCGGCCCGGGCCAGGCAGCCCCAGACCCTTCTTCTGGATGTTCGTGGACAATCTGGTACTGAACAAAGAAGACCAGGACGTCGCGTCTCGCTTCCTGGAGATGGAGCCCACCACCATCCCAGATGTCCGTGGCGGATCCCTACAGAATGCTGTCCGCGTGTGGAGCAACATTCCGGCCATAAGG---AGCAGGCAGCGGGTTCTGGTTTCGGAAGAAGAATTGTCCCTGCTGACTCAGAACAGGCAGAGCTCGAAGCTCGCGGCCAAGTGGCCCACCAAGCTGGTGAAGAACTGCTTTCTCCCCCTAAGAGAATATTTCAAGTATTTTTCAACCGAACTCACTTCCTCTTTA

>RhesusMacaque3L

GCGGCCATCCCAGCCCTGGACCCAGAGGCGGAGCCCAGCATGGACGTGATCTTGGTGGGATCCAGTGAGCTCTCAAGCCCCGTTTCGCCCGGGGAAGGCAGAGATCTGATTGCATATGAAGTCAAGGTTAACCAGCGAAATATAGAAGACATCTGTCTCTGCTGCGGAAGTCTCCAGGTTCACACGCAGCACCCTCTGTTTGAGGGAGGGATATGCGCCCCGTGTAAGGACAAGTTCCTGGATGCCCTCTTCCTGTACGACGATGATGGGTACCAATCCTACTGCTCCATCTGCTGCTCCGGAGAGACGCTGCTCATCTGCGGAAACCCCGATTGCACCCGATGCTACTGCTTTGAGTGTGTGGATAGCCTGGTCGGCCCCGGGACCTCGGGGAAGGTGCATGCCATGAGCAACTGGGTGTGCTTCCTGTGCCTGCCCTTCTCCCGAAGCGGGCTGCTGCAACGCCGGAGGAAGTGGCGCGGTCAGCTCAAGGCCTTCTACGACCGAGAGTCGGAGAGTCCCCTTGAGATGTTTGAAACCGTGCCTGTGTGGAGGAGAGAGCCGGTCCGCGTGCTGTCTCTTTTTGAAGACATCAAGAAAGAGCTGACGAGTTTGGGCTTTTTGGAAAGCGGTTCTGACCCGGGACAGCTGAAGCATCTGGATGATGTCACAGACACAGTGAGGAAGGATGTGGAGGAGTGGGGGCCCTTTGATCTCCTGTACGGCGCCACACCTCCCCTGGGCCACACCTGCGACCATCCTCCCAGCTGGTACCTGTTCCAGTTCCACCGGCTCCTGCAGTACGTGCGGCCCGGGTCAGGCAGCCCCAGACCCTTCTTCTGGATGTTCGTGGACAATCTGGTACTGAACAAAGAAGACCAGGACGTCGCGTCTCGCTTCCTGGAGATGGAGCCCACCACCATCCCAGATGTCCGTGGCGGATCCCTACAGAATGCTGTCCGCGTGTGGAGCAACATTCCGGCCATAAGG---AGCAGGCAGCGGGTTCTGGTTTCGGAAGAAGAATTGTCCCTGCTGACTCAGAACAGGCAGAGCTCGAAGCTCGCGGCCAAGTGGCCCACCAAGCTGGTGAAGAACTGCTTTCTCCCCCTAAGAGAATATTTCAAGTATTTTTCAACCGAACTCACTTCCTCTTTA

>CrabEatingMacaque3L

GCGGCCATCCCAGCCCTGGACCCAGAGGCCGAGCCCAGCATGGACGTGATCTTGGTGGGATCCAGTGAGCTCTCAAGCCCCGTTTCGCCCGGGGAAGGCAGAGATCTGATTGCATATGAAGTCAAGGTTAACCAGCGAAATATAGAAGACATCTGTCTCTGCTGCGGAAGTCTCCAGGTTCACACGCAGCACCCTCTGTTTGAGGGAGGGATATGCGCCCCGTGTAAGGACAAGTTCCTGGATGCCCTCTTCCTGTACGACGATGATGGGTACCAATCCTACTGCTCCATCTGCTGCTCCGGAGAGACGCTGCTCATCTGCGGAAACCCCGATTGCACCCGATGCTACTGCTTTGAGTGTGTGGATAGCCTGGTCGGCCCCGGGACCTCGGGGAAGGTGCATGCCATGAGCAACTGGGTGTGCTTCCTGTGCCTGCCCTTCTCCCGAAGCGGGCTGCTGCAACGCCGGAGGAAGTGGCGCGGTCAGCTCAAGGCCTTCTACGACCGAGAGTCGGAGAGTCCCCTTGAGATGTTTGAAACCGTGCCTGTGTGGAGGAGAGAGCCGGTCCGCGTGCTGTCTCTTTTTGGAGACATCAAGAAAGAGCTGACGAGTTTGGGCTTTTTGGAAAGCGGTTCTGACCCGGGACAGCTGAAGCATCTGGATGATGTCACAGACACAGTGAGGAAGGATGTGGAGGAGTGGGGGCCCTTTGATCTCCTGTACGGCGCCACACCTCCCCTGGGCCACACCTGCGACCATCCTCCCAGCTGGTACCTGTTCCAGTTCCACCGGCTCCTGCAGTACGTGCGGCCCGGGCCAGGCAGCCCCAGACCCTTCTTCTGGATGTTCGTGGACAATCTGGTACTGAACAAAGAAGACCAGGACGTCGCGTCTCGCTTCCTGGAGATGGAGCCCACCACCATCCCAGATGTCCGTGGCGGATCCCTACAGAATGCTGTCCGCGTGTGGAGCAACATTCCGGCCATAAGG---AGCAGGCAGCGCGTTCTGGTTTCGGAAGAAGAATTGTCCCTGCTGACTCAGAACAGGCAGAGCTCGAAGCTCGCGGCCAAGTGGCCCACCAAGCTGGTGAAGAACTGCTTTCTCCCCCTAAGAGAATATTTCAAGTATTTTTCAACCGAACTCACTTCCTCTTTA

>Colobus3L

GCGGCCATCCCAGCCCTGGACCCGGAGGCCGAGCCCAGTATGGACGTGATCTTGGTGGGATCCAGTGAGCTCTCAAGCCCCGTTTCACCTGGGGCAGGCAGAGATCTTATTGCATATGAAGTCAAGGTTAACCAGCGAAATATAGAAGACATCTGTCTCTGCTGCGGAAGTCTCCAGGTTCACACGCAGCACCCTCTGTTTGAGGGAGGGATGTGCACCCCATGTAAGGACAAGTTCCTGGATGCCCTCTTCCTGTATGACGATGACGGGTACCAGTCCTACTGCTCCATCTGCTGCTCCGGAGAGACGCTGCTCATCTGCGGAAACCCGGACTGCACCCGATGCTACTGCTTTGAGTGTGTGGATAGCCTGGTCGGCCCCGGGACCTCGGGGAAGGTGCATGCCATGAGCAACTGGGTGTGCTTCCTGTGCCTGCCCTTCTCCCGAAGCGGGCTGCTGCAACGCCGGAGGAAGTGGCGCGGTCAGCTCAAGGCCTTCTACGACCGAGAGTCGGAGAGTCCCCTTGAGATGTTTGAAACCGTGCCTGTGTGGAGGAAAGAGCCGGTCCGGGTGCTGTCTCTTTTTGAAGACATCAAGAAAGAGCTGACGAGTTTGGGCTTTTTGGAAAGCGGTTCTGACCTGGGACAGCTGAAGCATTTGGATGATGTCACAGACACAGTGAGGAAGGATGTGGAGGAGTGGGGGCCCTTCGATCTCCTGTACGGCGCCACACCTCCCCTGGGCCACACCTGCGACCATCCTCCCAGCTGGTACCTGTTCCAGTTCCACCGGCTCCTGCAGTACGCGCGGCCCGGGCCGGGCAGCCCCAGGCCCTTCTTCTGGATGTTCGTGGACAATCTGGTACTGAACAAAGAAGACCAGGACATCGCCTCTCGCTTCCTGGAGATGGAGCCAACCACCATCCCAGATGTCCGCGACGGATCCCTACAGAATGCCGTCCGCGTGTGGAGCAACATTCCAGCCATAAGG---AGCAGGCAGTGGGCTCTGGTTTCGGAAGAAGAATTGTCCCTGCTGATACAGAACAGGCAGAGCTCGAAGCTCGCGGCCAAGTGGCCCACCAAGCTGGTGAAGAACTGCTTTCTCCCTCTAAGAGAATATTTCAAGTATTTTTCAACCGAACTCACTTCCTCTTTA

>AGM3L

GCGGCCATCCCAGCCTTGGACCCAGAGGCCGAGCCCAGCATGGATGTGATCTTGGTGGGATCCAGTGAGCTCTCAAGCCCCGTTTCGCCCAGGGAAGGCAGAGATCTGATTGCATATGAAGTCAAGGTTAACCATCGAAATATAGAAGACATCTGTCTCTGCTGCGGAAGTCTCCAGGTTCACACGCAGCACCCTCTGTTTGAGGGAGGGATGTGCGCCCCGTGTAAGGACAAGTTCCTGGATGCCCTCTTCCTATATGACGATGACGGGTACCAATCGTACTGCTCCATCTGCTGCTCCGGAGAGACGCTGCTCATCTGCGGAAACCCCGACTGCACCCGGTGCTACTGCTTCGAGTGTGTGGATAGCCTGGTCGGCCCCGGGACCTCAGGGAAGGTGCATGCCATGAGCAACTGGGTGTGCTTCCTGTGCCTGCCCTTCTCCCGAAGCGGGCTGCTGCAACGCCGGAGGAAGTGGCGCGGTCAGCTCAAGGCCTTCTACGACCGAGAGTCGGAGAGTCCCCTTGAGATGTTTGAAACCGTGCCTGTGTGGAGGAGAGAGCCGGTCCGGGTGCTGTCTCTTTTTGAAGACATCAAGAAGGAGCTGACGAGTTTGGGCTTTTTGGAAAGCGGTTCTGAGCCGGGACAGCTGAAGCATCTGGACGATGTCACAGACACAGTGAGGAAGGATGTGGAGGAGTGGGGGCCCTTCGATCTCCTGTACGGCGCCACACCTCCCCTGGGCCACACCTGCGACCATCCTCCCAGCTGGTACCTGTTCCAGTTCCACCGGCTCCTGCAGTACGCGCGGCCCGGACCAGGCAGCCCCAGACCCTTCTTCTGGATGTTCGTGGACAATCTGGTACTGAACAAAGAAGACCAGGACGTCGCGTCTCGCTTCCTGGAGATGGAGCCCACCACCATCCCAGATGTCCGCGGCGGATCCCTACAGAATGCTGTCCGCGTGTGGAGCAACATTCCGGCCATAAGGAGCAGCAGGCAGTGGGCTCTGGTTTCGGAAGAAGAATTGTCCCTGCTGACTCAGAACAAGCAGAGATCGAAGCTCGCGGCCAAGTGGCCCACCAAGCTGGTGAAGAACTGCTTTCTCCCCCTAAAAGAATATTTCAAGTATTTTTCAACCGAACTCACTTCCTCTTTA

>RedColobus3L

GCGGCCATCCCAGCCCTGGACCCGGAGGCTGAGCCCAGTATGGACGTGATCTTGGTGGGATCCAGTGAGCTCTCAAGCCCCGTTTCATCTGGGGCAGGCAGAGATCTTATTGCGTATGAAGTCAAGGTTAACCAGCGAAATATAGAAGACATCTGTCTCTGCTGCGGAAGTCTCCAGGTTCACACGCAGCACCCTCTGTTTGAAGGAGGGATGTGCACCCCATGTAAGGACAAGTTCCTGGATGCCCTCTTCCTGTACGATGATGACGGGTACCAATCCTACTGCTCCATCTGCTGCTCCGGAGAGACGCTGCTCATCTGCGGAAACCCTGATTGCACCCGATGCTACTGCTTTGAGTGTGTGGATAGCCTGGTCGGCCCCGGGACCTCCGGGAAGGTGCATGCCATGAGCAACTGGGTGTGCTTCCTGTGCCTGCCCTTTTCCCGAAGCGGGCTGCTGCAACGCCGGAGGAAGTGGCGCGGTCAGCTCAAGGCCTTCTATGACCGAGAGTCGGAGAGTCCCCTTGAGATGTTTGAAACCGTGCCTGTGTGGAGGAGAGAGCCCGTCCGGGTGCTGTCTCTTTTTGAAGACATCACGAAAGAGCTGACGAGTTTGGGCTTTTTGGAAAGCGGTTCCGACCCGGGACAGCTGAAGCATCTGGATGACGTCACAGACACAGTGAGGAAGGATGTGGAGGAGTGGGGGCCCTTCGGTCTCCTGTACGGCGCCACACCTCCCCTGGGCCACACCTGCGAGCATCCTCCCAGTTGGTACCTGTTCCAGTTCCACCGGCTCCTGCAGTACGCGCGGCCCGGGCCGGGCAGCCCCAGGCCCTTCTTCTGGATGTTCGTGGACAATCTGGTACTGAACAAAGAAGACCAGGACGTCGCCTCTCGCTTCCTGGAGATGGAGCCAACCACCATCCCAGATGTCCGCGGCGGATCCCTACAGAATGCTGTCCGCGTGTGGAGCAACATTCCAGCCATAAGG---AGCAGGCAGTGGGCTCTGGTTTCGGAAGAAGAATTGTCCCTGCTGACACAGAACAGGCAGAGCTCGAAGCTCGCGGCCAAGCGGCCCACCAAGCTGGTGAAGAACTGCTTTCTCCCCCTAAGAGAATATTTCAAGTATTTTTCAACCGAACTCACTTCCTCTTTA

>PhilipineTarsier3L

GCAGCAGCCCCAGTCCTGGACCTGGAGGCTGAATGTAGCTTGGACGTGATCCTGGTAGGCTCCAGCGAGCTGTCGACTTCCTCTTCACCCAGGCTGGGCAGAGATCACATTGCATATGAAGTCAAGGTTAACCAGCGAAACATAGAAGACATCTGCCTCTGCTGTGGAAGTTTCCTCGTTCACACACAGCACCCTCTGTTTGAGGGAGGGATGTGTGCCCCATGTAAGGACAAGTTCCTGGACACACTTTTCCTGTATGACGAGGACGGGTACCAATCCTACTGTTCCATCTGCTGCTCGGGAGAGACACTGCTCATCTGTGAAAACCCCGATTGCACCCGATGCTACTGTTTTGAGTGTTTGGACACCCTGGTCAGCCCAGGGACCTCGGAGAAAGTCCATGCCATGAGTAACTGGGTGTGCTTCCTGTGCCTGCCTTTCACCCGCAGTGGCCTCCTGCAGAGGAGGAGGAAGTGGCGAGGCCAACTGAAGGCCTTCTATGACCGTGAGTCGGAGAGTTCTCTTGAAATGTACAAAACCGTGCCTGTGTGGAAGAGAGAACCAGTGCGGGTGCTGTCCCTTTTTGGGGACATCAAGAAAGAGCTGATGAGTTTGGGCTTTGTGGAAACCGGTTCTGACCCAGGAAGACTGAGGCATTTGGATGATACCACCAACATAGTGAGGAGGAACGTGGAAGAGTGGGGTCCATTCCATCTCCTGTATGGTGCAACACCTCCCTTGGGCCACACCTGCGACCGTCCTCCCGGCTGGTACCTGTTCCAGTTCCACCGGCTCCTGCAGTACGCACGGCCCCAGCCTGGCAGCCCGCAGCCCTTCTTCTGGATGTTTGTGGACAACGTCATGCTGACCAGAGAAGACCGGGCCATTGCAAGTCGTTTCCTGGAGACAGAGCCTGTGACCATCCCGGACATCCATGGCAGAGCCCTCCAGAATGCTGTGTGTGTATGGAGTAATATCCCTGCGGTAAGG---AGCAAGCACTCAGCCCTGGTTTCAGAAGAGGAGTTGTCCCTGCTGGCTCAGGACAGACAGAGAGCAAAGCTTCCCACCCAGGGGCCCACCAAACTGGTGAAGAACTGCTTTCTCCCCCTAAGAGAATATTTCAAATATTTTTCAACAGAACTCACTTCCTTCTTA

>OwlMonkey3L

GCAGCCACCCCAGCCCTGGACCTGGAGGCCGAGCCCAGCATGGACGTGATCTTGGTGGGATCCAGTGGGCTCTCAAGCCCTGTTCCCCCGGCCACAGGCAGAGAGCTTATTGCATATGAAGTCAAGGTTAACCAGCGGAACATAGAAGAGATCTGCCTCTGCTGCGGAAGCTTCCAGGTTCACACGCGGCACCCTCTGTTTGAGGGAGGGATGTGCGCTCCATGTAAGGACAAGTTCCTGGACGCCCTCTTCCTGTACGACGAGGACGGGTACCAGTCCTACTGCTCCATCTGCTCCTCTGGGGGGACGCTGCTCATCTGCGAAAACCCCGATTGCACCCGATGCTACTGCTTCGAGTGTGTGGATATCCTGGTCGGCCCCGGGACCTCGGGGAAGGTGCACGCCATGAGCAACTGGGTGTGCTTCCTGTGCCTGCCGTTCTCCCGCAGCGGGCTGCTCCAGCGCCGGAAGAAGTGGCGTAGCCATCTCAAGGCCTTCTACGACCAGGAGTCGAAGAGTCCACTTGAGATGTTCGAAACCCTGCCTGTGTGGAGGAGACAGCCGGTGCGGGTGCTGGCCCTTTTTGGGGACATCAAGAAAGAGCTGACGAGTCTGGGCTTTTTGGGAAGCGGTTCTGACCAGGGGCAGCTGAAGCACTTGGATGACGTCACAGACGTAGTGAGGAAGGACGTGGAGGAGTGGGGCCCGTTCGACCTCCTGTATGGCGCCACACCTGCCCCCGGCCACGCCTGTGACCATCCTCCTGCCTGGTACCTGTTCCAGTTCCACCGGCTCTTGCAGTACGCTCGGCCC---TCGGGCAGCACCCGGCCCTTCTTCTGGATGTTCGTGGACCATCTGAGGCTAAACAAAGAAGACCAAGACCTCGTGTATCGCTTCCTGGAGATGGAGCAGGCCACCATCCCGGATGTCTGCGGCGGAGTCCTGCAGGGCGCTGTCCGAGTGTGGAGCAACATCCCAGCCATCAGG---AGCAGGCACTCGGCCTTGCCTTCAGAAGAAGAATTGTCCGTGCTGGCTCAGAACCGACAGAGCTCCAAGCTCCCCACCCAGCGGCCCATCAAGCTGCTGAAGAGCTGCTTTCTCCCCCTAAGAGAGTATTTCAAGTATTTTTCAACAGAACTCACTTCCTCTTCA

>Sapajou3L

GCAGCCACCCCAGCCCTGGACCTGGAGGCCGAGCCCAGCATGGATGTGATCTTGGTGGGATCCAGTGAGCTCTCAAGCCCCGTTCCCGCGGCCACAGGCAGAGAGCTTATTGCATATGAAGTCAAGGTGAACCAGCGGAACATAGAAGAGATCTGCCTCTGCTGCGGAAGCTTCCAGGTTCACACGCAGCACCCTCTGTTTGAGGGAGGGATGTGCGCCCCGTGTAAGGACAAGTTCCTGGACGCCCTCTTCCTGTACGACGATGACGGATACCAATCCTACTGCTCCATCTGCTCCTCCGGGGGCACGCTGCTCATCTGCGAAAACCCCGATTGCACCCGATGCTACTGCTTCGAGTGTGTGGATATCCTGGTCGGCCCCGGGACCTCGGGGAAGGTGCACGCCATGAGCAACTGGGTGTGCTTCCTGTGCCTGCCCTTCTCCCGCAGCGGGCTGCTCCAGCGCCGGAAGAAGTGGCGCAGCCACCTCAAGGCCTTCTACGACCAGGAGTCGAAGAGTCCCGTTGAGATGTTCGAAACCCTGCCCGTGTGGAGGAGACAGCCGGTGCGGGTGCTGGCCCTTTTTGGGGACATCAAGAAAGAGCTGACGAGTCTGGGCTTTTTGGGAAGCGGTTCTGACCAGGGGCGACTGAAGCACTTGGATGATGTCACAGACGTAGTGAGGAAGGACGTGGAAGAGTGGGGCCCGTTCGACCTCCTGTATGGTGCCACACCTGCCCCCGGCCACGCCTGTGACCATCCTCCCGCCTGGTACCTGTTCCAGTTCCACCGGCTCCTGCAGTACGCACGGCCC---TCGGGCAGCCTCCGGCCCTTCTTCTGGATGTTCGTGGATCATCTGATGCTAAACAAAGAGGACCAAGACCTCGTGTGTCGCTTCCTGGAGATGGAGCAGGCCACCATCCCGGATGTCCGCGGCGGAGTCCTGCAGAGTGCTGTCCGAGTGTGGAGCAACATCCCAGCCATCAGGAGCAGCAGGCACTCAGCCCTGCCTTCGGAAGAAGAATTGTCCGGGCTGGCTCAGAACCGACAGAGCTCCAAGCTTCCCGCCCAGCGGCCCACCAAGCTGCTGAAGAACTGCTTTCTCCCCCTAAGAGAATATTTCAAGTATTTTTCAACAGAACTCACTTCCTCTTCA

>Marmoset3L

GCAGCCACCCCAGCCCTGGACCTGGAGGCAGAGCCCAGCGTGGATGTGATCTTGGTGGGATCCAGTGAGCTGTCAAGCCCCGTCCCCACGGCCACAGGCAGAGAGCTTACTGCATACGAAGTCAAGGCTAACCGGCGGGACATAGAAGAGATCTGCCTCTGCTGCGGAAGCTTCCAGGTTCACACGCAGCACCCTCTGTTTGAGGGAGGGATGTGCGCCCCATGTAAGGACAAGTTCCTGGACGCCCTCTTCCTGTACGACGATGATGGGTACCAATCCTACTGCTCCATCTGCTCCTCCGGGGGGACGCTGCTCATCTGCGAAAACCCCGATTGCACCCGATGCTATTGCTTCGAGTGTGTGGATATCCTGGTCGGCCCCGGGACCTCAGGGAAGGTGCACGCCATGAGCAACTGGGTGTGCTTCCTGTGCCTGCCCTTCTCCCGCAGCGGGCTGCTCCAGCGCCGGAAGAAGTGGCACAGCCACCTCAAGGCCTTCTATGACCAGGAGTCGAAGAGTCCGCTTGAGATGTTCGAAACCCTGCCTGTGTGGAGGAGACAGCCCATGCGGGTGCTGGCCCTTTTTGGGGACATCAAGAAAGAGCTGACGAGTCTGGGCTTTCTGGGAAGCGGTTCTGACCAGGGGCGACTGAAGCACTTGGATGATGTCAGAGACGTAGTGAGGAAGGACGTGGAAGAGTGGGGCCCGTTCGACCTCCTGTATGGCGCCACACCTGCCCCCGGCCACGCCTGTGACCATCCTCCCGCCTGGTACCTGCTCCAGTTCCACCGGCTCCTGCAGTACGCACGGCCC---TCGGGCAGCCCCCGGCCCTTCTTCTGGATGTTCGTGGACCATCTGATGCTAAACAAAGACGACCAAGACCTCGTGTGTCGCTTCCTGGAGATGGAGCAGGCCACCATCCCGGATGTCCGTGGCGGAGTCCTGCAGAGCGCCGTCCGAGTGTGGAGCAACATCCCAGCCATCAGGAGCAGCAGGCACTCGGCCCTGCCTTCGGAAGAAGAATTGTCCCTGCTGGCTCAGAACCGACAGAGCTCCAAGCTCCCCGCCCAGCGGCCCACCAAGCTGCTGAAGAACTGCTTTCTCCCCCTAAAAGAATATTTCAAGTATATTTCAACAGAACTCACTTCCTCTTCA

>Sifaka3L

GCAGCCACCCCAGTCCTAGACCTAGAGGCTGAGCCCAGTGTGGACGTGATCCTGGTGGGCTCCGG------------CCCGCCGTCACCCGGGCGGGGCAGAGATCTCATTGCGTATGAAGTCCAGGTGAACCAGAGAAACATAGAAGACATCTGCCTCTGCTGCGGAAGCTTCCGGGTTCACACGCAGCACCCTCTGTTTGAGGGGGGAATGTGCGCTCCGTGTAAGGACAAGTTCCTGGACACCCTCTTCCTGTACGACGACGATGGGTACCAGTCCTACTGCTCCATCTGCTGCGCGGGAGAGACGCTGCTCATCTGCGAAAACCCTGATTGCACCCGATGCTACTGTTTCGAGTGTGTCGATAGCCTGGTCAGCCCGGGGACCTCGGAGAAGGTTCACGCCATGAGCAACTGGGTGTGCTTCCTGTGCCTGCCCTTCTCCCGCAGCGGGCTGCTGCAGCGGAGAAGGAGGTGGCGGGCCGGGCTGAAGGCCTTCTGCGACCGCGAGTCGGAGAGTTCCCTGGAGACGTACAAAACCGTGCCTGTGTGGAAGAGAGAGCCGGTGCGCGTGCTGTCCCTCTTCGGGAACATTAAAAGAGAGCTAACGAGTCTGGGGTTTCTGGAGGCCGGCTCTGGCCCCGGAAGGCTGAAGCACTTGGACGACGTCACGGACGTGGTGAGGAGGGATGTGGAAGACTGGGGCCCGTTTGATCTCGTGTACGGCGCGACGCCTCCCCTGGGCCACACCTGCTACCGTCCCCCCGGCTGGTACCTGTTCCAGTTCTGCCGGCTCCTGCAGTACGCGAGGCCCCCGCCGGGCAGCCCGAAGCCCTTCTTCTGGATGTTCGTGGACAATCTGGTGCTGAGCCAGGAAGACGTGGACGCGGCGGCTCGATTCCTGGAGACGGAGCCGGTGACCATCCAGGACGTCTGTGGCAGAGCCCTGCAGGACGCCGTGCGAGTGTGGAGCAACGTCCCGGCCGTGAGGAGCAGCAAGCACTCGGCCCCCGTCTCCGAAGAGGAACTGTCCCTGCTGGCTCAGGACAGGCAGAGGGCAAGGCTGCCCCCCGGGACGCCCGCCAGGCTGGTGAAGAACTGCTTTCTCCCCCTAAGAGAATATTTCAAGTATTTTTCAGCAGAGCTCACTTCCTCTTTG

>Bushbaby3L

GCAGTCACCCCCCTCCTAAGCCTGGAAGCTGAGGCCGGCATGGACGTGATCCTGGTAGGCTCCAGTGAGCTGTCGGCTCCCCCTTCACCCAAGCTGGGCAGAGATCTGATTGCATATGAAGTCAAAGTTAACCAGAGAAACATAGAAGACATCTGCCTGTGCTGTGGAAGCTTCCAGGTTCACACCCAGCACCCTCTGTTTGAGGGAGGAATGTGTACCTCGTGTAAGGACAAGTTTCTGGGCGCCCTCTTCCTGTACGATGATGACGGGTACCTGTCCTACTGCTCCATCTGCTGCTCCGGAGAGACGCTGCTCATCTGCGAAAACCCCGATTGCACCCGGTGCTACTGTTTCGAGTGTGTGGACACCCTGGTGAGCCCGGGGACTTCGGAGAAGGTTCAAGCCATGAGCAACTGGGTGTGCTTCCTGTGCCTGCCCTTCTCCCGCAGTGGGCTGCTGCGGCGCCGGCGCAAGTGGCGGGCCAGGCTGAAGGCCTTCTGTGACCGTGAGTCTGAGAGTCCCCTTGAGATGTATAAGACTGTGCCTGTGTGGAAAAGAGAGCCAGTGCGAGTGCTGTCTCTCTTTGGGGACATCAAGAGAGAGCTAACCAGTTTGGGATTCTTGGGGACCAGCTGTGGCCCAGGCAGGCTGAAGCACCTGGATGACGTCACGAACGTTGTGAGGAGGGATGTGGAGGAGTGGGGCCCCTTTGACCTCGTGTACGGCTCAACACCGCCCCTGGGCCACACCTGCTACCATCCGCCTGGCTGGTACCTGTTCCAGTTCCACCGGCTCCTGCAGTACGCACGGCCCCCACCAGGCAGCCCACGGCCCTTCTTCTGGATGTTCGTGGACAACCTAGTGCTGGGCACAGATGACCAGGCCACAGCCACTCGCTTCCTGGAGACAGAGCCTGTGACCATCCAGGACCTCAGTGGCAGAGCCCTCCAGGACGCCGTGCATGTGTGGAGCAATGTCCCTGCTGTGAGG---AGCAAGCACGCGGCTCTGGGCCCTCAGGAAGAGCTGTCCCTGCTGGTTCAGGACAGGCAGAGGATGGGGCCCTACAGCCAGAGCTCTGCCAGGCTGGTGAAGAGCTGCTTCCTCCCACTGAGAGAGTATTTCAAGTATTTTTCGACAGAACTCTCTTCCTCCTTG
